# The Copy-Number Events in Skull Base Chordoma Stratify Tumours into Four Biologically Coherent Groups

**DOI:** 10.64898/2026.03.17.712307

**Authors:** Szymon Baluszek, Paulina Kober, Renata Woroniecka, Natalia Maławska, Michał Wągrodzki, Jacek Kunicki, Tomasz Mandat, Beata Grygalewicz, Mateusz Bujko

**Affiliations:** Department of Molecular and Translational Oncology, Maria Sklodowska-Curie National Research Institute of Oncology, Warsaw, Poland; Cytogenetic Laboratory, Maria Sklodowska-Curie National Research Institute of Oncology, Warsaw, Poland; Department of Cancer Pathomorphology, Maria Sklodowska-Curie National Research Institute of Oncology, Warsaw, Poland; Department of Neurosurgery, Maria Sklodowska-Curie National Research Institute of Oncology, Warsaw, Poland

**Keywords:** skull base chordoma, copy-number classification, chr9/*CDKN2A* loss, chr7/*CDK6* gain, chr2/*E2F6* gain, sacoma cytogenetics

## Abstract

Chordoma, a rare sarcoma of notochordal origin, exhibits slow growth and local aggressiveness. While copy-number (CN) events are recognized as key chordoma drivers, no comprehensive classification, based on CN, has yet been developed. Here, we establish a robust, reproducible genomic subtyping of chordoma, based on CN events. Two independent skull base chordoma cohorts (N=32,N=71) were analyzed, utilizing distinct analytical platforms, DNA methylation microarrays and whole-genome sequencing, both controlled for B-allele frequencies. Samples were clustered using unsupervised hierarchical methods. The CN events defined four consistent molecular clusters across both cohorts: **C1** (CN-stable), **C9** (chromosomal losses, especially of chr9/*CDKN2A*), **C7** (chr7 gain), and **C2** (gains of chr2 and chr7). The findings were validated in fluorescence *in situ* hybridization (FISH) with concordance of 84-89%. The CN clusters explain 31-33% of the RNA-sequencing transcriptional variance. Moreover, the **C2** cluster showed up-regulation of Sonic Hedgehog signaling and clusters **C2** and **C9** were enriched in cell-cycle-related genes. The proposed CN clusters correlate with existing chordoma classificators e.g. chromosomal instability (CIN), mutation burden, immune score, and methylation clusters. Furthermore, comparison with over 2,000 sarcomas highlighted CN patterns more common in chordoma (i.e. chr1q, chr2, chr7 gains and chr1p, chr3, chr9, chr10, chr13, chr14, chr18 losses) but also revealed shared aberrations, e.g. chr22 loss shared with Gastrointestinal Stromal Tumours (GISTs). This study provides a unifying classification for skull base chordoma, linking distinct genomic architectures to specific transcriptional programs and potential therapeutic vulnerabilities.

## Introduction

Chordoma is a rare, slow-growing sarcoma, arising from remnants of the notochord.[35] It has a low incidence, estimated at between 0.18 and 0.84 per 1,000,000 persons per year, with a notable predilection for males and patients in their fifth and sixth decades of life.[4] These tumours develop along the axial skeleton, most commonly in the sacrum (29-45%), skull base (26-32%), and mobile spine. Despite its indolent growth, chordoma is locally aggressive and invasive, leading to significant morbidity due to its proximity to critical neurovascular structures. The prognosis is challenging with median survival of 6.3 years and 5- and 10-year survival rates of 67.6% and 39.9%, respectively.[22] This disconnect between a slow proliferative rate and a relentlessly aggressive clinical course suggests that the underlying molecular drivers of chordoma are distinct from those of many other cancers. The search for therapeutic options is hindered by a lack of driver mutation in almost half of chordoma cases.[2, 31]

One of the first examples, pointing to the importance of CN events in chordoma, was discovery of a susceptibility locus on chr6q27 in familial chordoma. It contains a duplicated *TBXT* gene, encoding brachyury, a transcription factor, intimately linked to notochord development.[42] The brachyury amplification is less common in sporadic chordoma, occurring in 27% of cases.[31] Central role of brachyury in chordoma development is well established e.g., histone 3.3 lysine 27 demethylases (KDM6A/B) inhibitors limit chordoma cell lines growth; this effect was shown to reduce *TBXT* expression and ectopic brachyury expression was protective against the effects of these demethylases knock-out.[9]

Another well-established cytogenetic event in chordoma is loss of *CDKN2A/B* locus on chr9p21. There is some controversy surrounding its prognostic effect i.e., Horbinski *et al.*[13] observed worse prognosis in chr9p loss of heterozygosity (LOH) but not in chr9p21 homozygous loss; meanwhile both Bai *et al.*[2] and Passeri *et al.*[26] observed that *CDKN2A/B* loss significantly shortened progression-free survival but not overall survival. Because *CDKN2A/B* protein products are involved in cell-cycle regulation, *CDKN2A/B* loss was an inclusion criterion in a recently concluded phase II clinical trial of palbociclib (cyclin-dependent kinase 4/6 inhibitor) in chordoma.[33] We have previously shown that *CDKN2A/B* loss is associated with decreased infiltration of immune cells and up-regulation of cell-cycle-related genes.[5] Of note, loss of *PTEN* function (including copy-number event on chr10q23) was associated with both *CDKN2A/B* loss and worse prognosis.[41]

A number of other CN events in chordoma have been described in the literature. Loss of chr1p36 was proposed to be early in chordoma oncogenesis as its loss was found in up to 90% of cases.[18, 27] Loss of short arm of chr3 was also frequent (69%) and associated with loss of histone 3.3 lysine 36 trimethylation and a better prognosis in sacral chordomas.[46] In a recent study the loss of chr14q and chr18p were associated with poor outcomes.[15] The authors did not find association of chr22 loss with overall survival, which was the case in a different cohort.[2] Although, most reported CN events in chordomas were losses, chr7 can be gained in up to 70% of these tumours, leading to up-regulation of c-MET oncogene.[37]

The abovementioned studies span 3 decades of research, where different experimental and statstical approaches were used. These inconsistencies may at least partly explain the differences in the results. To address this, we have assessed CN events in two independent cohorts of patients, utilizing different experimental and computational approaches, while controlling for single nucleotide ploymorphisms (SNPs) B-allele frequencies (BAF). The results were subsequently clustered, yielding 4 consistent groups across both experiments. The CN findings from one cohort were validated by fluorescence *in situ* hybridization (FISH) experiments at loci for *ALK*, *EGFR*, *CDKN2A*, and *PTEN*. Moreover, we have combined these results with RNA-sequencing data available for these patients, reconstructing potential gene-expression effects of the observed CN events. Finally, the results were compared with other groupings and observations in chordoma[2, 3] and more broadly, sarcoma.[23]

## Materials and Methods

### Patients and tissue samples

Samples from 32 patients with skull base chordoma, who were treated with transnasal and/or transoral endoscopic surgery at the Department of Neurosurgery, Maria Sklodowska-Curie National Research Institute of Oncology, Warsaw, in years 2014–2020 were included. This patient population was already described. Data from this study were deposited to Gene Omnibus (GSE230168).[5]

Moreover, 71 of 80 whole-genome sequencing of skull base chordoma samples, published by Bai *et al.* and available from dbGaP at phs002301, passed quality control and were utilized as a validation dataset.[2] For comparison with other sarcomas a MSK-IMPACT dataset, available from www.cbioportal.org, comprising of 2,138 samples was utilized.[23]

### DNA isolation and processing

As described previously,[5] DNA was extracted with AllPrep DNA/RNA/miRNA Universal Kit (Qiagen). The concentration of nucleotides was measured spectrophotometrically, using NanoDrop 2000 (Thermo Scientific) and with fluorescence-based method using QuantiFluor Dye kit (Promega) and Quantus (Promega) instrument. Isolated total DNA was stored at -20° C. DNA was bisulfite converted with EZ-96 DNA Methylation kit (Zymo Research) and used for genome-wide DNA methylation profiling with Methylation EPIC (Illumina) BeadChip microarrays (Infinium HD Methylation Assay Reference Guide, Illumina). Laboratory procedures were performed by the Eurofins Genomics service provider.

Moreover, the BAFs of common single nucleotide polymorphisms (SNPs) were extracted from DNA sequencing of 664 cancer-related genes using the SeqCap EZ Custom Enrichment Kit (Roche, Basel, Switzerland). A complete list of genes was described in detail previously.[40] Methods and somatic variants called have been described elsewhere[6] and somatic variants from this study are available from Zenodo (doi: 10.5281/zenodo.16751776).

### Data analysis

We have previously demonstrated that in order to properly assess CN events in patient-derived samples, one must control for BAF.[28] To this end, CN segments, generated by *conumee*[34] were examined along with BAF, extracted from panel DNA sequencing data, aligned to hg38p11 genome with *bwa*[19] for SNPs from *annovar*.[38] This led to extraction of chromosomes for which CN status was obvious and they were utilized as a reference to adjust conumee-derived DNA quantities. The WGS samples were also aligned to hg38p11 genome and facets was utilized to extract the CN events; for 7 samples the cluster of segments assigned diploid status was modified after plot examination. Virtual karyotypes for all samples are available as and the identified CN events in the Supplementary File 1 and the segments in the Supplementary File 2. Scripts, utilized for the analysis are freely available on GitHub (https://github.com/SBaluszek/CNchordoma2025). Analysis was performed in R(v4.2.2) and data were visualized in ggplot2 package.

For both groups, samples were clustered, utilizing agglomerative hierarchical algorithm (Ward D) on binary distance, assuming 6 possible CN states (normal, LOH, heterozygous loss, hemizygous loss, amplification, and at least 4-copy amplification) in each autosomal chromosomal band.[21] Next, these distances were utilizaed to construct uniform manifold approximation projection (UMAP) for dimensionality reduction, and correlation between events in cytogenetic bands was calculated, utilizing Kendall correlation test. Subsequently, patterns of CN events were compared with *COSMIC* CN signatures with cosine similarity.[32]

The global effects of CN events on gene expression were assessed with canonical-component analysis, as implemented in vegan package.[11] For each cytogenetic band, gene expression signature was extracted, utilizing *GSEAlm* package[25] and differentially expressed genes (DEGs) between clusters were identified with *DESeq2*[20] with p-adjusted threshold of 0.05. Next, for selected genes, correlation between CN events and gene expression was tested with Kendall correlation test. To assess biological processes, a global sample-level approach was applied first, running *mlm* algorithm (*decoupleR* package[1]) on gene expression with *PROGENy* signatures.[29] Next, fgsea[16] was utilized to extract significant signatures among Reactome and Gene Ontology terms (*msigdbr* package[12]). Finally, *STRINGdb* (v12.0) was utilized to extract protein-protein interactions (PPI) within sets of DEGs between clusters.[30]

### Fluorescence *in situ* hybridization

Fluorescence *in situ* hybridization analysis was performed on 19 formalin-fixed paraffin-embedded (FFPE) tumours. The FFPE specimens were prepared with a Pretreatment Reagent Kit (Wuhan Healthcare Biotechnology, Wuhan, Hubei, China) according to the manufacturer’s protocol. For the confirmation of gains of chr2 and chr7, the following probes were used: *ALK* break apart (BAP) (Vysis Abbott Molecular, Downers Grove, IL, USA) and *EGFR*/CEN7 (Zytovision, GmbH Bremerhaven, Germany). For the verification of the losses of chromosomes chr9 and chr10, the following probes were applied: *CDKN2A*/CEP9 (Vysis Abbott Molecular, Downers Grove, IL, USA) *PTEN*/CEN10 (Zytovision, GmbH Bremerhaven, Germany). The FISH results were visualized, using a fluorescence microscope, Axioskop2 (Carl Zeiss, Jena, Germany), documented by the BioView Allegro-Plus automated scanning and analysis system (Rehovot, Israel). Independent of the probe, a cut-off point was established at 20%.

### Comparison with other classifications and tumour types

The clustering results were compared with existing clusters, identified previously by our group[5] and Bai *et al.*[3] as well as average tumour ploidy, fraction of genome, affected by a CN change, and mutation burden (MB). As it came from two different dataset modalities, we have normalized mutational burden by dividing by interquartile range and subtracting the median. Raw number of mutations was also tested. For samples, where RNA-sequencing was available *ESTIMATE*[44] was utilized to assess tumour purity, immune, and stromal score. Transcriptional clusters, described by Bai *et al.* were imputed for other samples, utilizing *XGboost*.[8] Association between clustering and these features was tested with van der Vaerden test or exact Fisher test (wherever appropriate) with Dunn test and Exact Fisher test as post-hoc analysis, wherever appropriate. For comparison with other sarcomas a MSK-IMPACT dataset, available from www.cbioportal.org, comprising of 2,138 samples was utilized.[23] We did not posses BAF for this dataset but utilized homozygous deletion cases to select an appropriate cut-off. All samples were clustered as described above. Next, differences in CN frequencies between chordoma and other sarcomas were assessed with exact Fisher test. In order to explore differences and similarities within the band data, non-negative matrix factorization was applied.[10]

## Results

### Copy-number events can be used to split samples into four groups

In the group, treated in our institution, DNA relative quantities, extracted from DNA methylation arrays, utilizing *conumee* package[34] were adjusted for BAFs, extracted from a targeted gene DNA sequencing panel. Overall, recurrent losses of chr1p, chr3, chr4, chr9, chr10, chr13, chr14, chr18, chr21, and chr22 were common in these samples. Gains were most common in chr1q, chr2 and chr7. Hierarchical clustering allowed to group the events into 4 groups:

- **C1**, with relatively few CN events (5 samples);
- **C9**, characterized by loss of chr1p, chr3, chr9, chr10, chr13, chr14, chr18, chr21, and chr22 (8 samples);
- **C7** with common gain of chr7, with concomitant loss of chr1p, chromosomes chr3, chr10, chr13, chr14, and chr18 (11 samples);
- **C2** with characteristic gain of chr1q, chr2 and chr7, paired with losses of chromosomes chr9, chr13, chr14, and chr18 (8 samples).

Overall, the most common copy number events were a loss of chr1p and chromosomes chr3, chr4, chr9, chr10, chr13, chr14, and chr18 along with gain of chr1q, chr2, and chr10 (Figure 1a).

**Figure 1.**
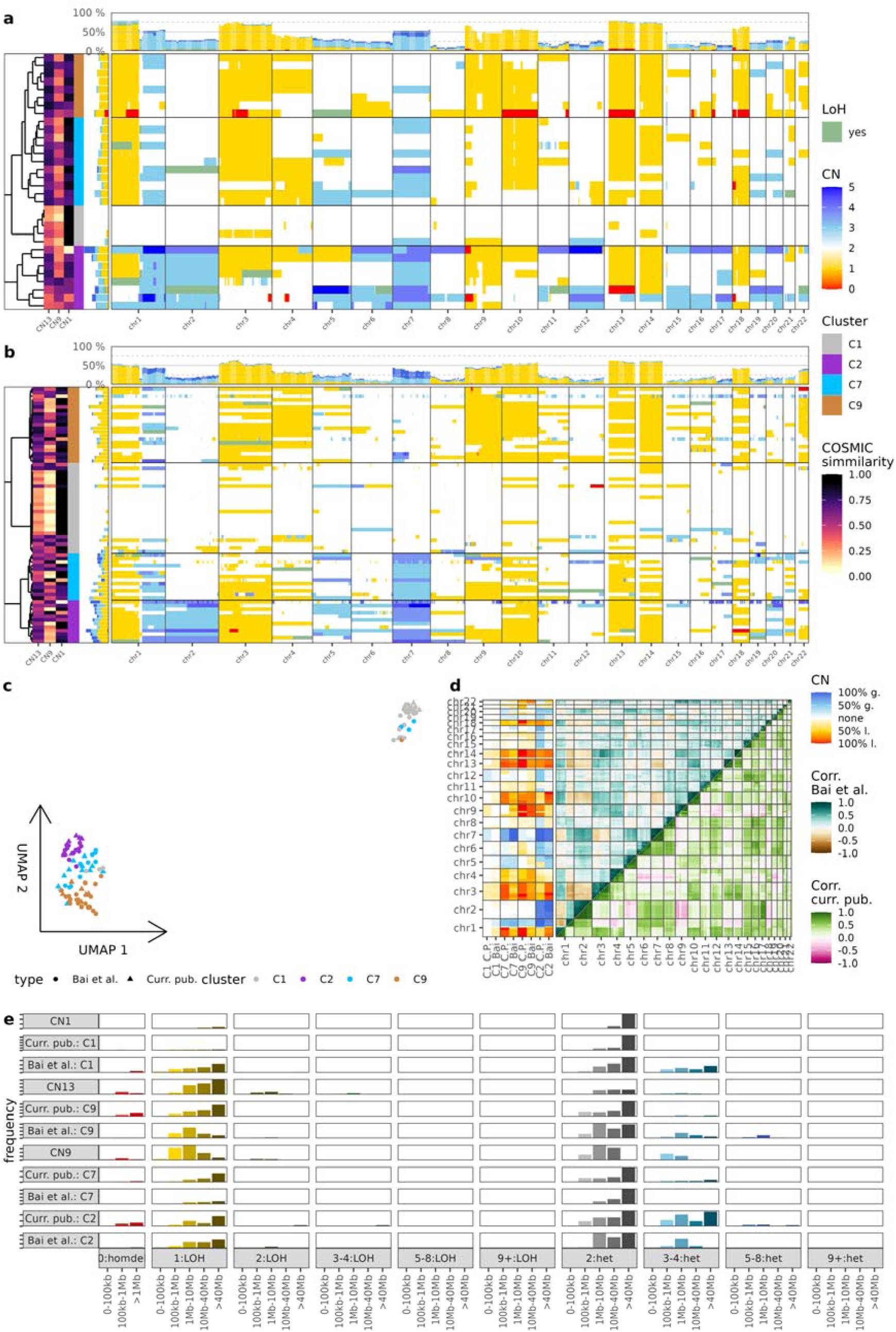
Copy-number events in human chordoma; **a)** Data from the current study, 32 skull-base chordoma samples, conumee-generated segments were corrected with panel DNA-sequencing BAFs; **b)** Data from Bai et al., 71 skull-base chordoma samples, facets-generated segments from WGS; **c)** UMAP of the samples from both sources; **d)** CN events in each cluster and their correlation with each other in both studies; **e)** comparison of CN event profiles with COSMIC signatures.

Samples from Bai *et al.* were strikingly similar in terms of CN events. The identified clusters were as follows:

- **C1**, with relatively few CN events (25 samples);
- **C9**, characterized by loss of chr1p chr3, chr4, chr9, chr10, chr13, chr14, chr18, and chr22 (21 samples);
- **C7** with relatively common gain of chr7 and chr1q, with concomitant loss of chr1p, chr3, chr10, chr13, and chr14 this cluster had less CN events overall (13 samples);
- **C2** with characteristic gain of chr1q, chr2, and chr7, paired with loss of chromosomes chr3, chr10, chr13, chr14, and chr18 (12 samples).

Overall, the most common copy number events where a loss of chr1p and chromosomes chr3, chr4, chr9, chr10, chr13, chr14, chr18, and chr22 along with gain of chr1q, chr2, and chr10 (Figure 1b). All identified CNs are available in Supplementary Material 1 and Supplementary Table 2.

The UMAP of chromosomal band CN events from both sample sources (without any integration) is shown in Figure 1c. Clearly, CN cluster and not sample origin distinguishes the observations. Nonetheless, some differences were observed. *In explicite*, chr7 gain, without chr2 gain (**C7**) was never accompanied by full chr9 gain in our data and it occurred in 5/13(38%) cases, described by Bai *et al.* Moreover, in cluster **C2** chr9 but not chr10 was frequently lost in the current study while chr10 but not chr9 was lost in the Bai *et al.* samples. Importantly, chr22 loss was a feature of **C9** cluster in the current study and of **C2** and **C9** clusters in Bai *et al.* These differences are clearly visible in Figure 1d. It also depicts correlation between CN events. Clearly, due to chromosome-wide events, bands within one chromosome were correlated with a distinction of chr1, whose short arm tends to be lost and long arm tends to be gained. Moreover, most loss and gain events tend to be correlated with each other. The exception are bands frequently gained in chordoma (chr1q, chr2, chr7) negatively correlated with chromosomes frequently lost (e.g. chr3, chr9, chr10, chr18). The general observation that gains and losses are correlated is corroborated by comparison of CN events with COSMIC signatures. Chordomas most resemble *CN1* (diploid genome), *CN13* (chromosomal LOH), and *CN9* (focal LOH - diploid and chromosomal instability); consult Figure 1e.

### Fluorescence *in situ* hybridization experiments corroborate the findings

For all the samples, where state of the FFPE slides allowed for FISH (19), probes for *ALK* (chr2), *EGFR* (chr7), *CDKN2A* (chr9), and *PTEN* (chr10) were hybridized with the samples. These genes of interest were selected due to probe availability and proven track record. The FISH result was considered concordant with the computational results if at least 20% of cells displayed the same abnormality. This was achieved in 16 (84%) samples in the case of *ALK* and 17 (89%) samples for the remaining probes. Figure 2 shows representative samples for each cytogenetic abnormality, along with comparison between CN clusters, computed abnormalities, and FISH results.

**Figure 2.**
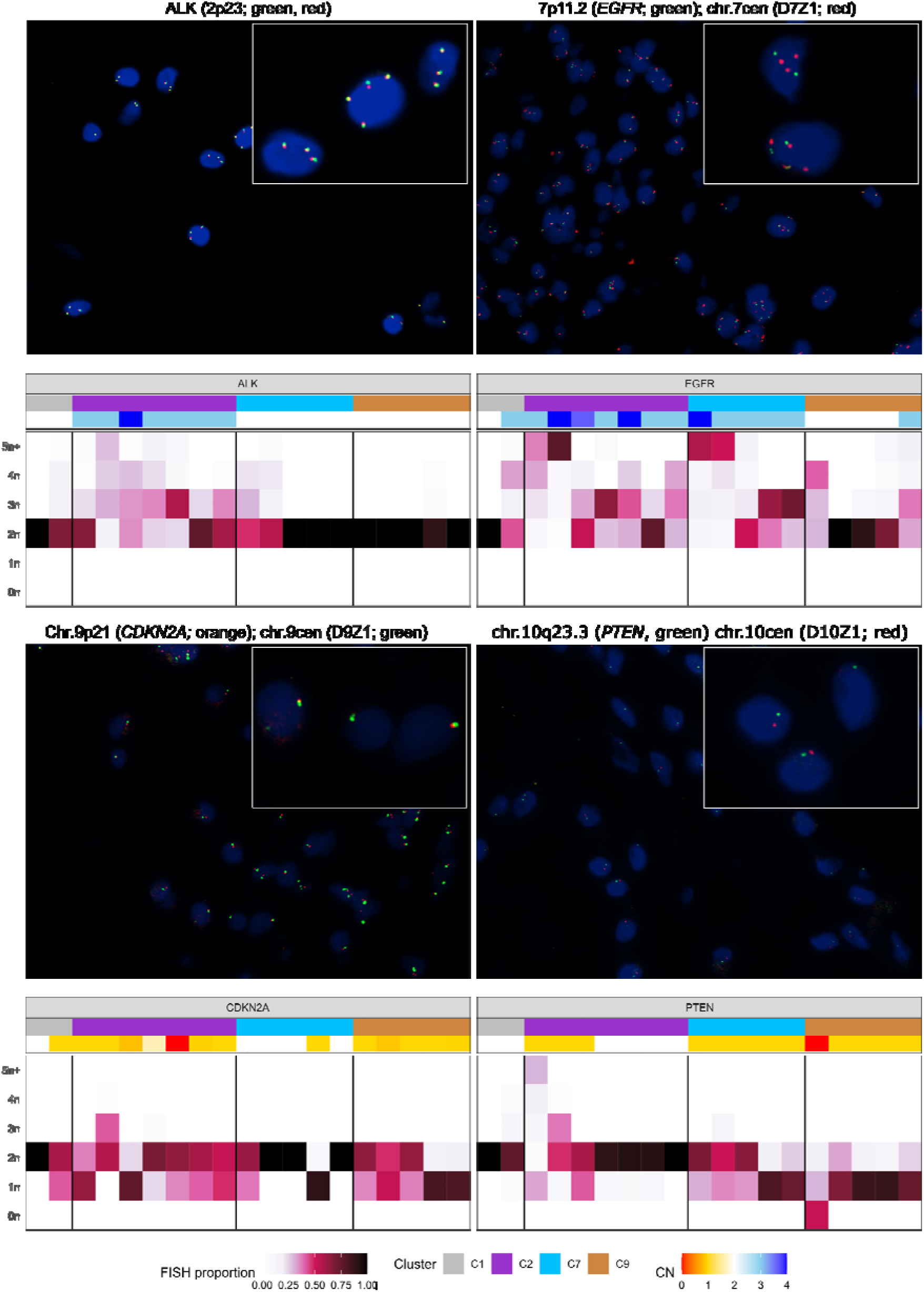
Fluorescence in situ hybridization on available subset of 19 formalin-fixed paraffin-embedded chordoma samples. Experiments were performed with probes specific for chr.2p23 (ALK yellow), chr.7p11.2 (EGFR green, D7Z1 red), chr.9p21 (CDKN2A red, D9Z1 green), and chr10q23.3 (PTEN green, D10Z1 red). For each probe, top panels shows representative sample hybridization, bottom depict fraction of cells assigned to CN events for each probe.

### Chordoma copy-number clusters shape gene expression

To demonstrate effect of CN events on chordoma gene expression, RNA-sequencing data, accompanying both studies were analyzed. Constrained component analysis was performed to assess global effects of CN changes; this method allows for measuring the variance in the constrained dimensions and the remaining, unexplained variance in the data. The CN clusters, along with chr2, chr7 gains and chr9, chr10 losses were used as the constraining variables while VST-normalized RNA-seq counts were used for the dimensionality reduction matrix. As a result CN events explained 31.26% and 33.45% of variance in the current publication and Bai *et al.* data, respectively. The resultant biplots are shown in Figure 3a and Supplementary Figure 1a, respectively. They both show relatively good separation of samples from each cluster, based on gene-expression data.

**Figure 3.**
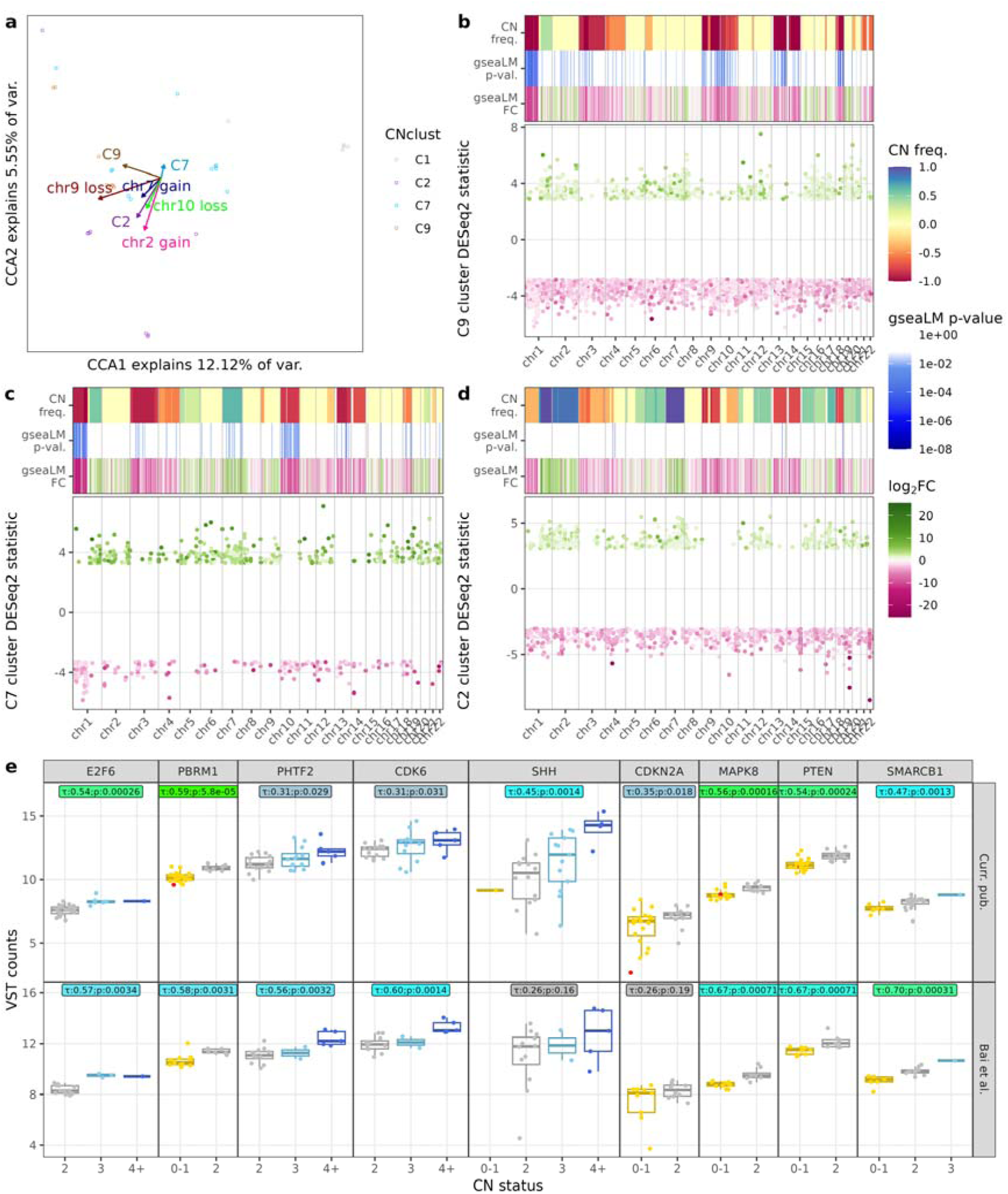
Copy-number events in the context of RNA-sequencing data; **a)** canonical componenent analysis of samples from the current study; **b)** gene expression and CN events in **C9**from the current study; **c)** gene expression and CN events in **C7**from the current study; **d)** gene expression and CN events in **C2**from the current study; **e)** gene expression correlation with CN events for selected genes;

Next, establishing local effects of CN events on gene expression was undertaken; DEGs between **C1** (CN-stable tumours) and each cluster were established[20] and *GSEAlm*[25] was used to calculate signature of gene expression for each chromosomal band. In the **C9** cluster, genes from chr1p, chr3, chr9, chr10, chr11, chr13, chr14, and chr18 had an overall lower expression with over-representation of downregulated DEGs (Figure 3b). Meanwhile, in the **C7** cluster, lowered expression on bands in chr1p, chr10, chr13, and chr18 was observed with higher expression of genes from bands on chr7. Note corresponding over-representation of DEGs (upregulated genes on chr7 and downregulated on chr1p, chr10, chr13, and chr18, Figure 3c). For the **C2** cluster few bands had significantly different GSEAlm-normalized expression but clear over-representation of genes from chr3, chr9, chr10, chr13, and chr14 among the upregulated DEGs is striking (Figure 3d). All of these observations were validated in the dataset from Bai *et al.* (Supplementary Figure 1b-d). The general tendencies are similar, although with fewer significant differences, probably due to lower number of samples.

Moreover, CN events effect on selected genes of interest, was tested (Figure 3e). First, *E2F6* encodes a transcription factor, involved in cell-cycle regulation that forms multimeric complexes capable of binding to *MYC* and brachyury binding sites.[24] It is located on chr2 and its expression is correlated with gain of its *locus*. Another gene of interest is *PBRM1*, already described in chordoma context, whose expression was significantly lowered by the loss on chr3.[2] A correlation of expression with chr7 gain was observed for *PHTF2* (a putative homeobox domain) and *CDK6* (target for a recently finished clincal trial in chordoma). We have also seen a correlation of *SHH* expression with CN event on chr7 in our data. Likewise, in the current study but not these by Bai *et al.* a correlation was observed for *CDKN2A* expression and locus CN status. Two genes of interest, located on chr10, whose expression was correlated with CN event, were *MAPK8* and *PTEN*. Finally, *SMARC1B*, whose biallelic loss is a hallmark of poorly-differentiated chordoma,[36] had expression correlating with CN status in both datasets. Of note, we did not observe strong correlation of CN events at the brachyury locus with its expression. Another gene of interest in chordoma is *EGFR* as it is over-expressed in this tumour and was implicated in mesenchymal to epithelial transition. Interestingly, its expression was not correlated with locus CN status in our data and only weak correlation was seen in data from Bai *et al.* (Supplementary Figure 1e).

### Functional consequences of copy-number events

At the next step more general relationships between CN clusters and tumour biology were inferred from genes expression profile. First, *PROGENy[1, 25]* top 500 gene scores were utilized to build multivariate linear model (MLM) in the *DecoupleR* package.[1, 30] The resultant scores for samples are shown in Figure 4a. Two pathways had scores signifficantly different between clusters in both datasets - VEGF and MAPK. The *post-hoc* analysis indicated that their signaling was lower in **C1** cluster than in **C2** (both datasets). We have observed more functional effects in our dataset (e.g. TRAIL and NFKB signaling lower in **C1** cluster than **C9**) that are consistent with the previously made observations that chr9 loss is associated with lower immune cells infiltration.

**Figure 4.**
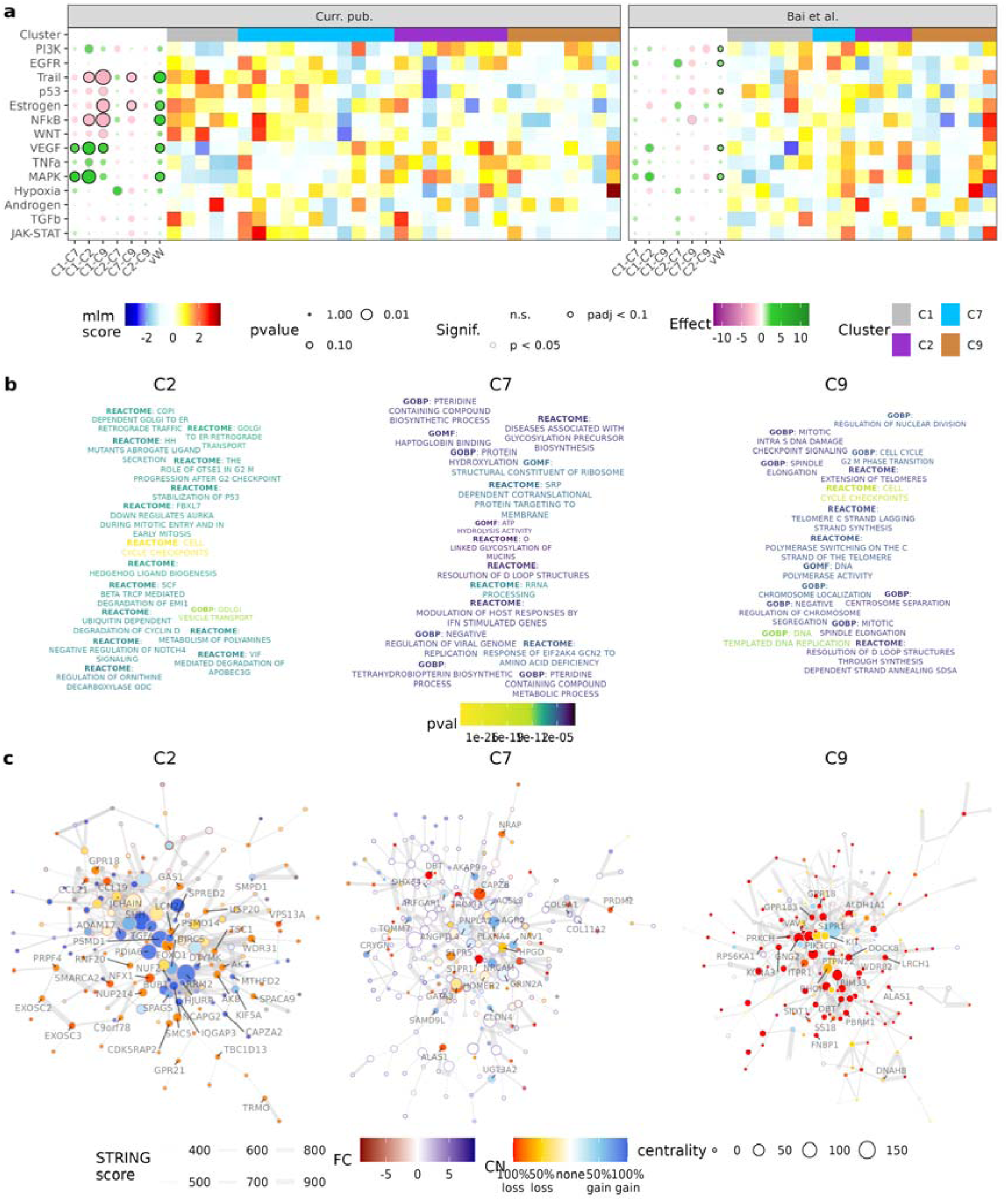
Functional gene analysis; **a)** Enrichment (enricheR mlm) on the Progeny gene activity scores from RNA-sequenced samples; **b)** Reactome and Gene Ontology enrichment (fgsea), top terms are shown as a word-cloud; **c)** STRINGdb-derived net of DEGs in each cluster.

Second, differences between clusters were enriched in terms from Reactome, Gene Ontology: Molecular Function, and Gene Ontology: Biological Process. In general, **C2** cluster showed tendency to express genes, associated with sonic hedgehog protein, regulation of cell-cycle and apoptosis. In the **C7** cluster terms, associated with gene translation and mRNA metabolisms tended to be over-expressed. Meanwhile, in the **C9** cluster, genes associated with cell-cycle and DNA polymerase activity were over-expressed. The most significant and specific terms are shown in the word-clouds in the Figure 4b. Full results of the functional analyses are available in Supplementary Table 3.

Finally, proteins, encoded by genes over-expressed in each cluster were enriched in interactions between them, revealing centrality of each protein (node). The resultant PPI-networks are shown in Figure 4c. Note that SHH and RRM2 are central in the **C2** network and located on chr7 and chr2, respectively. They are involved in sonic hedgehog signaling and deoxyribonucleotide synthesis. This is in line with finding that SHH pathway and cell-cycle are affected in this cluster. In **C7** cluster a sparser network revealed proteins, involved in cell-cell adhesion (e.g. NRCAM, collagens) and lipid metabolism (e.g. S1PR1, S1PR5, ANGPTL4). Meanwhile the **C9** network was dominated by genes, associated with G-protein signaling (e.g. DOCK8, PIK3CD, GNG2) and SWI/SNF chromatin remodelling complex (e.g. SS18, PBRM1). There were very few over-expressed genes here, probably reflecting a metabolism oriented towards cell division rather than mRNA synthesis.

### Contextualization of the copy-number clusters

Several molecular classifications of chordoma has been previously described. For our samples, DNA methylation-based classifier[5] was readily available and metrics commonly assessed in chordoma were included in the analysis of all 103 samples. The CN clusters were significantly associated with ploidy, understood as *gains net losses*; it was higher in **C1** and **C2** clusters than in **C7** and **C9** (for both datasets). Fraction of genome affected by CN was lowest in **C1** cluster, higher in **C9,** and yet higher in **C2** and **C7** clusters (again for both datasets). The absolute MB was higher for tumours from **C7** cluster than in **C1** cluster (both datasets). The DNA methylation clusters were also associated with CN clusters, although *post-hoc* analysis did not yield significant results. See Figure 5a,b,e for the results.

**Figure 5.**
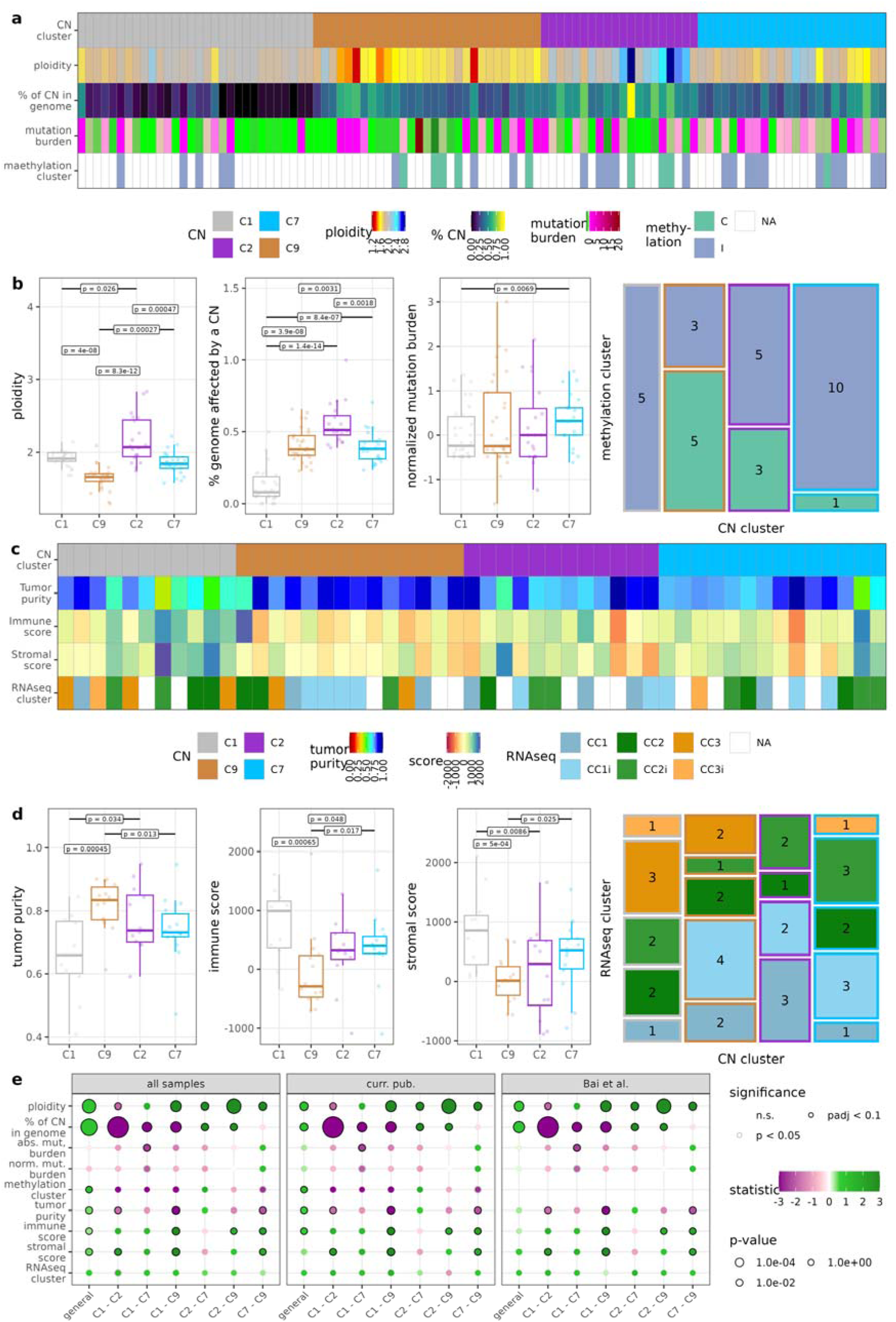
Cross-examination of the identified clusters with other features; **a)** CN burden, ploidy, MB, and DNA methylation cluster for all the samples (32 + 71 = 103); **b)** association of these features with CN clusters; **c)** tumour purity, ESTIMATE immune and stromal score, and RNA-sequencing cluster for samples with available RNA-sequencing (32 + 19 = 51); **d)** association of the RNA-sequencing features with CN clusters; **e)** statistical significance of the associations.

Moreover, for 51 samples, paired RNA-sequencing data was available. ESTIMATE, a computational tool for estimation of tumour infiltration by non-cancerous cell types was applied. A far lower tumour purity was seen in **C1** chordoma (in both datasets). This can be explained by high immune and stromal score in this cluster observed in our dataset. The immune score was also higher there in the **C2** and **C7** clusters than in **C9** cluster. We did not observe a significant association of RNA-sequencing-based clustering, described by Bai *et al.*[3] and CN clusters even after an imputation of this classification was performed on our data. See Figure 5c,d,e for the results.

### Chordoma copy-number events in a broader context of sarcomas

Despite abundance of data, no comparison of chordoma with other sarcomas in terms of CN events has yet been made. Here, data from MSK-IMPACT study were accessed (2,138 samples)[23] and combined with 103 chordoma samples, described above. Next samples were clustered and the result, along with CN event frequency can be seen in Figure 6a. It can be easily observed that some CN events common in sarcoma *en masse* are also common in chordoma (i.e. chr1p, chr3, chr4, chr9, chr10, chr13, chr14, chr18, and chr22 losses as well as chr7 gain). However, some CN events are significantly more common in chordoma - losses of chr1p, chr3, chr4, chr9, chr10, chr13, chr14, chr18, and chr21 as well as gains of chr1q, chr2 and chr7. Notably, losses of chr1q, chr2, chr7, chr17 as well as gains of chr3, chr4, chr8, and chr18 are significantly less common in chordoma, when compared to other sarcomas (see Figure 6b).

**Figure 6.**
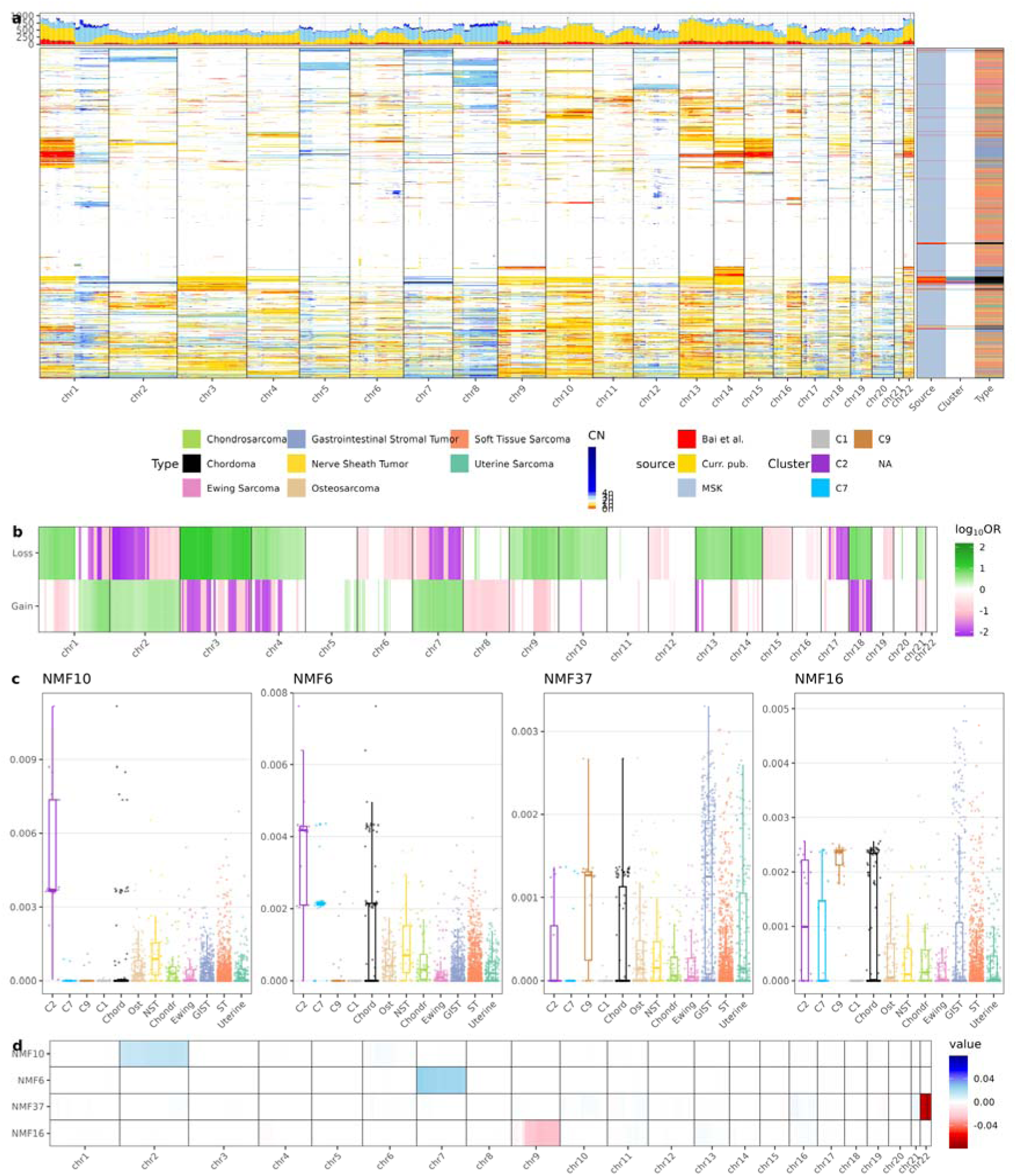
Integration of chordoma samples into the MSK-IMPACT data; **a)** CN events in the combined datasets; **b)** Significant differences between chordomas and other sarcomas; **c)** Non-negative matrix factorization selected themes activity in samples; **d)** Non-negative matrix factorization selected themes activity in chromosomal bands.

A method frequently employed to search for common topics in a large corpus of sparse data is NMF.[10] It decomposes a table of non-negative values into two matrices (of samples and features), each with activity of a specific topic. After cross-validation 42 dimensions were selected for the CN dataset of MSK-IMPACT and chordomas combined. Several factors (topics) were significantly different between chordoma and other samples. Notable factors were NMF10, associated with cluster **C2**, NMF6; associated with clusters **C2** and **C7**; NMF37, associated with cluster **C9**; and NMF37, associated with clusters **C2** and **C9** (see Figure 6c). Notably, NMF37 was also high in GISTs. Association of factors with CN events is shown in Figure 6d. Clearly, factors NMF10, NMF6, NMF37, and NMF16 are associated with chr2 gain, chr7 gain, chr22 loss, and chr9q loss respectively. The observation of common theme (loss of chr22) helps to explain why some chordomas cluster together with GIST (Figure 5a); loss of chr22 has been already described as a feature of GIST.[17]

## Discussion

This study presents a comprehensive analysis of CN alterations in skull base chordoma, utilizing two independent patient cohorts and distinct analytical platforms to establish a robust and reproducible genomic classification. By the analysis of genetic data of 32 tumour samples from patients treated in our center four distinct and consistent CN-based clusters were identified — **C1** (CN-stable), **C9** (characterized by widespread chromosomal losses, including chr9), **C7** (defined by chr7 gain), and **C2** (marked by gains of chr2 and chr7 and various chromosomal losses). They not only unify previously disparate cytogenetic observations but also correlate with specific, actionable biological pathways. The **C1** cluster, characterized by relatively few CN events, likely represents an early genomic state, while the progression through **C9** or **C7** clusters both lead to the **C2** cluster that is associated with a characteristic chr2 gain that is not observed in the other clusters (states). It may therefore serve as a proxy for high chromosomal instability in chordoma.

The selected recurrent CN events were examined by FISH, to verify the identified CN clusters. The 84-89% concordance achieved between computational CN calls and FISH validation demonstrates the reliability of our analytical approach. Our analysis across two independent cohorts using different experimental platforms (methylation arrays with targeted sequencing vs. whole-genome sequencing) strengthens the generalizability of our findings. The consistency of clustering patterns, despite different technical approaches, suggests that the identified CN signatures represent genuine biological phenomena rather than technical artefacts.

Recent studies have begun to establish molecular subtyping systems for chordoma using various approaches. RNA sequencing-based studies have identified molecular subtypes associated with chromatin remodeling defects,[2, 39] DNA methylation analysis has revealed distinct epigenetic clusters, associated with immune cells infiltration,[5, 14, 47] and proteomic analysis has identified three molecular subtypes based on bone microenvironment, mesenchymal, and epithelial transition patterns.[43] Our CN-based classification provides a complementary and potentially orthogonal molecular stratification system. First, Zhang *et al.* observed that chordomas with high CIN score (many CN events) lead to worse overall and progression-free survival.[45] The four clusters identified in this work propose a more granular classification with **C1** samples having fewest events and **C2** having the most events and the remaining two clusters being two alternative intermediate states. Second, a few studies have described clusters that were highly infiltrated by immune cells.[5, 14, 47] We have seen much higher immune infiltration and stromal score in **C1** than any other cluster. The relation of CN clusters with methylation clusters is less straight forward, however. The chordoma C was characterized by a malignant pattern of global DNA methylation and signatures of rapid cell cycling.[5] These features, while not present in **C1** cluster, were shared by a varying fraction of other cluster samples.

Our identification of recurrent chromosomal losses involving chr1p, chr3, chr4, chr9, chr10, chr13, chr14, chr18, and chr22, along with gains of chr1q, and chr7, aligns remarkably well with established literature. Le *et al.*[18] previously reported similar patterns of large copy number losses in sporadic chordomas and our findings corroborate the frequent loss of chr1p36 described in up to 90% of cases by Riva *et al.*[27] The prominence of chr7 gain, observed in our **C2** and **C7** clusters, is consistent with previous reports showing chr7 gain in up to 70% of chordomas, leading to c-MET up-regulation.[37] The frequent loss of *CDKN2A/B* locus, that has previously been associated with shortened progression-free survival in chordoma is a back-bone of our classification.[3, 26]

Gain of chr2 and existence of **C2** cluster were not previously described. Clearly, chr2 gains were present in previous works.[2, 15, 45] It almost always occurs in a context of chr7 gain (7/7 and 11/12 in data from current publication and Bai *et al.*, respectively). It is less clear, how it relates to chr9 and chr10 losses - chr2 gain was accompanied by chr9 loss in 6/7 and 3/12 and by chr10 loss in 2/7 and 8/12 in data from current publication and Bai *et al.*, respectively. It had clear transcriptomic consequences - activation of MAP kinase pathway, probably due to SHH activation, leading to rapid cell cycling. The upregulation of Sonic Hedgehog signaling in the **C2** cluster is consistent with the known role of this pathway in chordoma pathogenesis.[7] The correlation between *E2F6* expression and chr2 gain is particularly intriguing, as E2F6 can bind to both MYC and brachyury binding sites, potentially linking chromosomal gain to transcriptional dysregulation of key chordoma drivers. Notably, **C2** cluster had higher ESTIMATE immune score than **C9** cluster, further supporting its separate identity.

Our constrained component analysis revealed that CN events explain 31-33% of transcriptional variance, indicating that chromosomal alterations are major drivers of chordoma gene expression programs. This finding is particularly significant given that nearly half of chordomas lack identifiable driver mutations, supporting the thesis that CN alterations may serve as primary pathogenic mechanisms in this malignancy.[2, 31] As previously described, the enrichment of cell cycle-related genes in clusters with CDKN2A/B loss (**C2** and **C9**) provides mechanistic insight into how these CN events drive tumour progression.[5] This was posited to lead to susceptibility to CDK4/6 inhibitors such as palbociclib, which showed modest antitumour activity in a recently completed phase II trial.[33] Notably, *CDK6* lies on chr7 and one of four responders after 6 cycles in this trial harboured chr7 gain. Role of chr7 gain in predicting response to CDK4/6 inhibitors remains to be seen.

The comparison of chordoma CN patterns with other sarcomas from the MSK-IMPACT cohort reveals both shared and distinct features. While some CN events are common across sarcomas, chordomas show significantly higher frequencies of losses involving chromosomes chr1p, chr3, chr4, chr9, chr10, chr13, chr14, chr18, and chr21, as well as gains of chromosomes chr1q, chr2, and chr7. This pattern suggests that chordomas may have unique selective pressures or developmental origins that predispose to specific chromosomal instabilities. The observation that some chordomas cluster with GIST based on chr22 loss is particularly interesting as chr22 loss associated with shortened overall survival in chordoma[2] but also to a malignant phenotype in GIST.[17]

Several limitations should be acknowledged. First, our study focused exclusively on skull-base chordomas, and extrapolation to sacral, mobile spine, or extra-axial chordomas requires validation. Second, while we identified transcriptional consequences of CN events, the functional impact of these alterations on protein expression and cellular behavior requires further investigation. Third, the prognostic value of the CN clusters needs to be examined in larger cohorts. Future studies should investigate how our CN-based classification correlates with clinical outcomes, treatment response, overall and progression-free survival.

## Conclusions

This study provides the most comprehensive analysis to date of CN events in chordoma, identifying four reproducible molecular clusters with distinct genomic and transcriptional characteristics. Our findings demonstrate that chromosomal alterations are major drivers of chordoma biology, explaining nearly half of transcriptional variance and providing a framework for understanding the molecular heterogeneity of this rare malignancy. The consistency of our results across independent cohorts and validation by FISH analysis supports the clinical relevance of this molecular classification system. These insights provide a foundation for future precision medicine approaches in chordoma and highlight potential therapeutic vulnerabilities that warrant further investigation.

## Supporting information

GSEA results

Copy-number segmentation, visualization of raw results

Copy-number segmentation, segments

## Supplementary Material

Supplementary File 1

***SupplementaryFile1.pdf***, contains segments and BAFs for all samples.

Supplementary Table 2

***SupplementaryTable2.xlsx***, contains:

· *SegmentsCurrentPub* tab with CN events from the current publication;

· *SegmentsBai* tab with CN events from Bai *et al*.

Supplementary Table 3

***SupplementaryTable3.xlsx***, contains:

· *ProgenyCurrentPub* tab with Progeny scores for the current publication;

· *ProgenyBai* tab with Progeny scores forBai *et al.*;

· *fgseaCurrentPub* tab with fgsea results for the current publication;

· *fgseaBai* tab with fgsea results for Bai *et al*.

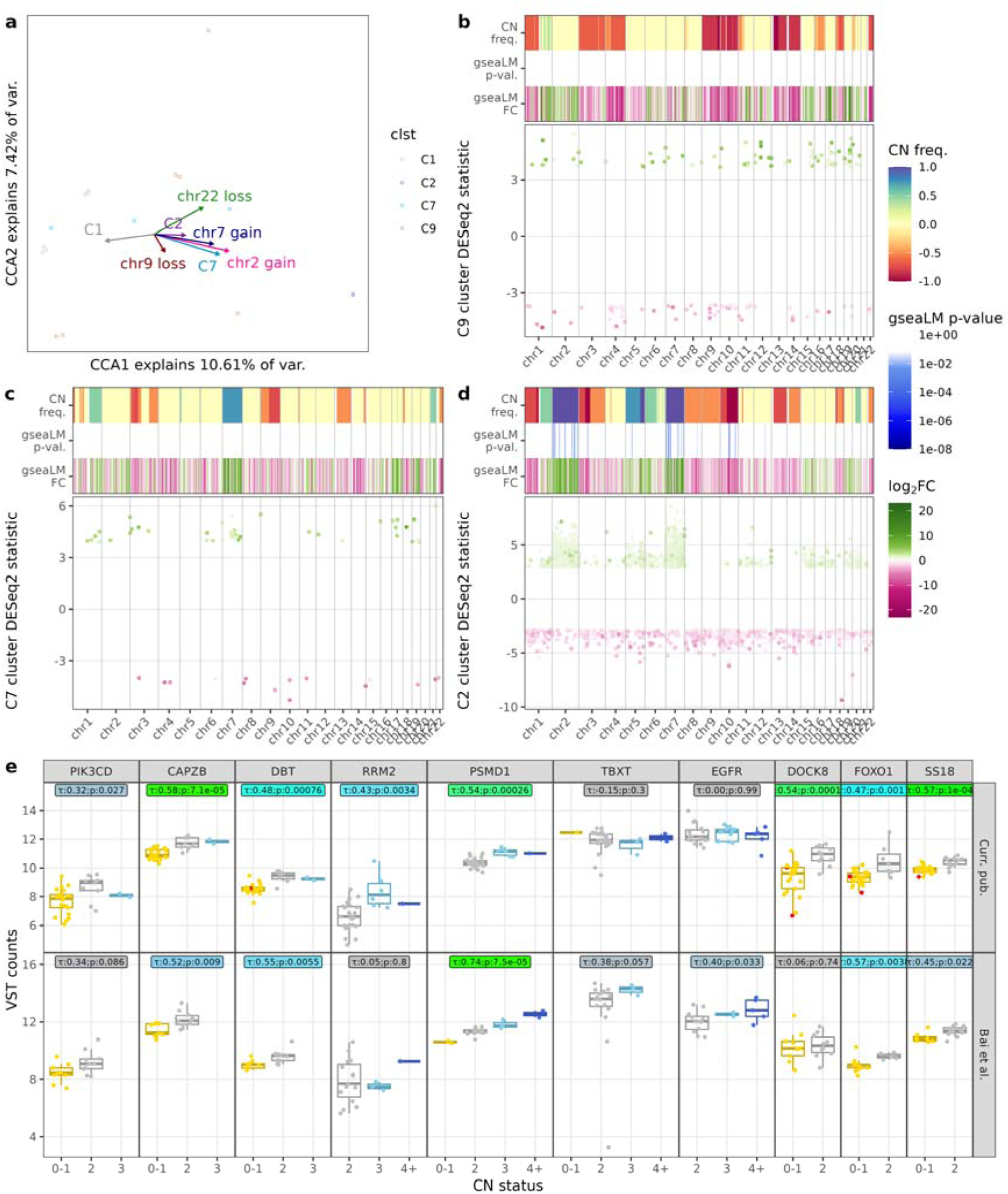
Copy-number events in the context of RNA-sequencing data; a) canonical componenent analysis of samples from Bai et al.; b) gene expression and CN events in **C9** from Bai et al.; c) gene expression and CN events in **C7** from Bai et al.; d) gene expression and CN events in **C2** from Bai et al.; e) gene expression correlation with CN events for selected genes.

