## Supplementary material for "The Copy-Number Events in Skull Base Chordoma Stratify Tumours into Four Biologically Coherent Groups": Copy-number segmentation, visualization of raw results

11/18

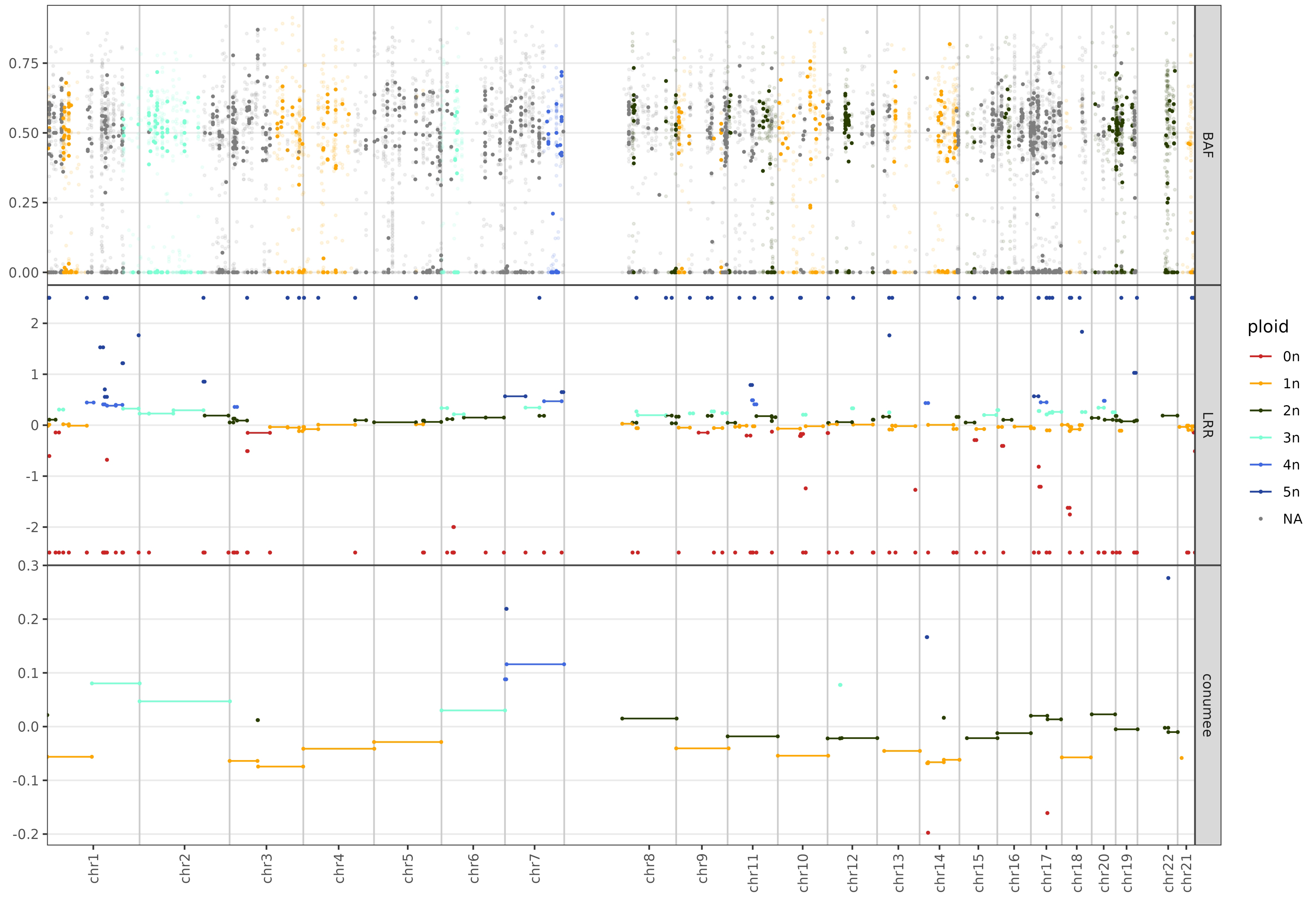
BAF
LRR
conumee
ploid
0n
1n
2n
3n
4n
5n
NA
chr1
chr2
chr3
chr4
chr5
chr6
chr7
chr8
chr9
chr11
chr10
chr12
chr13
chr14
chr15
chr16
chr17
chr18
chr20
chr19
chr22
chr21

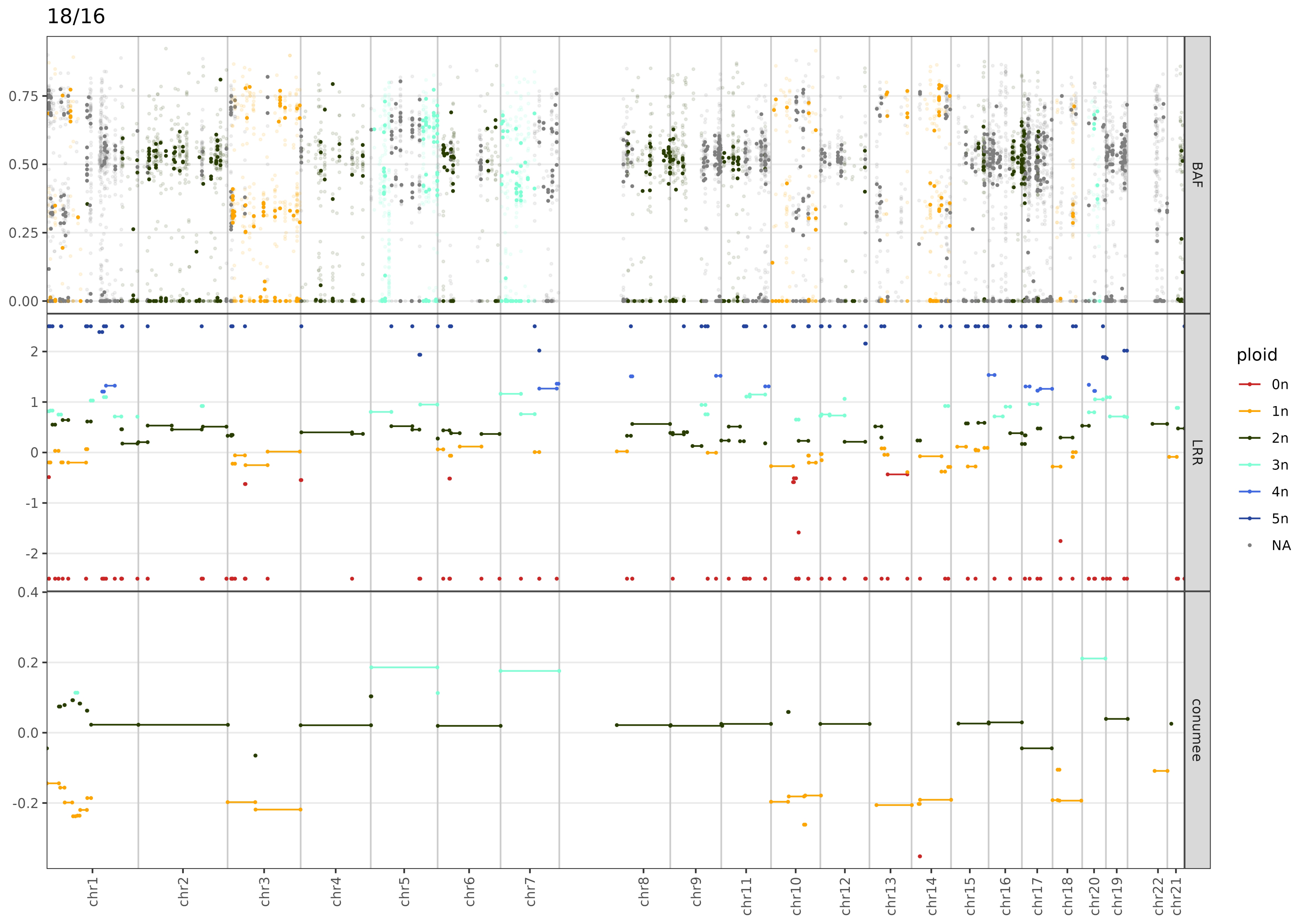

18/16
0.75
0.50
0.25
0.00
BAF
2
1
0
-1
-2
LRR
0.4
0.2
0.0
-0.2
conumee
chr1
chr2
chr3
chr4
chr5
chr6
chr7
chr8
chr9
chr11
chr10
chr12
chr13
chr14
chr15
chr16
chr17
chr18
chr20
chr19
chr22
chr21
ploid
0n
1n
2n
3n
4n
5n
NA

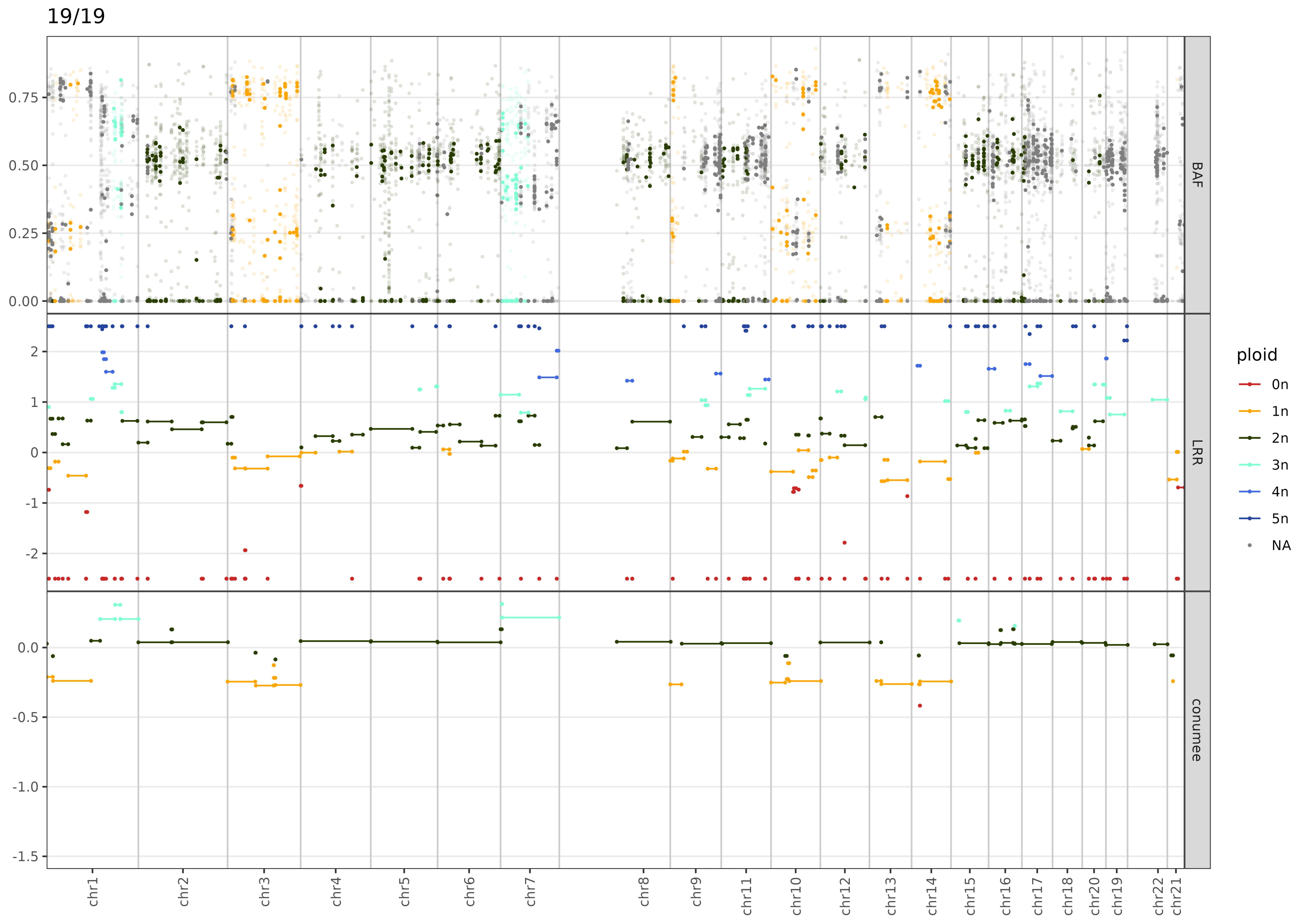

19/19
BAF
LRR
conumee
ploid
0n
1n
2n
3n
4n
5n
NA
0.75
0.50
0.25
0.00
2
1
0
-1
-2
0.0
-0.5
-1.0
-1.5
chr1
chr2
chr3
chr4
chr5
chr6
chr7
chr8
chr9
chr11
chr10
chr12
chr13
chr14
chr15
chr16
chr17
chr18
chr20
chr19
chr22
chr21

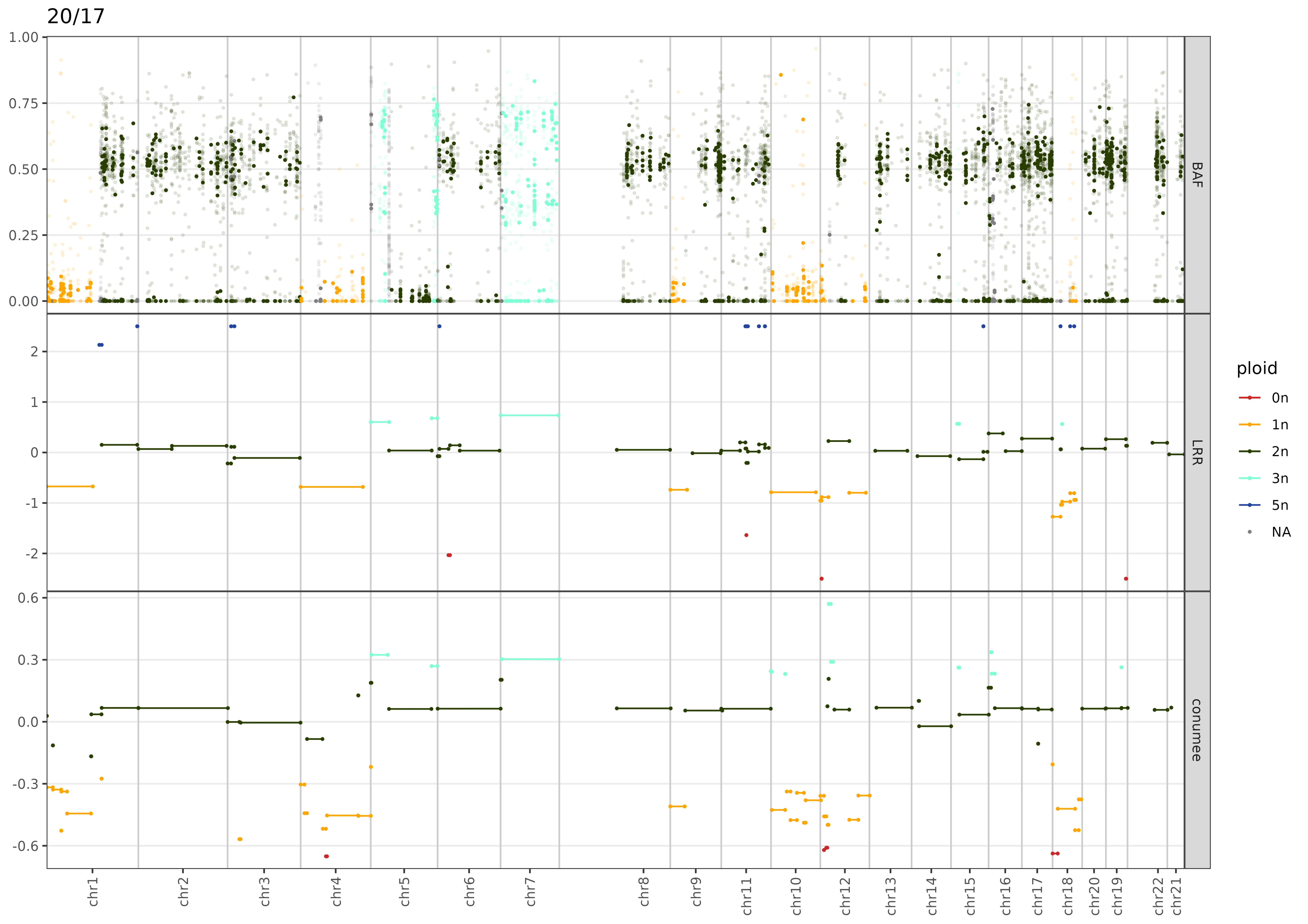
20/17
BAF
LRR
conumee
ploid
0n
1n
2n
3n
5n
NA
chr1
chr2
chr3
chr4
chr5
chr6
chr7
chr8
chr9
chr11
chr10
chr12
chr13
chr14
chr15
chr16
chr17
chr18
chr20
chr19
chr22
chr21

24/18

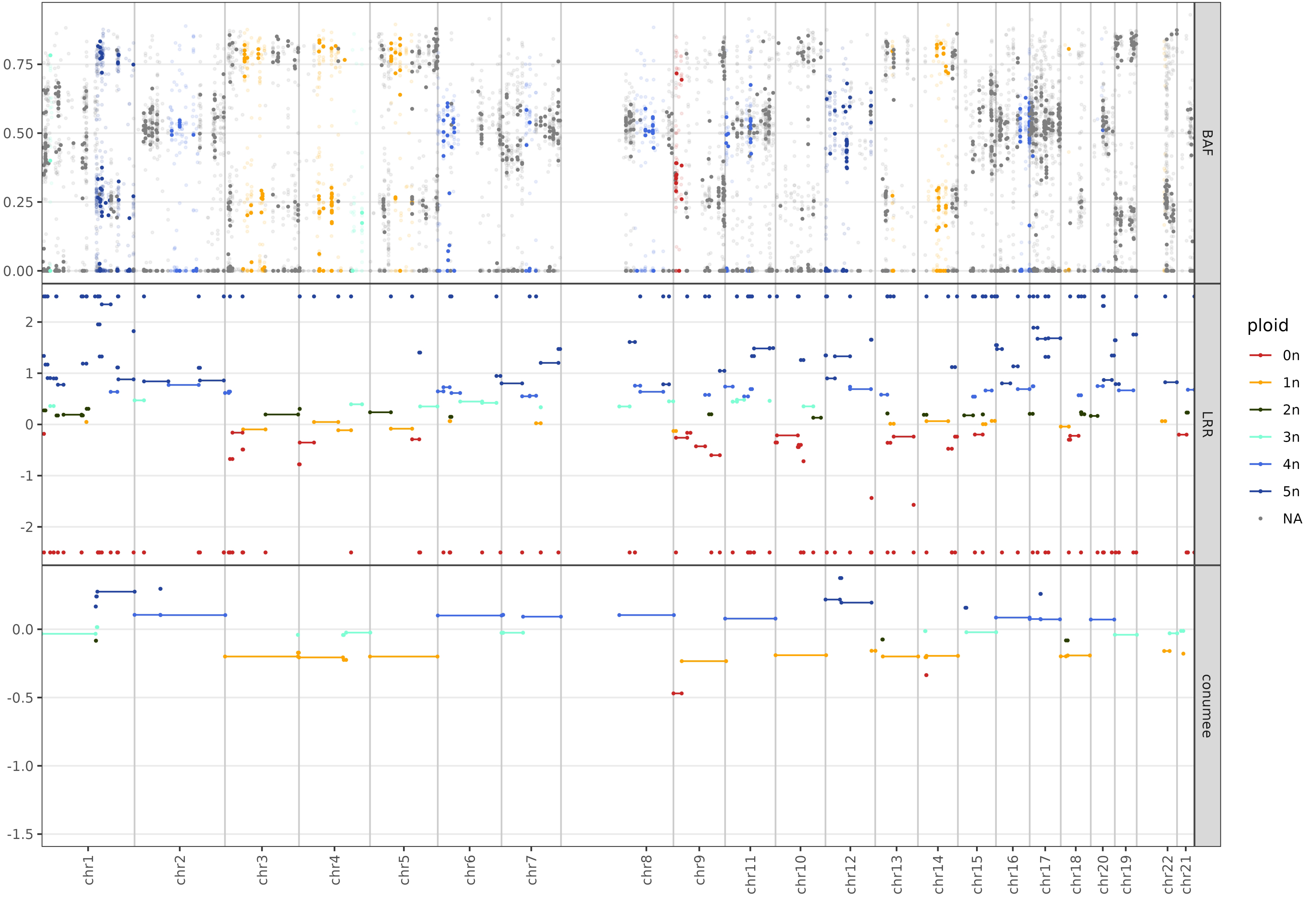

BAF
LRR
conumee
ploid
0n
1n
2n
3n
4n
5n
NA
0.75
0.50
0.25
0.00
2
1
0
-1
-2
0.0
-0.5
-1.0
-1.5
chr1
chr2
chr3
chr4
chr5
chr6
chr7
chr8
chr9
chr11
chr10
chr12
chr13
chr14
chr15
chr16
chr17
chr18
chr20
chr19
chr22
chr21

26/15

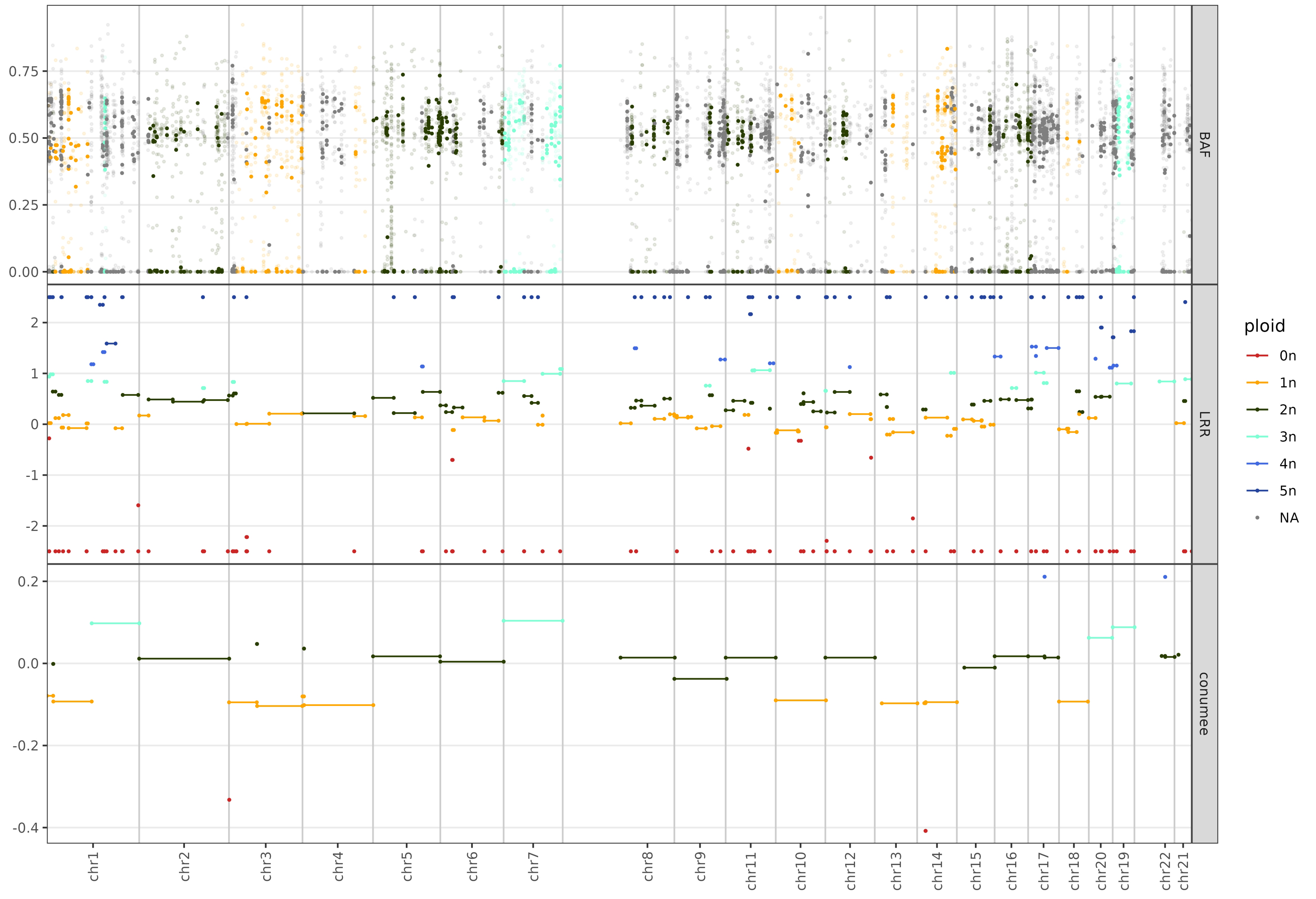

BAF
LRR
conumee
ploid
0n
1n
2n
3n
4n
5n
NA
0.75
0.50
0.25
0.00
2
1
0
-1
-2
0.2
0.0
-0.2
-0.4
chr1
chr2
chr3
chr4
chr5
chr6
chr7
chr8
chr9
chr11
chr10
chr12
chr13
chr14
chr15
chr16
chr17
chr18
chr20
chr19
chr22
chr21

26/16

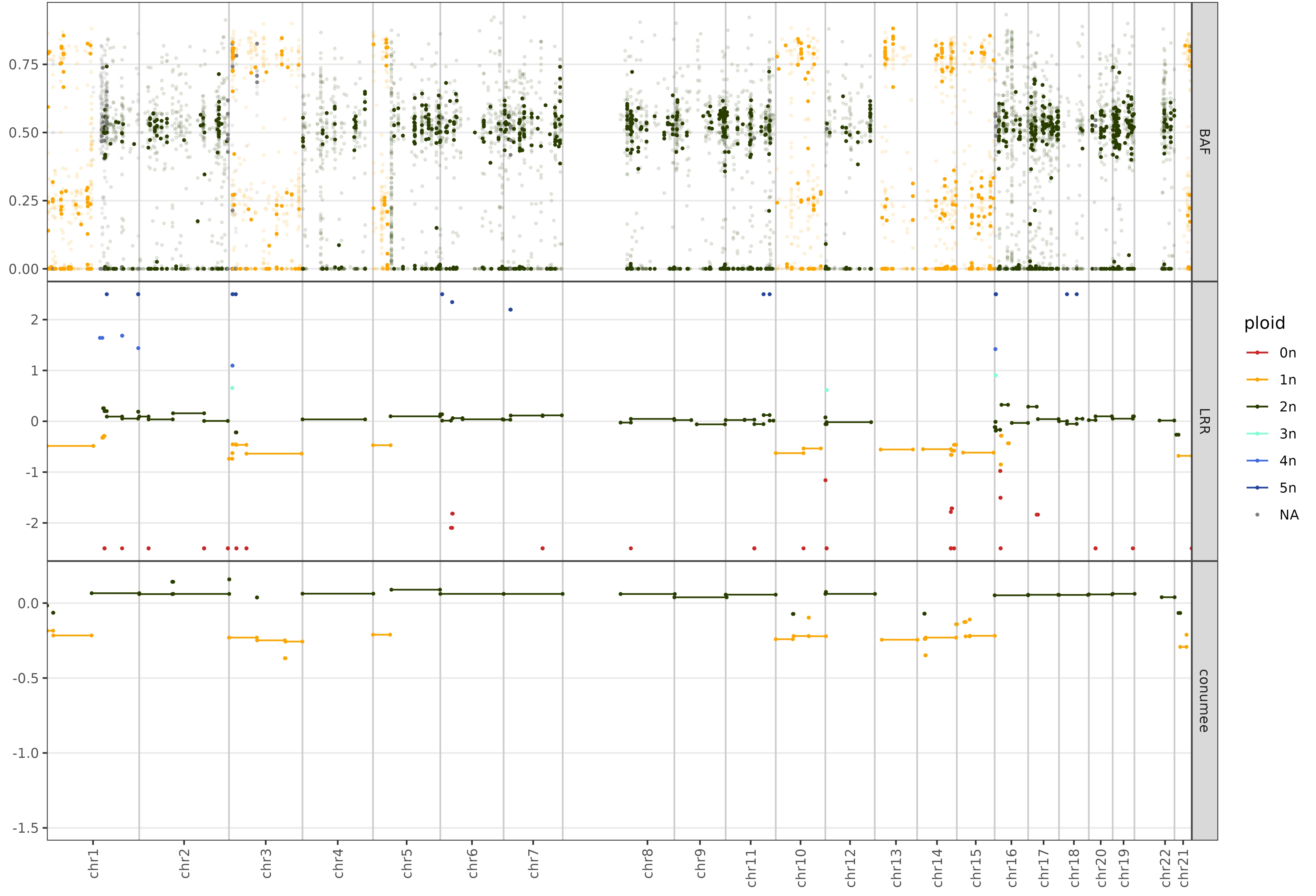

BAF
LRR
conumee
0.75
0.50
0.25
0.00
2
1
0
-1
-2
0.0
-0.5
-1.0
-1.5
ploid
0n
1n
2n
3n
4n
5n
NA
chr1
chr2
chr3
chr4
chr5
chr6
chr7
chr8
chr9
chr11
chr10
chr12
chr13
chr14
chr15
chr16
chr17
chr18
chr20
chr19
chr22
chr21

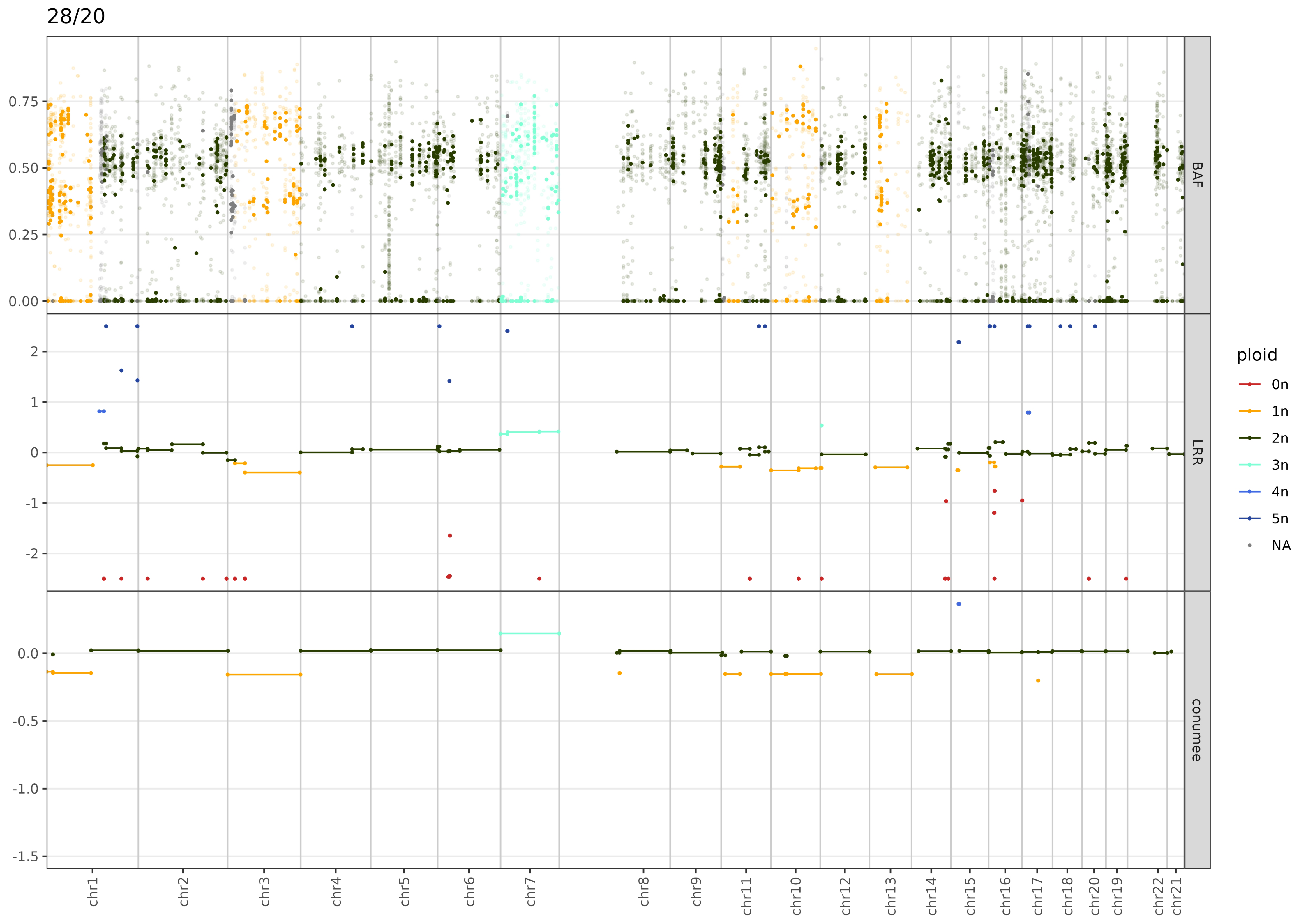

28/20
BAF
LRR
conumee
0.75
0.50
0.25
0.00
2
1
0
-1
-2
0.0
-0.5
-1.0
-1.5
chr1
chr2
chr3
chr4
chr5
chr6
chr7
chr8
chr9
chr11
chr10
chr12
chr13
chr14
chr15
chr16
chr17
chr18
chr20
chr19
chr22
chr21
ploid
0n
1n
2n
3n
4n
5n
NA

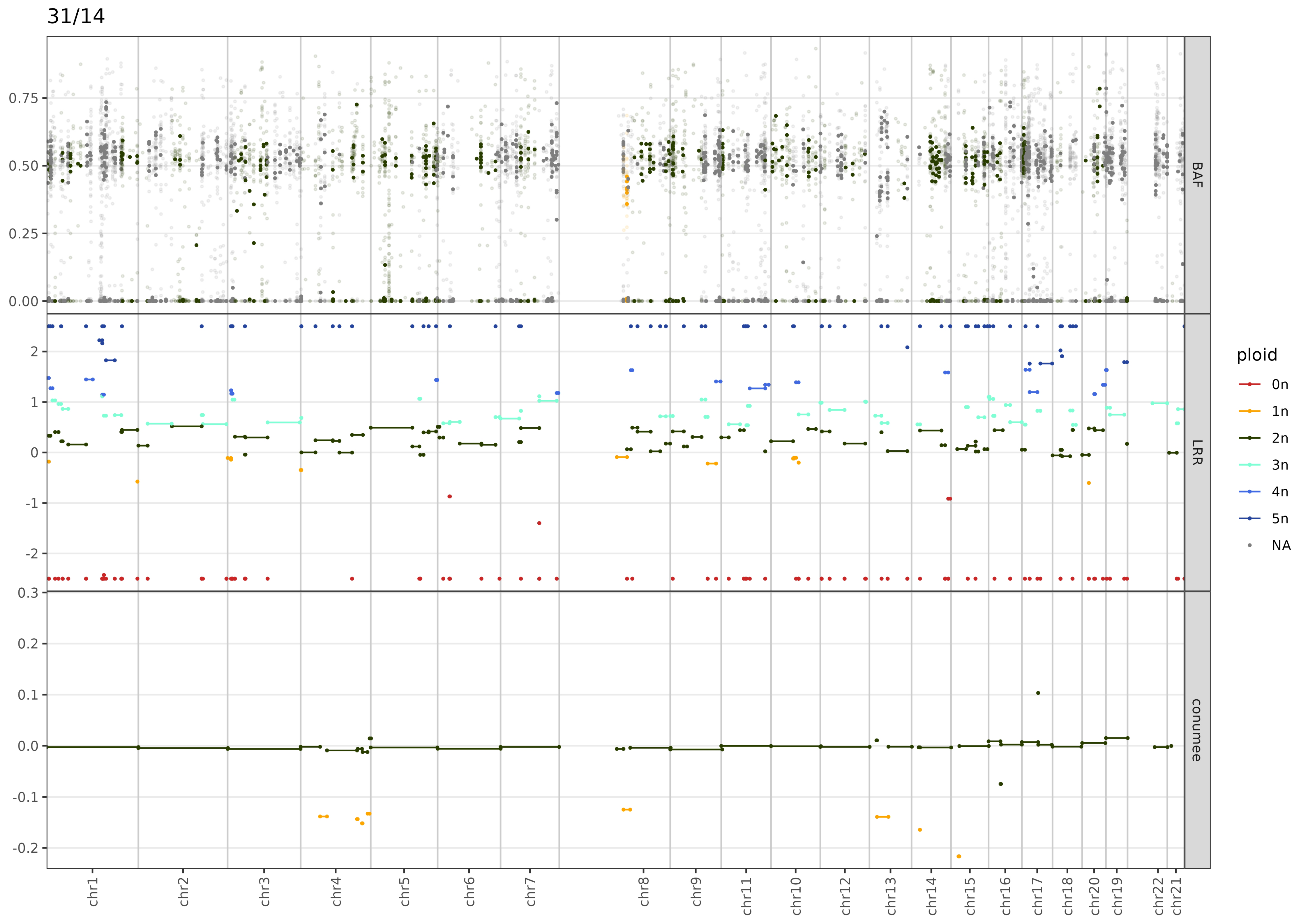

31/14
BAF
LRR
conumee
ploid
0n
1n
2n
3n
4n
5n
NA
chr1
chr2
chr3
chr4
chr5
chr6
chr7
chr8
chr9
chr11
chr10
chr12
chr13
chr14
chr15
chr16
chr17
chr18
chr20
chr19
chr22
chr21

31/20

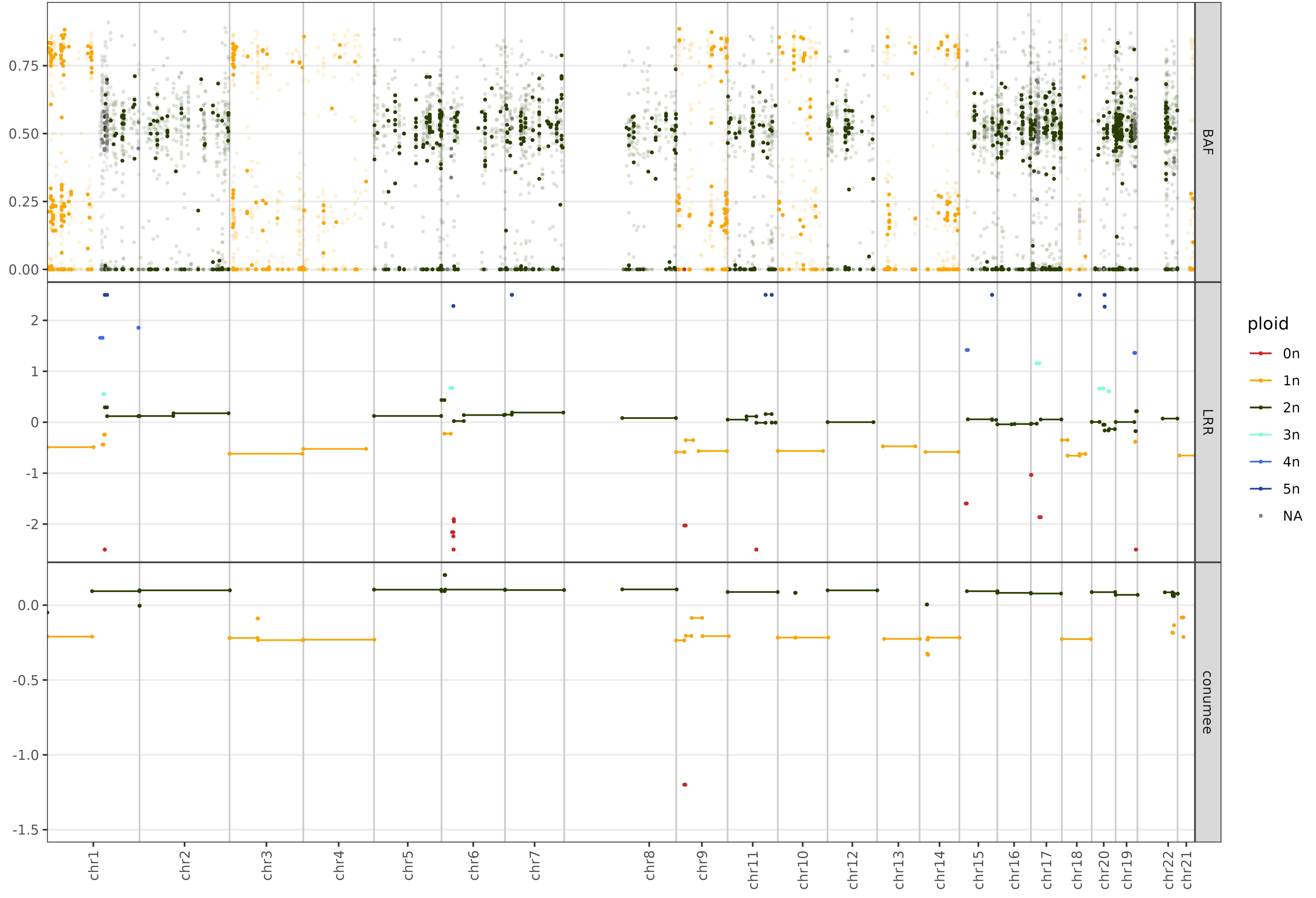
BAF
LRR
conumee
ploid
0n
1n
2n
3n
4n
5n
NA
0.75
0.50
0.25
0.00
2
1
0
-1
-2
0.0
-0.5
-1.0
-1.5
chr1
chr2
chr3
chr4
chr5
chr6
chr7
chr8
chr9
chr11
chr10
chr12
chr13
chr14
chr15
chr16
chr17
chr18
chr20
chr19
chr22
chr21

32/14

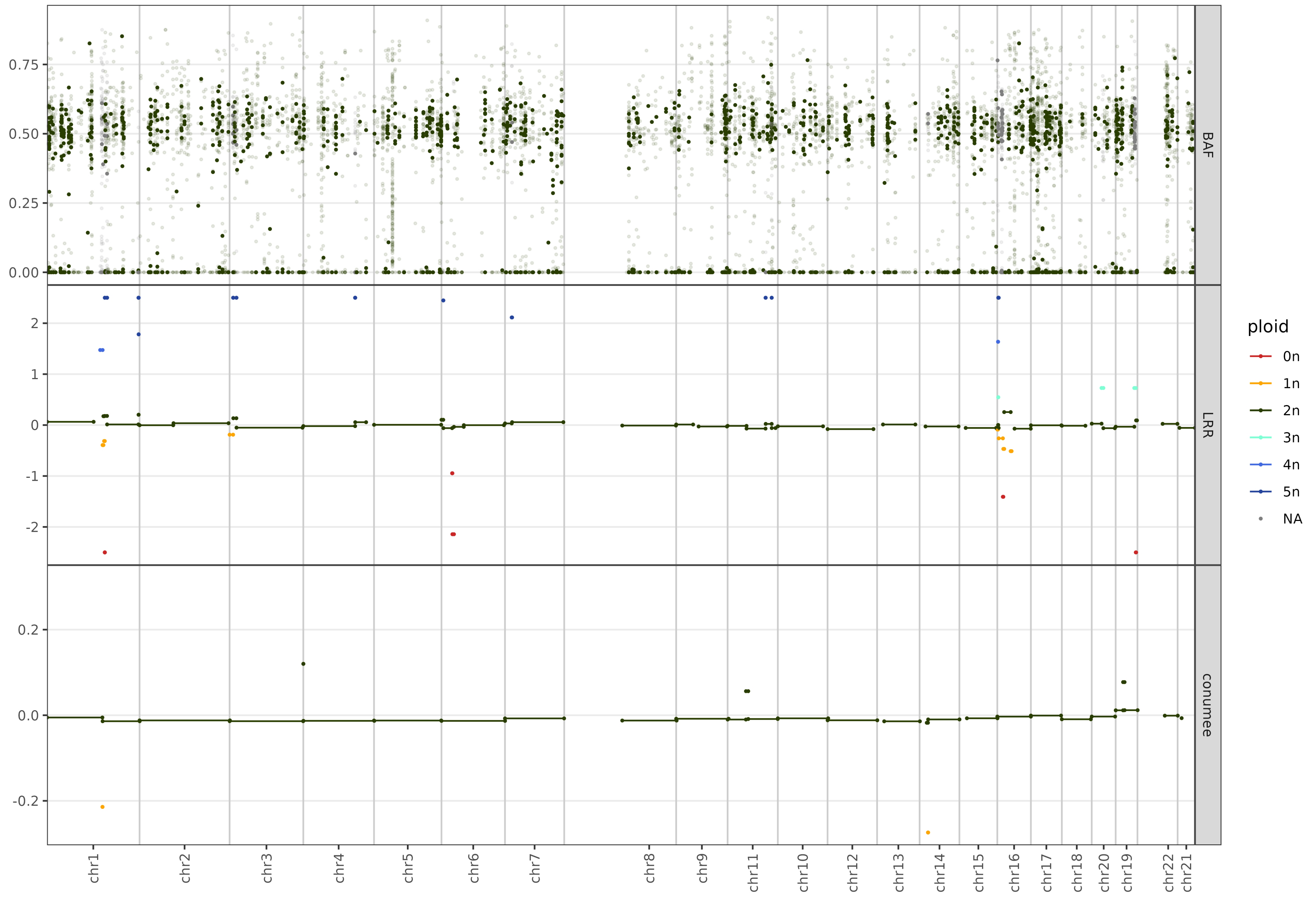

BAF
LRR
conumee
0.75
0.50
0.25
0.00
2
1
0
-1
-2
0.2
0.0
-0.2
chr1
chr2
chr3
chr4
chr5
chr6
chr7
chr8
chr9
chr11
chr10
chr12
chr13
chr14
chr15
chr16
chr17
chr18
chr20
chr19
chr22
chr21
ploid
0n
1n
2n
3n
4n
5n
NA

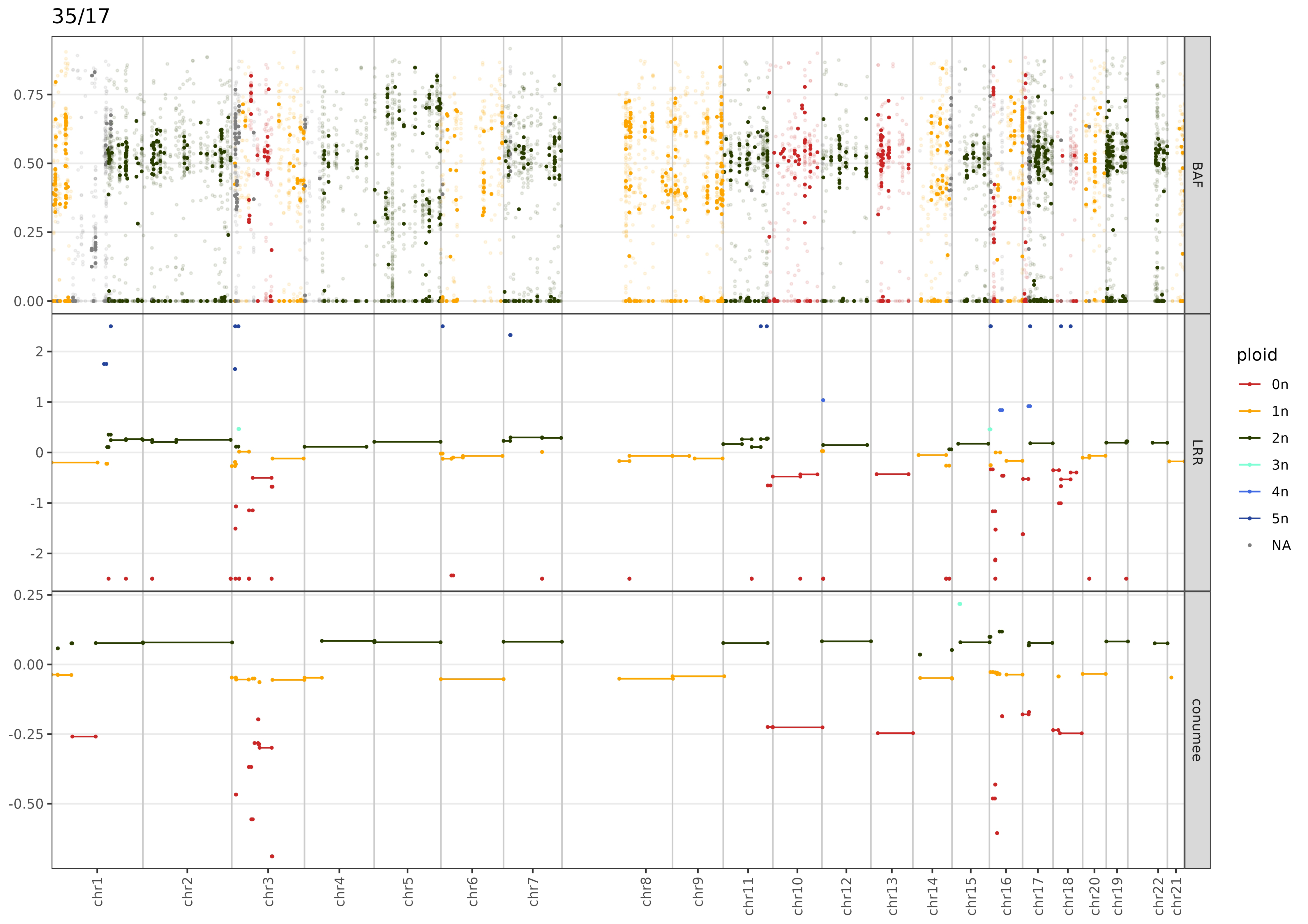

35/17
BAF
LRR
conumee
ploid
0n
1n
2n
3n
4n
5n
NA
0.75
0.50
0.25
0.00
2
1
0
-1
-2
0.25
0.00
-0.25
-0.50
chr1
chr2
chr3
chr4
chr5
chr6
chr7
chr8
chr9
chr11
chr10
chr12
chr13
chr14
chr15
chr16
chr17
chr18
chr20
chr19
chr22
chr21

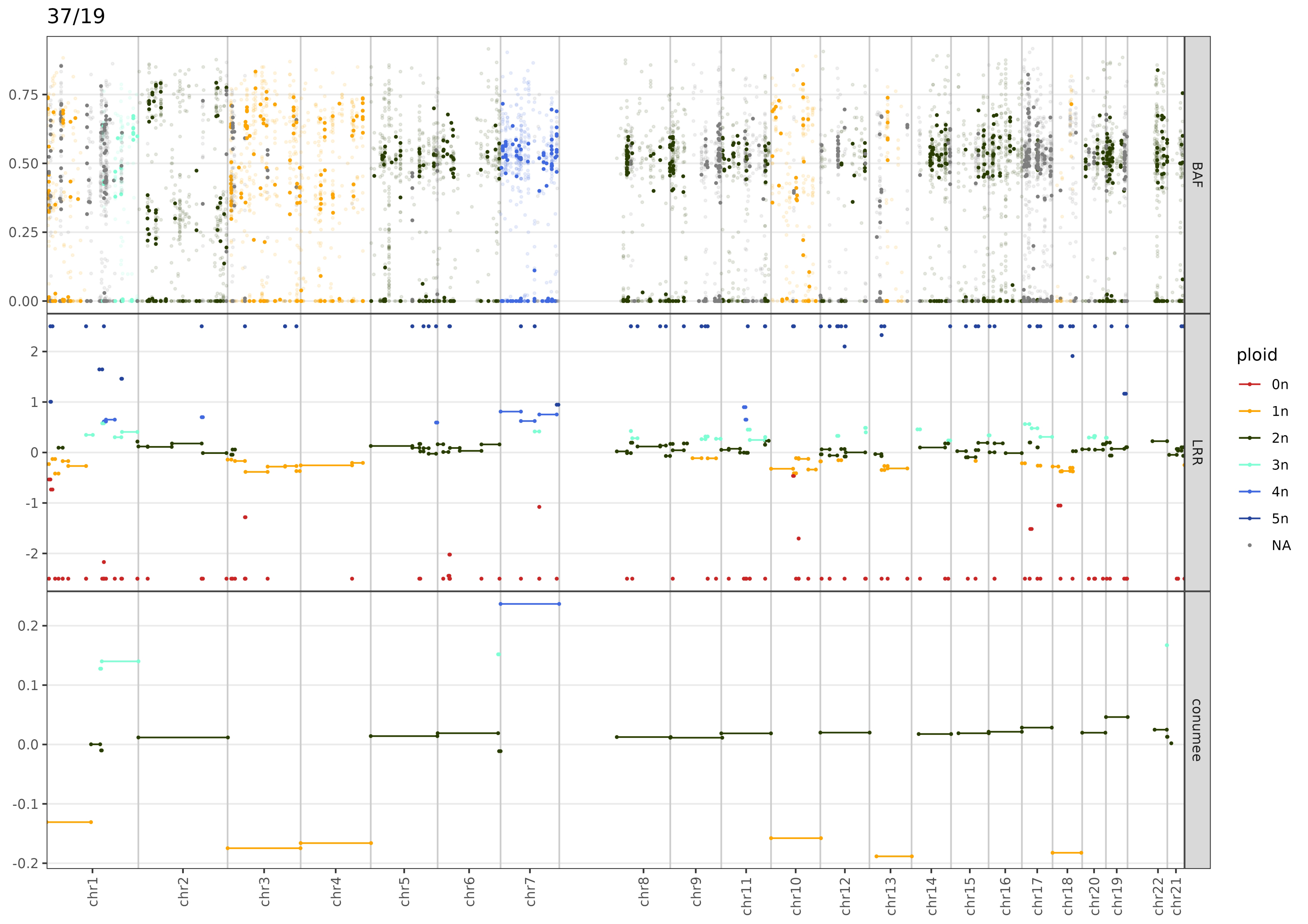

37/19
BAF
LRR
conumee
0.75
0.50
0.25
0.00
2
1
0
-1
-2
0.2
0.1
0.0
-0.1
-0.2
chr1
chr2
chr3
chr4
chr5
chr6
chr7
chr8
chr9
chr11
chr10
chr12
chr13
chr14
chr15
chr16
chr17
chr18
chr20
chr19
chr22
chr21
ploid
0n
1n
2n
3n
4n
5n
NA

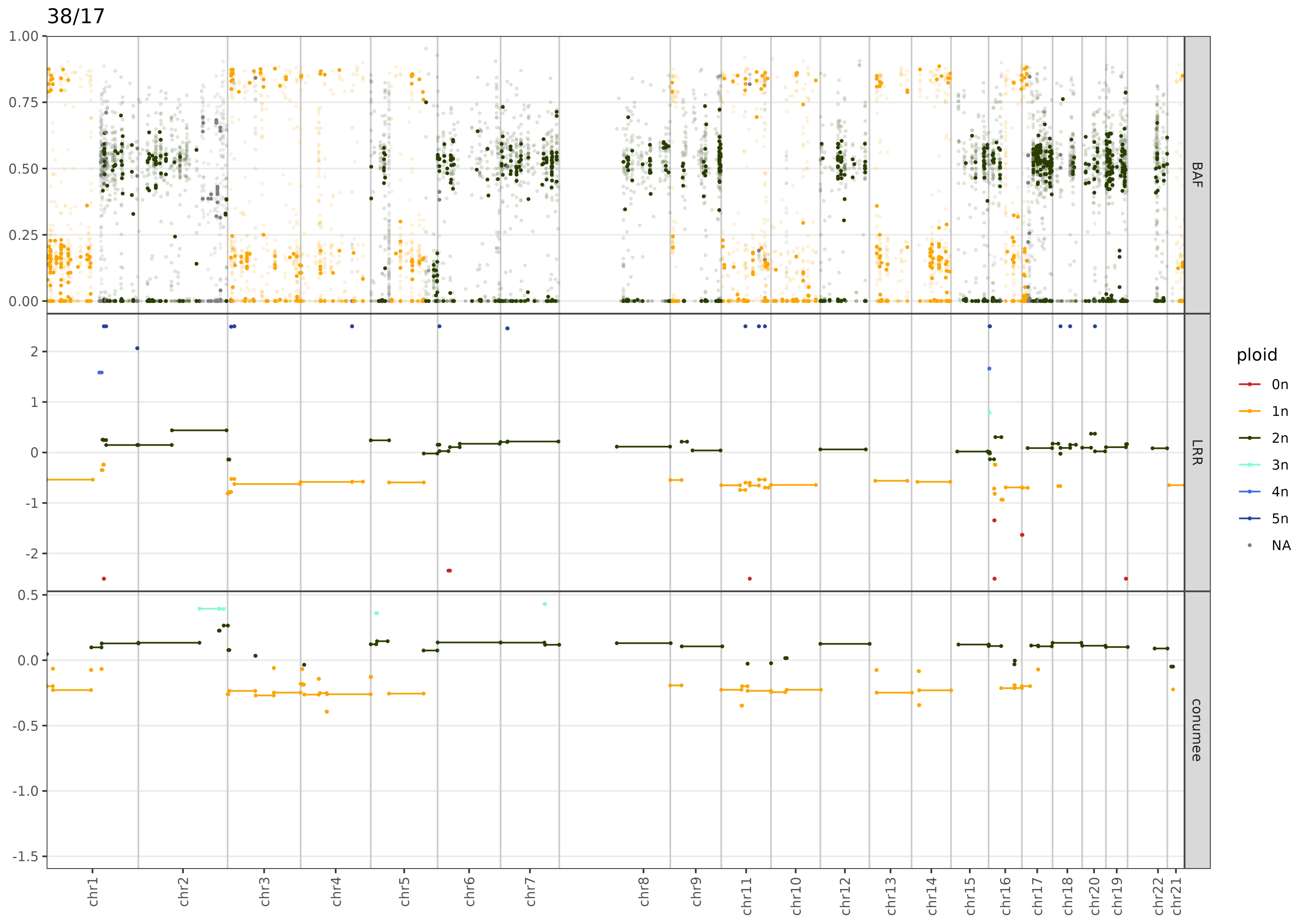

38/17
1.00
0.75
0.50
0.25
0.00
BAF
2
1
0
-1
-2
LRR
0.5
0.0
-0.5
-1.0
-1.5
conumee
chr1
chr2
chr3
chr4
chr5
chr6
chr7
chr8
chr9
chr11
chr10
chr12
chr13
chr14
chr15
chr16
chr17
chr18
chr20
chr19
chr22
chr21
ploid
0n
1n
2n
3n
4n
5n
NA

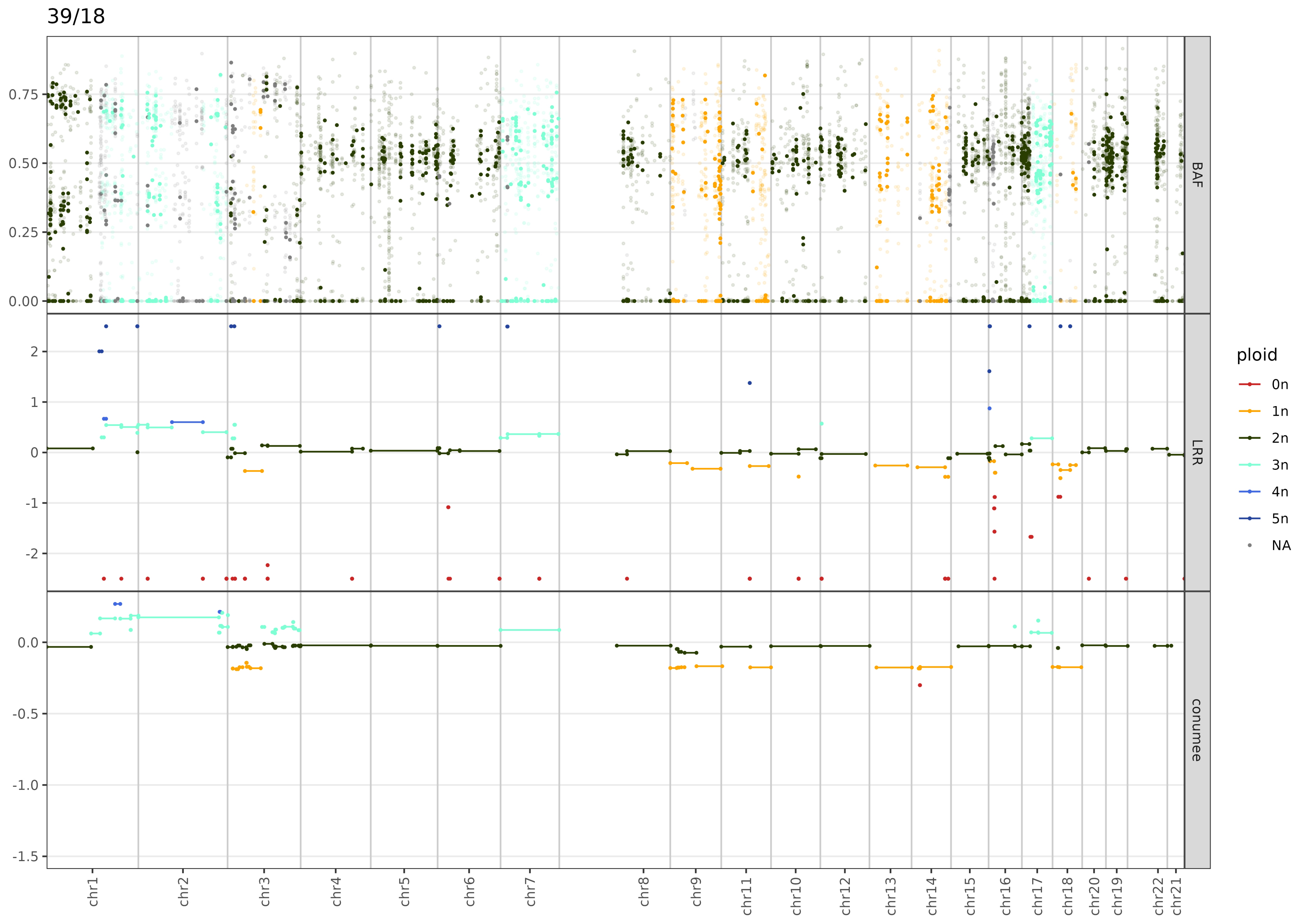

39/18
BAF
LRR
conumee
ploid
0n
1n
2n
3n
4n
5n
NA
chr1
chr2
chr3
chr4
chr5
chr6
chr7
chr8
chr9
chr11
chr10
chr12
chr13
chr14
chr15
chr16
chr17
chr18
chr20
chr19
chr22
chr21

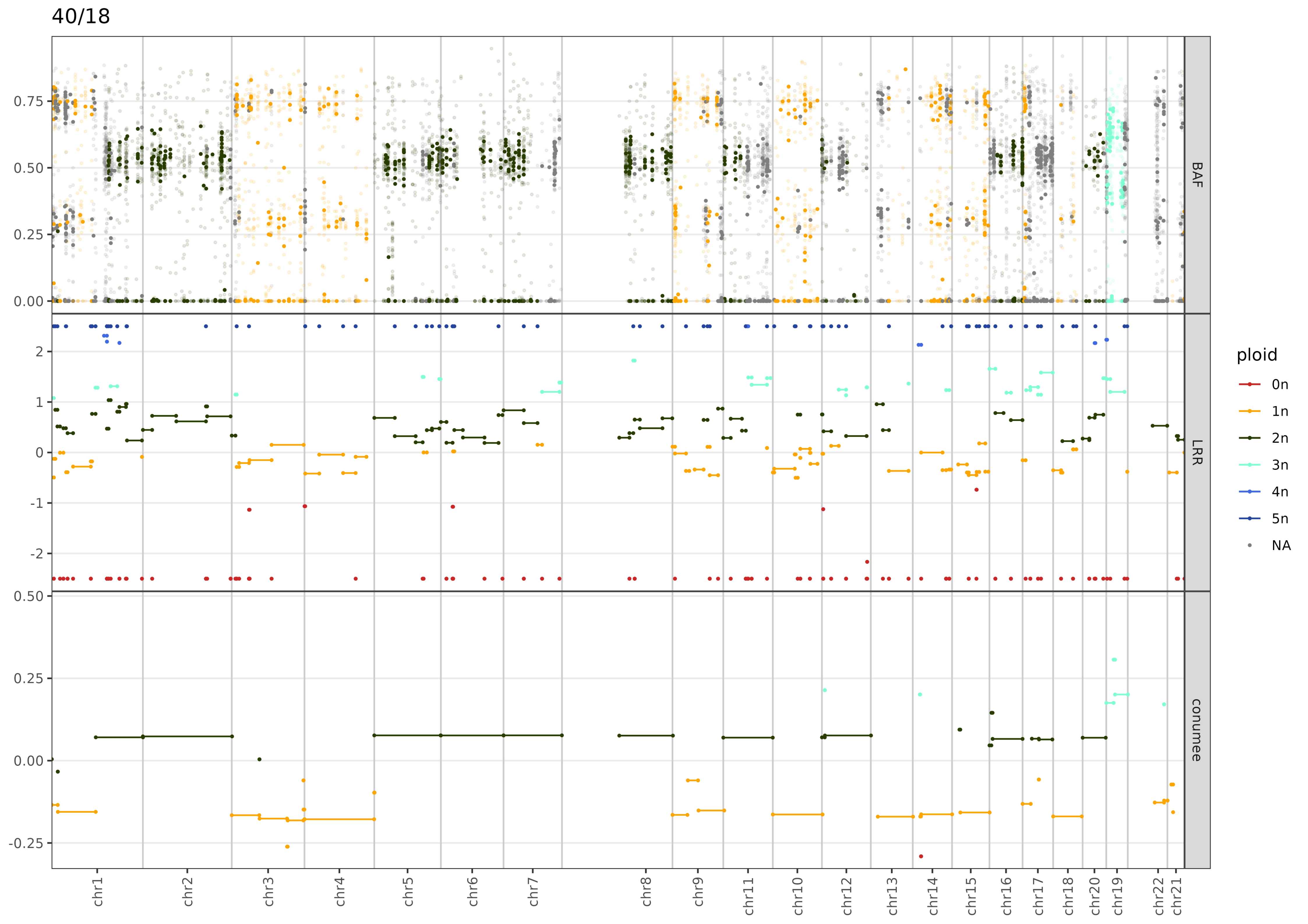

40/18
BAF
LRR
conumee
ploid
0n
1n
2n
3n
4n
5n
NA
0.75
0.50
0.25
0.00
2
1
0
-1
-2
0.50
0.25
0.00
-0.25
chr1
chr2
chr3
chr4
chr5
chr6
chr7
chr8
chr9
chr11
chr10
chr12
chr13
chr14
chr15
chr16
chr17
chr18
chr20
chr19
chr22
chr21

45/17

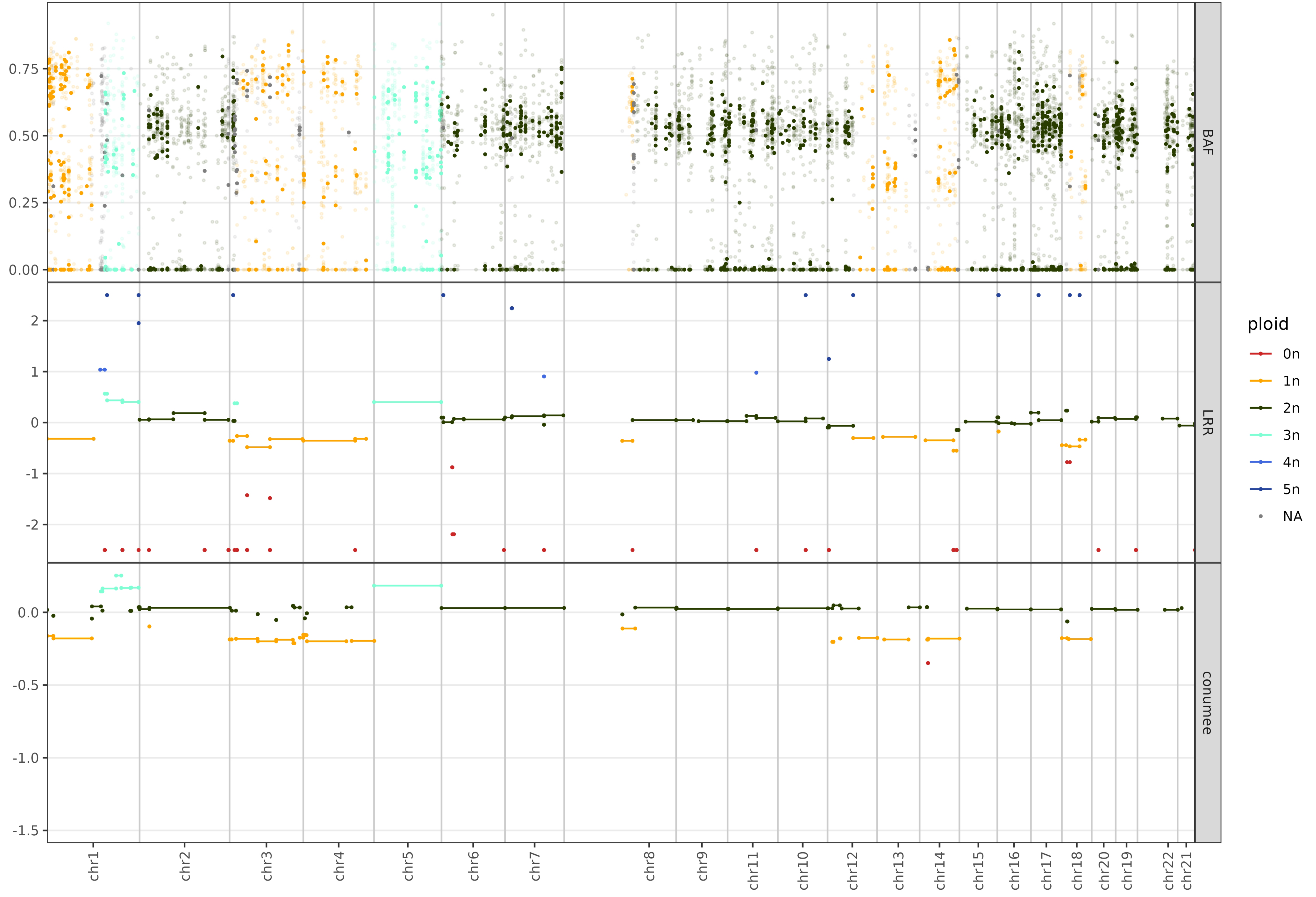

BAF
LRR
conumee
ploid
0n
1n
2n
3n
4n
5n
NA
0.75
0.50
0.25
0.00
2
1
0
-1
-2
0.0
-0.5
-1.0
-1.5
chr1
chr2
chr3
chr4
chr5
chr6
chr7
chr8
chr9
chr11
chr10
chr12
chr13
chr14
chr15
chr16
chr17
chr18
chr20
chr19
chr22
chr21

# 50/15

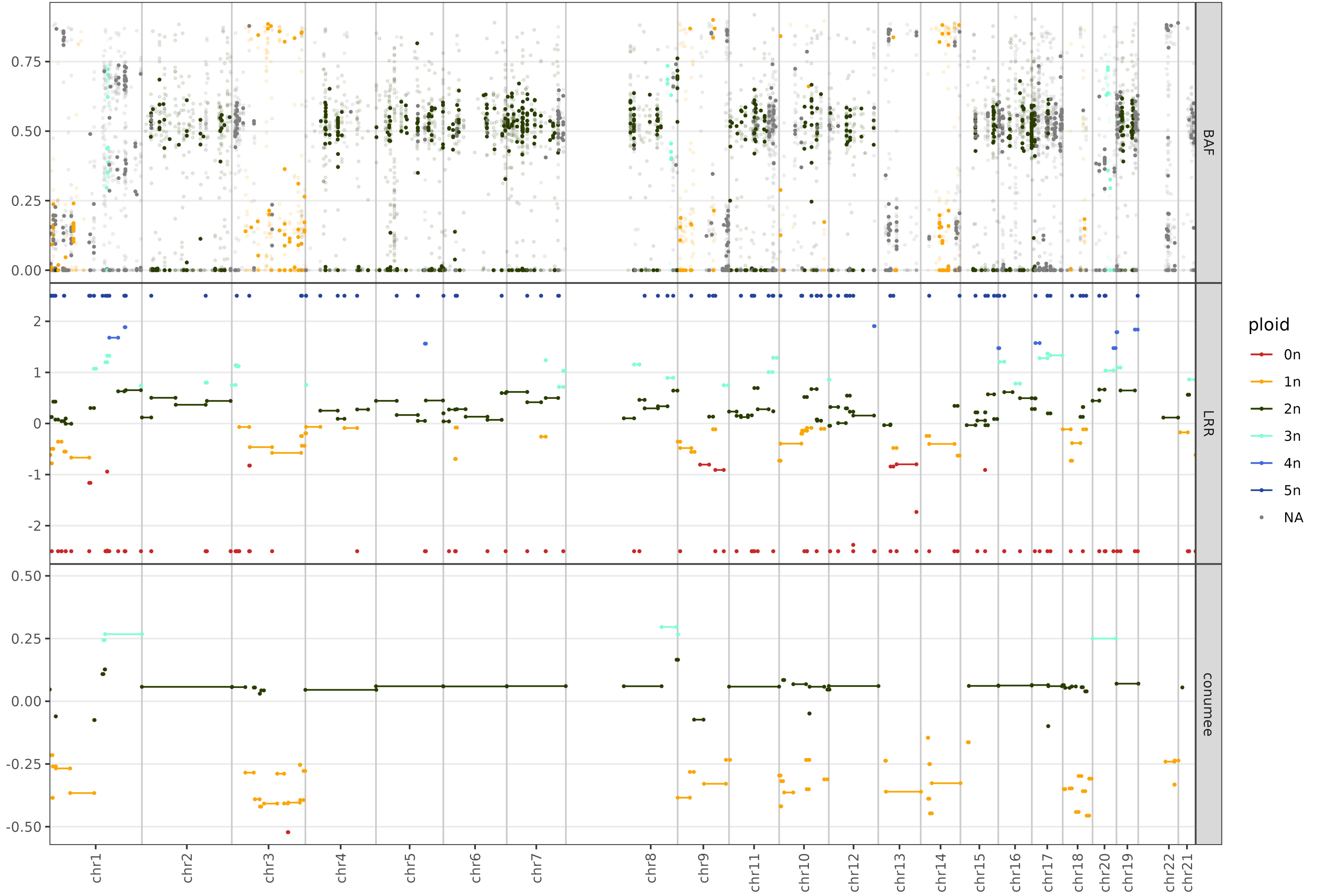

BAF
LRR
conumee
0.75
0.50
0.25
0.00
2
1
0
-1
-2
0.50
0.25
0.00
-0.25
-0.50
chr1
chr2
chr3
chr4
chr5
chr6
chr7
chr8
chr9
chr11
chr10
chr12
chr13
chr14
chr15
chr16
chr17
chr18
chr20
chr19
chr22
chr21
ploid
0n
1n
2n
3n
4n
5n
NA

50/19

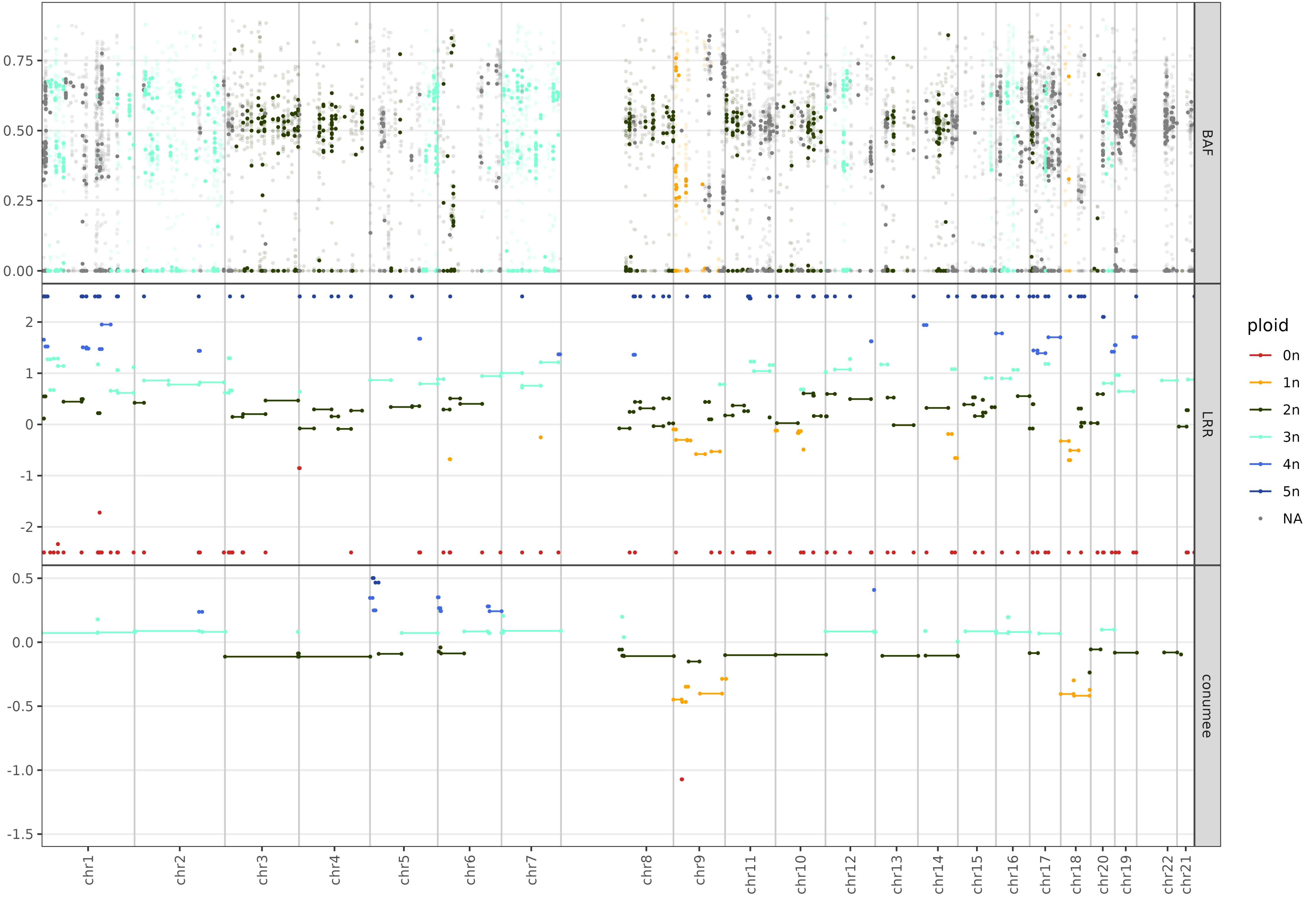

BAF
LRR
conumee
ploid
0n
1n
2n
3n
4n
5n
NA
0.00
0.25
0.50
0.75
2
1
0
-1
-2
0.5
0.0
-0.5
-1.0
-1.5
chr1
chr2
chr3
chr4
chr5
chr6
chr7
chr8
chr9
chr11
chr10
chr12
chr13
chr14
chr15
chr16
chr17
chr18
chr20
chr19
chr22
chr21

53/18

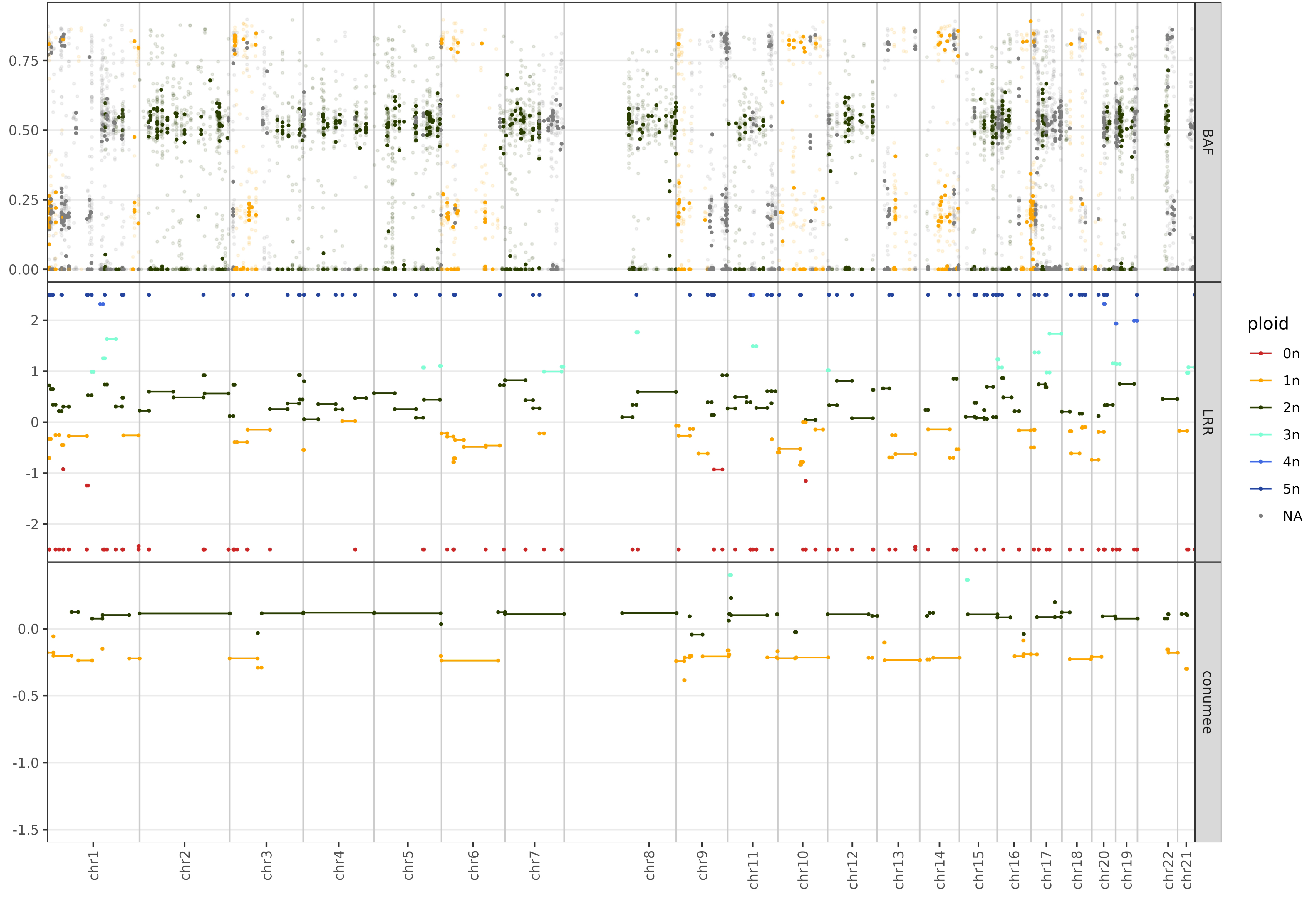

BAF
LRR
conumee
ploid
0n
1n
2n
3n
4n
5n
NA
0.75
0.50
0.25
0.00
2
1
0
-1
-2
0.0
-0.5
-1.0
-1.5
chr1
chr2
chr3
chr4
chr5
chr6
chr7
chr8
chr9
chr11
chr10
chr12
chr13
chr14
chr15
chr16
chr17
chr18
chr20
chr19
chr22
chr21

54/15

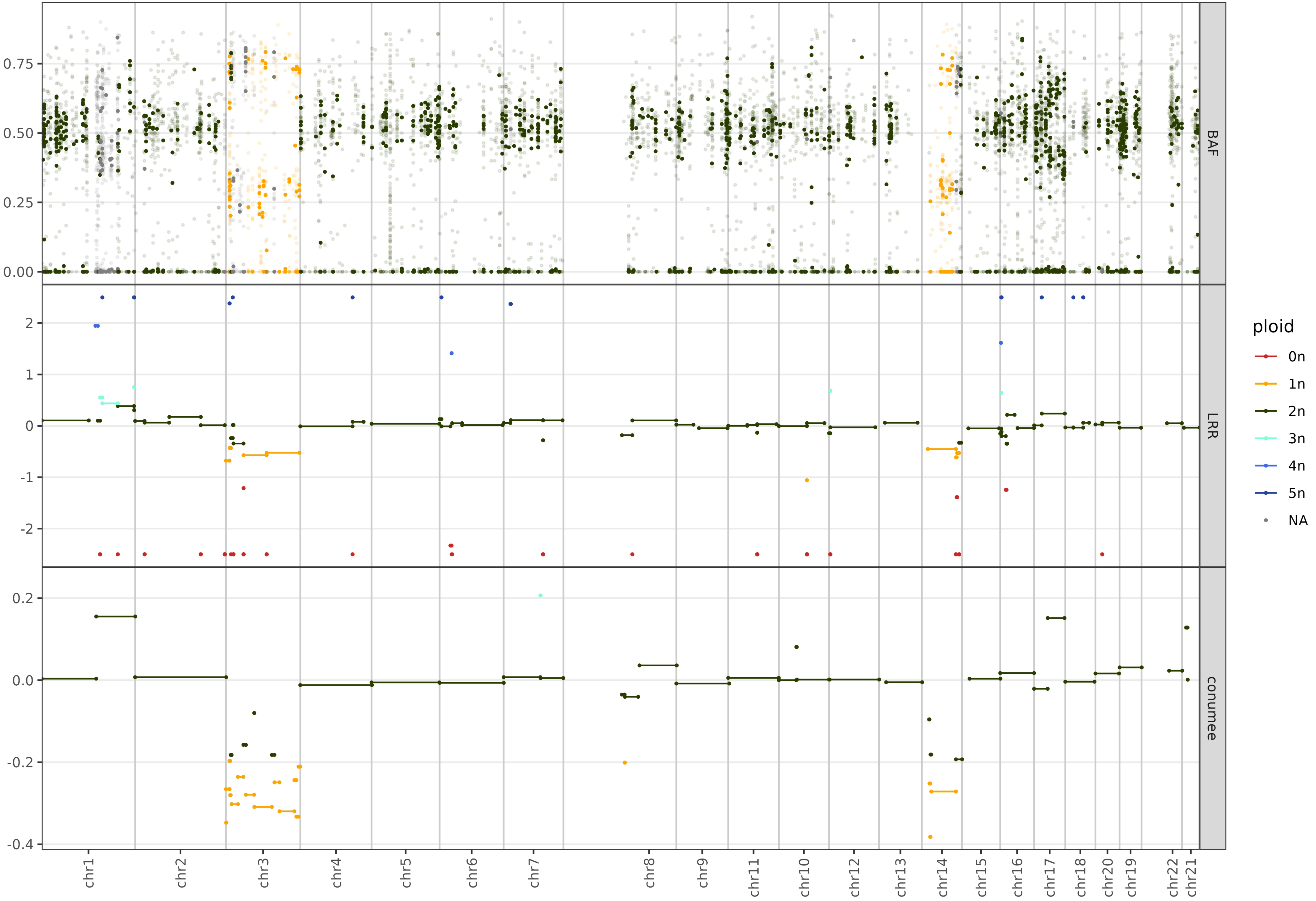

BAF
LRR
conumee
0.75
0.50
0.25
0.00
2
1
0
-1
-2
0.2
0.0
-0.2
-0.4
chr1
chr2
chr3
chr4
chr5
chr6
chr7
chr8
chr9
chr11
chr10
chr12
chr13
chr14
chr15
chr16
chr17
chr18
chr20
chr19
chr22
chr21
ploid
0n
1n
2n
3n
4n
5n
NA

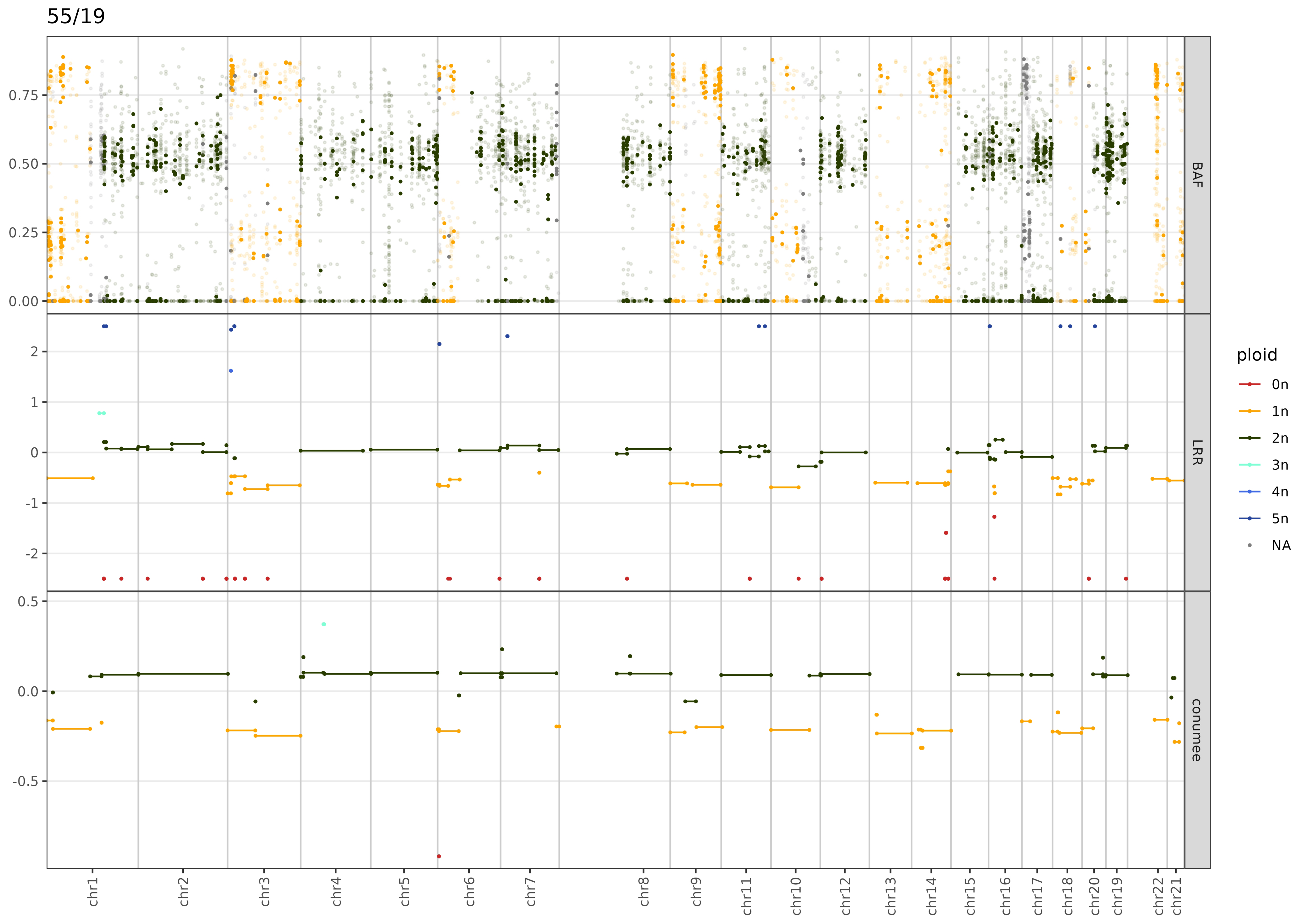

55/19
BAF
LRR
conumee
ploid
0n
1n
2n
3n
4n
5n
NA
0.75
0.50
0.25
0.00
2
1
0
-1
-2
0.5
0.0
-0.5
chr1
chr2
chr3
chr4
chr5
chr6
chr7
chr8
chr9
chr11
chr10
chr12
chr13
chr14
chr15
chr16
chr17
chr18
chr20
chr19
chr22
chr21

62/15

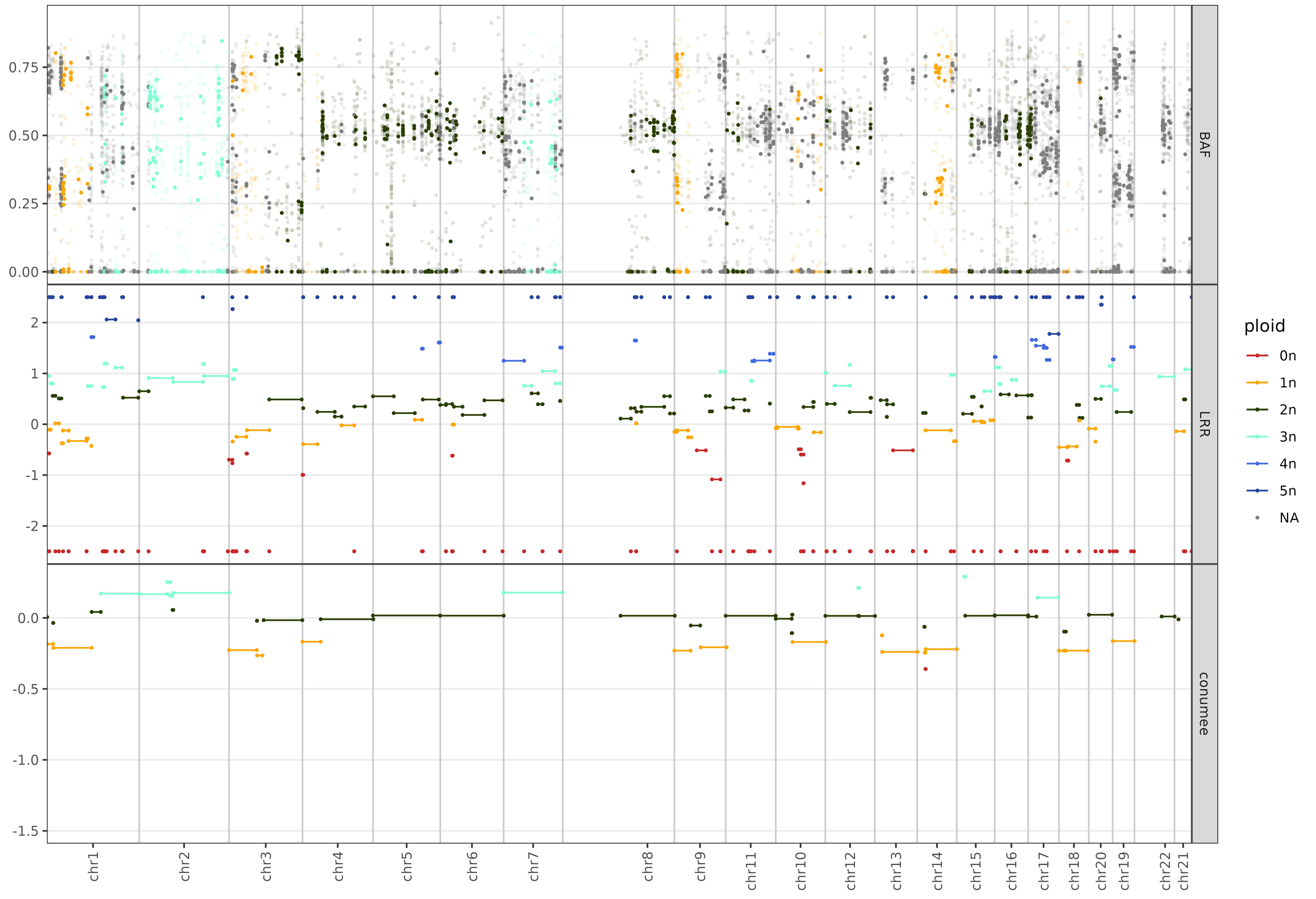

BAF
LRR
conumee
ploid
0n
1n
2n
3n
4n
5n
NA
0.75
0.50
0.25
0.00
2
1
0
-1
-2
0.0
-0.5
-1.0
-1.5
chr1
chr2
chr3
chr4
chr5
chr6
chr7
chr8
chr9
chr11
chr10
chr12
chr13
chr14
chr15
chr16
chr17
chr18
chr20
chr19
chr22
chr21

68/18

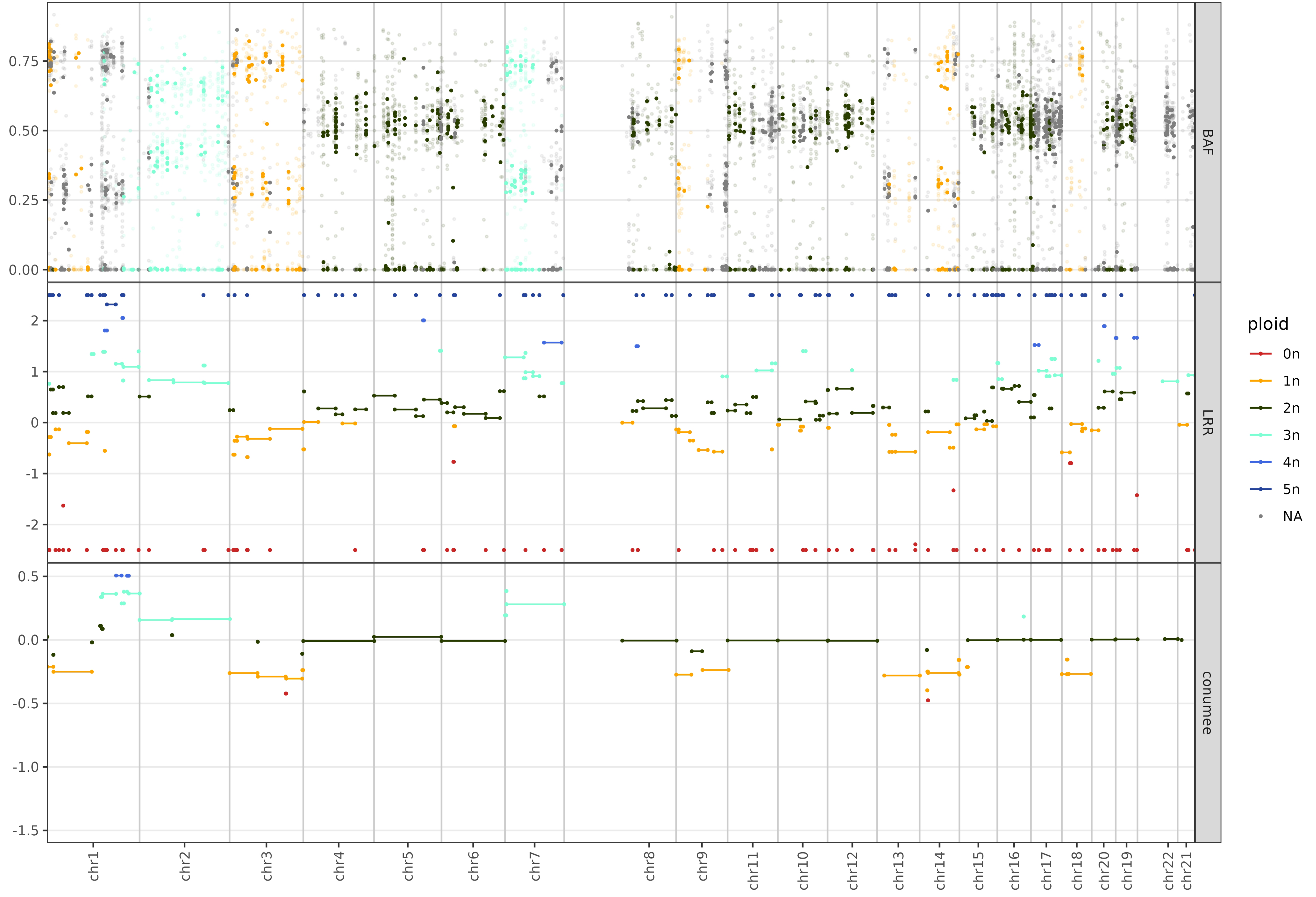

BAF
LRR
conumee
ploid
0n
1n
2n
3n
4n
5n
NA
0.75
0.50
0.25
0.00
2
1
0
-1
-2
0.5
0.0
-0.5
-1.0
-1.5
chr1
chr2
chr3
chr4
chr5
chr6
chr7
chr8
chr9
chr11
chr10
chr12
chr13
chr14
chr15
chr16
chr17
chr18
chr20
chr19
chr22
chr21

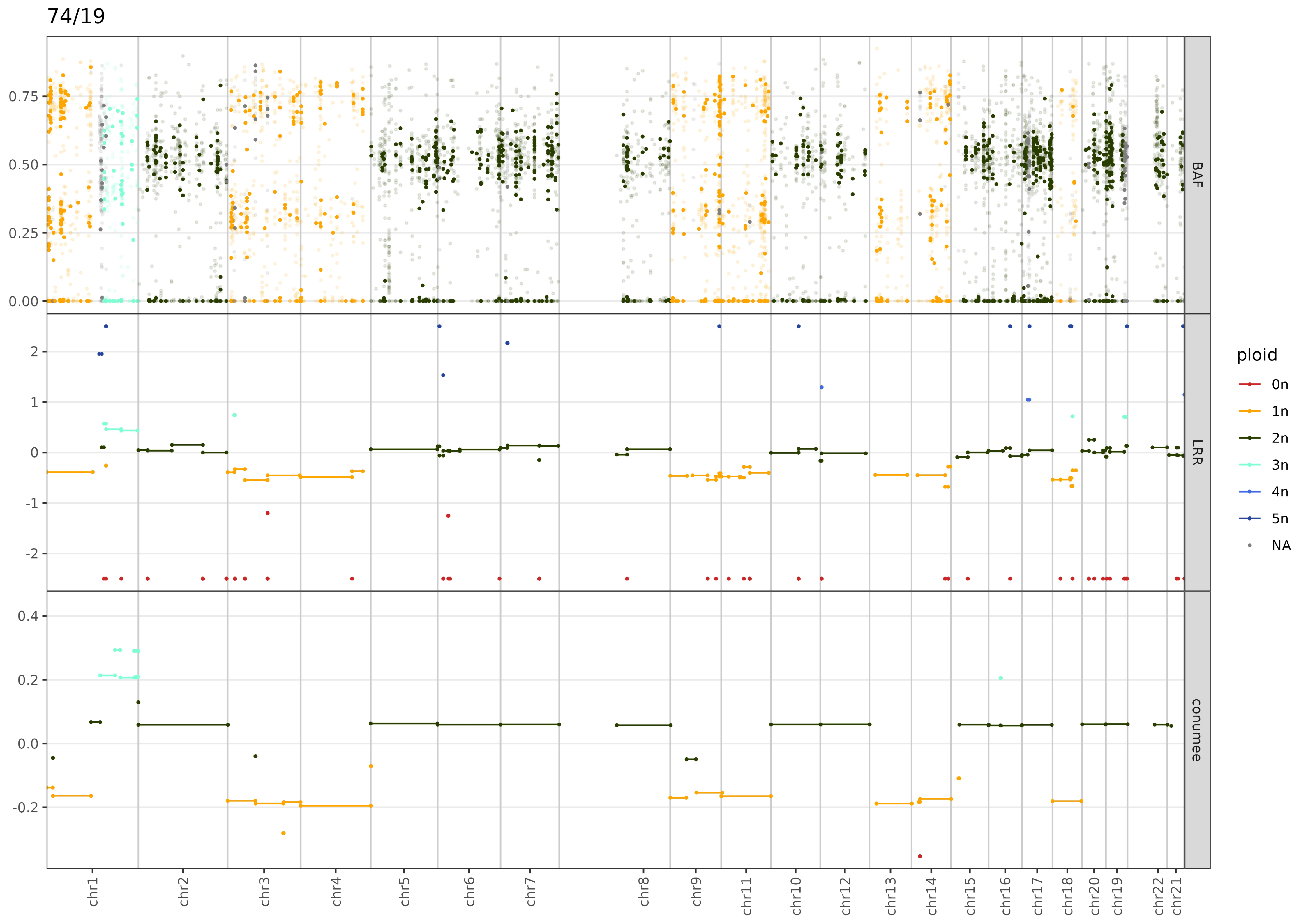

74/19
BAF
LRR
conumee
0.75
0.50
0.25
0.00
2
1
0
-1
-2
0.4
0.2
0.0
-0.2
chr1
chr2
chr3
chr4
chr5
chr6
chr7
chr8
chr9
chr11
chr10
chr12
chr13
chr14
chr15
chr16
chr17
chr18
chr20
chr19
chr22
chr21
ploid
0n
1n
2n
3n
4n
5n
NA

81/15

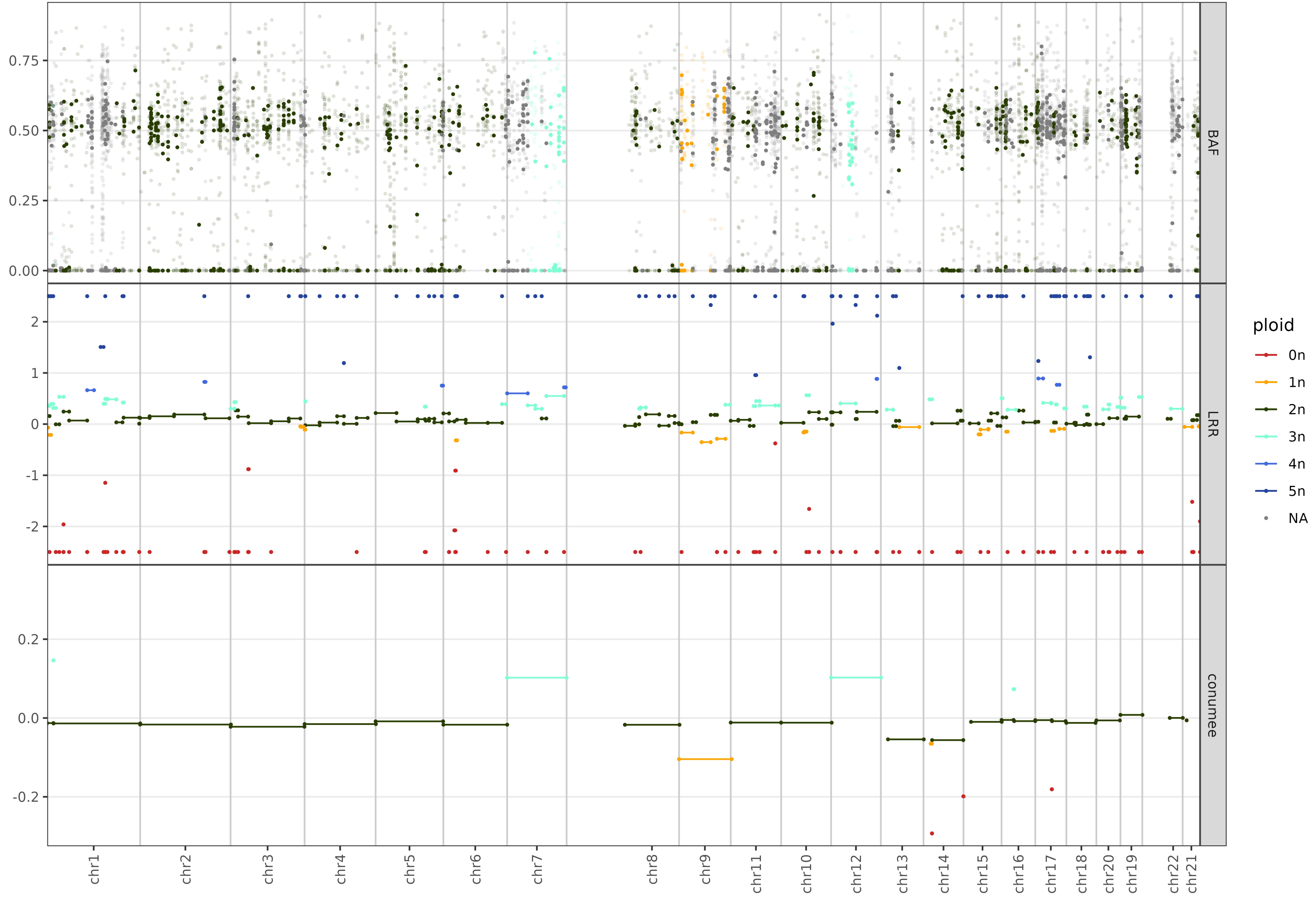

BAF
LRR
conumee
0.75
0.50
0.25
0.00
2
1
0
-1
-2
0.2
0.0
-0.2
chr1
chr2
chr3
chr4
chr5
chr6
chr7
chr8
chr9
chr11
chr10
chr12
chr13
chr14
chr15
chr16
chr17
chr18
chr20
chr19
chr22
chr21
ploid
0n
1n
2n
3n
4n
5n
NA

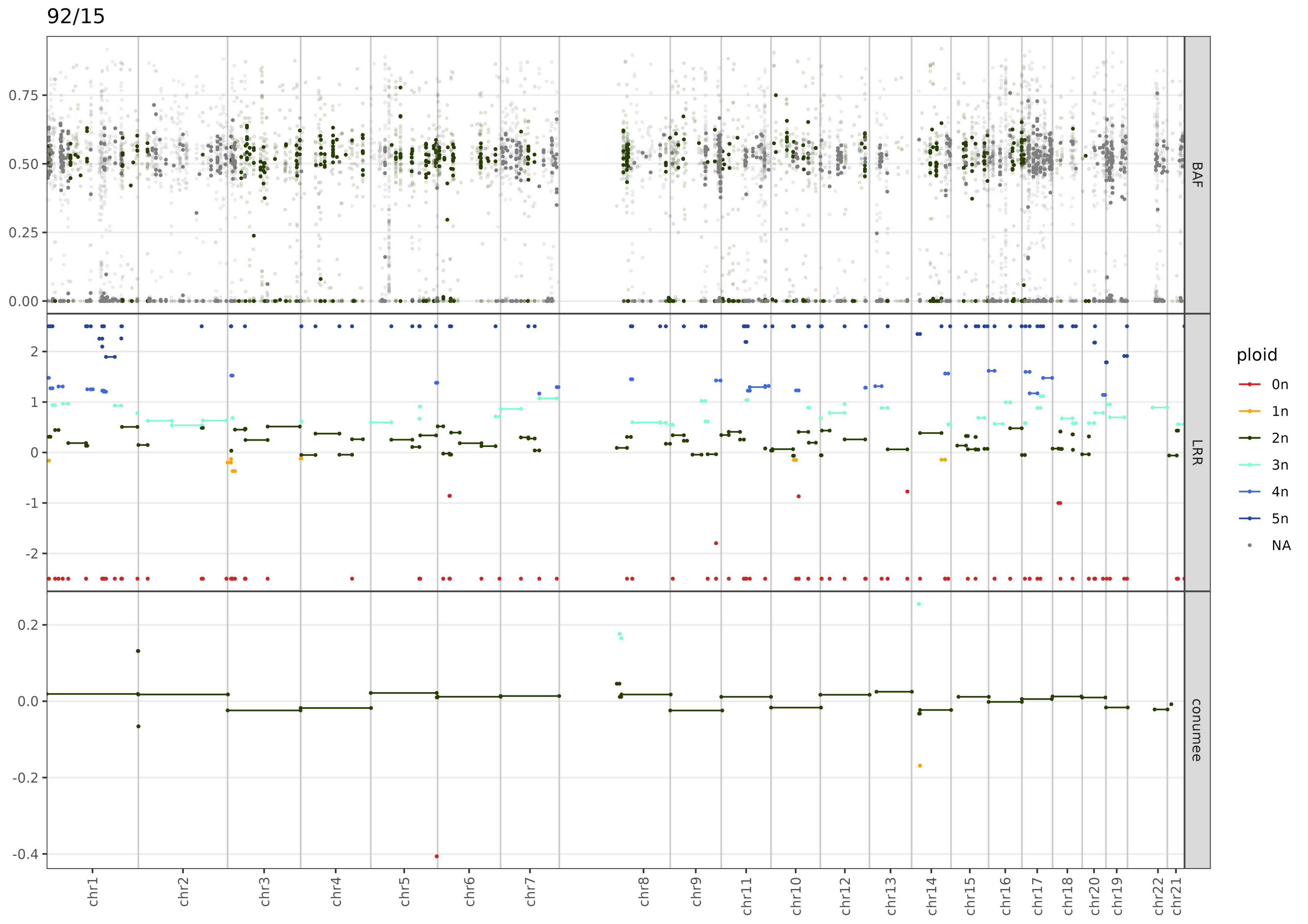

92/15
BAF
LRR
conumee
ploid
0n
1n
2n
3n
4n
5n
NA
0.75
0.50
0.25
0.00
2
1
0
-1
-2
0.2
0.0
-0.2
-0.4
chr1
chr2
chr3
chr4
chr5
chr6
chr7
chr8
chr9
chr11
chr10
chr12
chr13
chr14
chr15
chr16
chr17
chr18
chr20
chr19
chr22
chr21

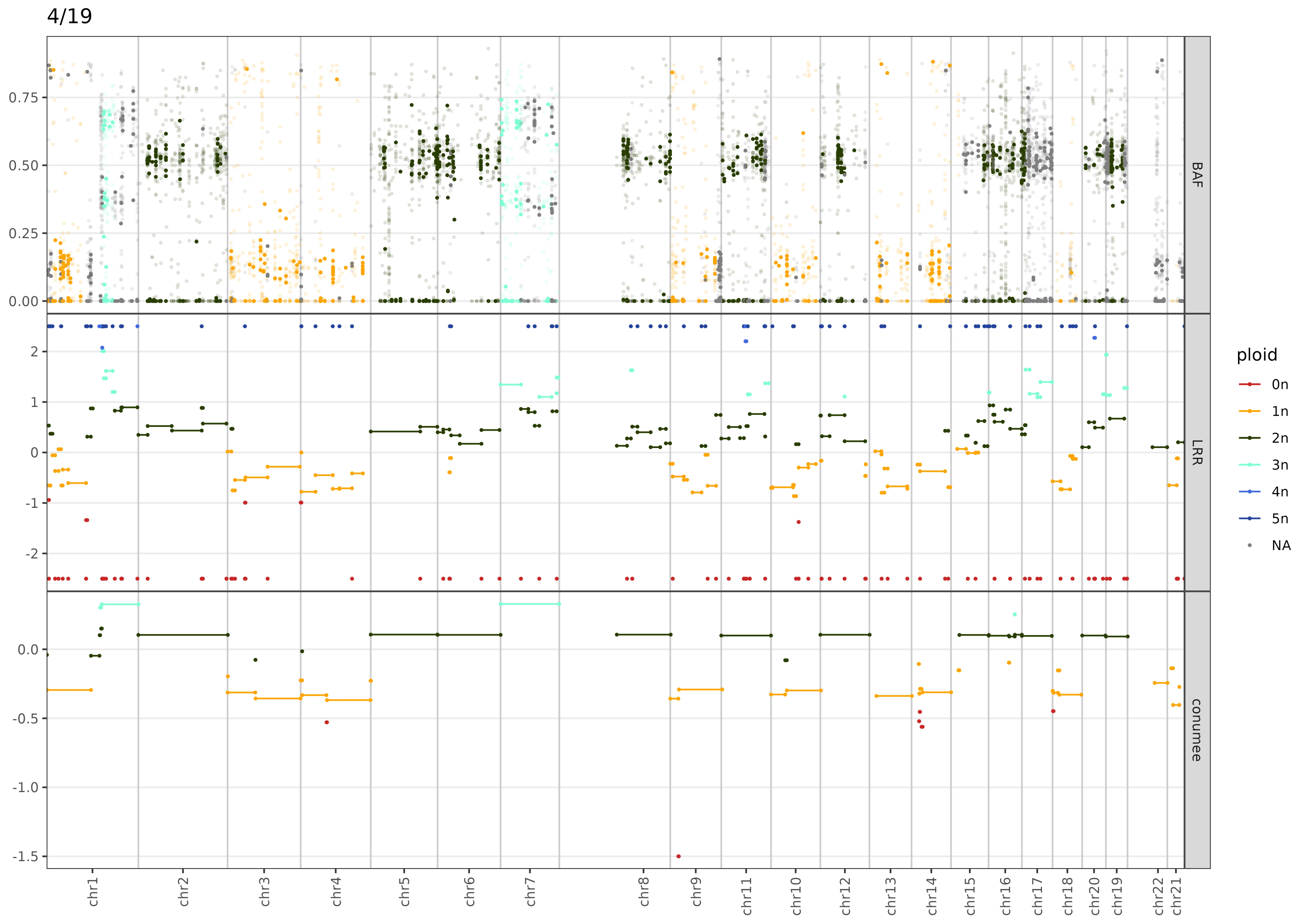

4/19
BAF
LRR
conumee
ploid
0n
1n
2n
3n
4n
5n
NA
0.75
0.50
0.25
0.00
2
1
0
-1
-2
0.0
-0.5
-1.0
-1.5
chr1
chr2
chr3
chr4
chr5
chr6
chr7
chr8
chr9
chr11
chr10
chr12
chr13
chr14
chr15
chr16
chr17
chr18
chr20
chr19
chr22
chr21

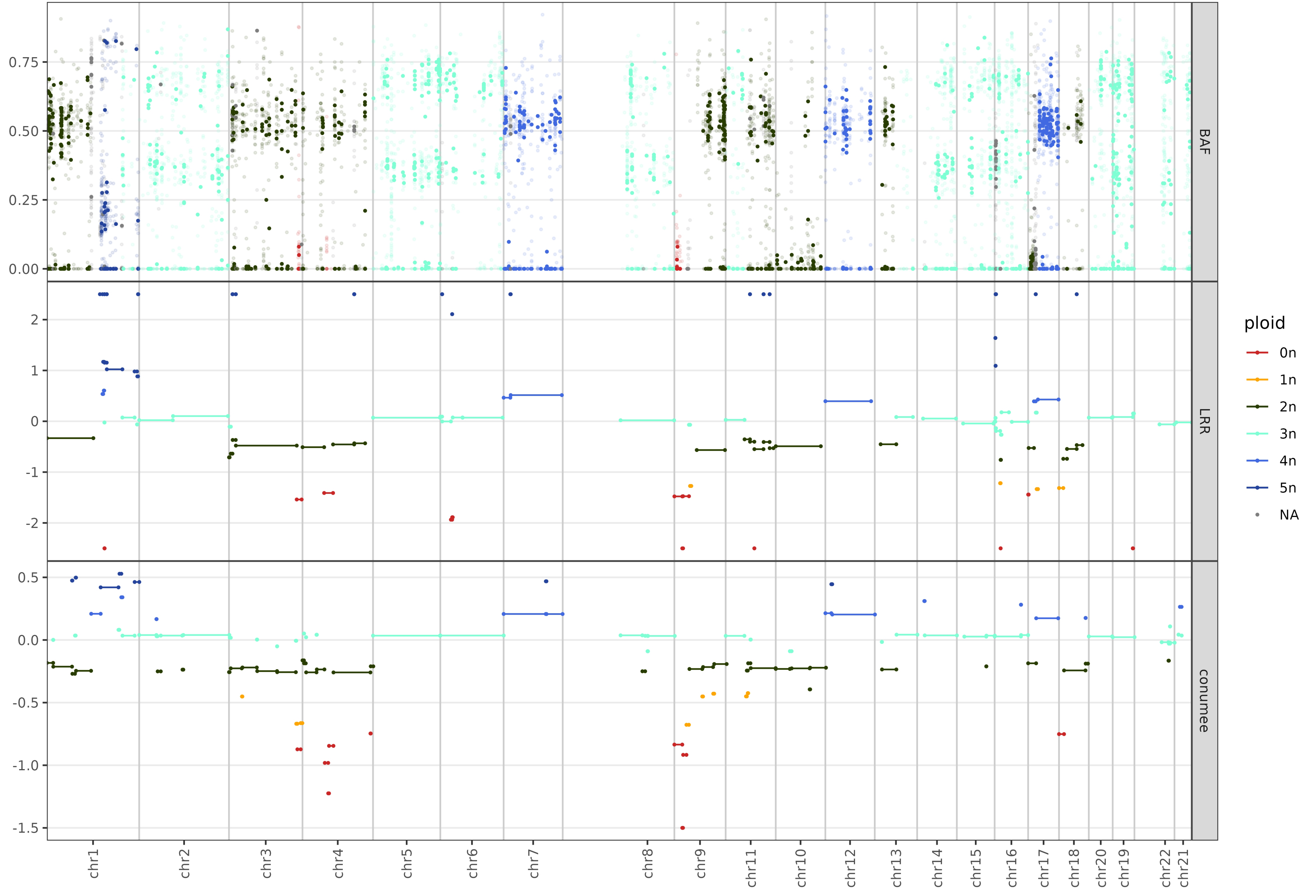

BAF
LRR
conumee
0.75
0.50
0.25
0.00
2
1
0
-1
-2
0.5
0.0
-0.5
-1.0
-1.5
chr1
chr2
chr3
chr4
chr5
chr6
chr7
chr8
chr9
chr11
chr10
chr12
chr13
chr14
chr15
chr16
chr17
chr18
chr20
chr19
chr22
chr21
ploid
0n
1n
2n
3n
4n
5n
NA

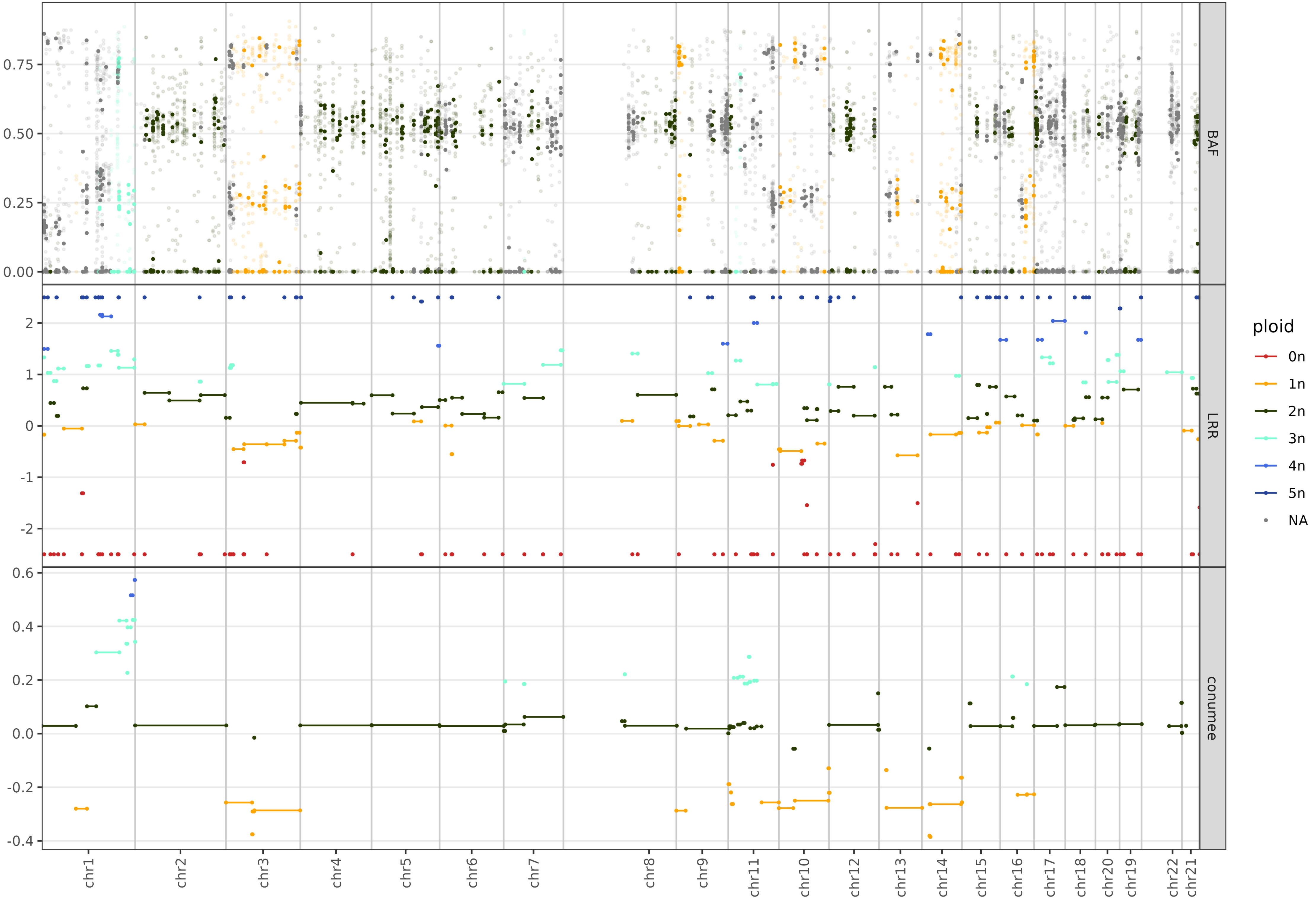

BAF
LRR
conumee
0.75
0.50
0.25
0.00
2
1
0
-1
-2
0.6
0.4
0.2
0.0
-0.2
-0.4
ploid
0n
1n
2n
3n
4n
5n
NA
chr1
chr2
chr3
chr4
chr5
chr6
chr7
chr8
chr9
chr11
chr10
chr12
chr13
chr14
chr15
chr16
chr17
chr18
chr20
chr19
chr22
chr21

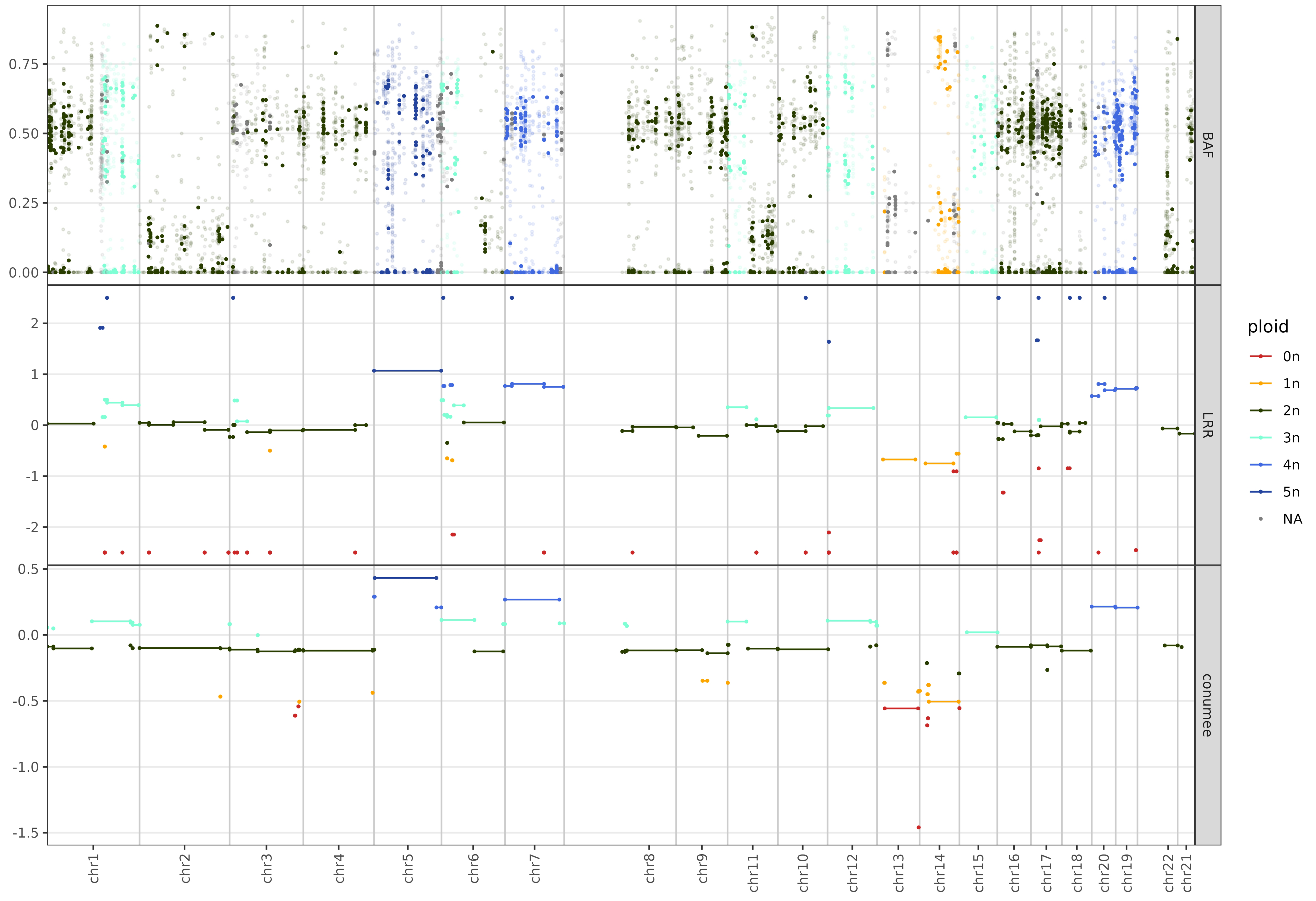
BAF
LRR
conumee
ploid
0n
1n
2n
3n
4n
5n
NA
chr1
chr2
chr3
chr4
chr5
chr6
chr7
chr8
chr9
chr11
chr10
chr12
chr13
chr14
chr15
chr16
chr17
chr18
chr20
chr19
chr22
chr21

9/20

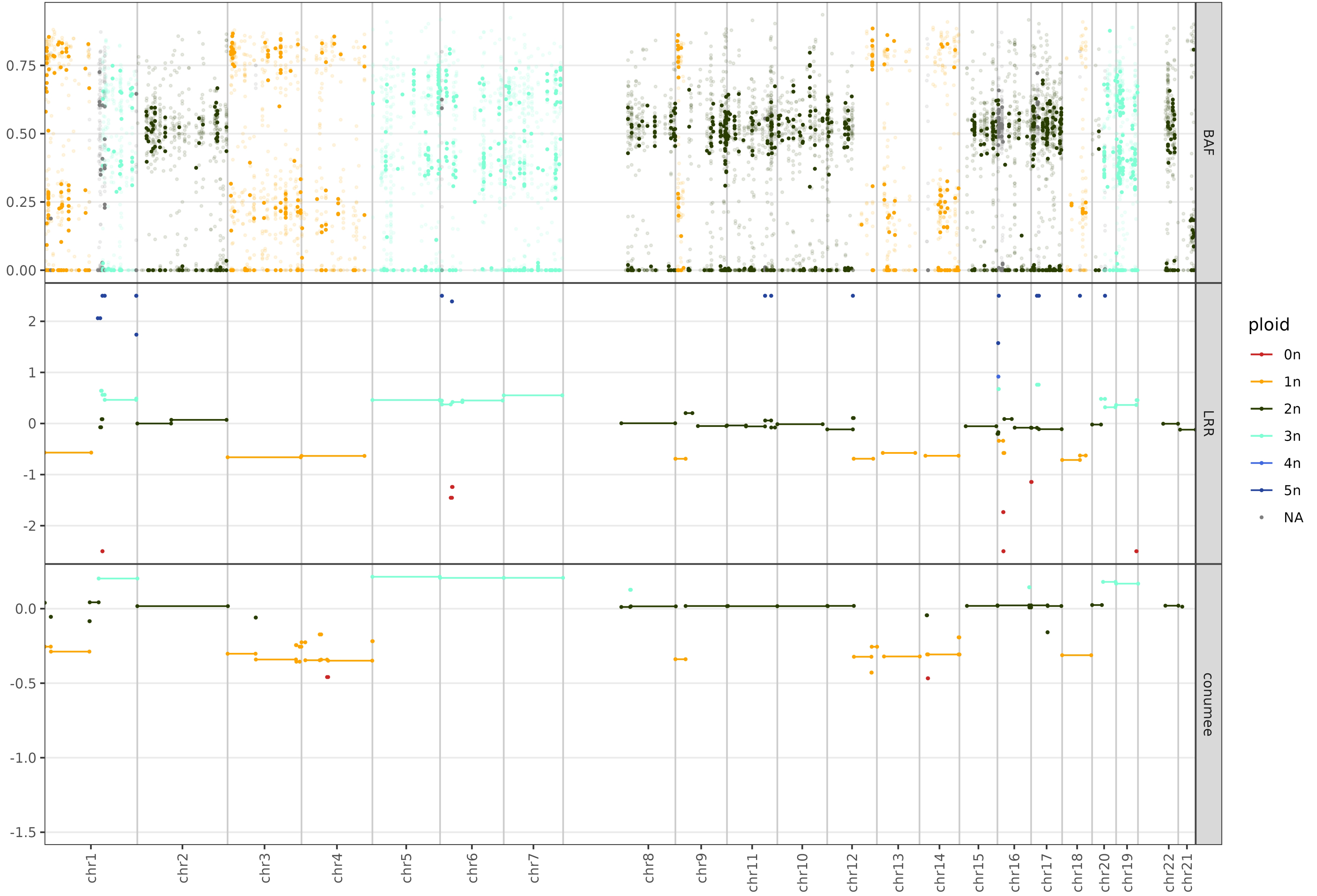
BAF
LRR
conumee
ploid
0n
1n
2n
3n
4n
5n
NA
0.00
0.25
0.50
0.75
-2
-1
0
1
2
-1.5
-1.0
-0.5
0.0
chr1
chr2
chr3
chr4
chr5
chr6
chr7
chr8
chr9
chr11
chr10
chr12
chr13
chr14
chr15
chr16
chr17
chr18
chr20
chr19
chr22
chr21

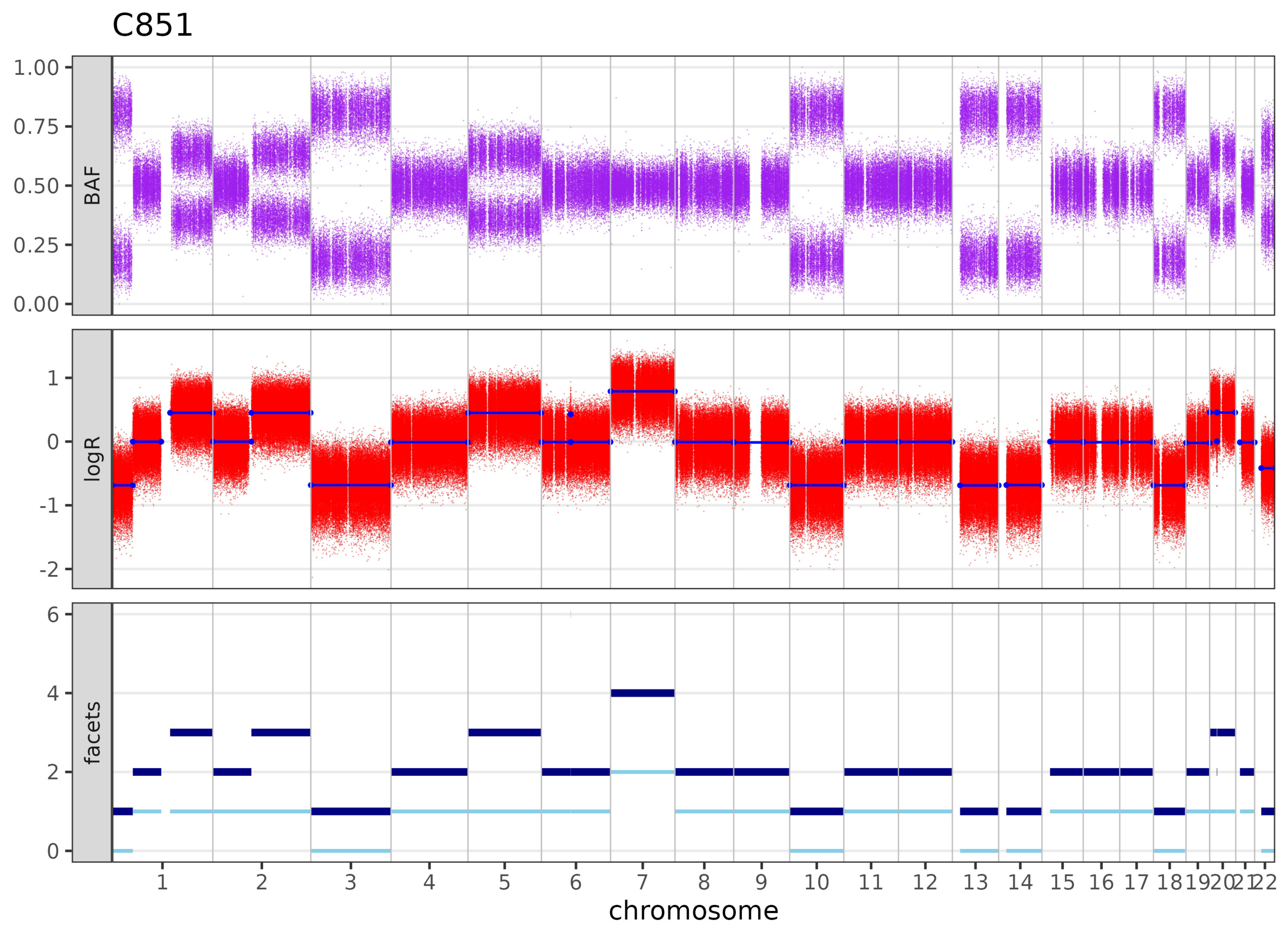
C851
BAF
1.00
0.75
0.50
0.25
0.00
logR
1
0
-1
-2
facets
6
4
2
0
1 2 3 4 5 6 7 8 9 10 11 12 13 14 15 16 17 18 19 20 21 22
chromosome

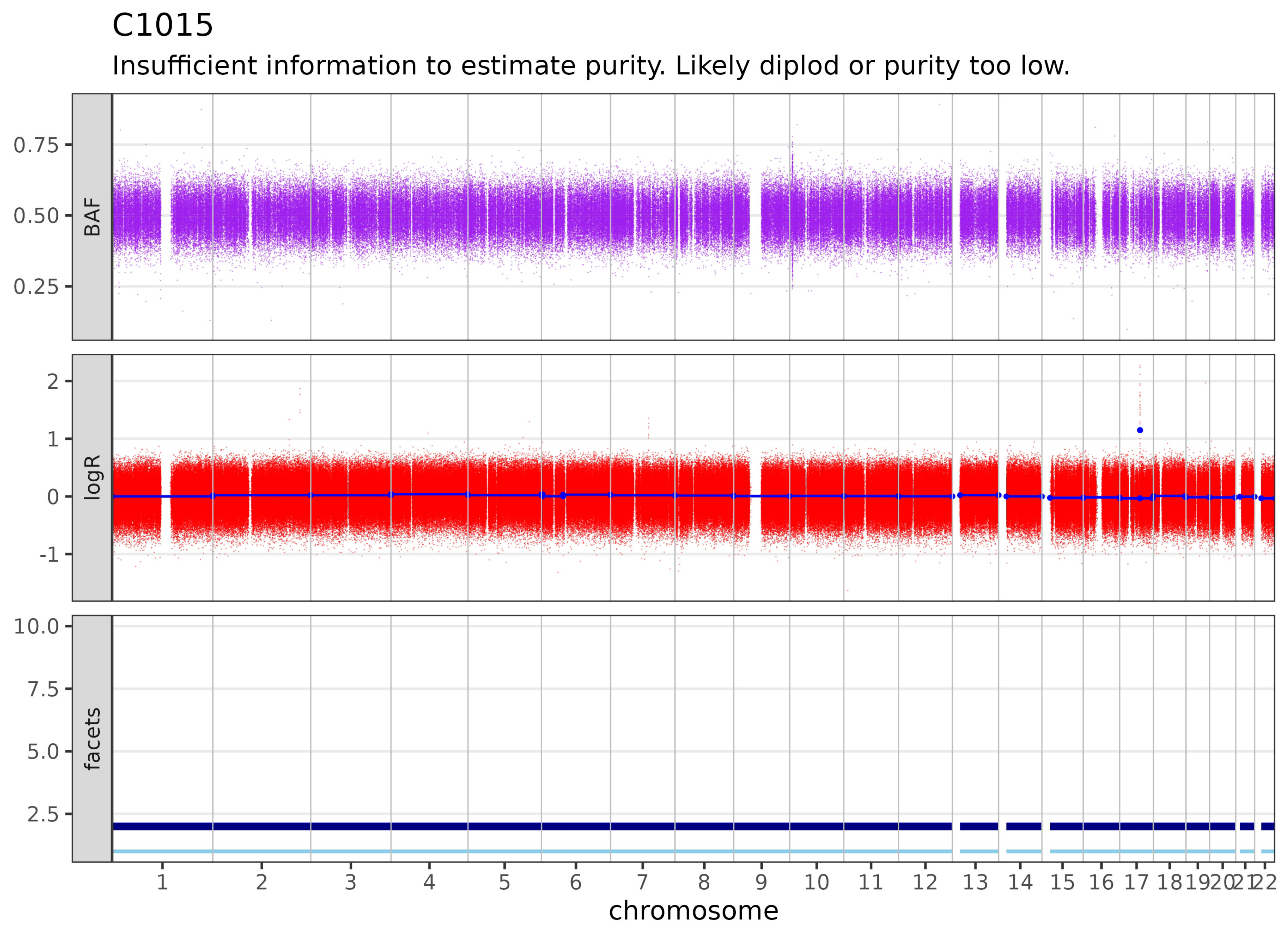
C1015
Insufficient information to estimate purity. Likely diplod or purity too low.
BAF
0.75
0.50
0.25
logR
2
1
0
-1
facets
10.0
7.5
5.0
2.5
1 2 3 4 5 6 7 8 9 10 11 12 13 14 15 16 17 18 19 20 21 22
chromosome

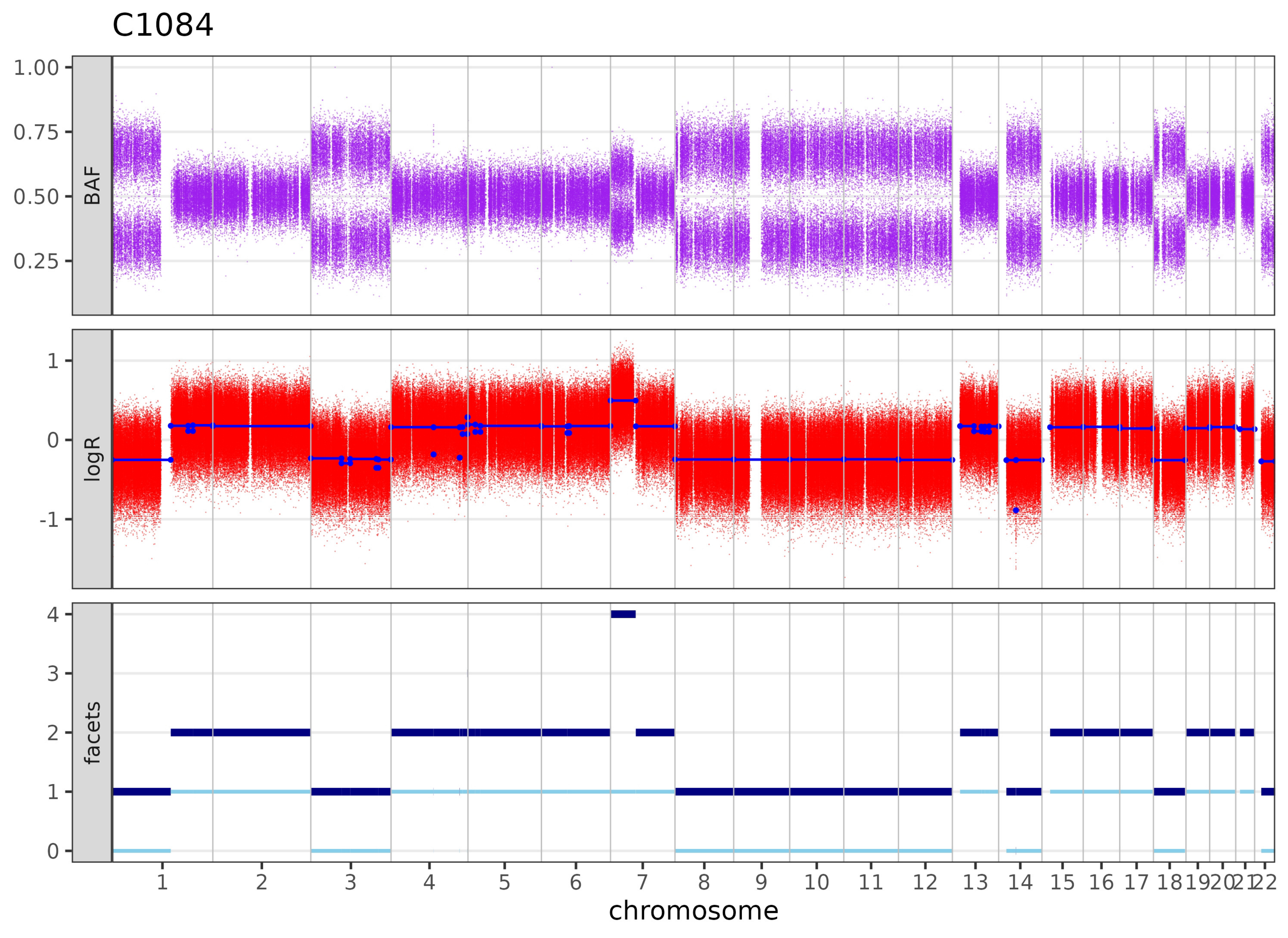

C1084
BAF
logR
facets
1.00
0.75
0.50
0.25
1
0
-1
4
3
2
1
chromosome

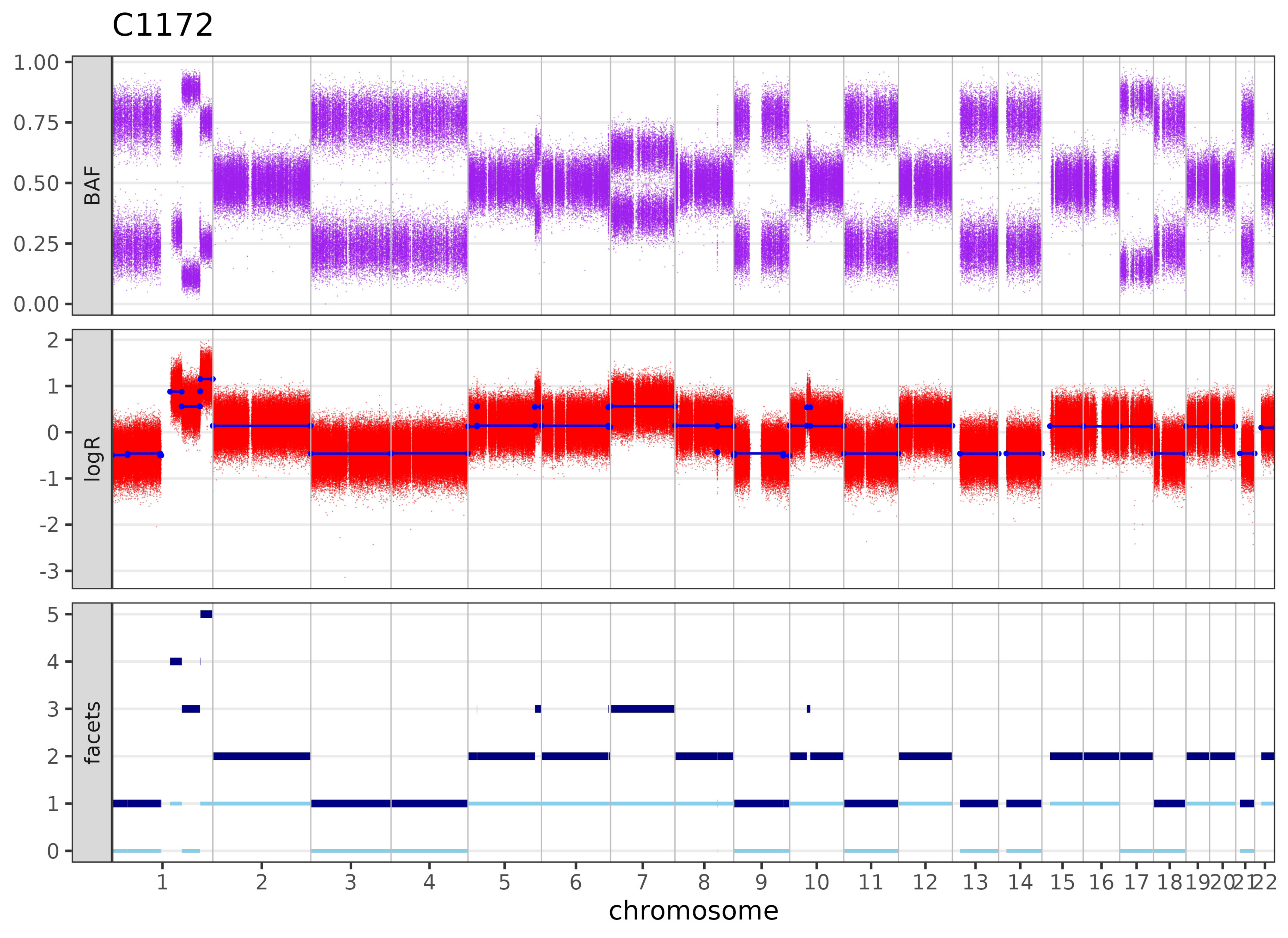

C1172
BAF
1.00
0.75
0.50
0.25
0.00
logR
2
1
0
-1
-2
-3
facets
5
4
3
2
1
0
1 2 3 4 5 6 7 8 9 10 11 12 13 14 15 16 17 18 19 20 21 22
chromosome

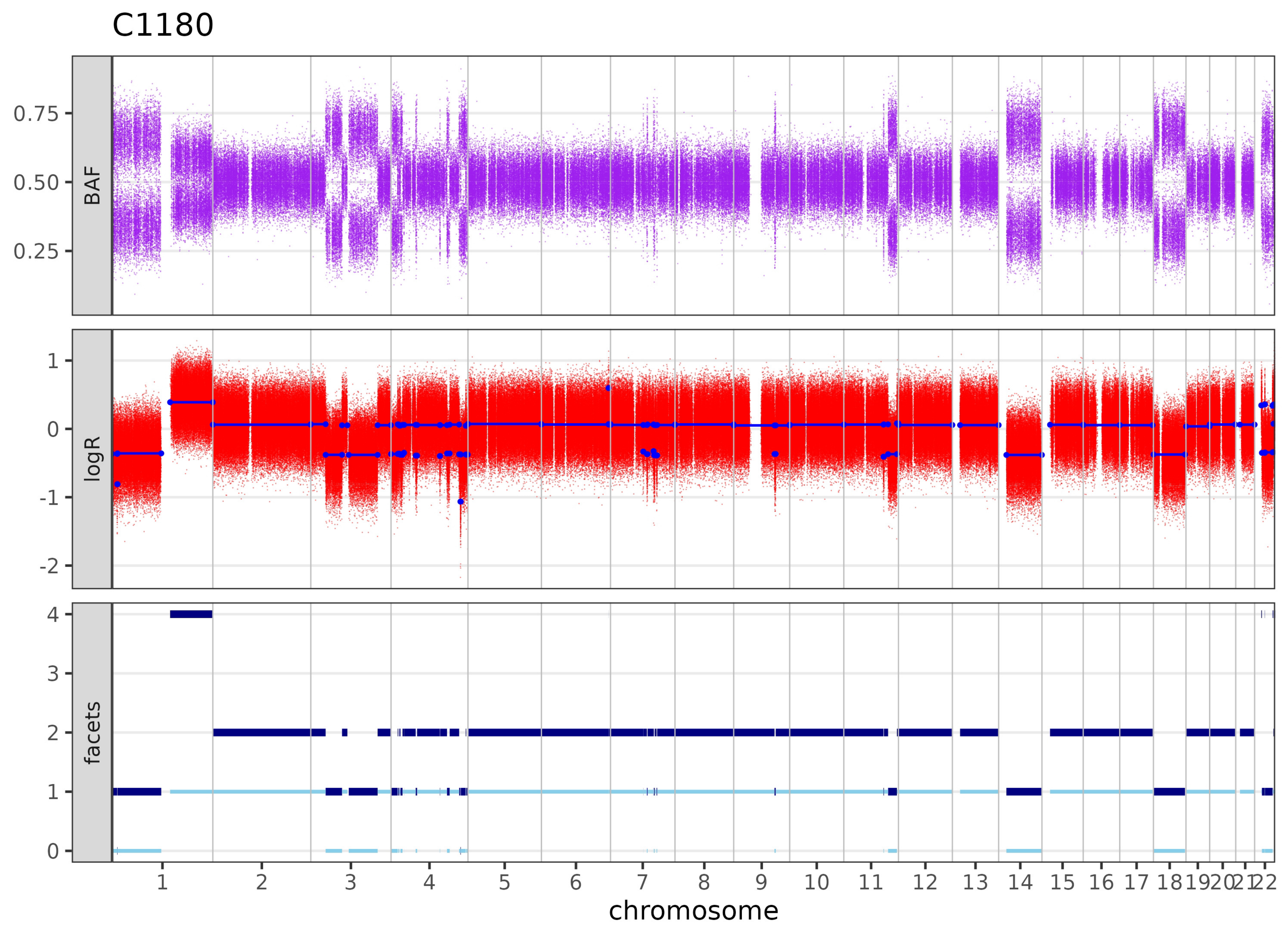

C1180
BAF
0.75
0.50
0.25
logR
1
0
-1
-2
facets
4
3
2
1
chromosome

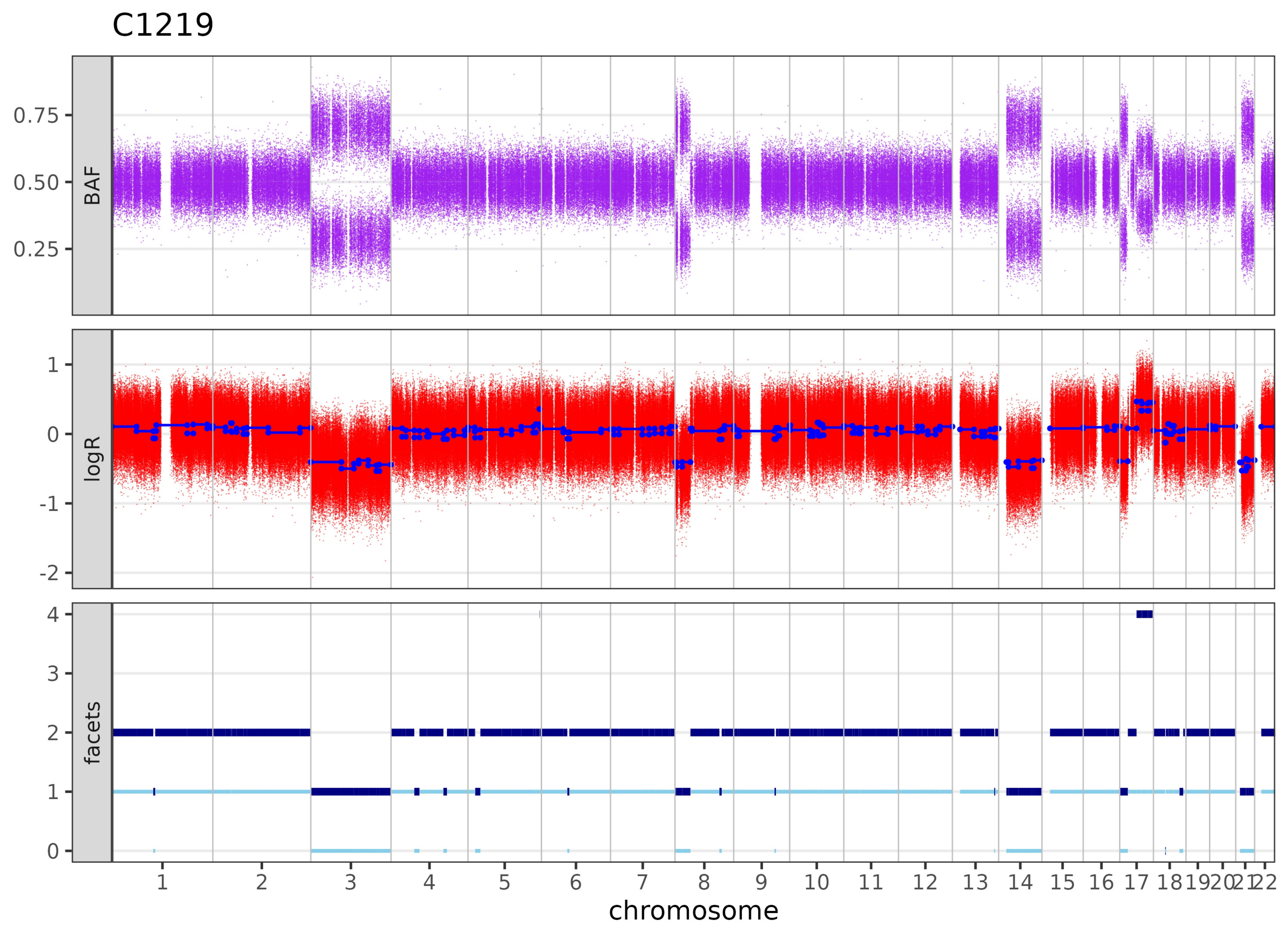
C1219
BAF
0.75
0.50
0.25
logR
1
0
-1
-2
facets
4
3
2
1
chromosome

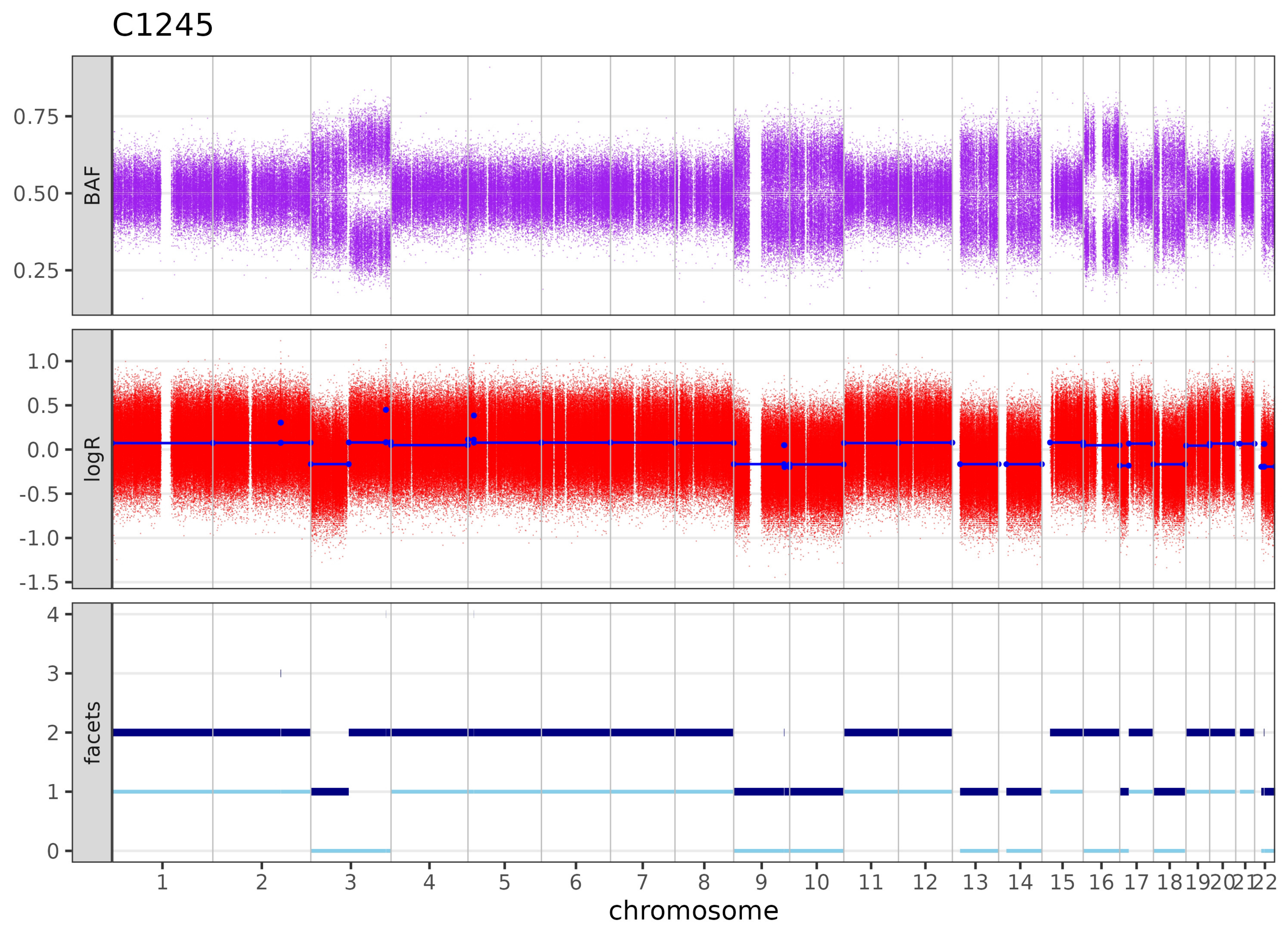
C1245
BAF
0.75
0.50
0.25
logR
1.0
0.5
0.0
-0.5
-1.0
-1.5
facets
4
3
2
1
chromosome

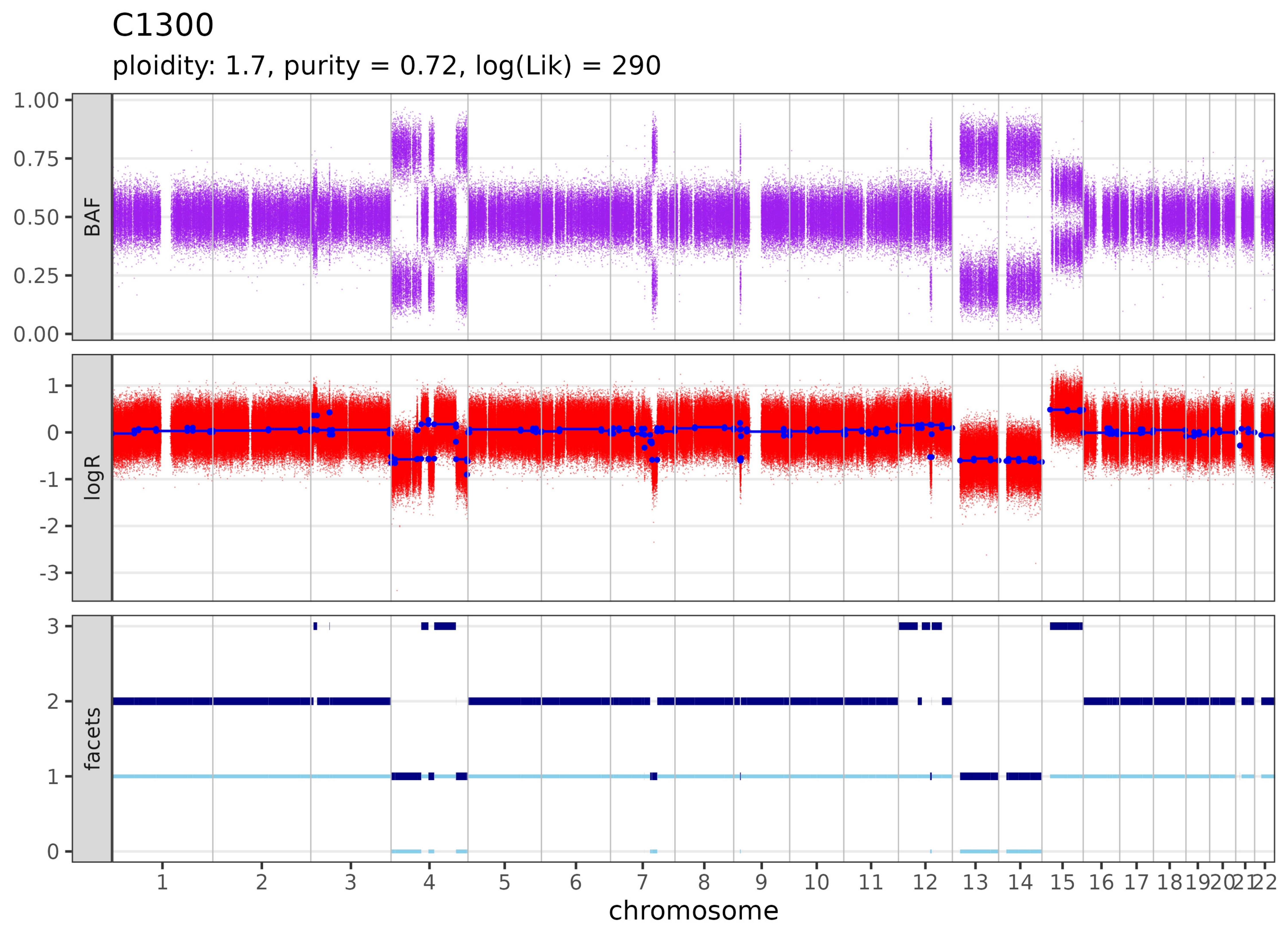

C1300
ploidity: 1.7, purity = 0.72, log(Lik) = 290
BAF
1.00
0.75
0.50
0.25
0.00
logR
1
0
-1
-2
-3
facets
3
2
1
0
1 2 3 4 5 6 7 8 9 10 11 12 13 14 15 16 17 18 19 20 21 22
chromosome

# C1303

Insufficient information to estimate purity. Likely diplod or purity too low.

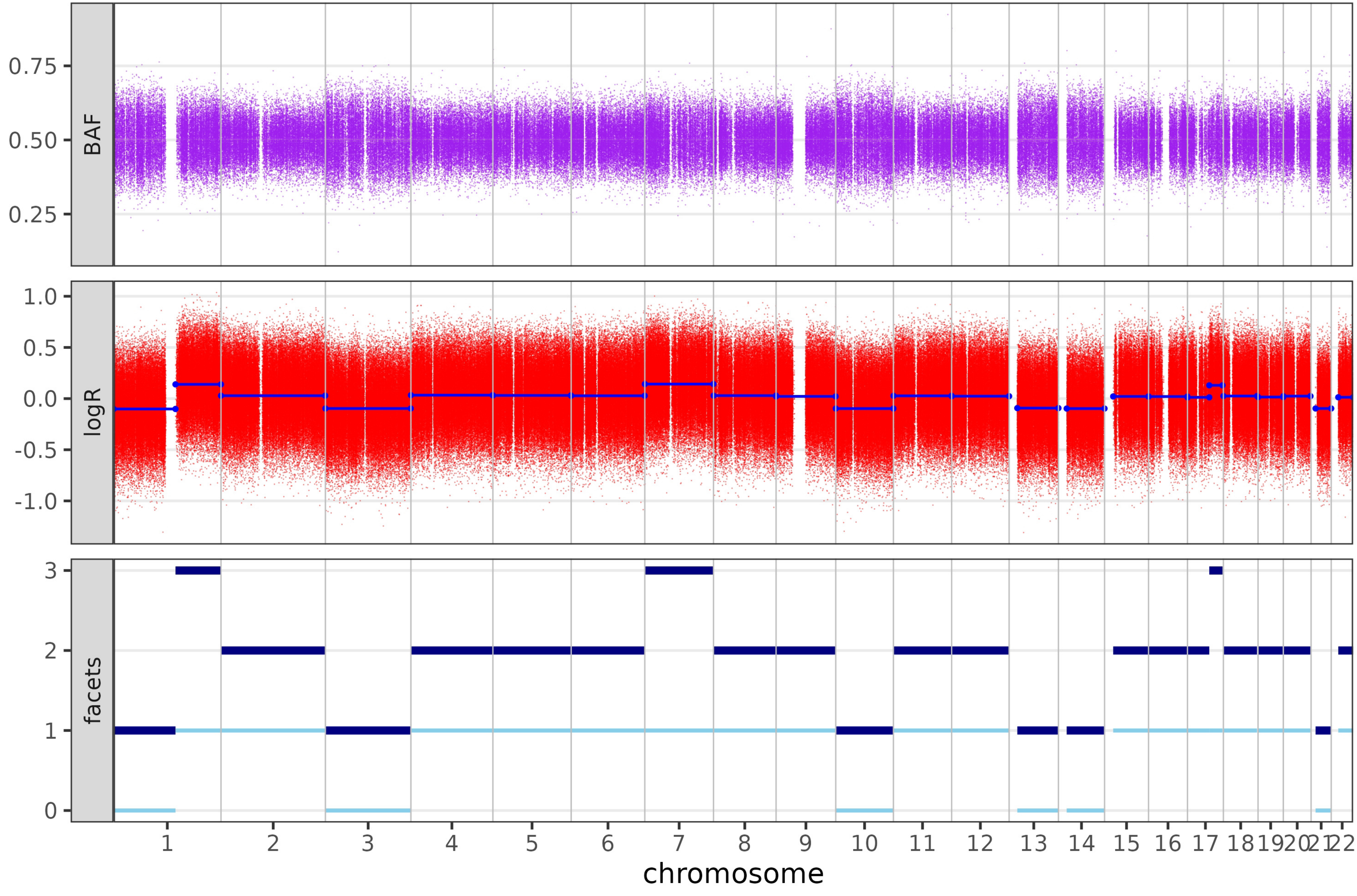

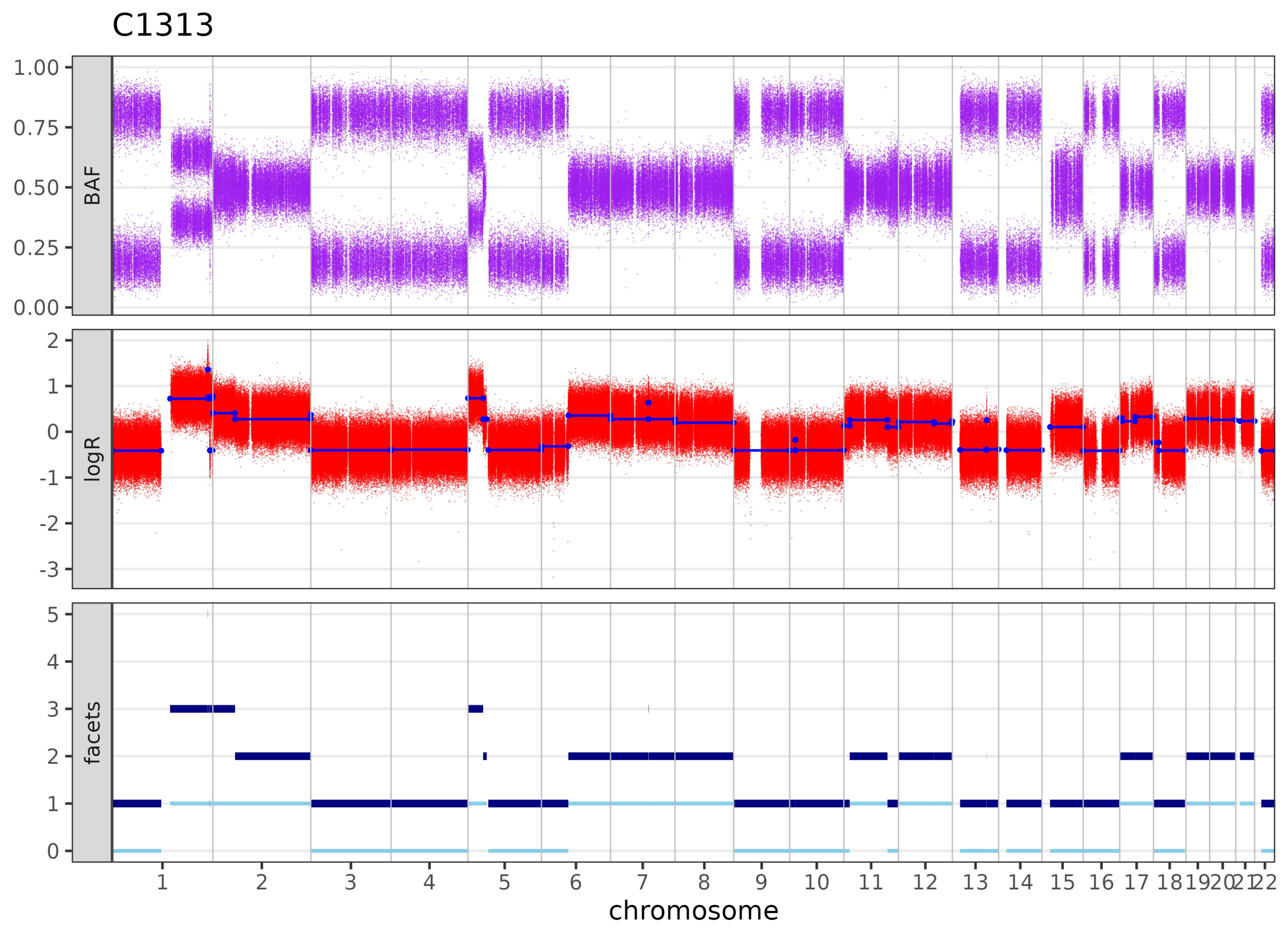
C1313
BAF
logR
facets
1.00
0.75
0.50
0.25
0.00
2
1
0
-1
-2
-3
5
4
3
2
1
0
1 2 3 4 5 6 7 8 9 10 11 12 13 14 15 16 17 18 19 20 21 22
chromosome

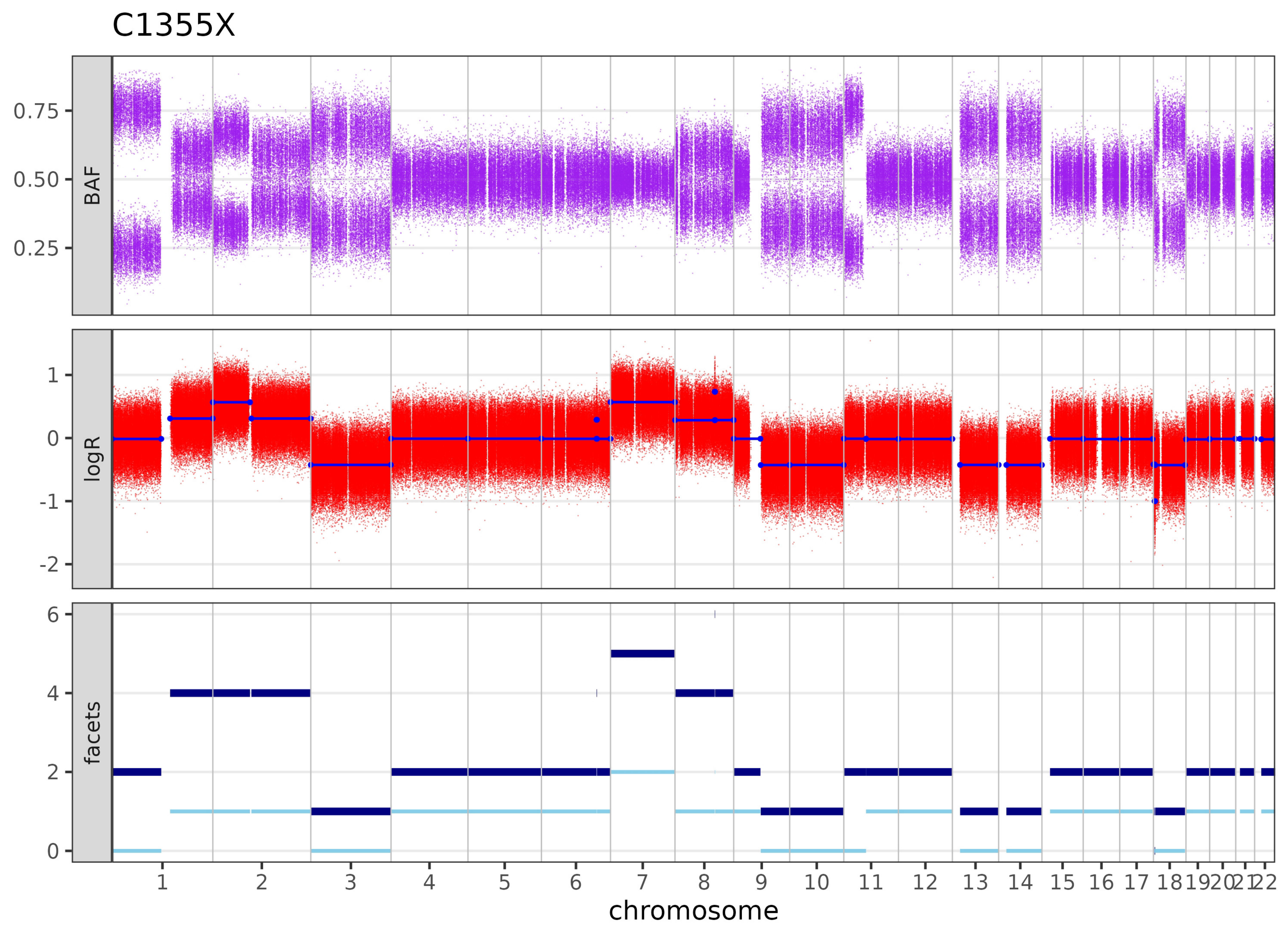

C1355X
BAF
0.75
0.50
0.25
logR
1
0
-1
-2
facets
6
4
2
0
1 2 3 4 5 6 7 8 9 10 11 12 13 14 15 16 17 18 19 20 21 22
chromosome

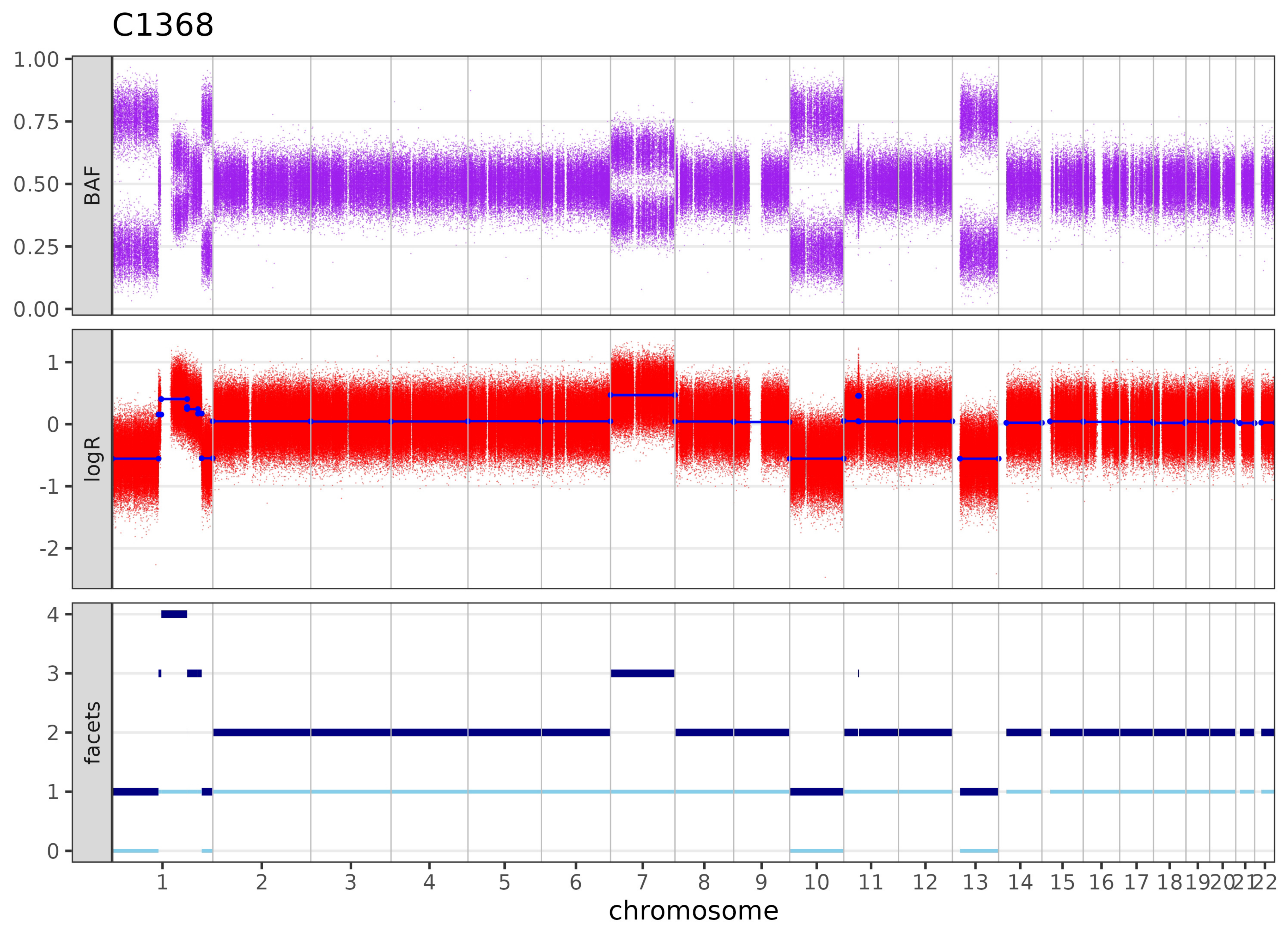

C1368
BAF
logR
facets
1.00
0.75
0.50
0.25
0.00
1
0
-1
-2
4
3
2
1
0
1 2 3 4 5 6 7 8 9 10 11 12 13 14 15 16 17 18 19 20 21 22
chromosome

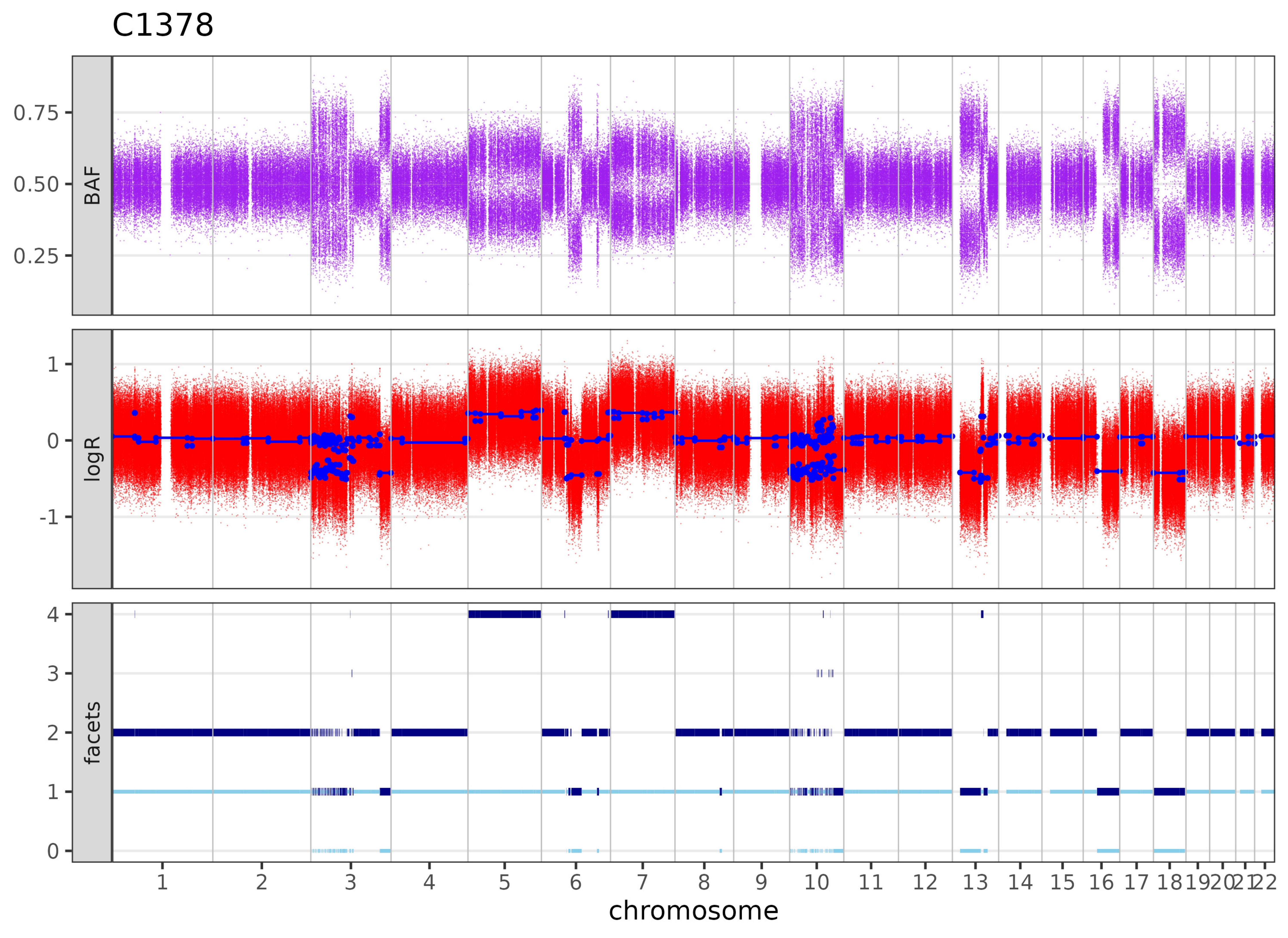

C1378
BAF
0.75
0.50
0.25
logR
1
0
-1
facets
4
3
2
1
chromosome

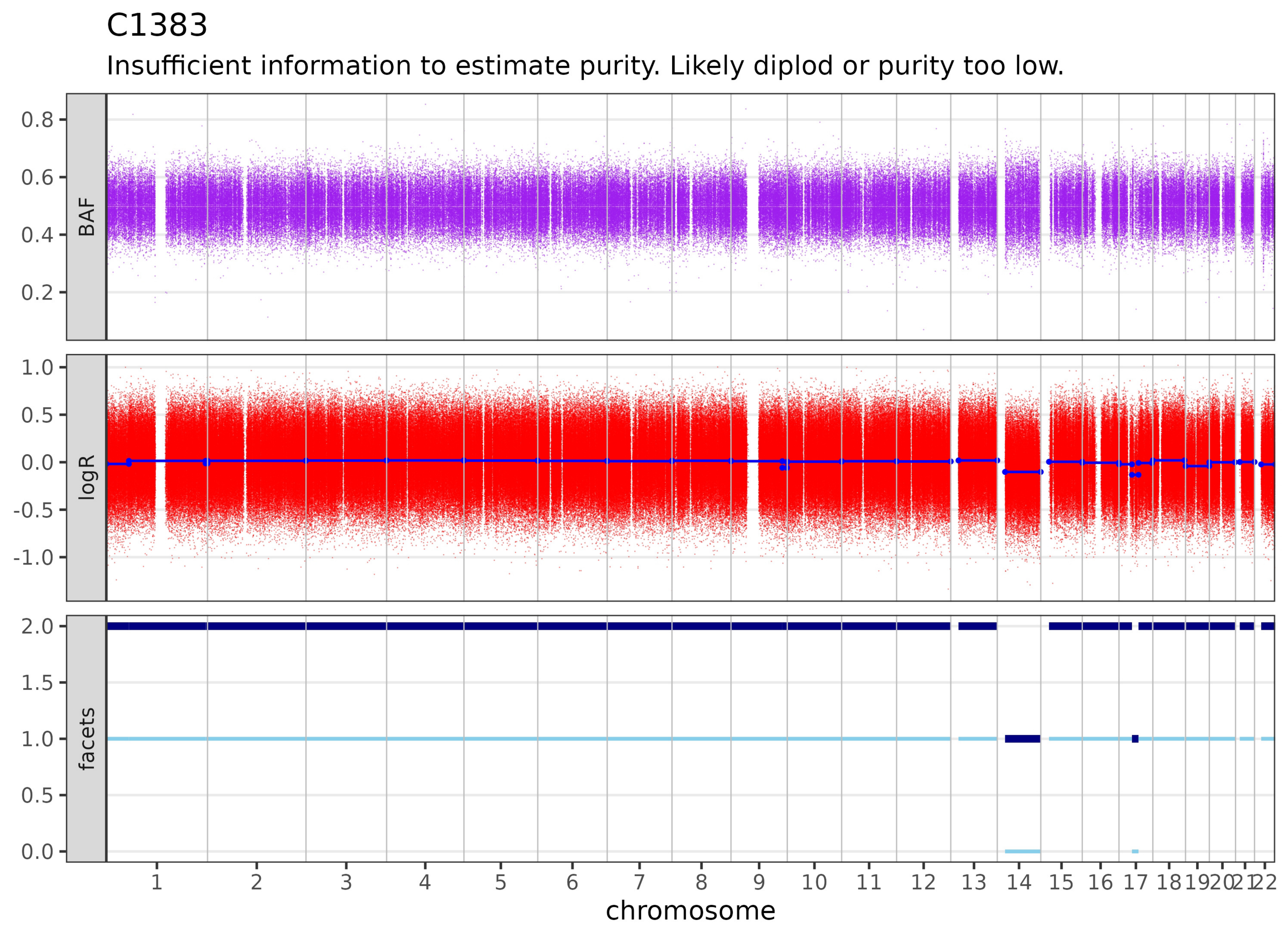
C1383
Insufficient information to estimate purity. Likely diplod or purity too low.
BAF
0.8
0.6
0.4
0.2
logR
1.0
0.5
0.0
-0.5
-1.0
facets
2.0
1.5
1.0
0.5
0.0
chromosome

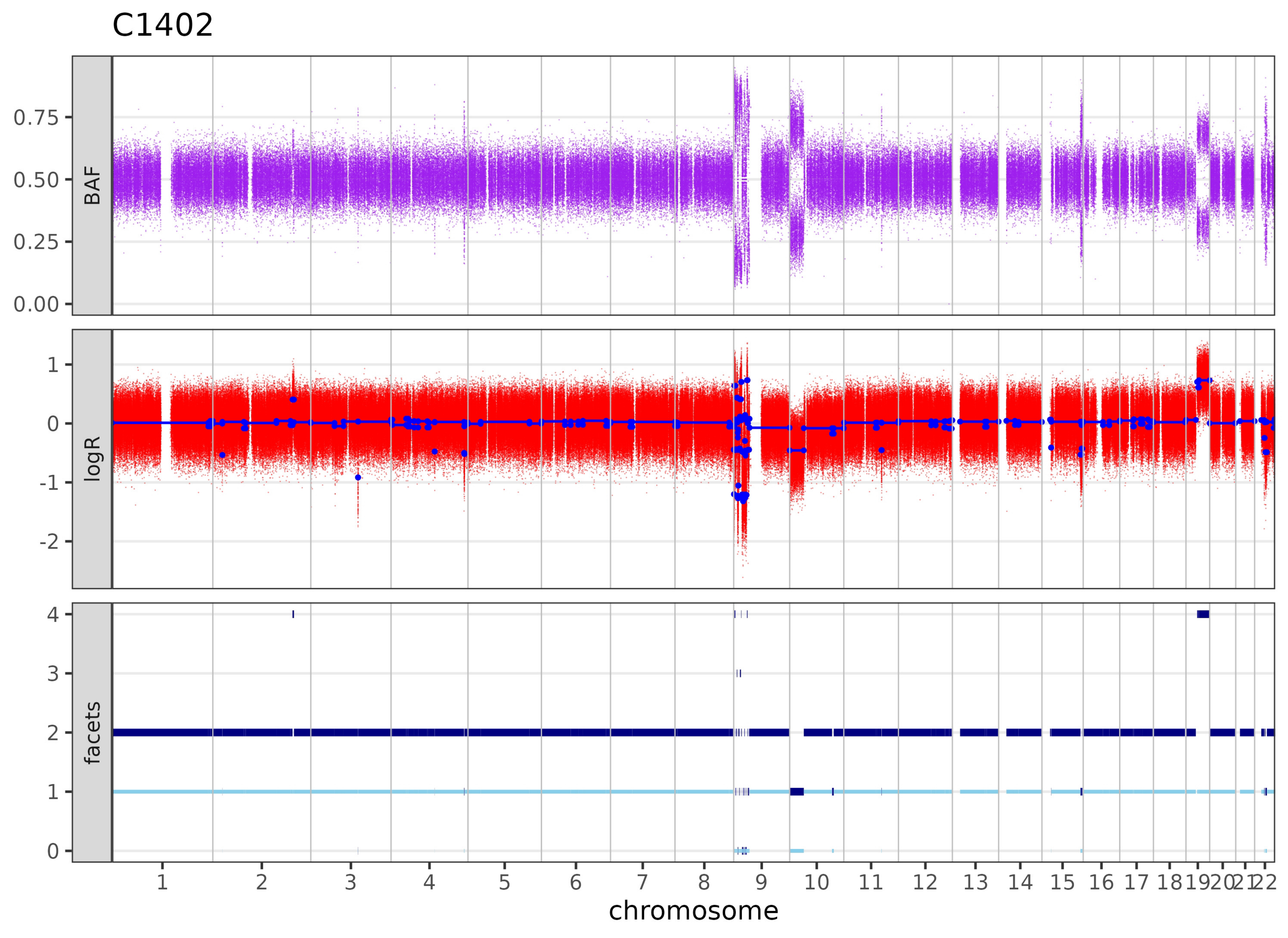

C1402
BAF
0.75
0.50
0.25
0.00
logR
1
0
-1
-2
facets
4
3
2
1
chromosome

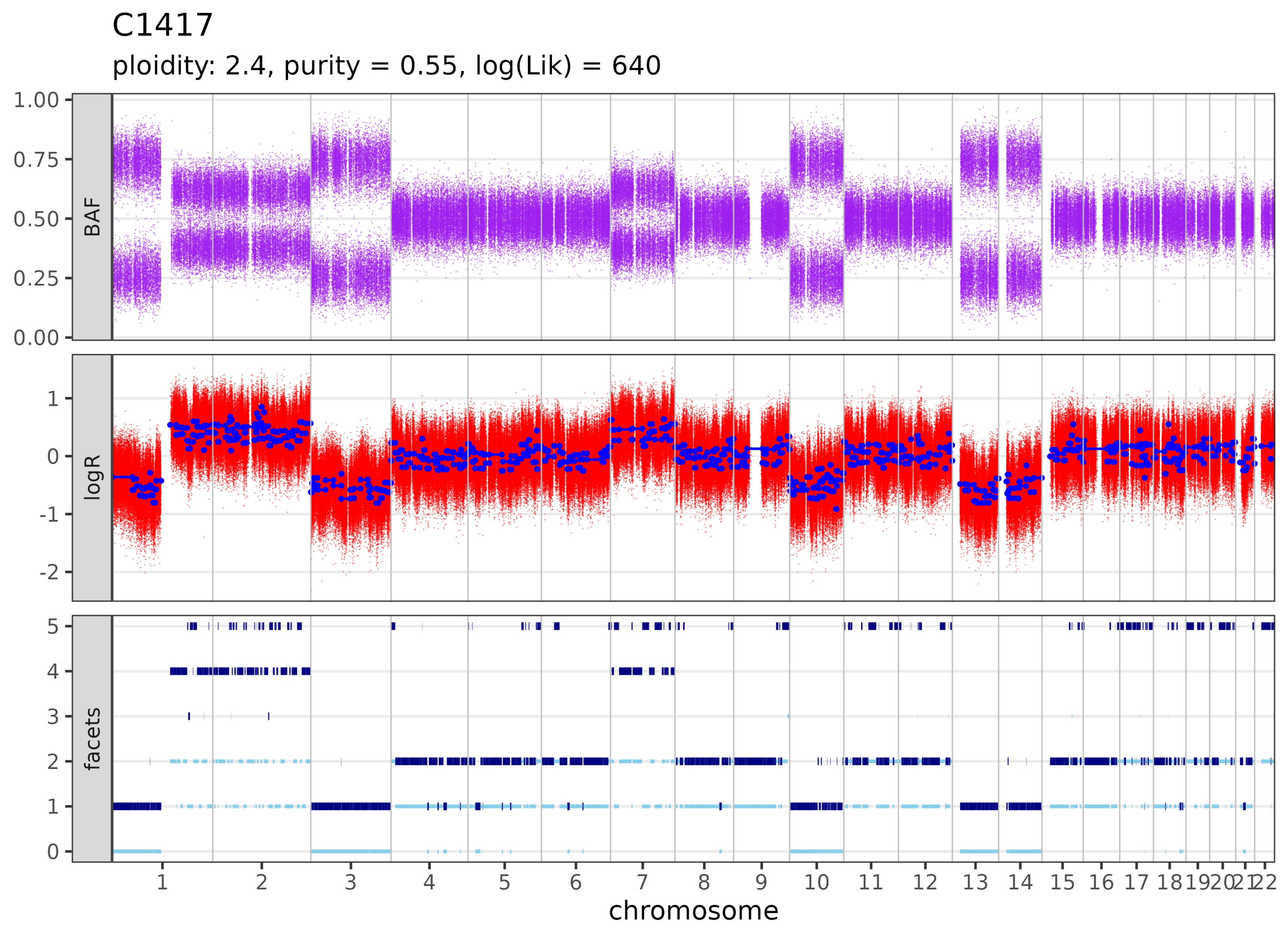

C1417
ploidity: 2.4, purity = 0.55, log(Lik) = 640
BAF
1.00
0.75
0.50
0.25
0.00
logR
1
0
-1
-2
facets
5
4
3
2
1
0
1 2 3 4 5 6 7 8 9 10 11 12 13 14 15 16 17 18 19 20 21 22
chromosome

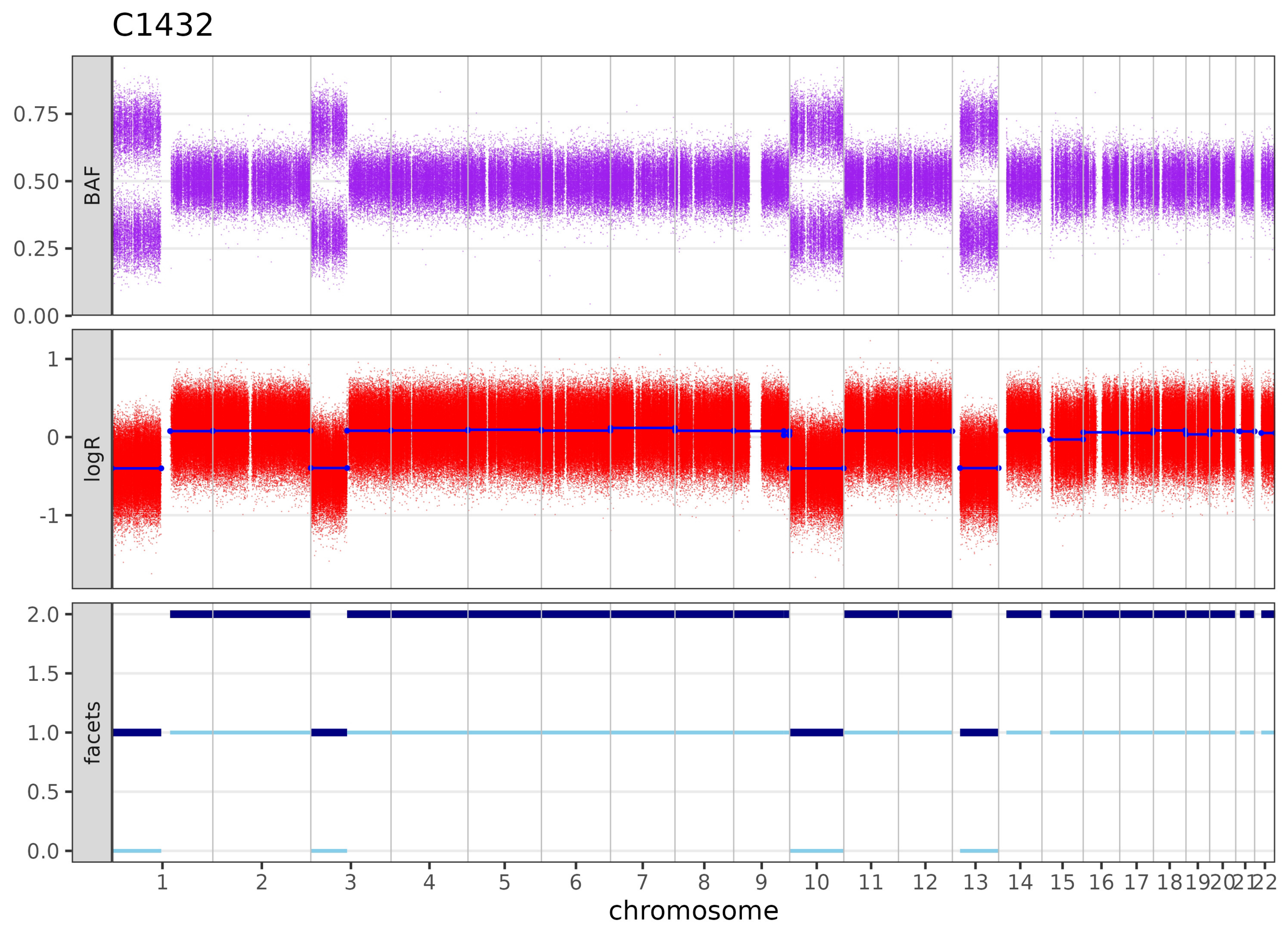

C1432
BAF
logR
facets
0.00
0.25
0.50
0.75
1
0
-1
0.0
0.5
1.0
1.5
2.0
1 2 3 4 5 6 7 8 9 10 11 12 13 14 15 16 17 18 19 20 21 22
chromosome

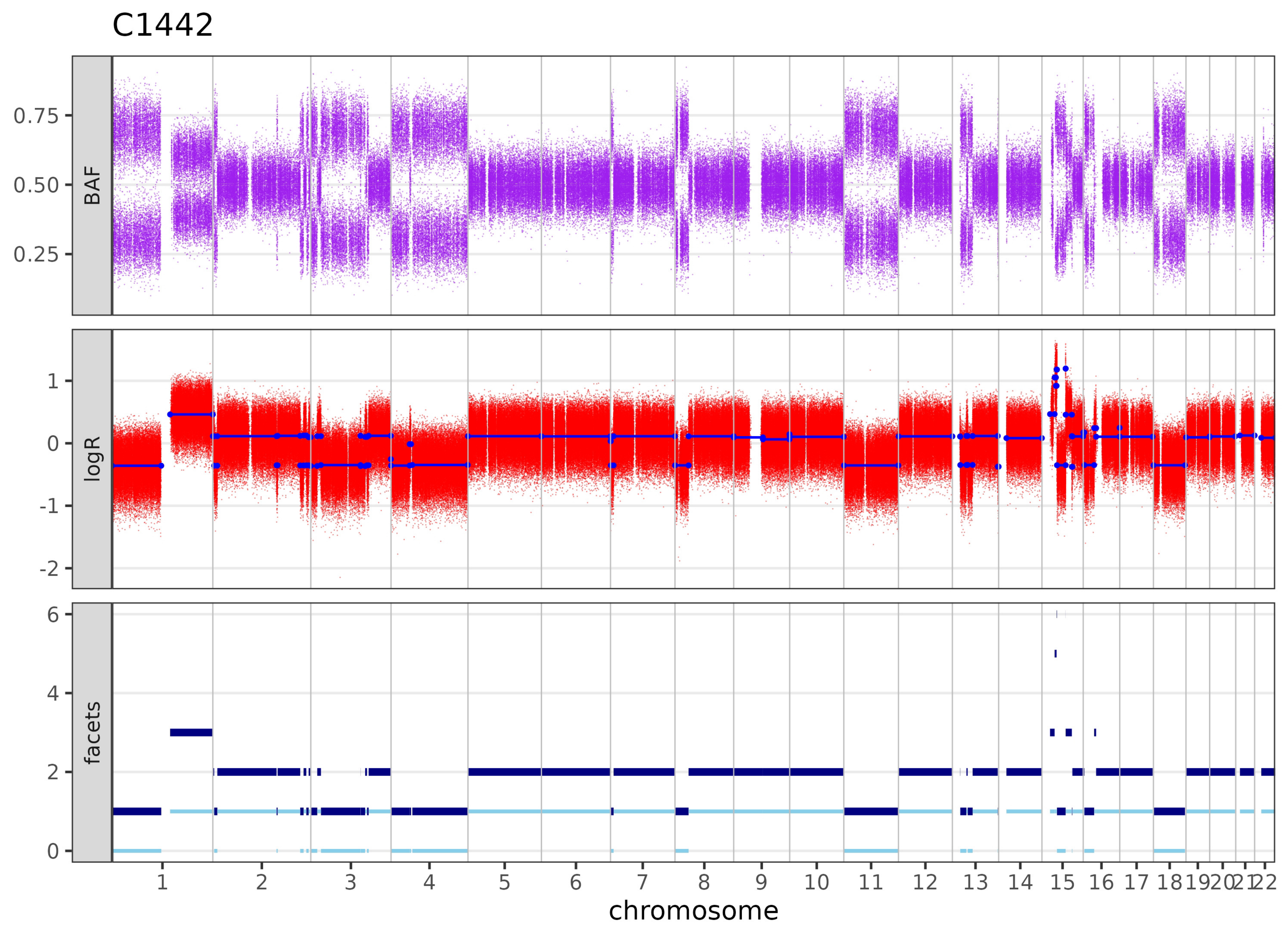

C1442
BAF
0.75
0.50
0.25
logR
1
0
-1
-2
facets
6
4
2
0
1 2 3 4 5 6 7 8 9 10 11 12 13 14 15 16 17 18 19 20 21 22
chromosome

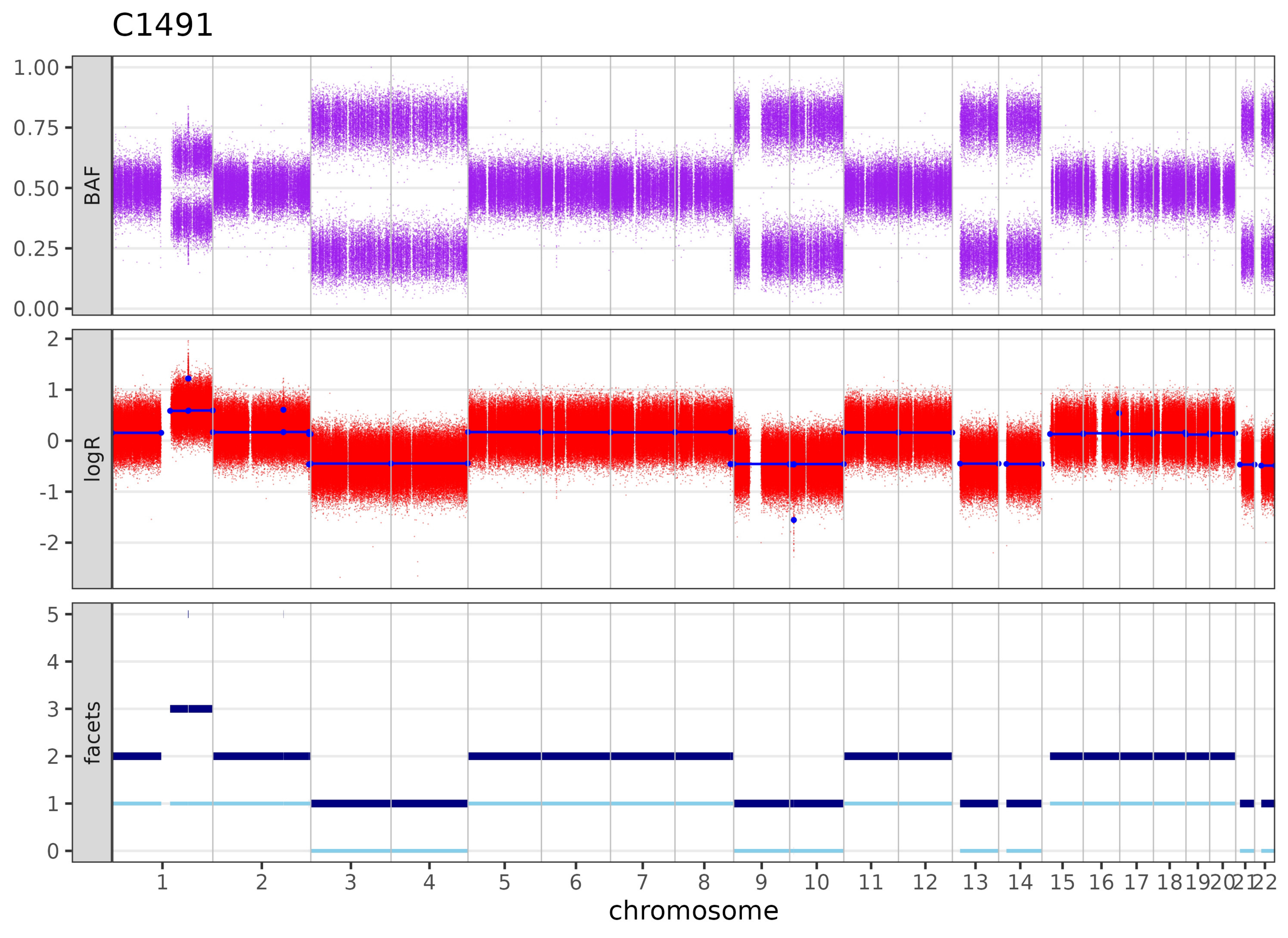

C1491
BAF
logR
facets
1.00
0.75
0.50
0.25
0.00
2
1
0
-1
-2
5
4
3
2
1
0
1 2 3 4 5 6 7 8 9 10 11 12 13 14 15 16 17 18 19 20 21 22
chromosome

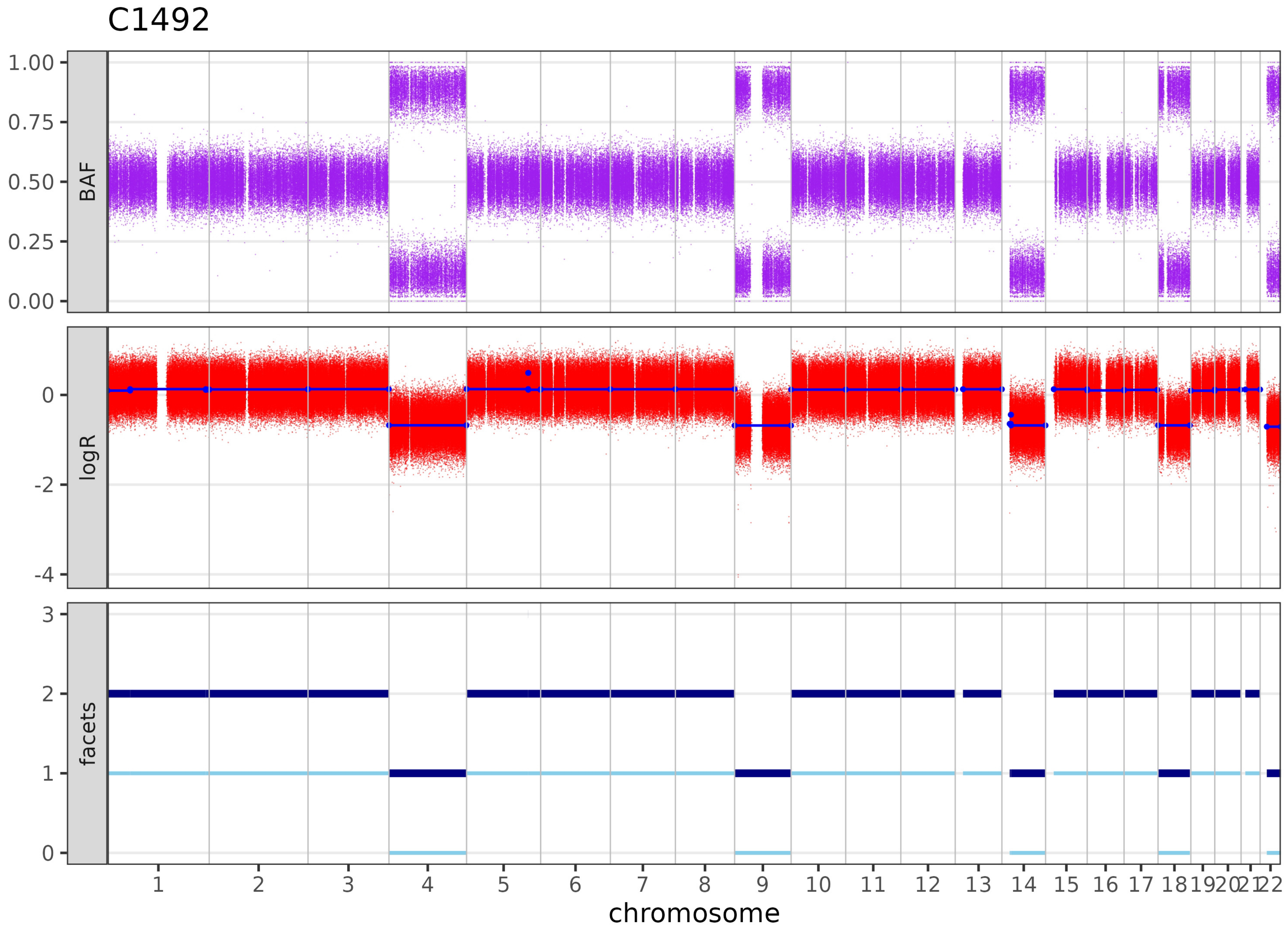
C1492
BAF
1.00
0.75
0.50
0.25
0.00
logR
0
-2
-4
facets
3
2
1
0
1 2 3 4 5 6 7 8 9 10 11 12 13 14 15 16 17 18 19 20 21 22
chromosome

C1537

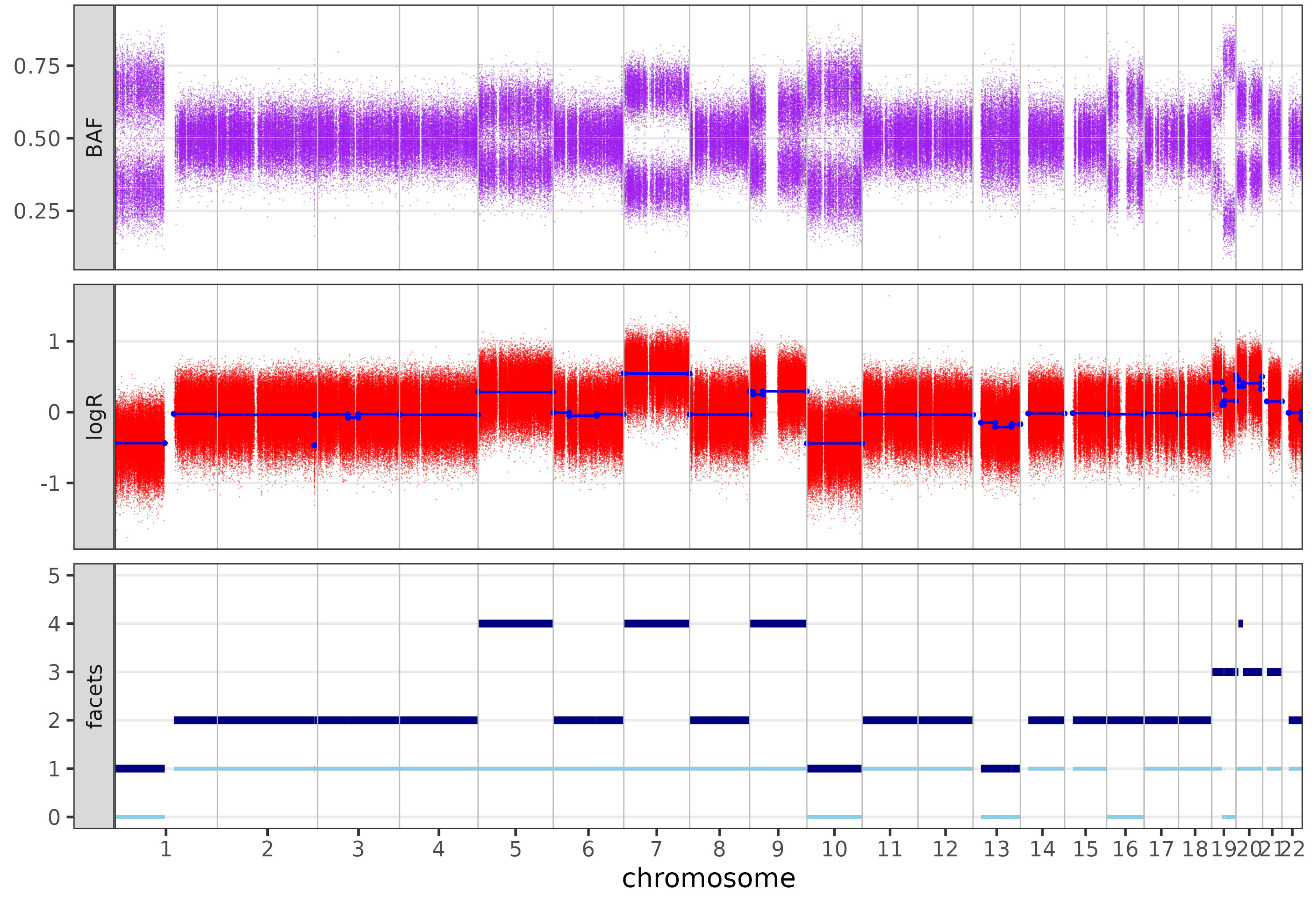

BAF
0.75
0.50
0.25
logR
1
0
-1
facets
5
4
3
2
1
chromosome

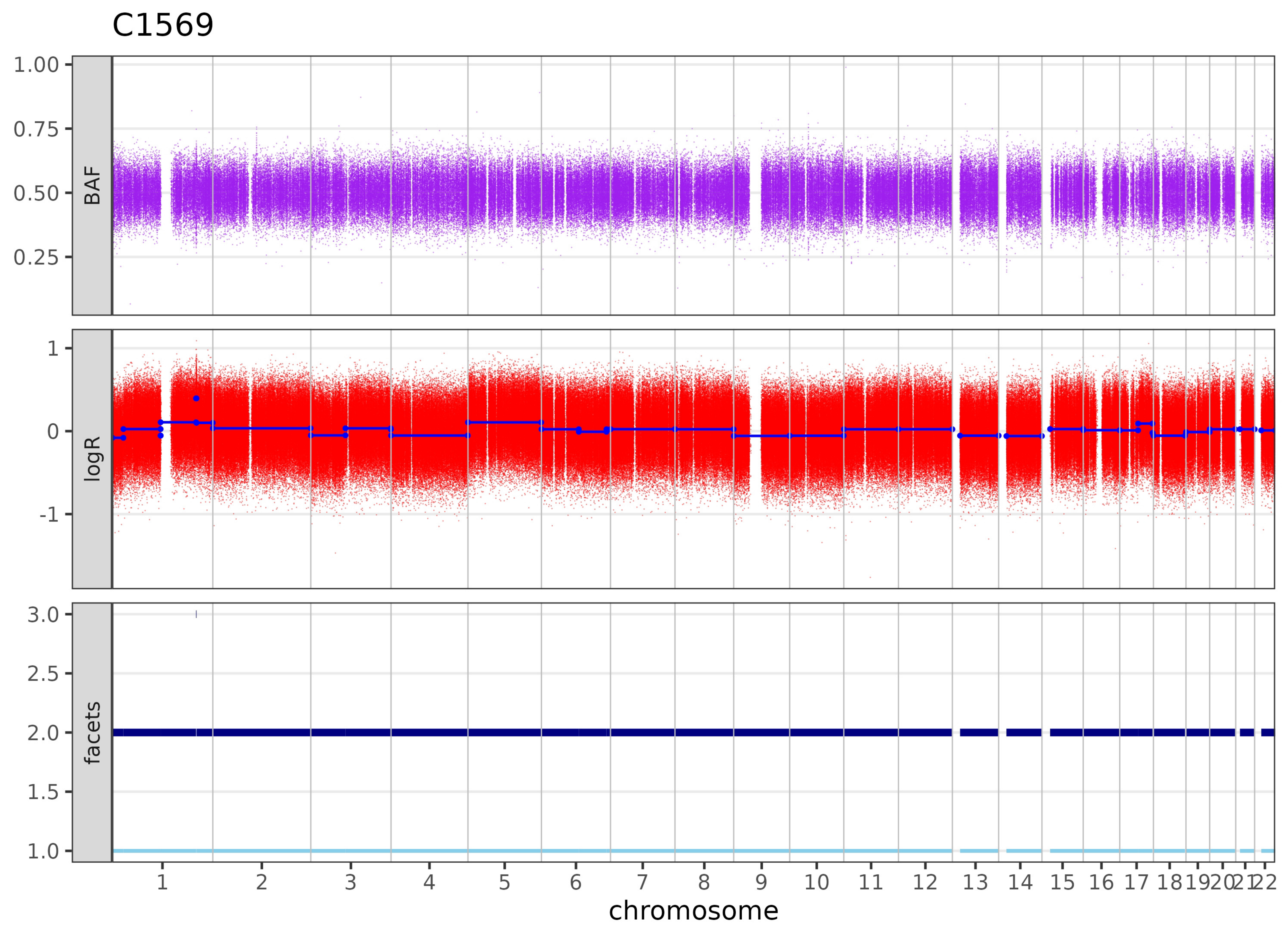

C1569
BAF
logR
facets
1.00
0.75
0.50
0.25
1
0
-1
3.0
2.5
2.0
1.5
1.0
1 2 3 4 5 6 7 8 9 10 11 12 13 14 15 16 17 18 19 20 21 22
chromosome

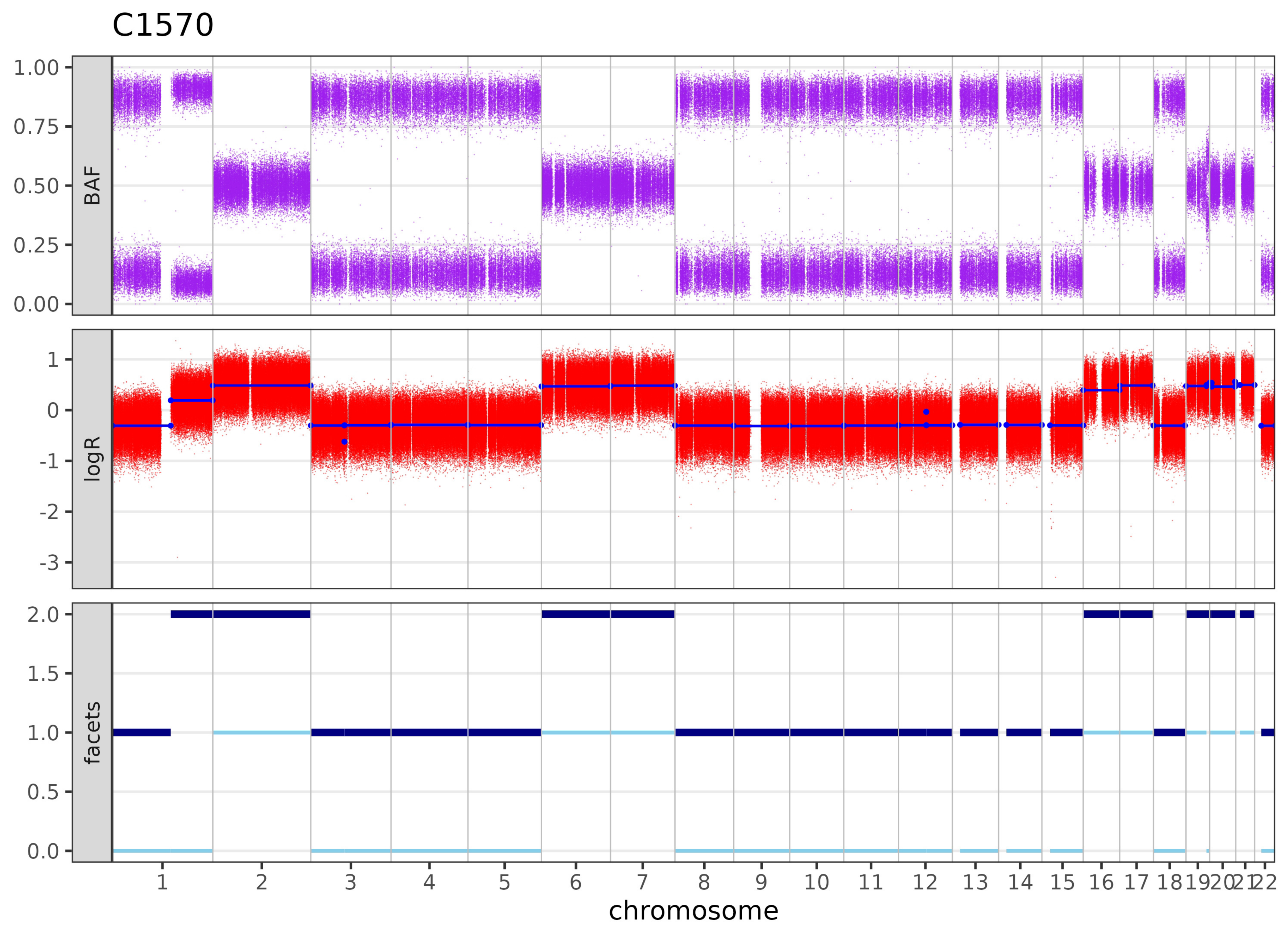

C1570
BAF
logR
facets
1.00
0.75
0.50
0.25
0.00
1
0
-1
-2
-3
2.0
1.5
1.0
0.5
0.0
1 2 3 4 5 6 7 8 9 10 11 12 13 14 15 16 17 18 19 20 21 22
chromosome

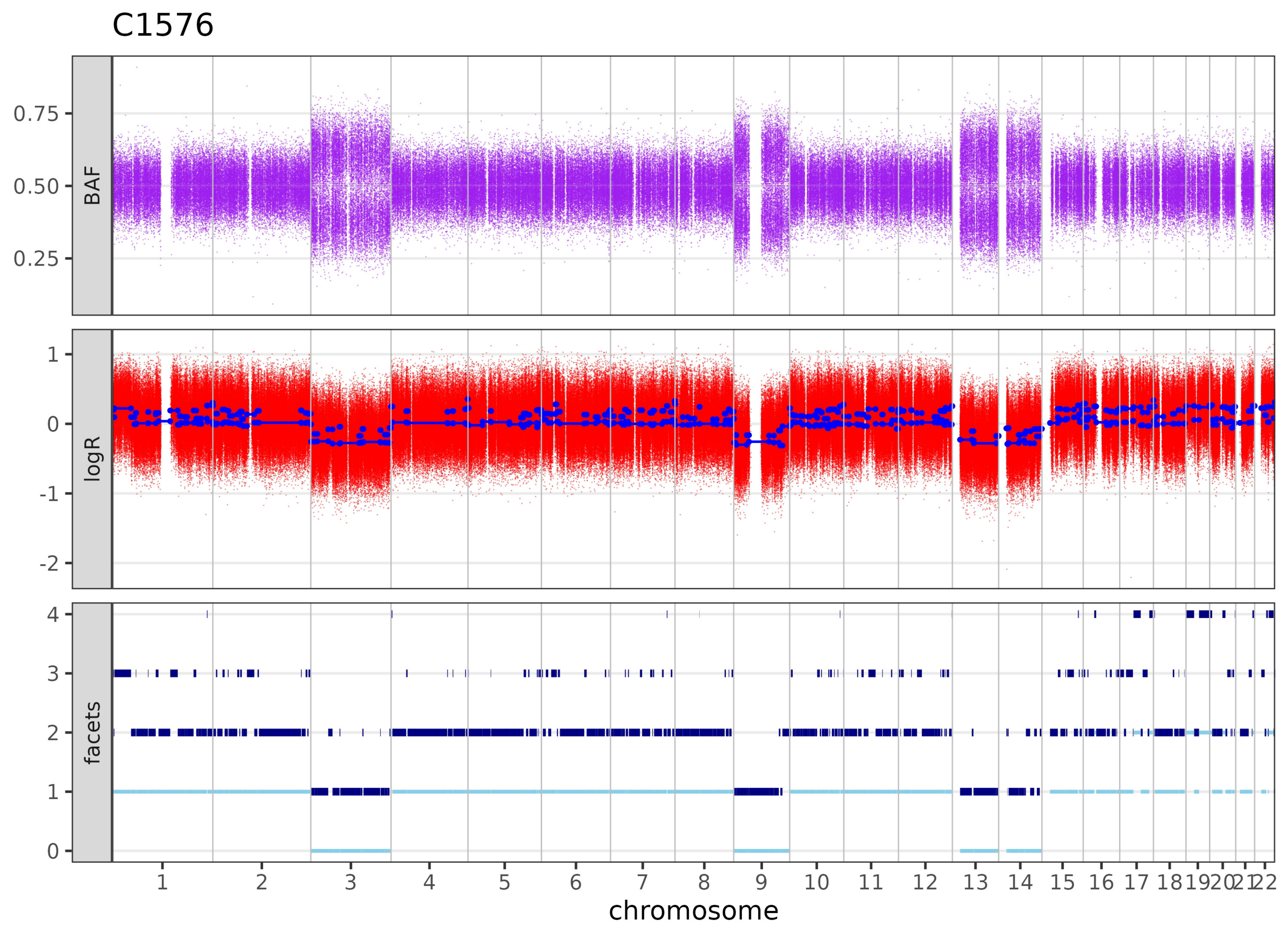

C1576
BAF
logR
facets
0.75
0.50
0.25
1
0
-1
-2
4
3
2
1
0
1 2 3 4 5 6 7 8 9 10 11 12 13 14 15 16 17 18 19 20 21 22
chromosome

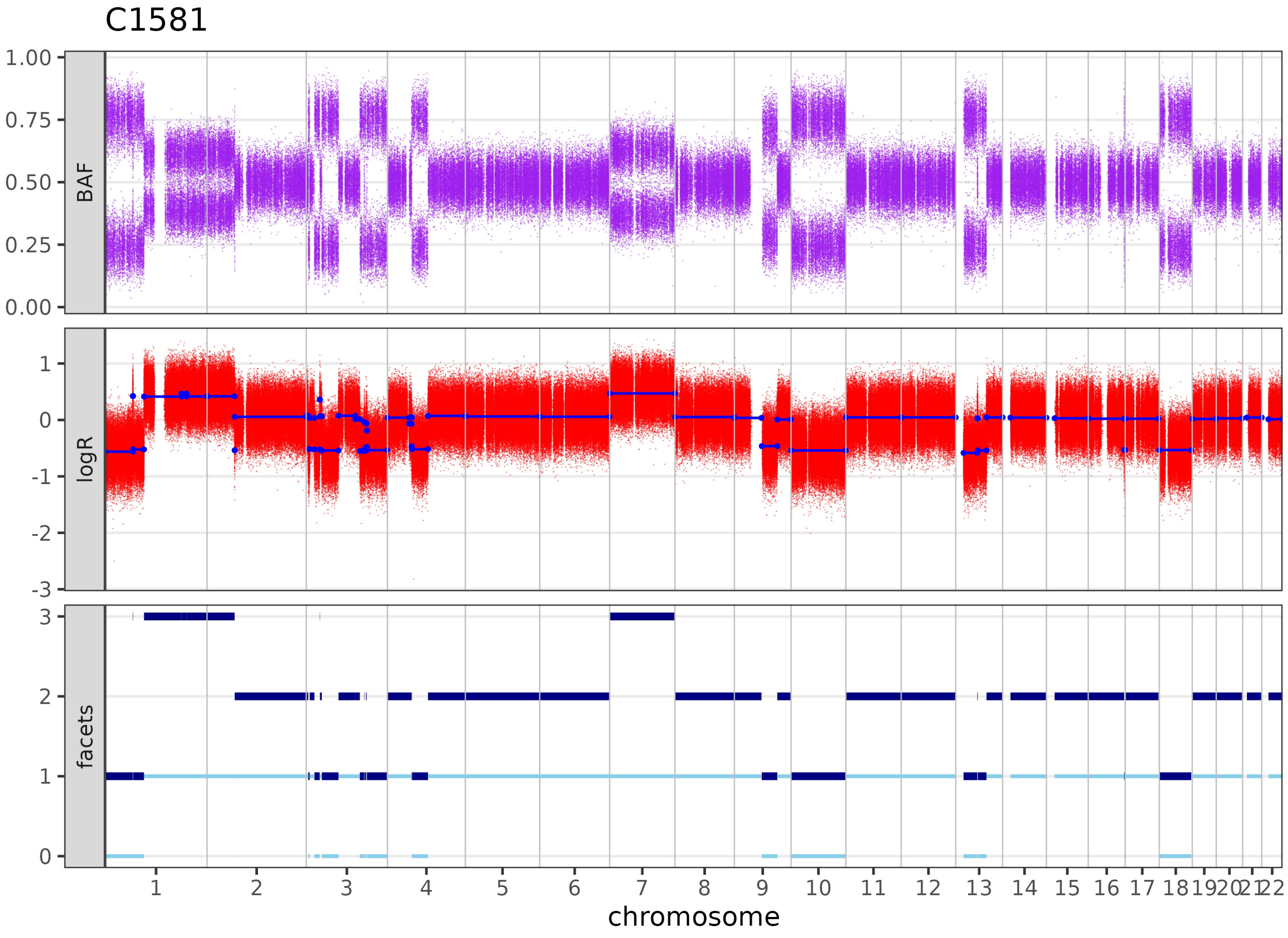

C1581
BAF
logR
facets
1.00
0.75
0.50
0.25
0.00
1
0
-1
-2
-3
3
2
1
0
1 2 3 4 5 6 7 8 9 10 11 12 13 14 15 16 17 18 19 20 21 22
chromosome

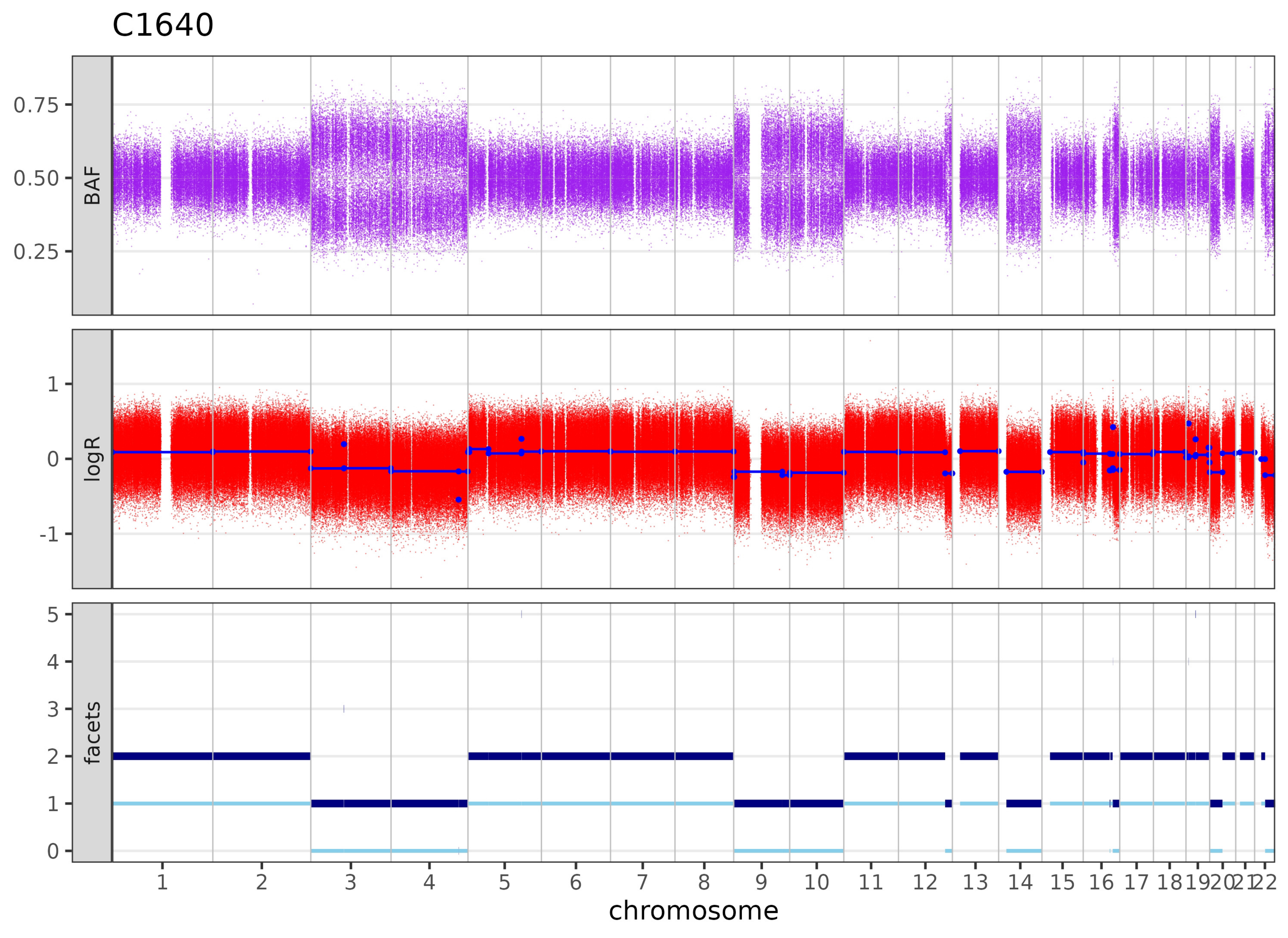
C1640
BAF
0.75
0.50
0.25
logR
1
0
-1
facets
5
4
3
2
1
chromosome

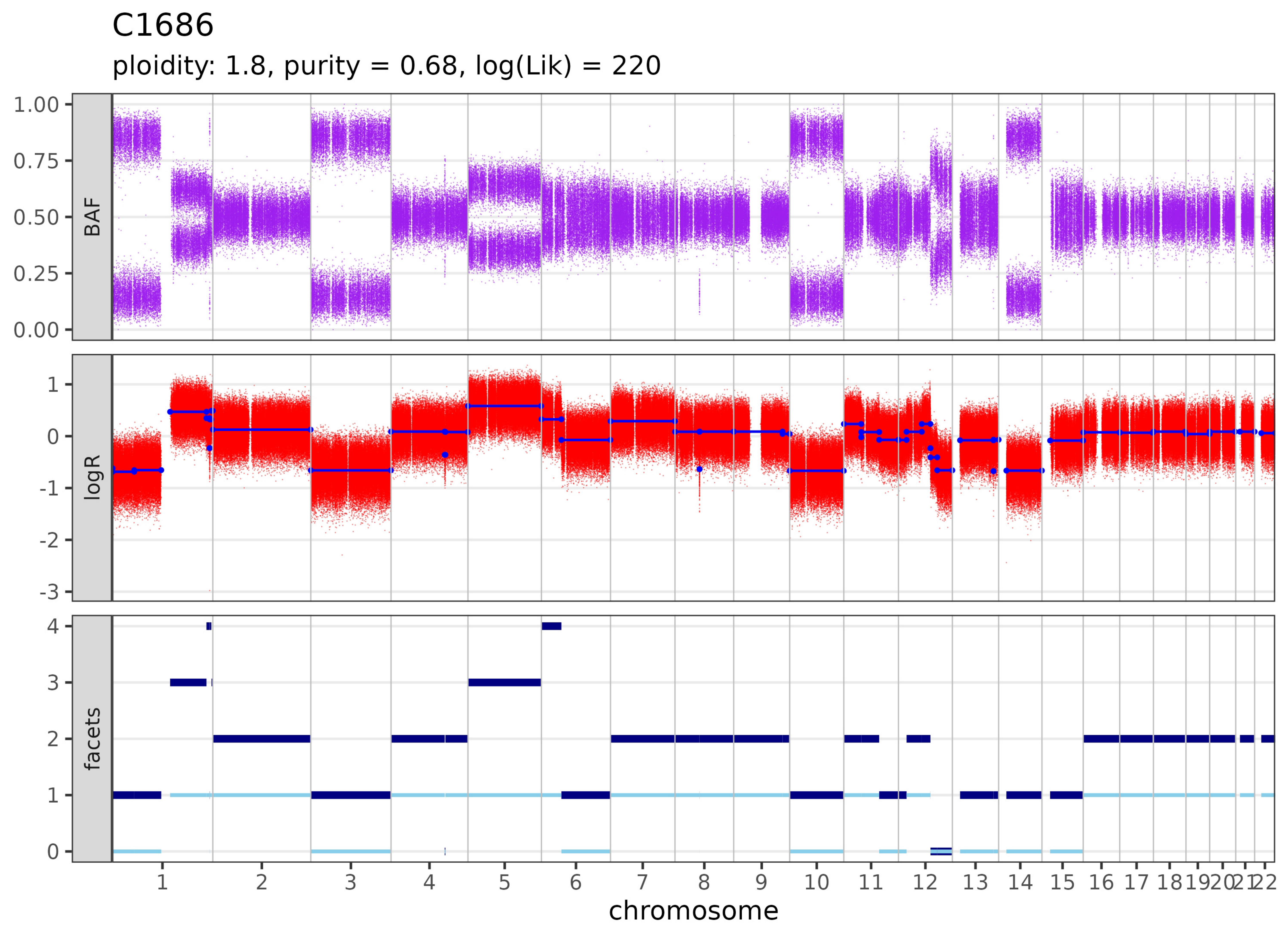

C1686
ploidity: 1.8, purity = 0.68, log(Lik) = 220
BAF
logR
facets
1.00
0.75
0.50
0.25
0.00
1
0
-1
-2
-3
4
3
2
1
0
1 2 3 4 5 6 7 8 9 10 11 12 13 14 15 16 17 18 19 20 21 22
chromosome

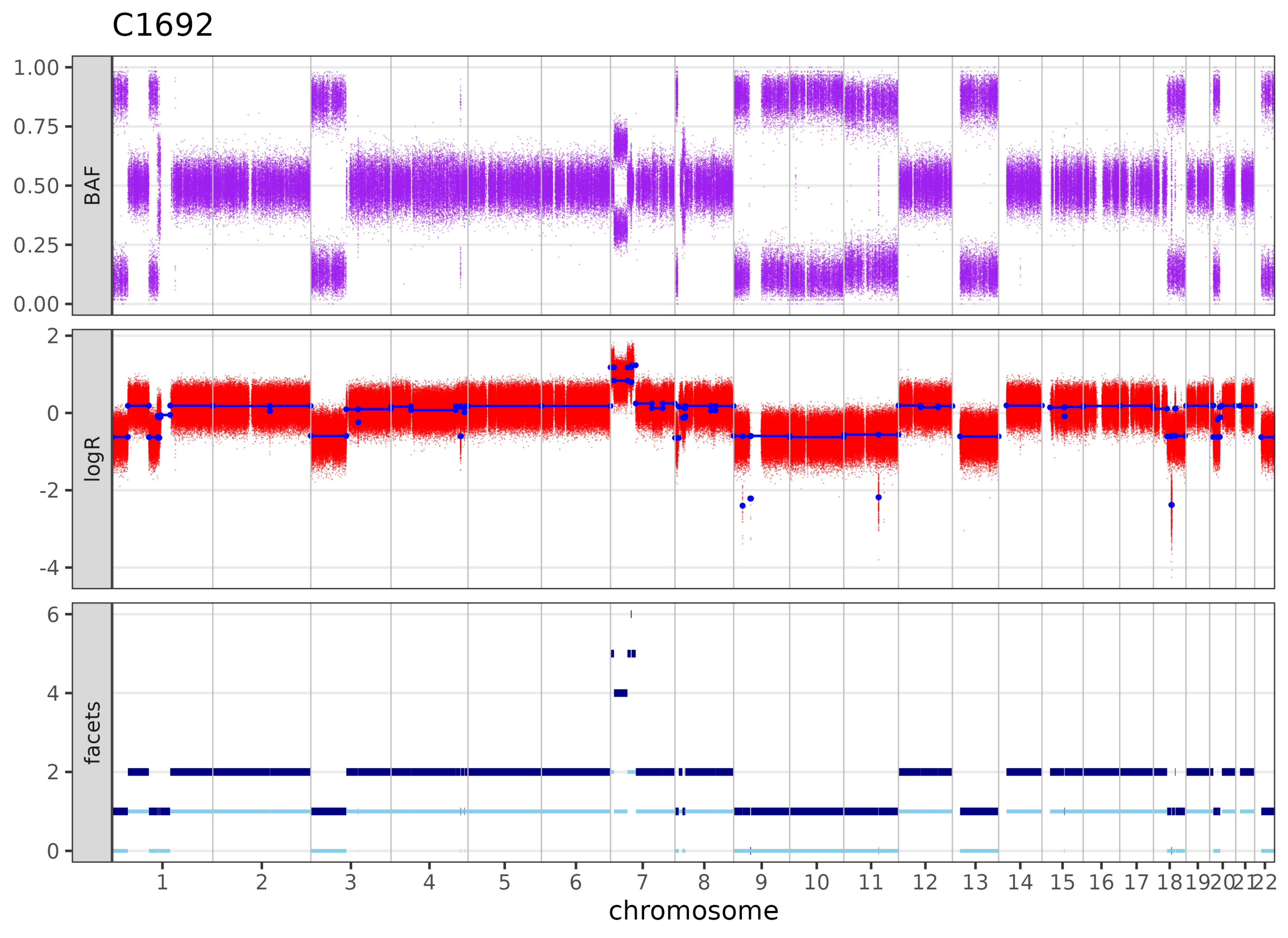
C1692
BAF
1.00
0.75
0.50
0.25
0.00
logR
2
0
-2
-4
facets
6
4
2
0
1 2 3 4 5 6 7 8 9 10 11 12 13 14 15 16 17 18 19 20 21 22
chromosome

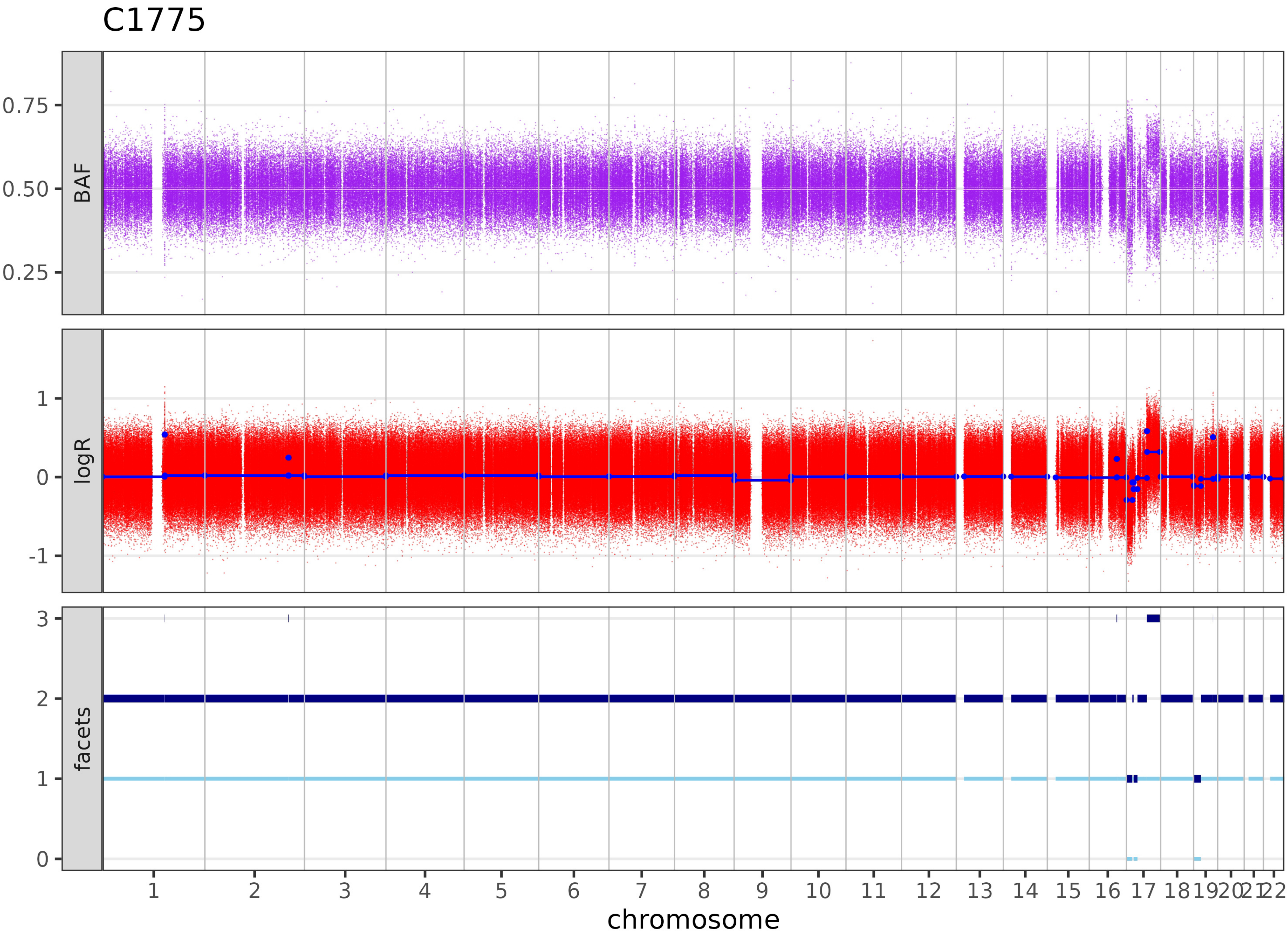
C1775
BAF
0.75
0.50
0.25
logR
1
0
-1
facets
3
2
1
chromosome

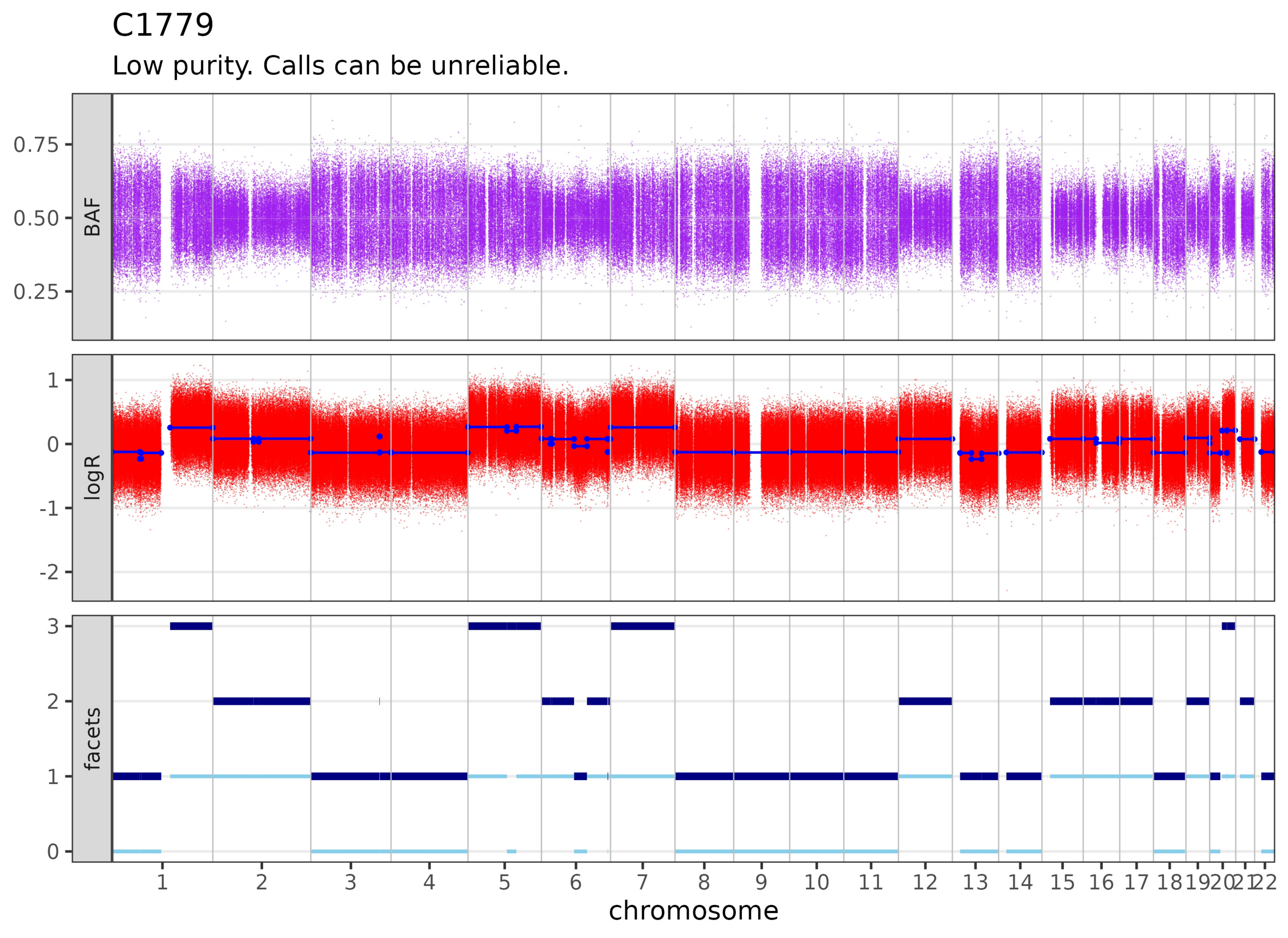
C1779
Low purity. Calls can be unreliable.
BAF
0.75
0.50
0.25
logR
1
0
-1
-2
facets
3
2
1
0
1 2 3 4 5 6 7 8 9 10 11 12 13 14 15 16 17 18 19 20 21 22
chromosome

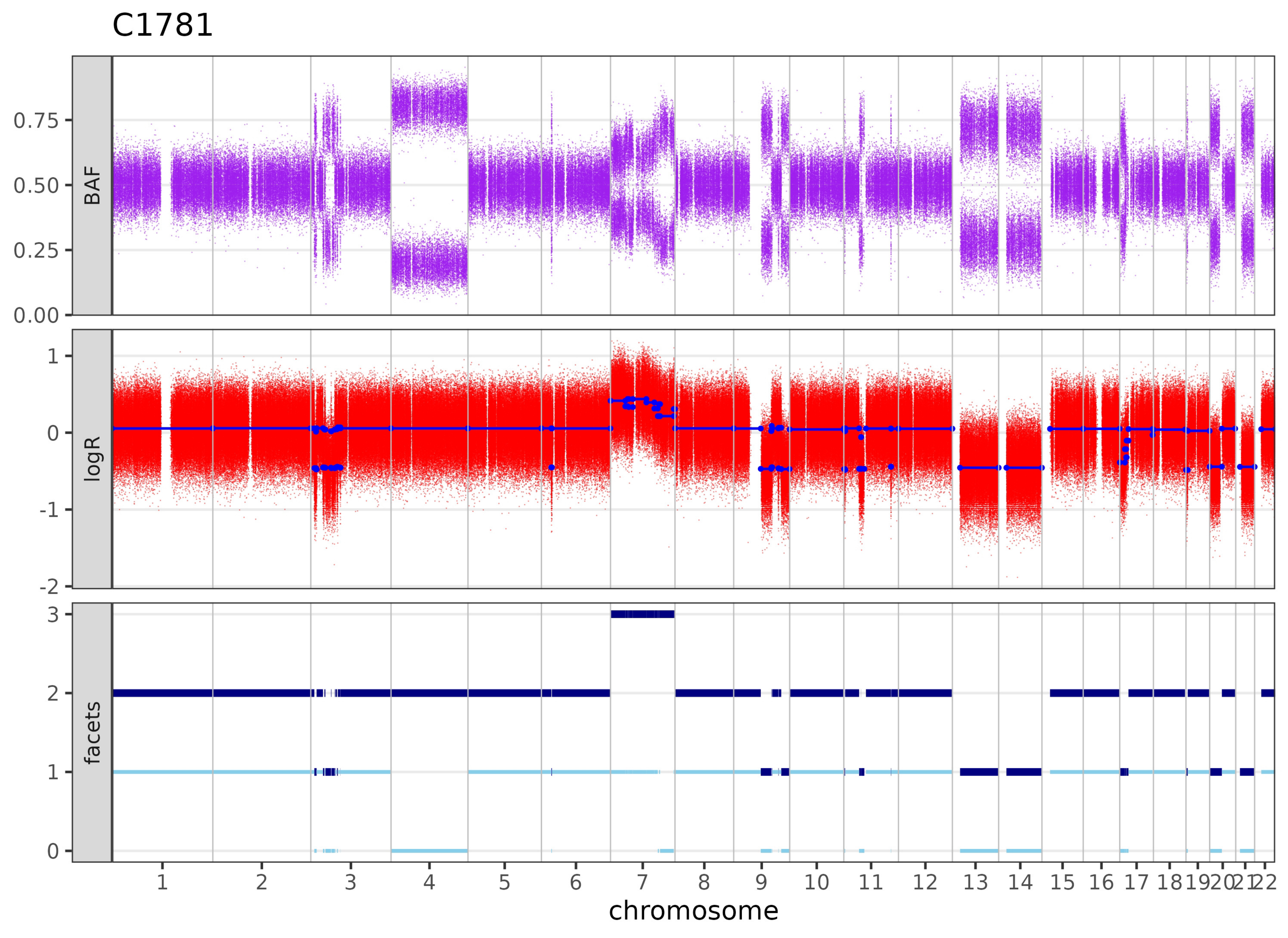
C1781
BAF
0.75
0.50
0.25
0.00
logR
1
0
-1
-2
facets
3
2
1
chromosome

# C1790

Insufficient information to estimate purity. Likely diplod or purity too low.

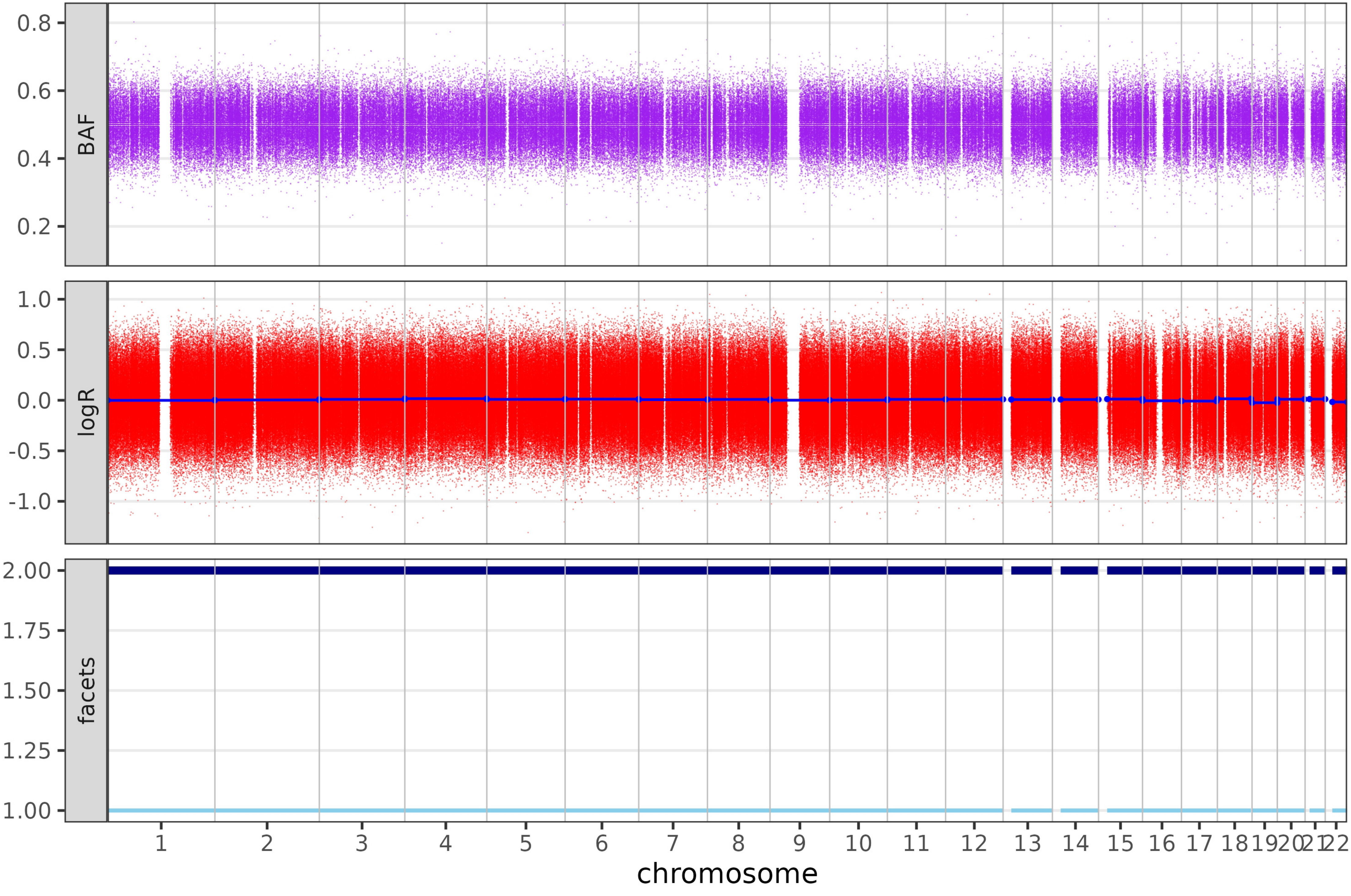

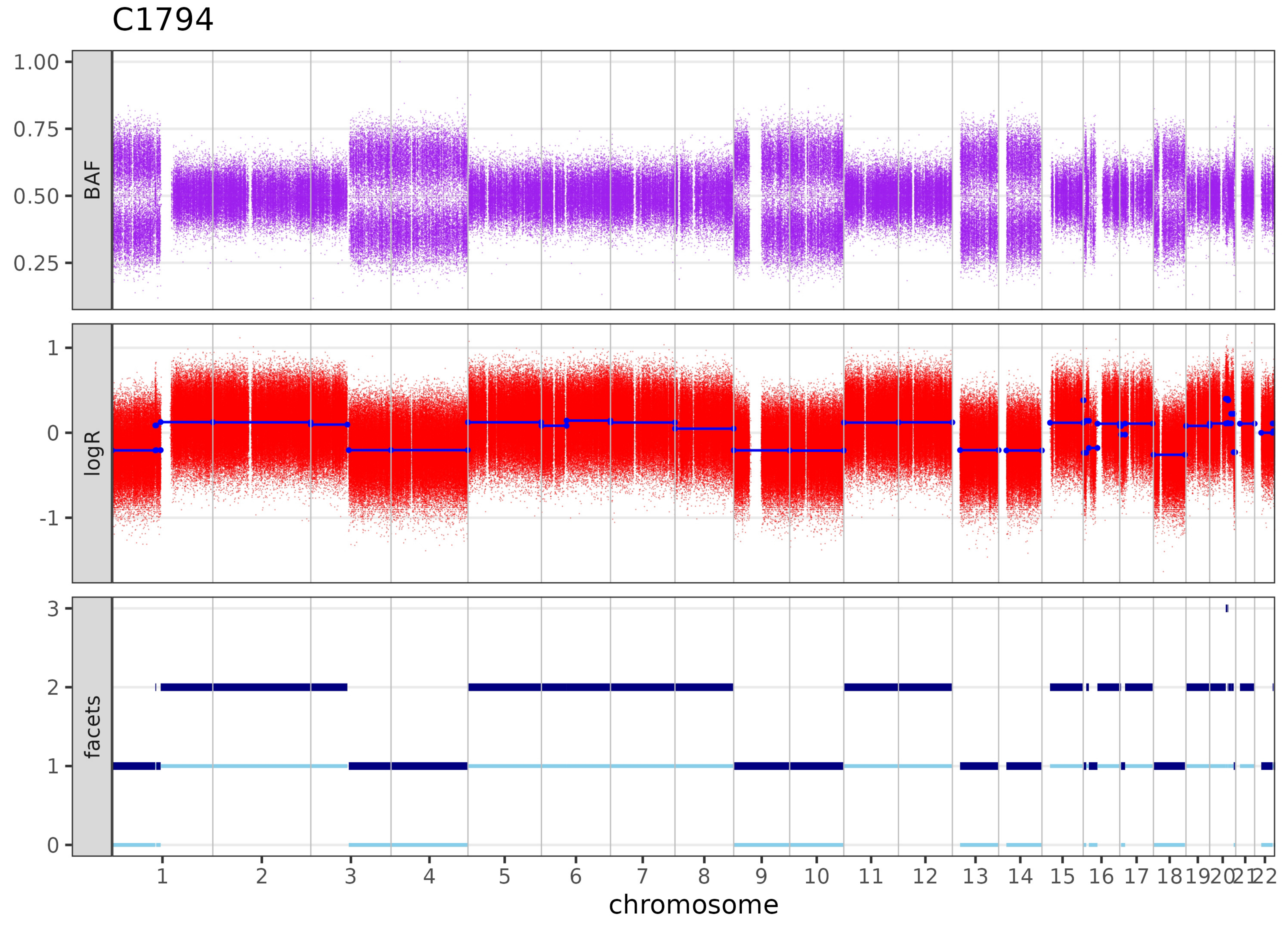

C1794
BAF
logR
facets
chromosome

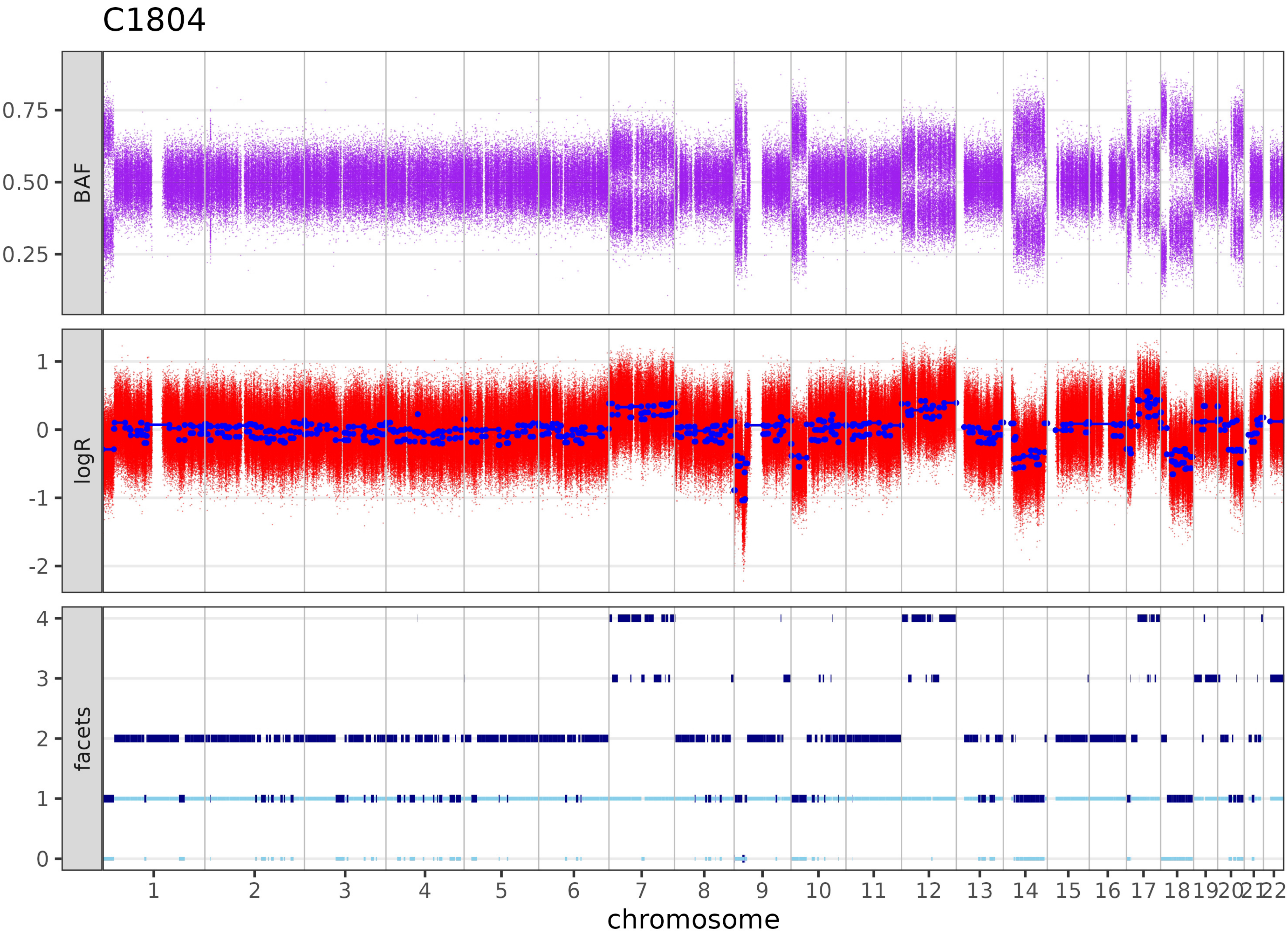

C1804
BAF
0.75
0.50
0.25
logR
1
0
-1
-2
facets
4
3
2
1
0
1 2 3 4 5 6 7 8 9 10 11 12 13 14 15 16 17 18 19 20 21 22
chromosome

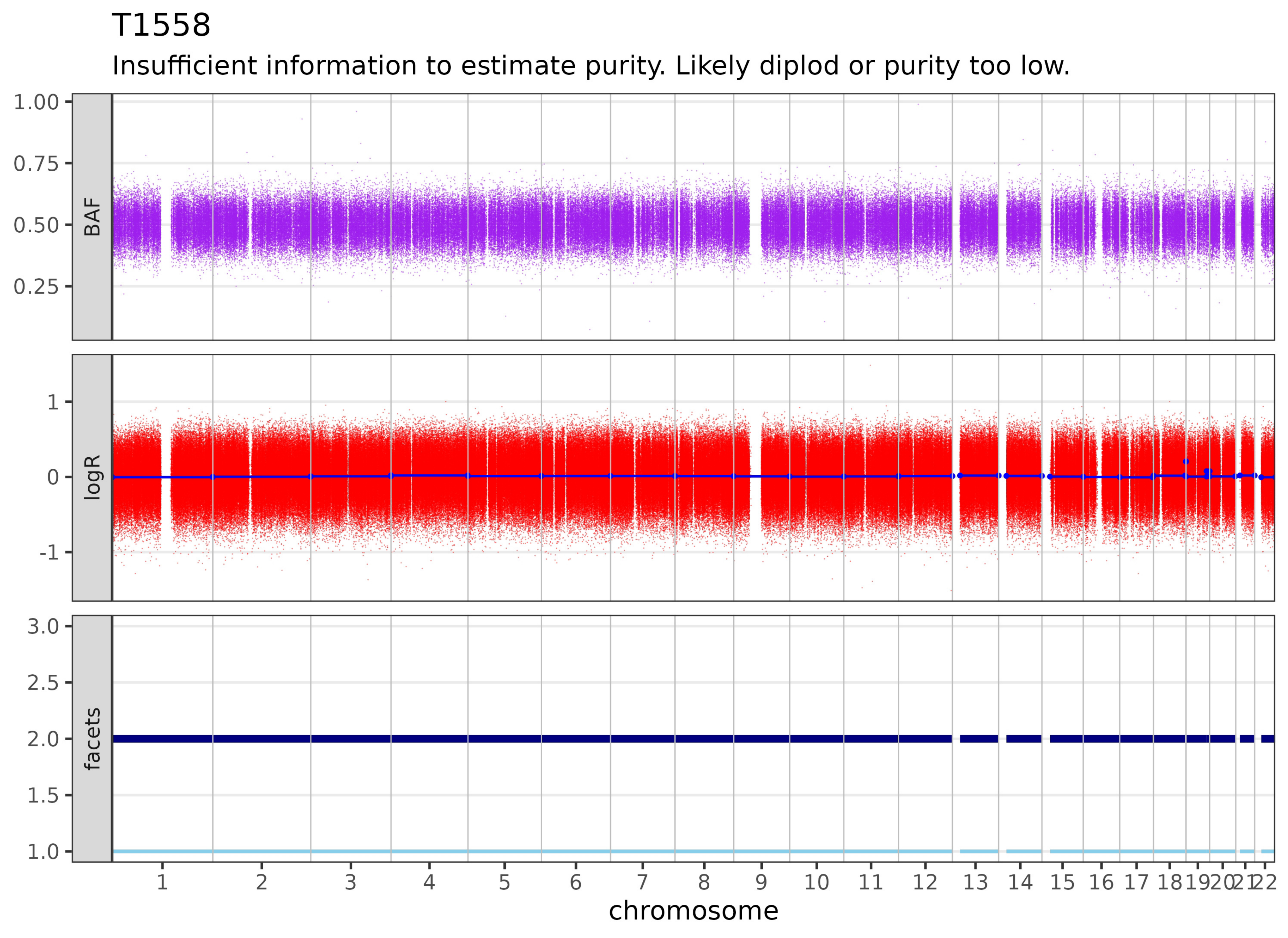
T1558
Insufficient information to estimate purity. Likely diplod or purity too low.
BAF
logR
facets
1.00
0.75
0.50
0.25
1
0
-1
3.0
2.5
2.0
1.5
1.0
1 2 3 4 5 6 7 8 9 10 11 12 13 14 15 16 17 18 19 20 21 22
chromosome

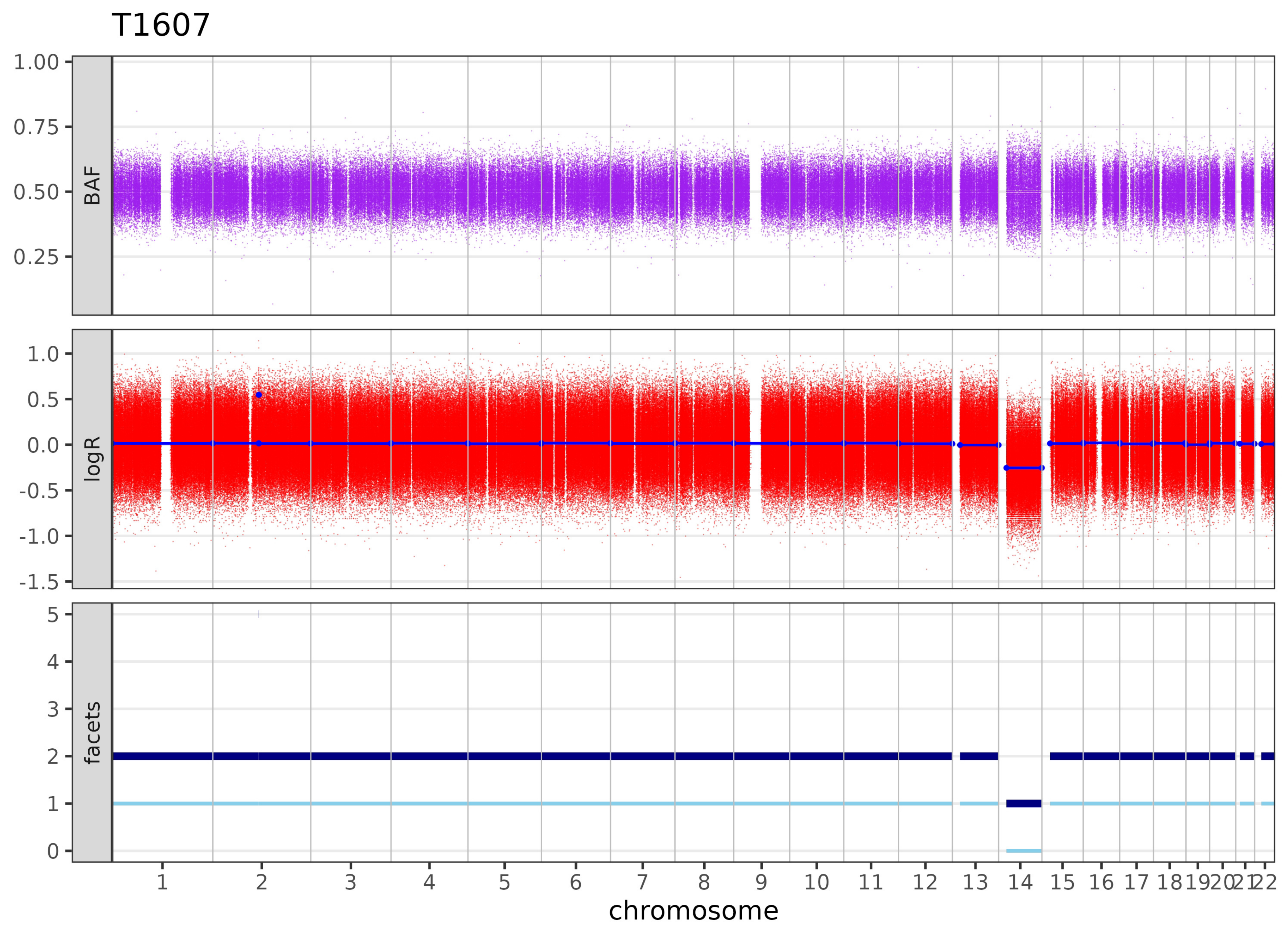

T1607
BAF
logR
facets
1.00
0.75
0.50
0.25
1.0
0.5
0.0
-0.5
-1.0
-1.5
5
4
3
2
1
0
1 2 3 4 5 6 7 8 9 10 11 12 13 14 15 16 17 18 19 20 21 22
chromosome

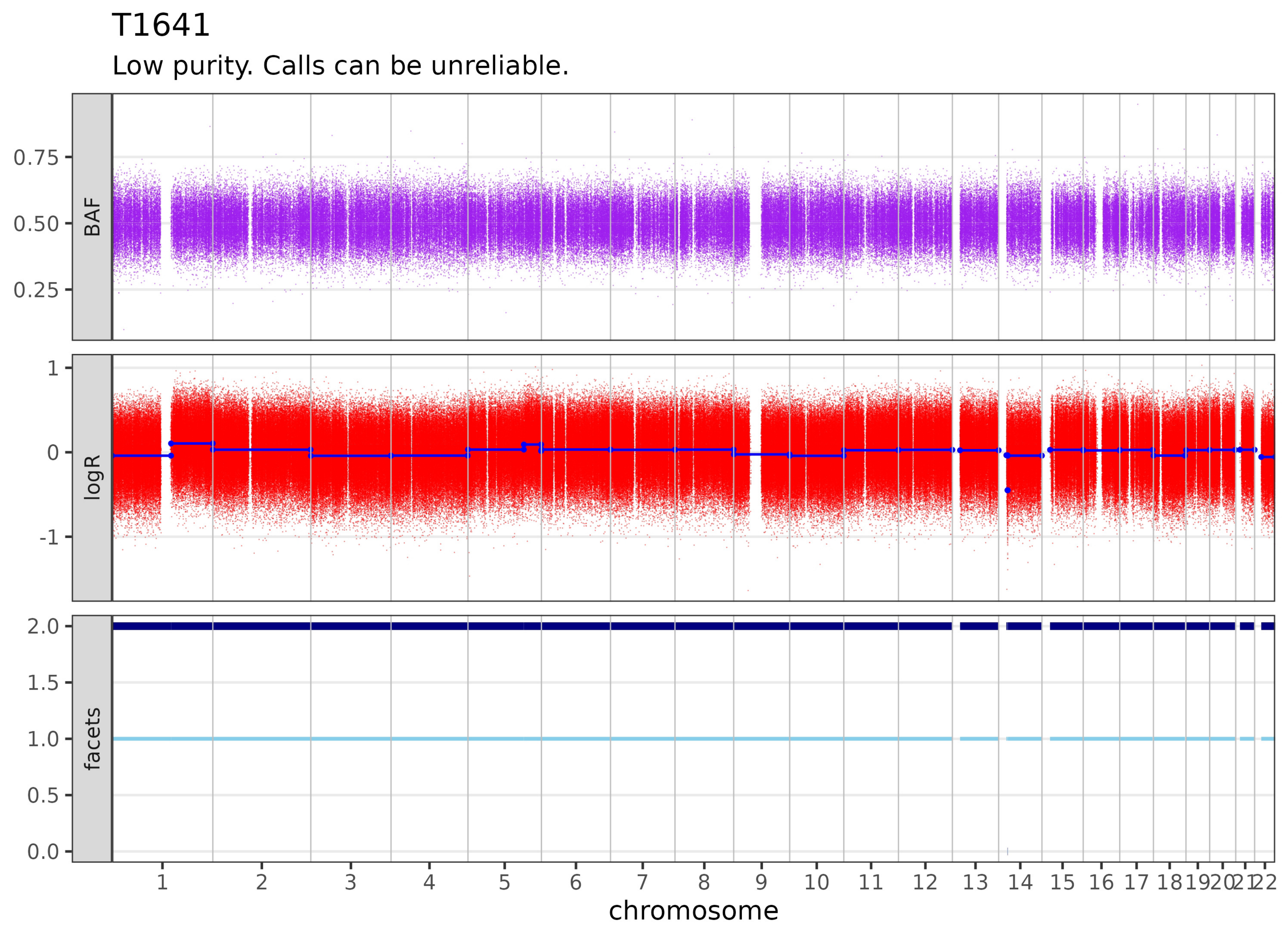

T1641
Low purity. Calls can be unreliable.
BAF
0.75
0.50
0.25
logR
1
0
-1
facets
2.0
1.5
1.0
0.5
0.0
1 2 3 4 5 6 7 8 9 10 11 12 13 14 15 16 17 18 19 20 21 22
chromosome

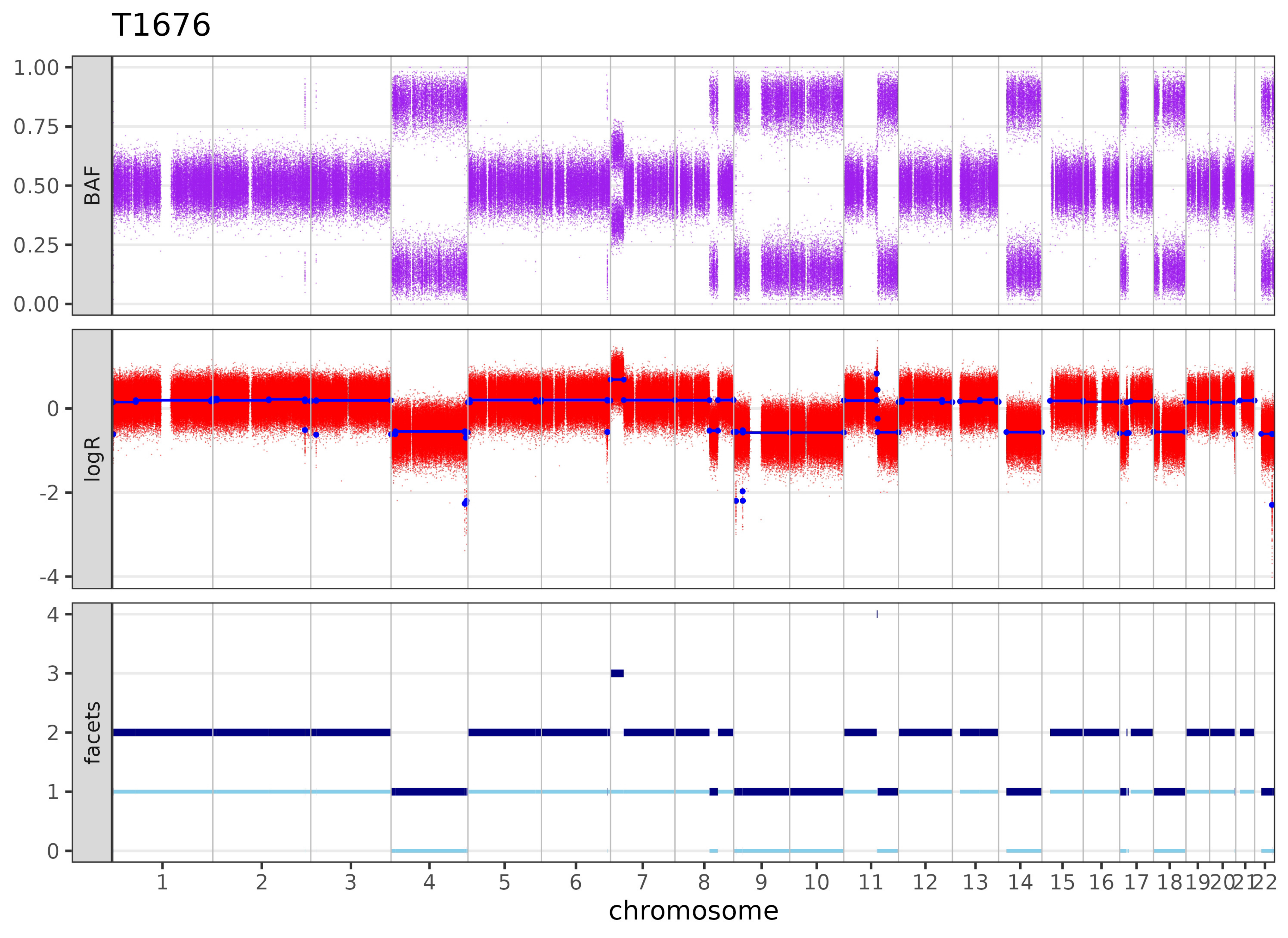
T1676
BAF
1.00
0.75
0.50
0.25
0.00
logR
0
-2
-4
facets
4
3
2
1
0
1 2 3 4 5 6 7 8 9 10 11 12 13 14 15 16 17 18 19 20 21 22
chromosome

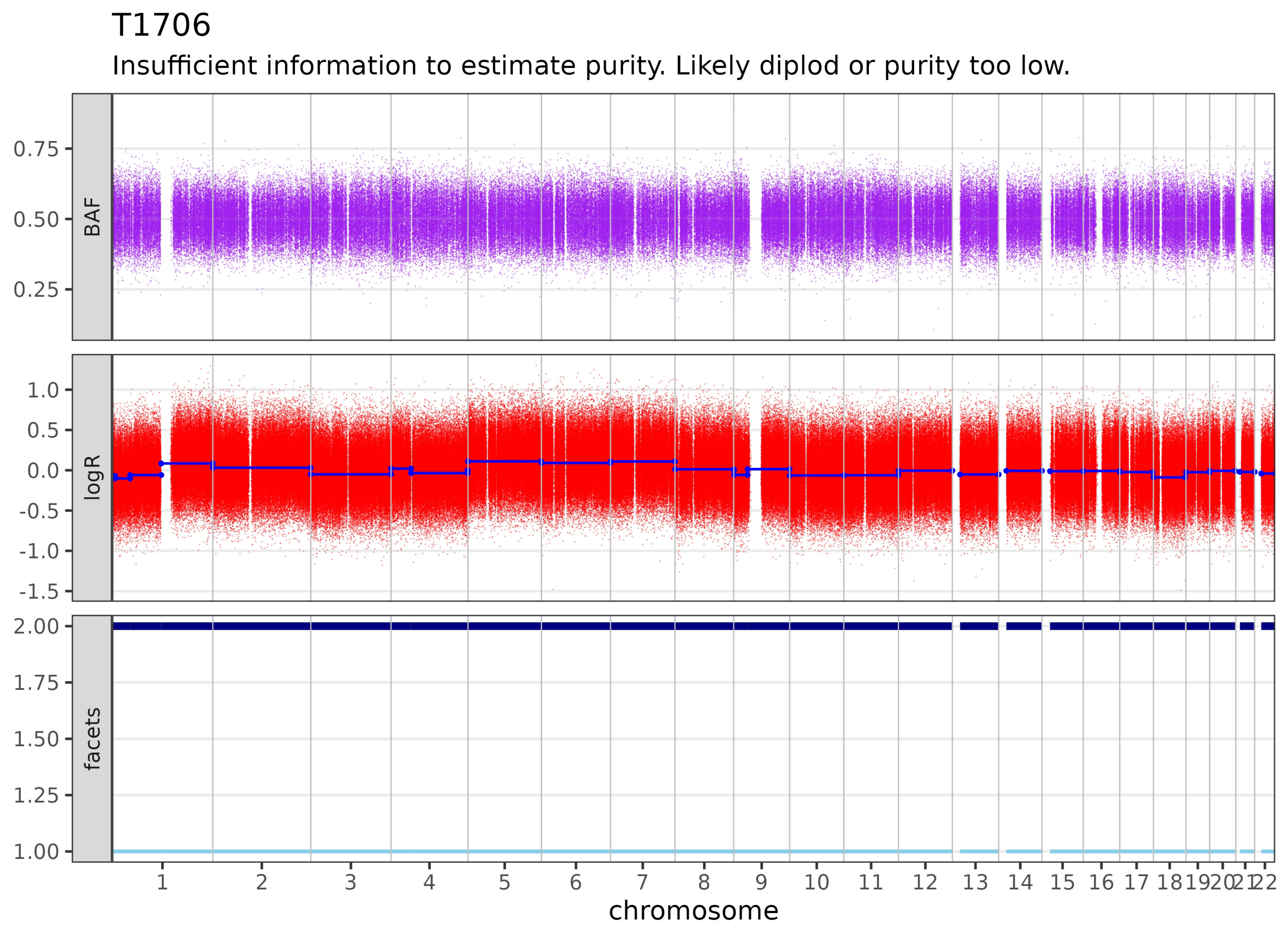
T1706
Insufficient information to estimate purity. Likely diplod or purity too low.
BAF
0.75
0.50
0.25
logR
1.0
0.5
0.0
-0.5
-1.0
-1.5
facets
2.00
1.75
1.50
1.25
1.00
1 2 3 4 5 6 7 8 9 10 11 12 13 14 15 16 17 18 19 20 21 22
chromosome

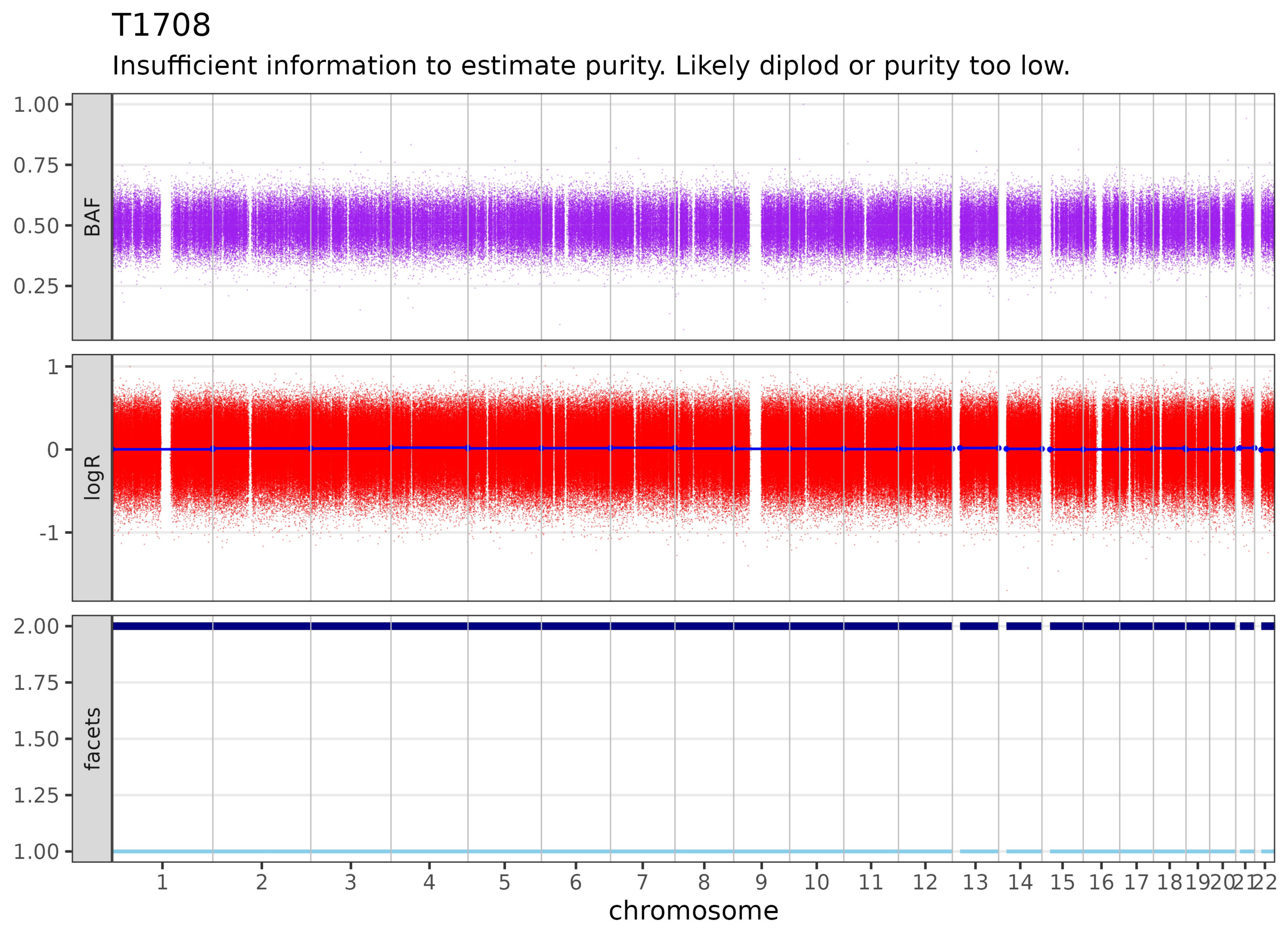

T1708
Insufficient information to estimate purity. Likely diplod or purity too low.
BAF
logR
facets
1.00
0.75
0.50
0.25
1
0
-1
2.00
1.75
1.50
1.25
1.00
1 2 3 4 5 6 7 8 9 10 11 12 13 14 15 16 17 18 19 20 21 22
chromosome

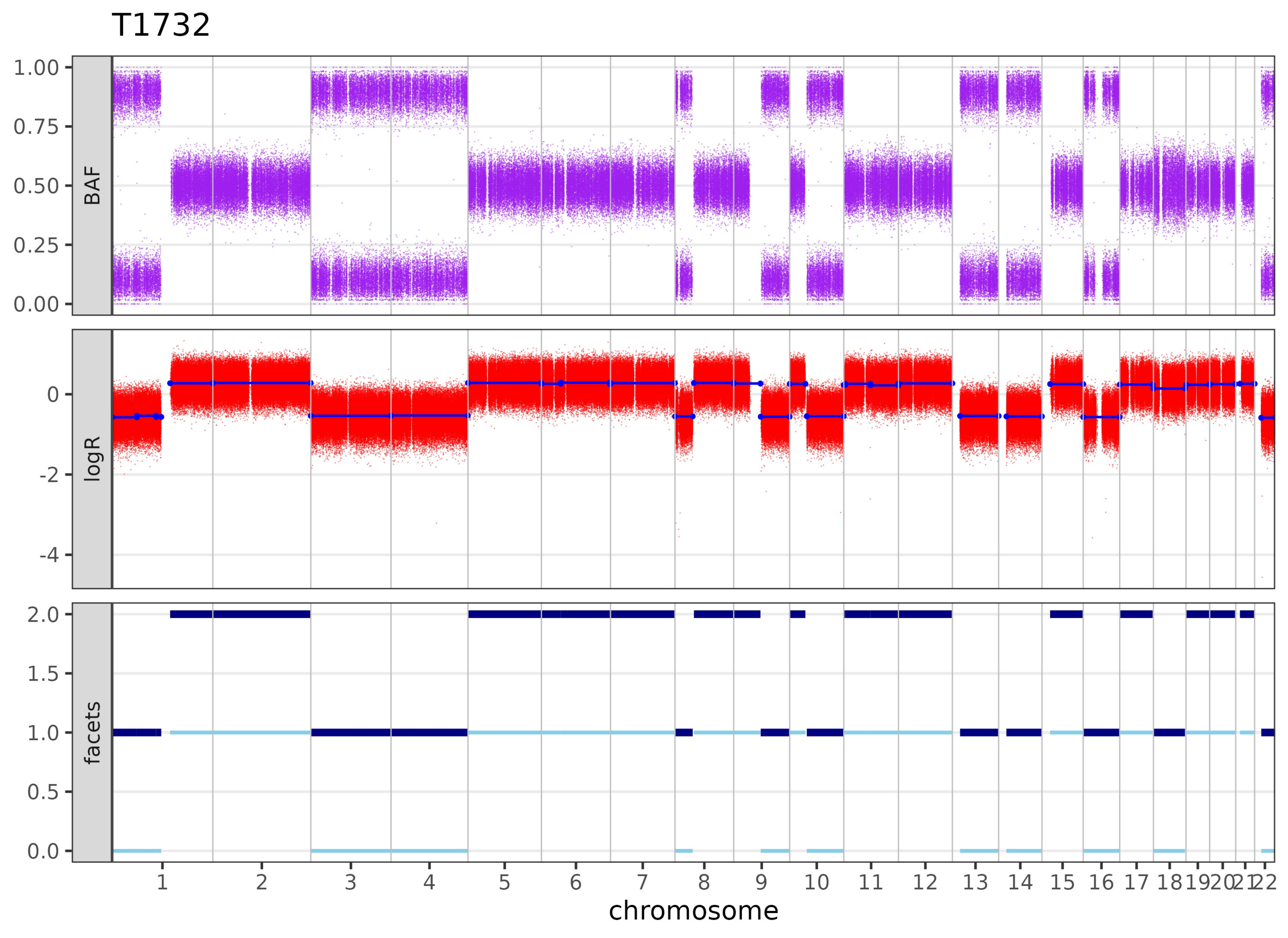

T1732
BAF
1.00
0.75
0.50
0.25
0.00
logR
0
-2
-4
facets
2.0
1.5
1.0
0.5
0.0
1 2 3 4 5 6 7 8 9 10 11 12 13 14 15 16 17 18 19 20 21 22
chromosome

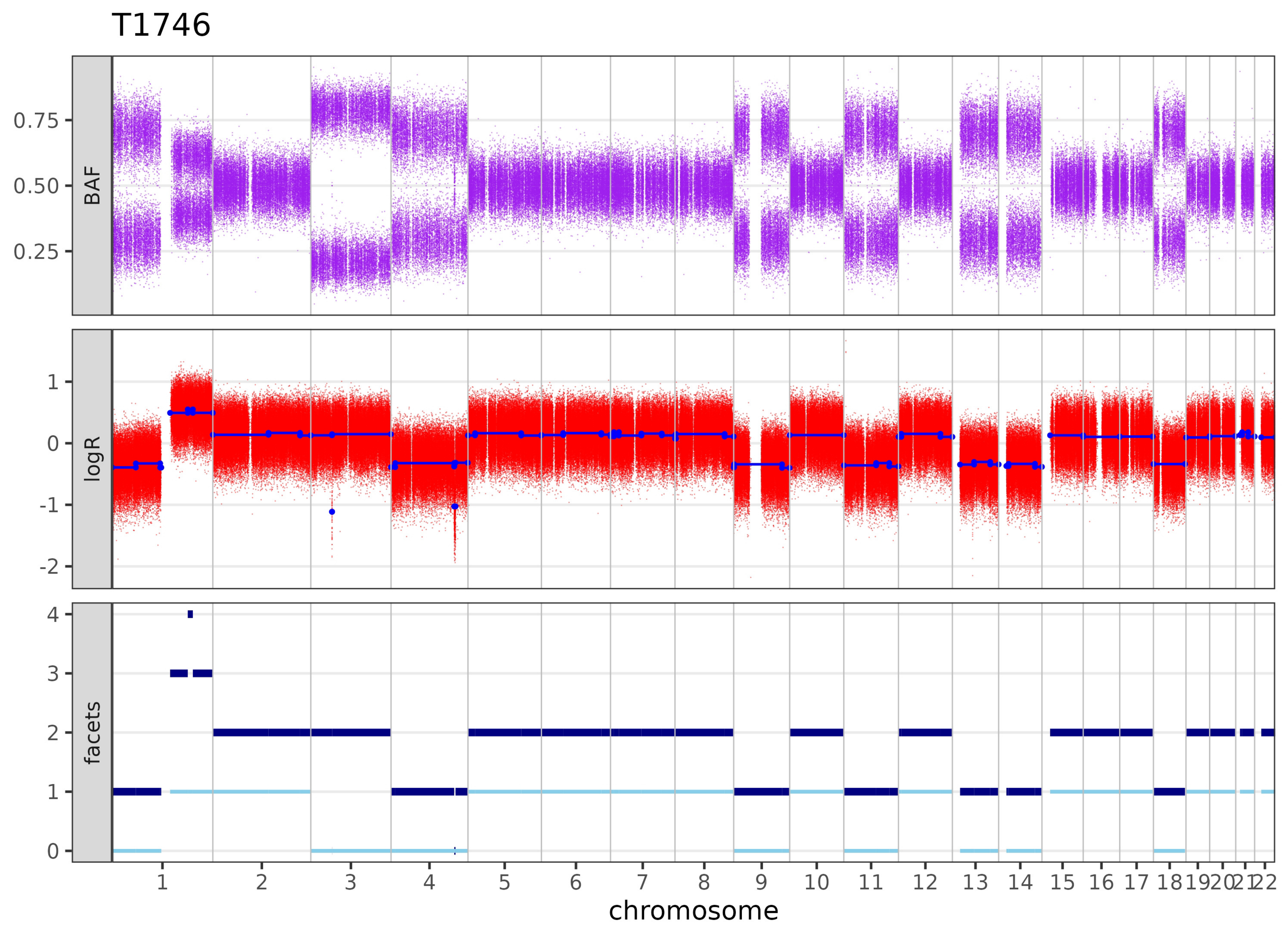

T1746
BAF
0.75
0.50
0.25
logR
1
0
-1
-2
facets
4
3
2
1
0
1 2 3 4 5 6 7 8 9 10 11 12 13 14 15 16 17 18 19 20 21 22
chromosome

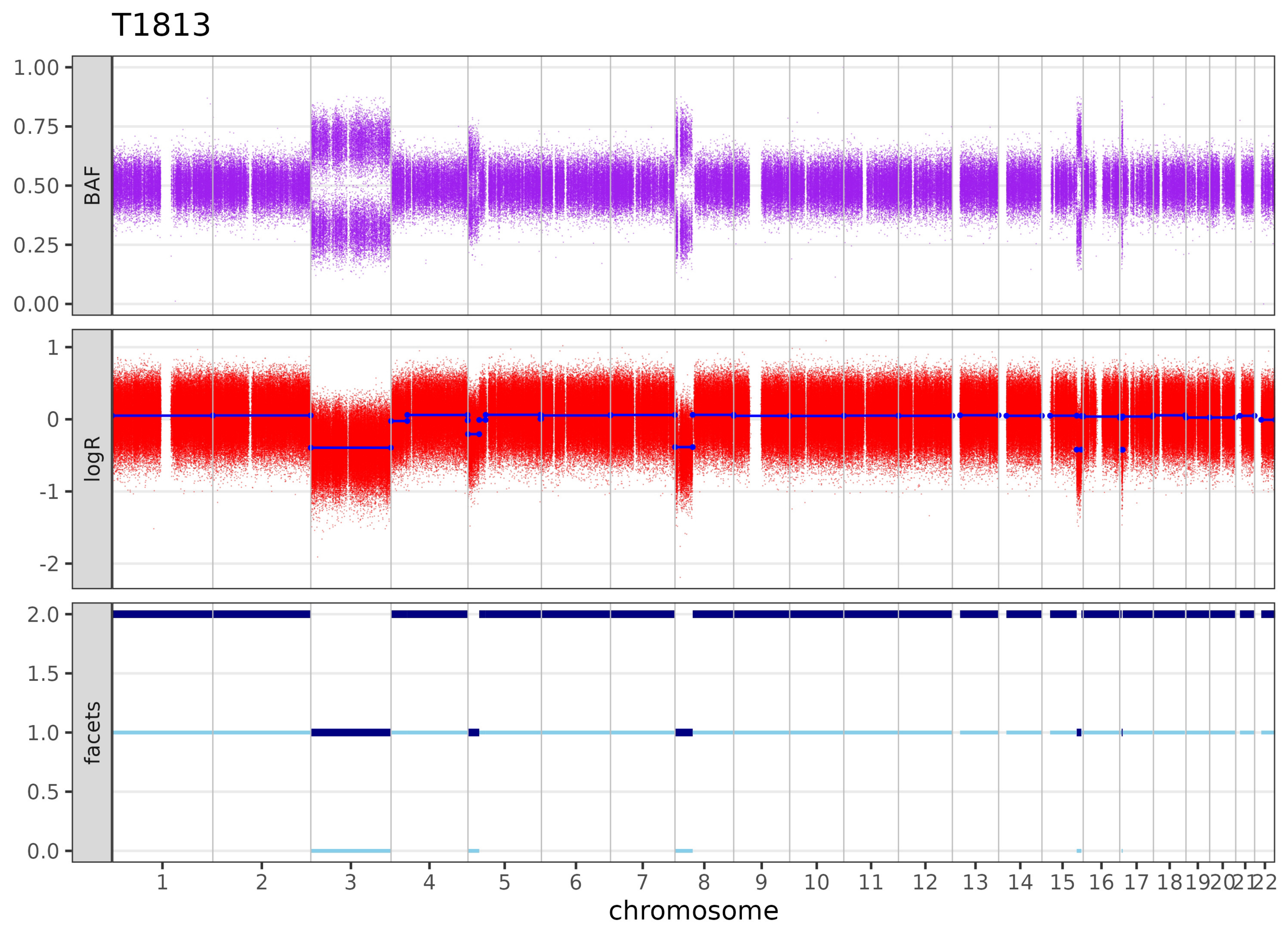

T1813
1.00
0.75
0.50
0.25
0.00
BAF
1
0
-1
-2
logR
2.0
1.5
1.0
0.5
0.0
facets
chromosome

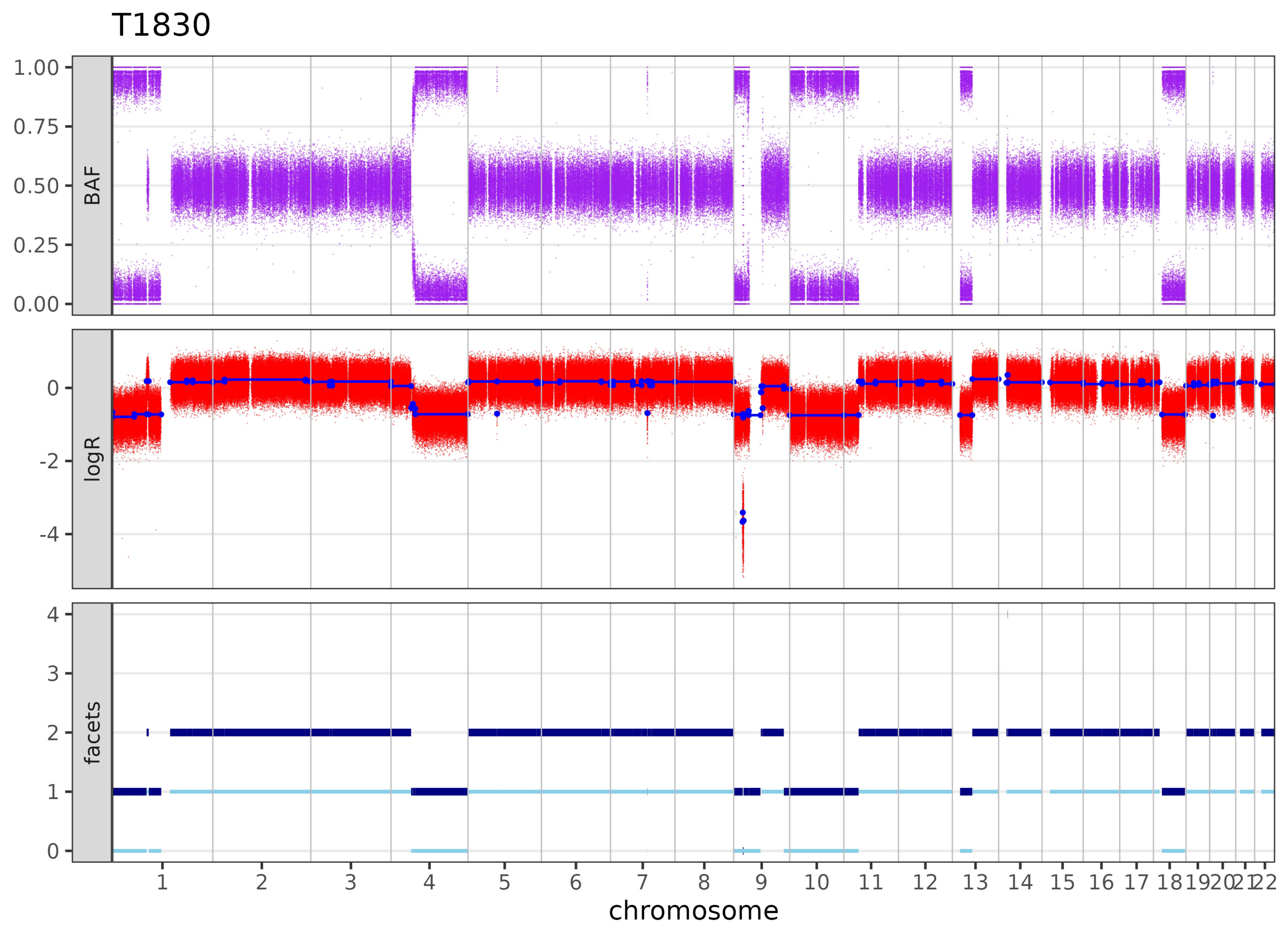

T1830
BAF
logR
facets
1.00
0.75
0.50
0.25
0.00
0
-2
-4
4
3
2
1
0
1 2 3 4 5 6 7 8 9 10 11 12 13 14 15 16 17 18 19 20 21 22
chromosome

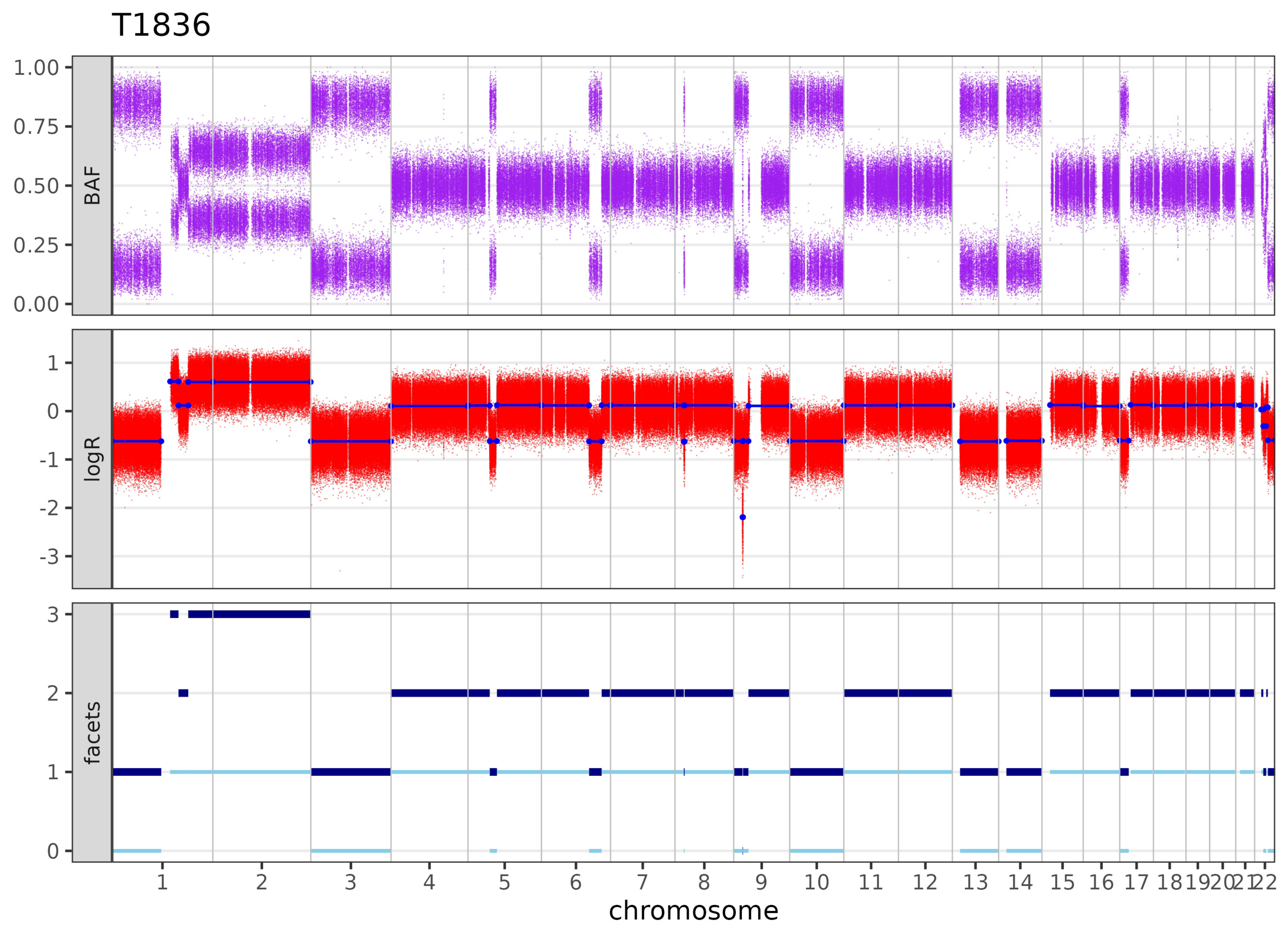

T1836
BAF
1.00
0.75
0.50
0.25
0.00
logR
1
0
-1
-2
-3
facets
3
2
1
0
1 2 3 4 5 6 7 8 9 10 11 12 13 14 15 16 17 18 19 20 21 22
chromosome

T1844

Low purity. Calls can be unreliable.

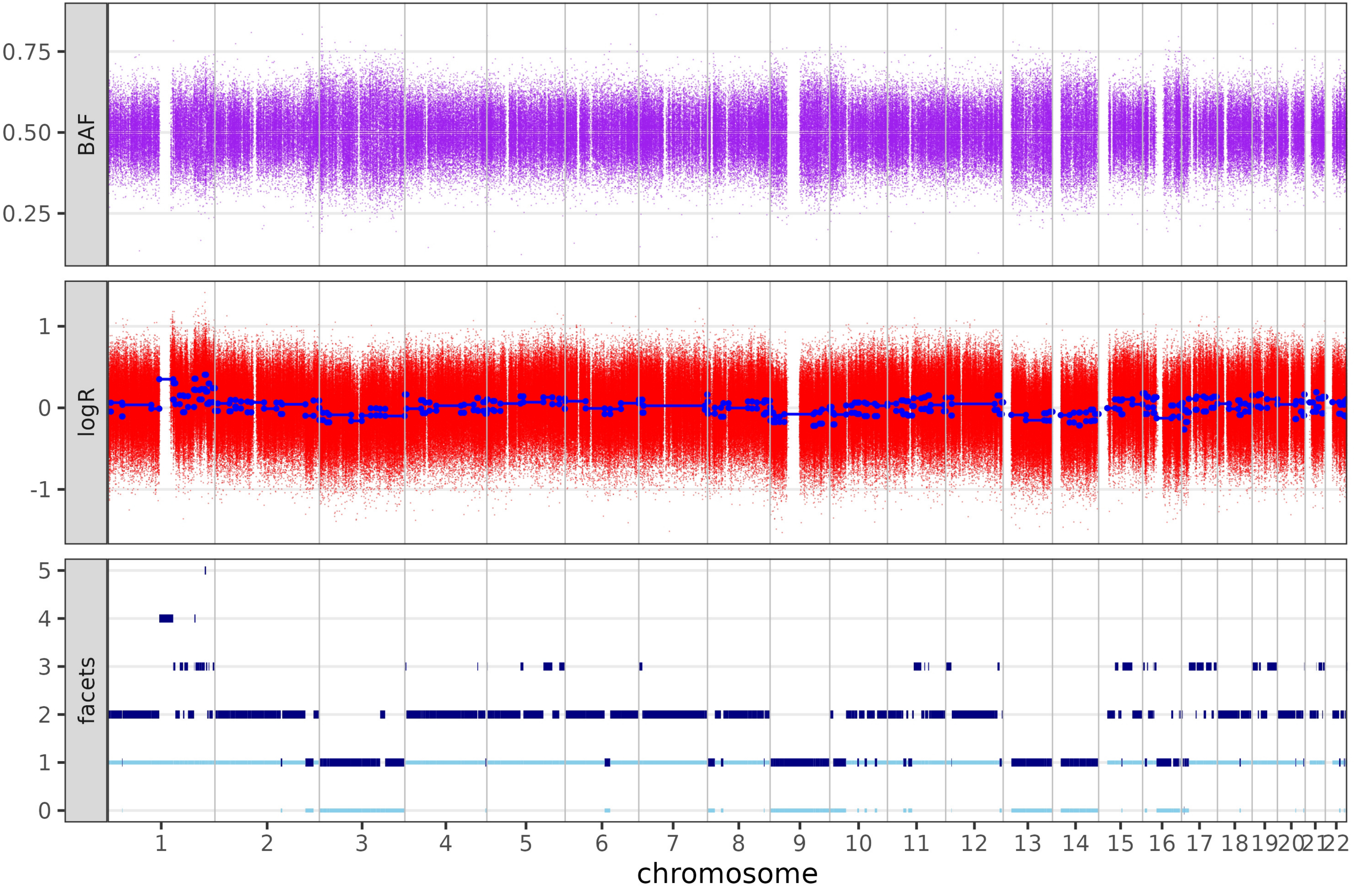

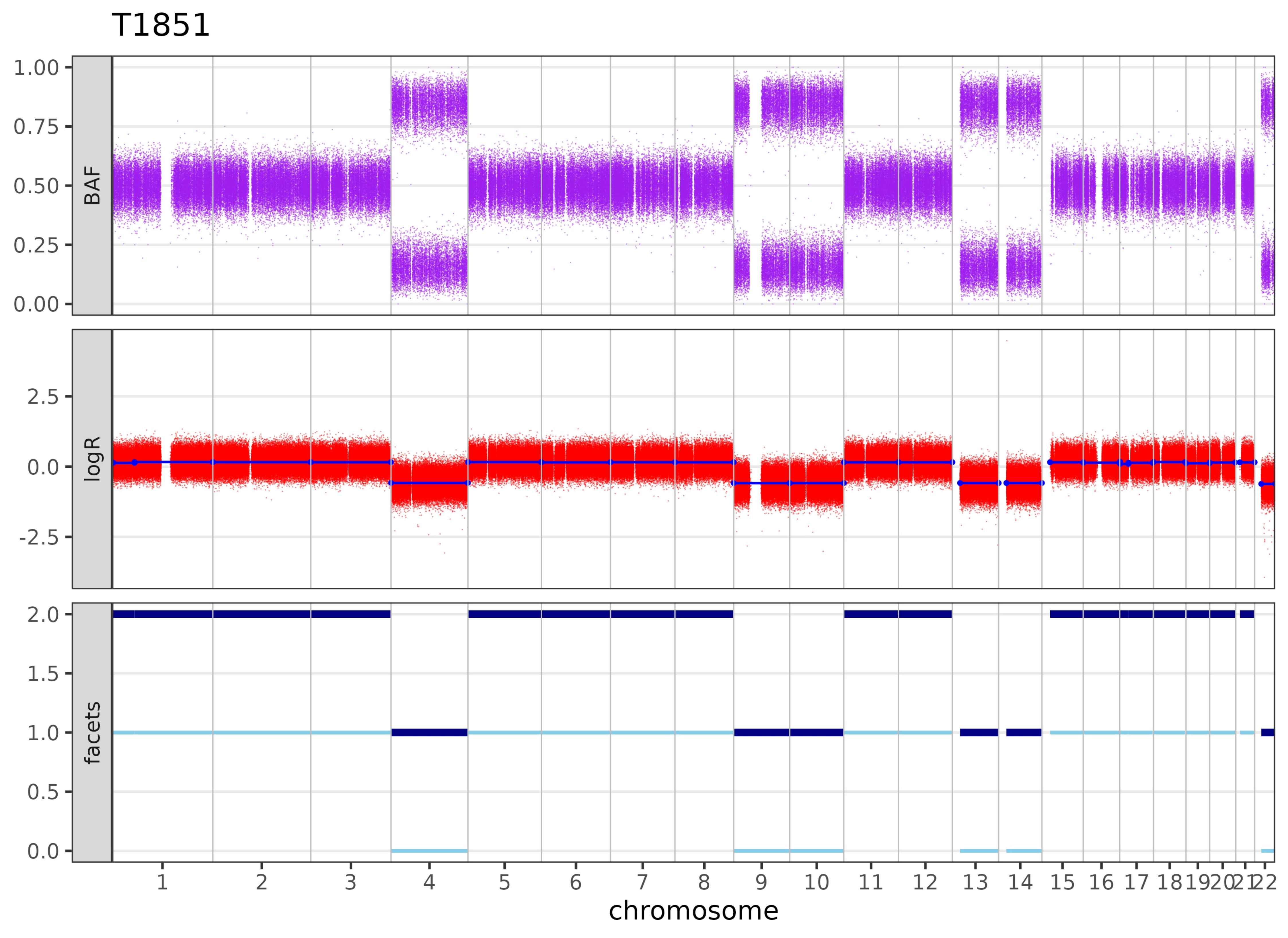

T1851
BAF
1.00
0.75
0.50
0.25
0.00
logR
2.5
0.0
-2.5
facets
2.0
1.5
1.0
0.5
0.0
1 2 3 4 5 6 7 8 9 10 11 12 13 14 15 16 17 18 19 20 21 22
chromosome

# T1853

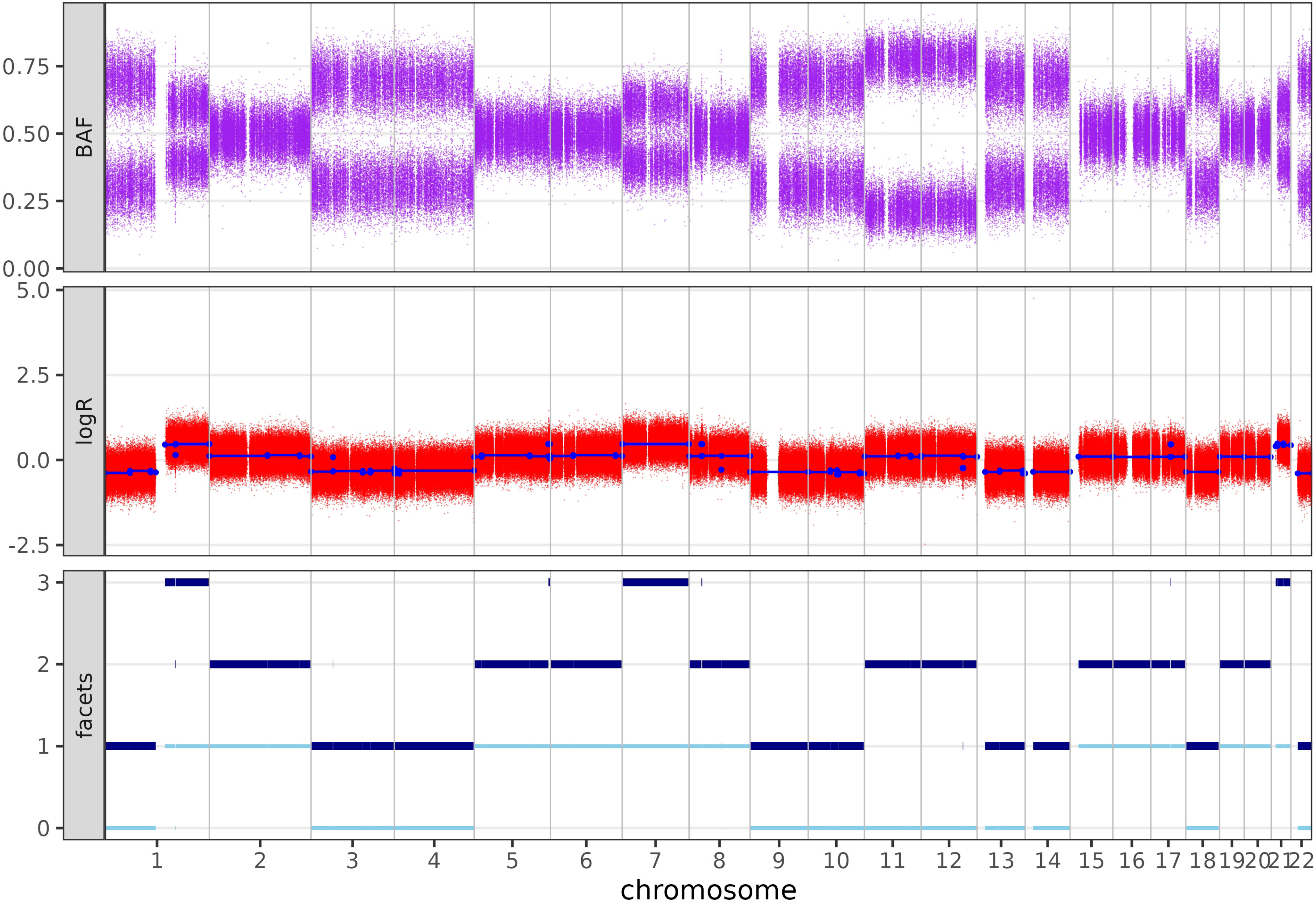
BAF
logR
facets
0.00
0.25
0.50
0.75
-2.5
0.0
2.5
5.0
0
1
2
3
1 2 3 4 5 6 7 8 9 10 11 12 13 14 15 16 17 18 19 20 21 22
chromosome

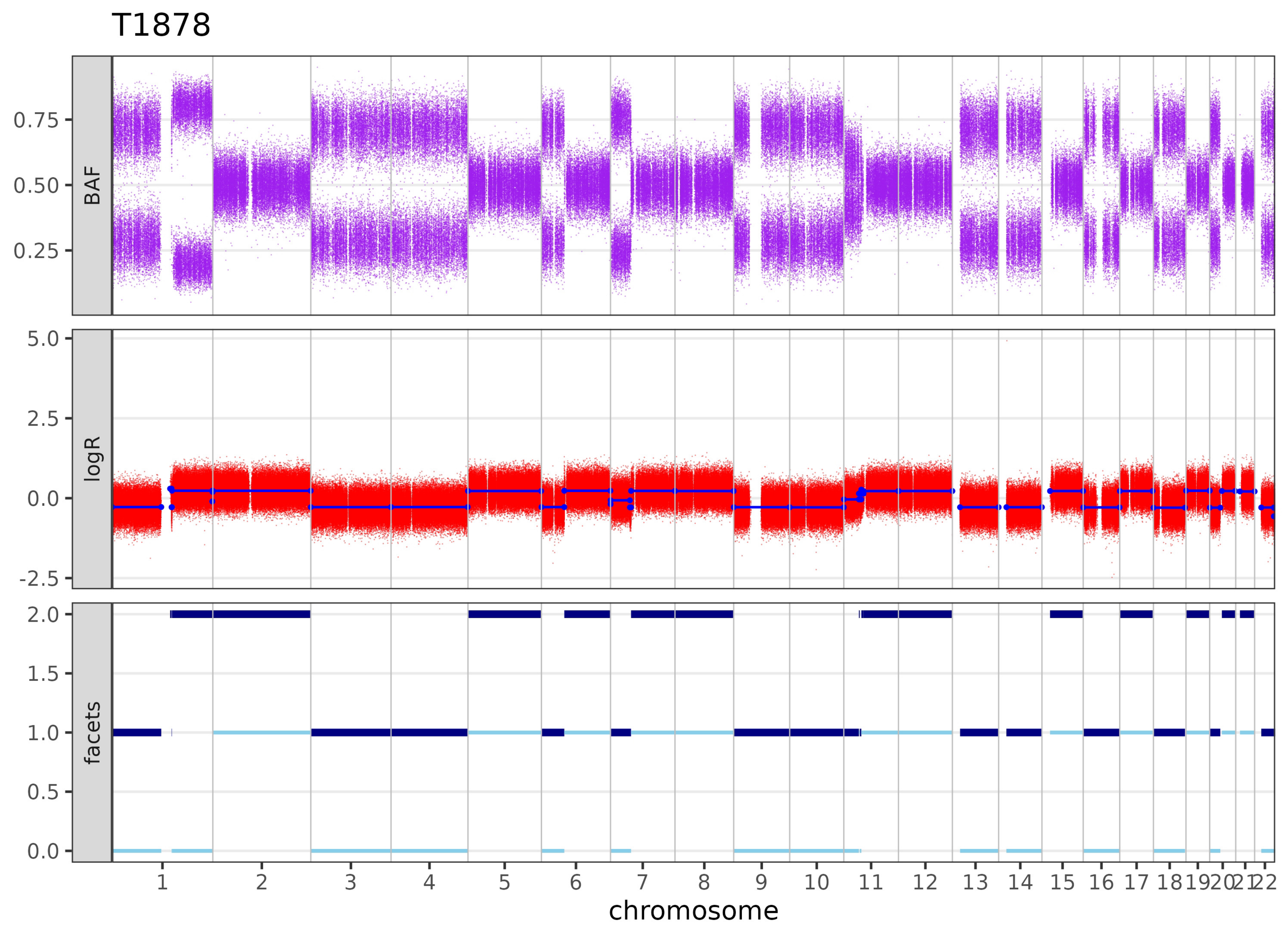
T1878
BAF
0.75
0.50
0.25
logR
5.0
2.5
0.0
-2.5
facets
2.0
1.5
1.0
0.5
0.0
chromosome

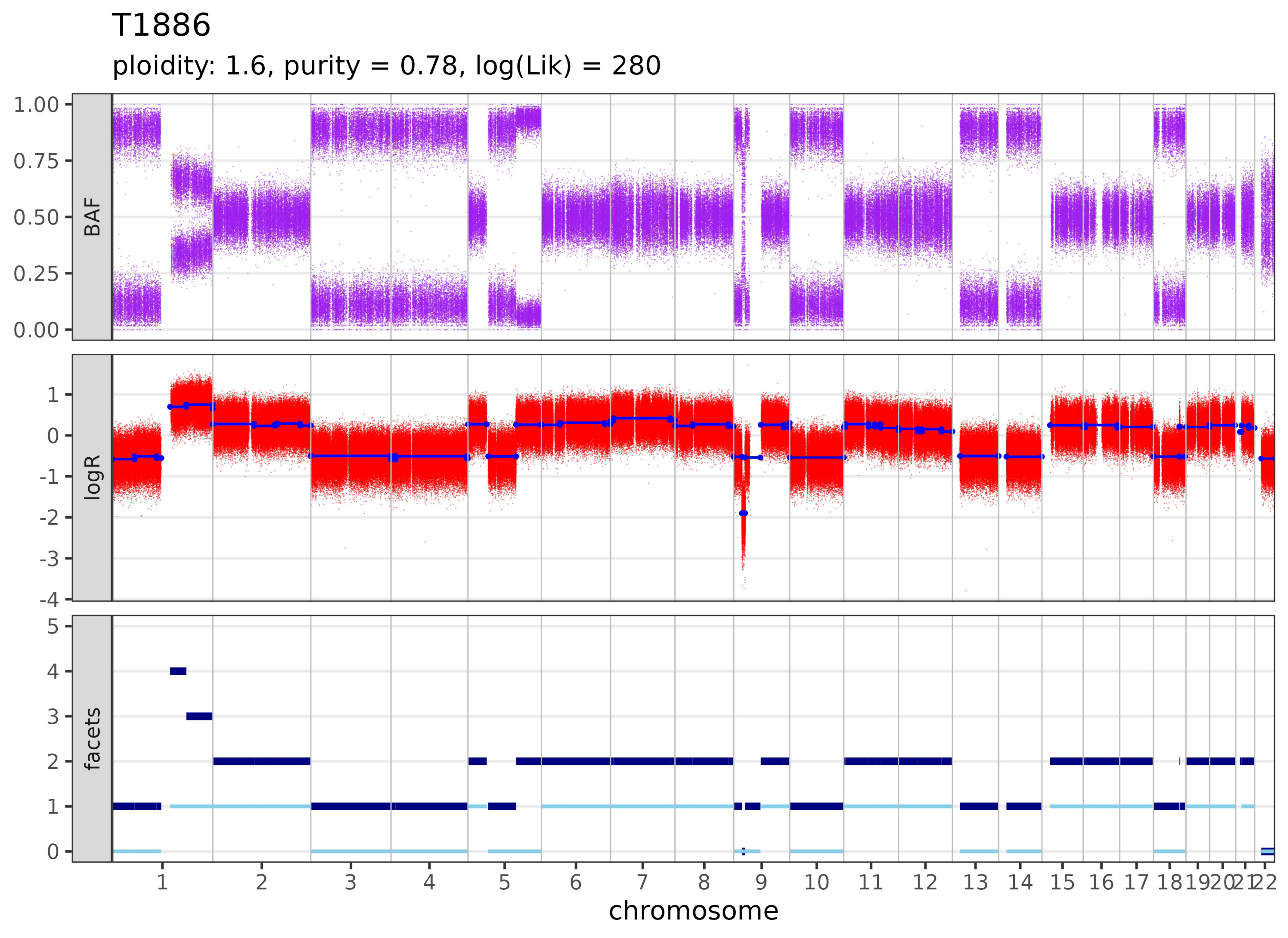

T1886
ploidity: 1.6, purity = 0.78, log(Lik) = 280
BAF
1.00
0.75
0.50
0.25
0.00
logR
1
0
-1
-2
-3
-4
facets
5
4
3
2
1
0
1 2 3 4 5 6 7 8 9 10 11 12 13 14 15 16 17 18 19 20 21 22
chromosome

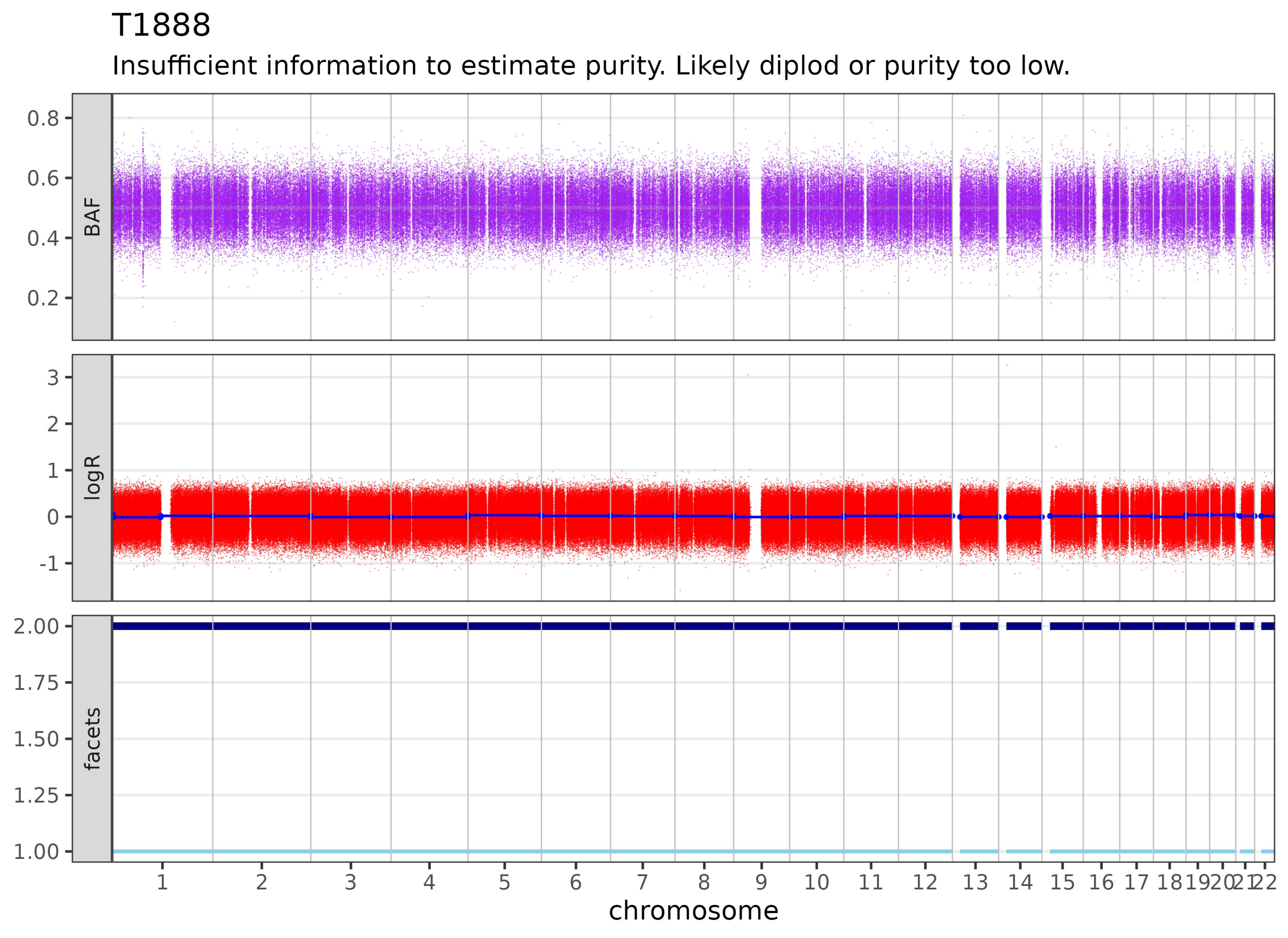
T1888
Insufficient information to estimate purity. Likely diplod or purity too low.
BAF
0.8
0.6
0.4
0.2
logR
3
2
1
0
-1
facets
2.00
1.75
1.50
1.25
1.00
1 2 3 4 5 6 7 8 9 10 11 12 13 14 15 16 17 18 19 20 21 22
chromosome

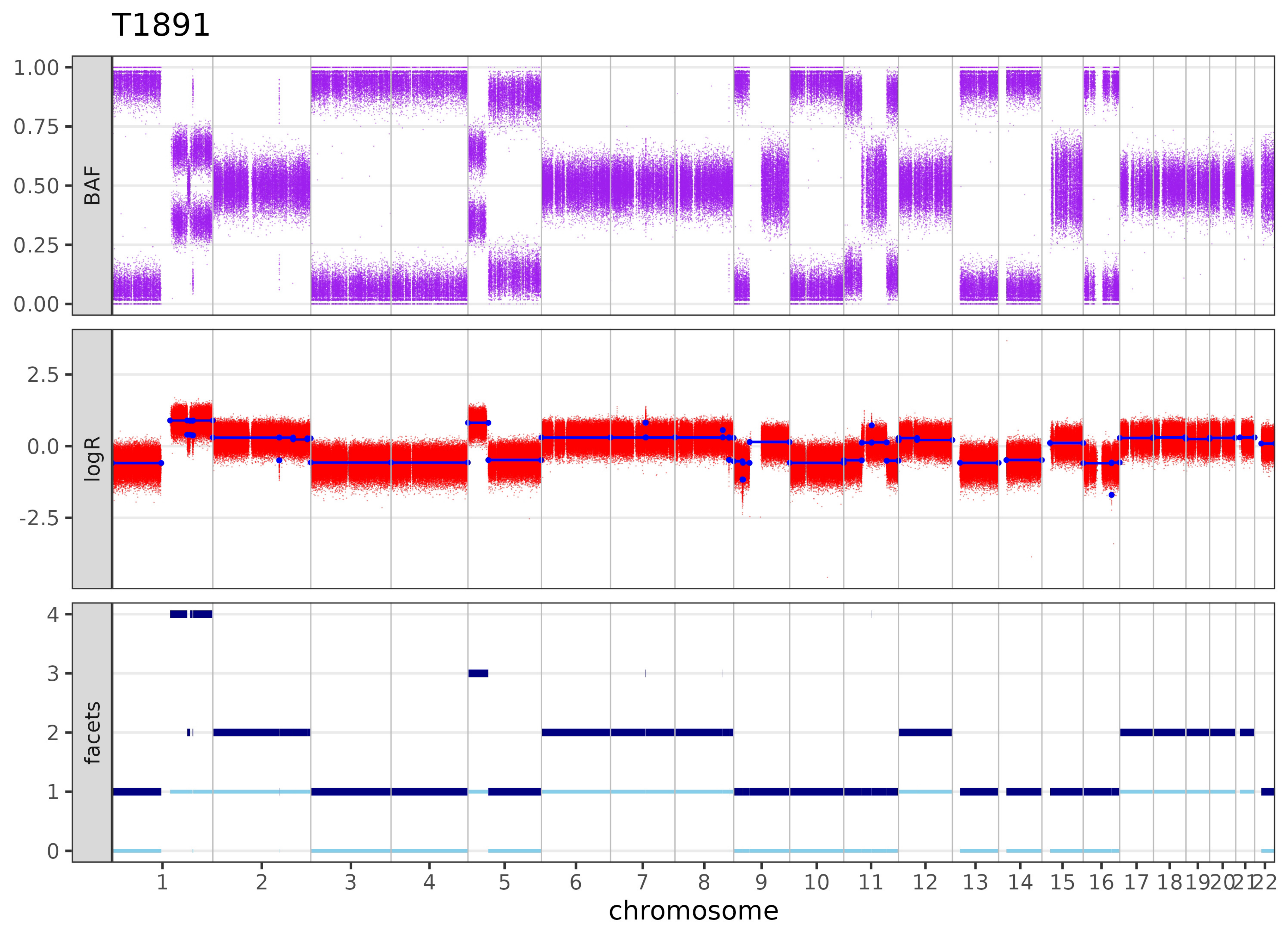

T1891
BAF
1.00
0.75
0.50
0.25
0.00
logR
2.5
0.0
-2.5
facets
4
3
2
1
chromosome

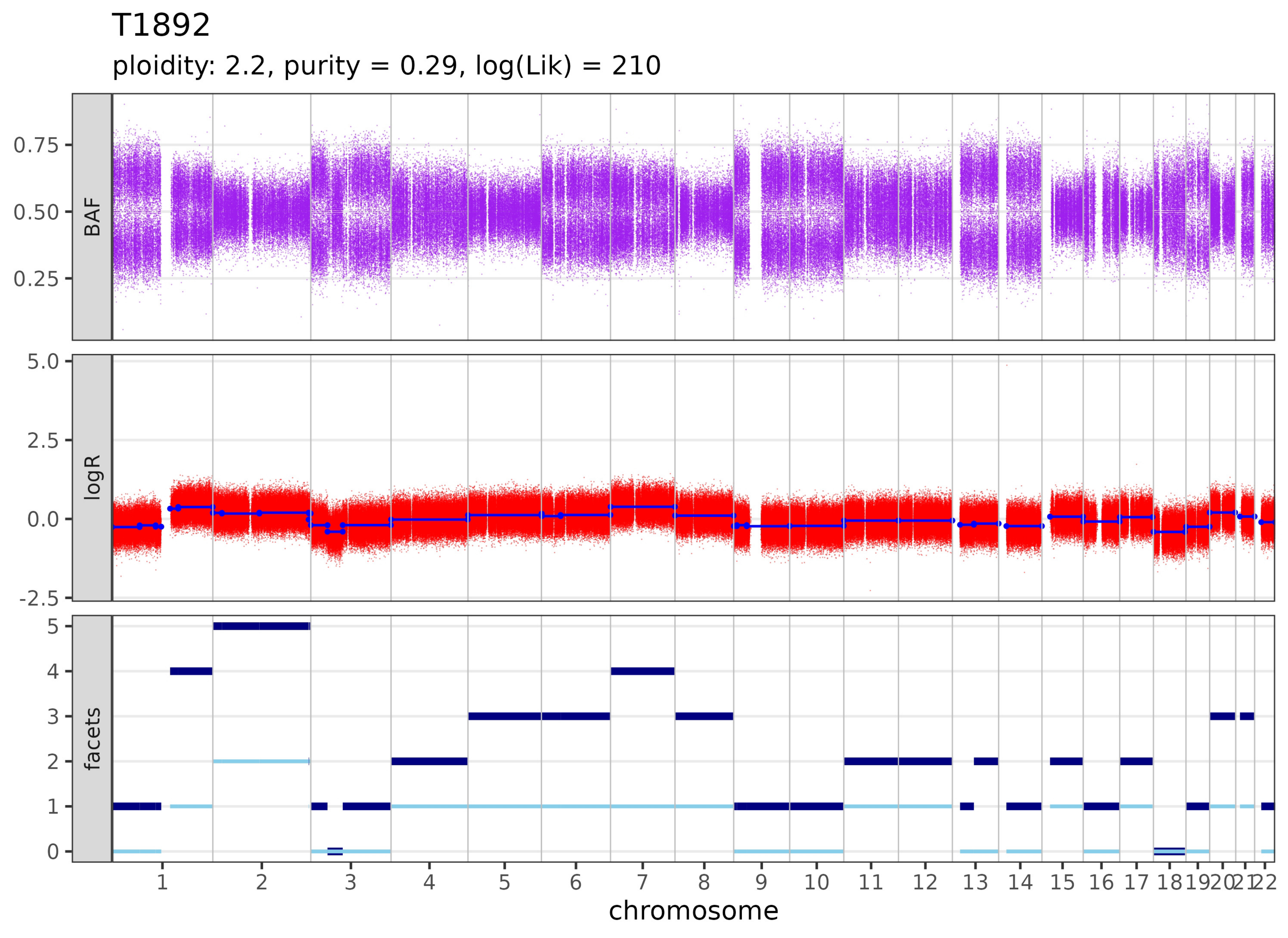
T1892
ploidity: 2.2, purity = 0.29, log(Lik) = 210
BAF
logR
facets
chromosome

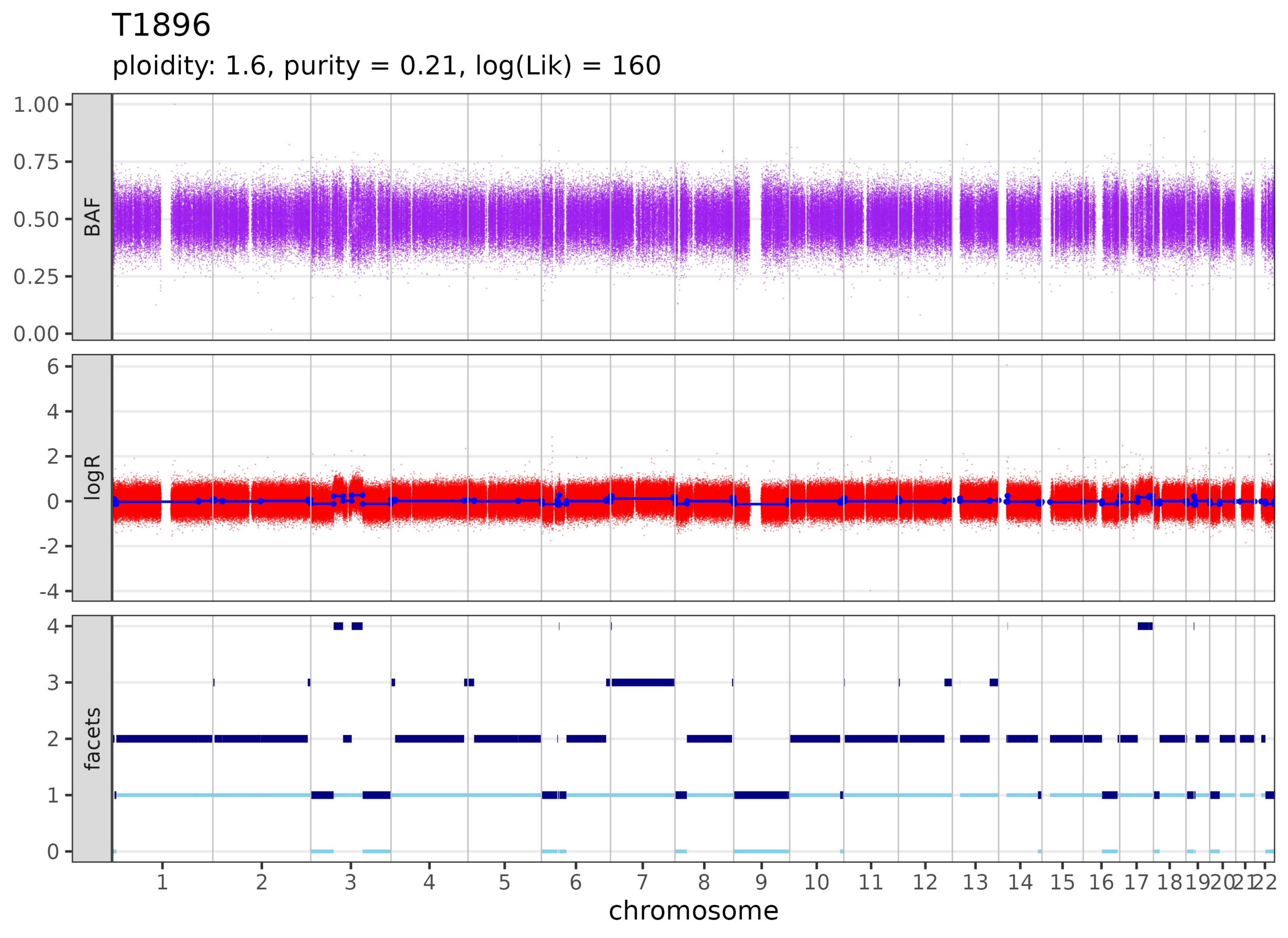
T1896
ploidity: 1.6, purity = 0.21, log(Lik) = 160
BAF
logR
facets
1.00
0.75
0.50
0.25
0.00
6
4
2
0
-2
-4
4
3
2
1
0
1 2 3 4 5 6 7 8 9 10 11 12 13 14 15 16 17 18 19 20 21 22
chromosome

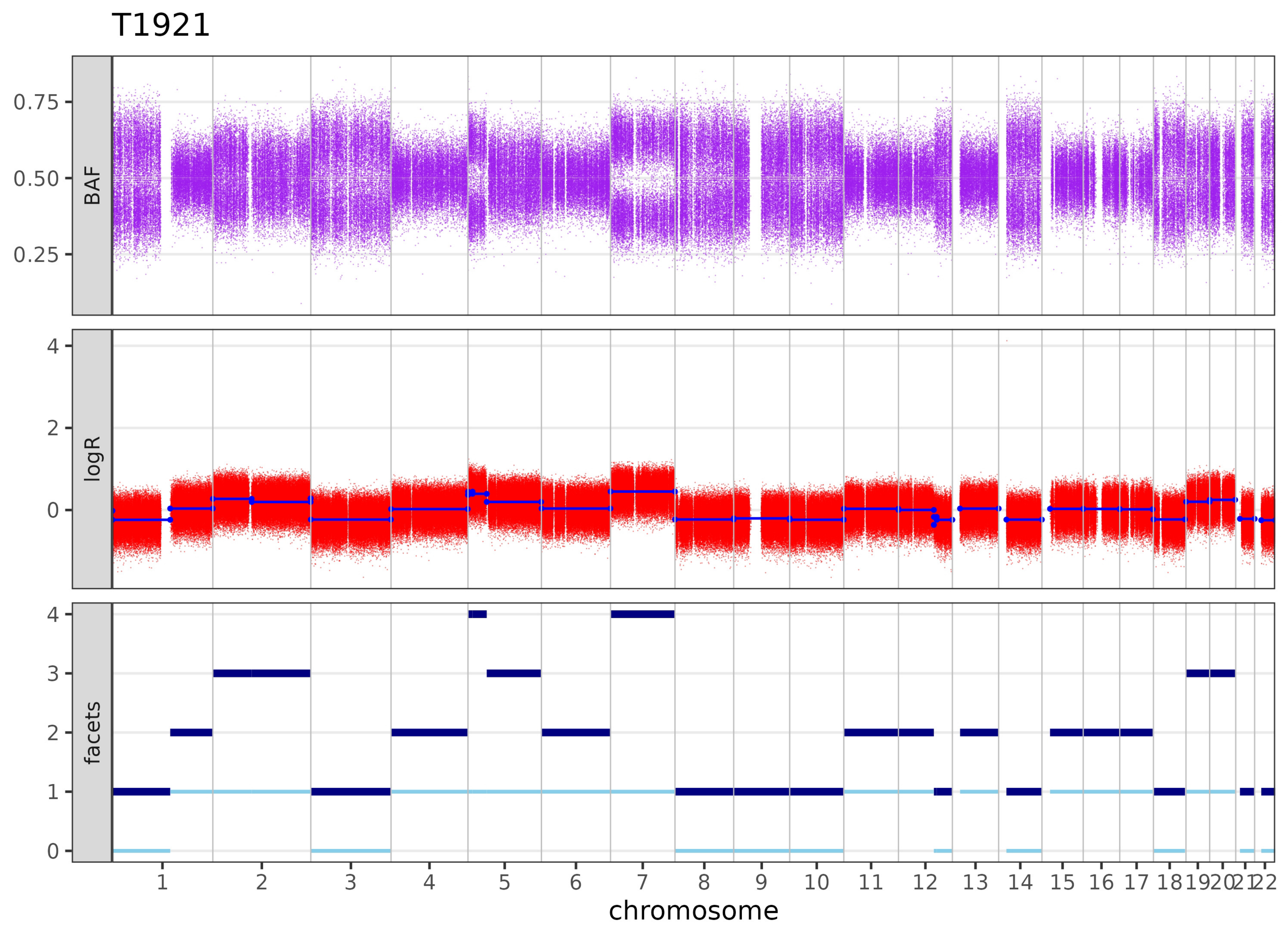

T1921
BAF
logR
facets
0.75
0.50
0.25
4
2
0
3
1
1 2 3 4 5 6 7 8 9 10 11 12 13 14 15 16 17 18 19 20 21 22
chromosome

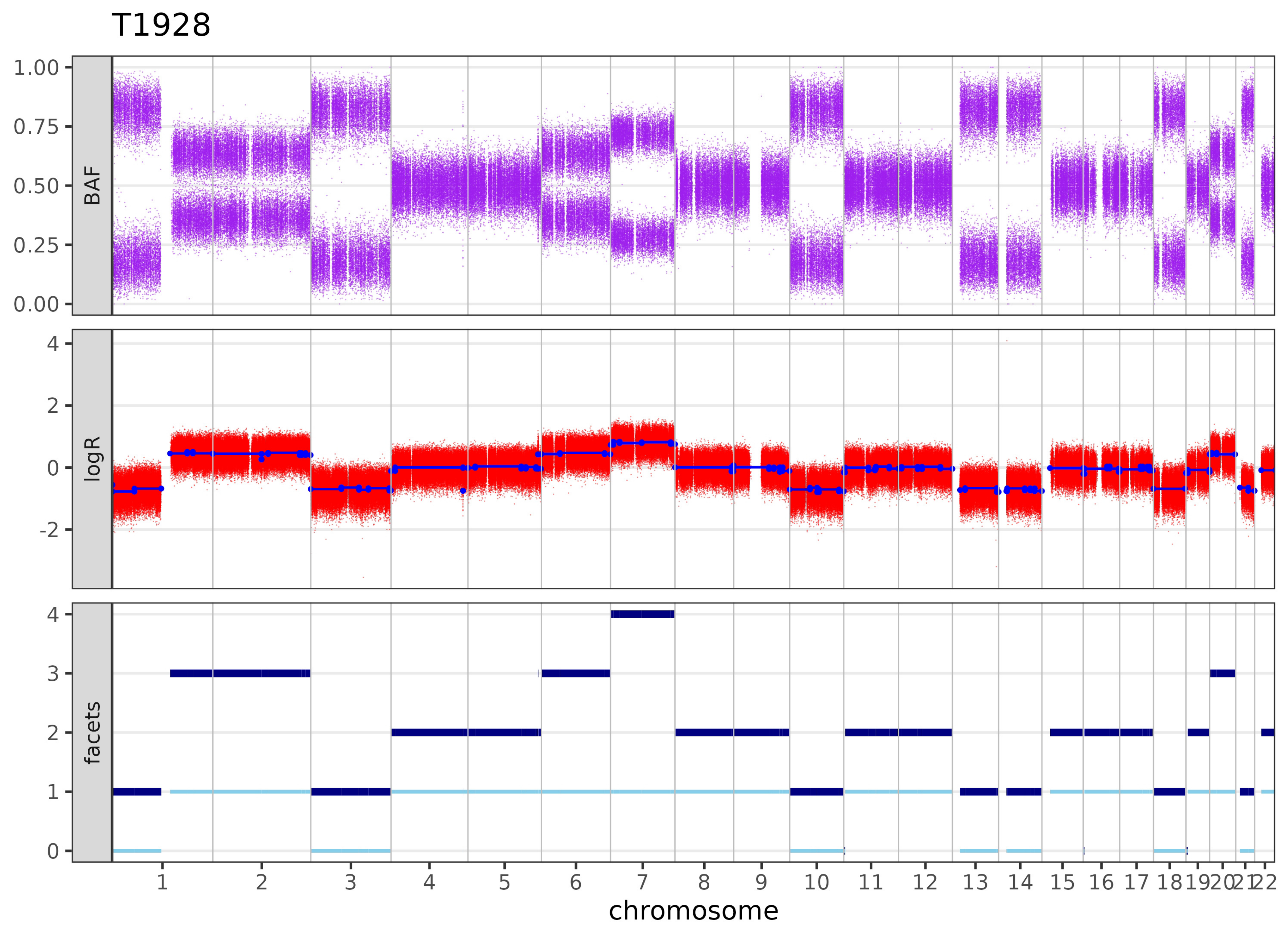

T1928
BAF
1.00
0.75
0.50
0.25
0.00
logR
4
2
0
-2
facets
4
3
2
1
0
1 2 3 4 5 6 7 8 9 10 11 12 13 14 15 16 17 18 19 20 21 22
chromosome

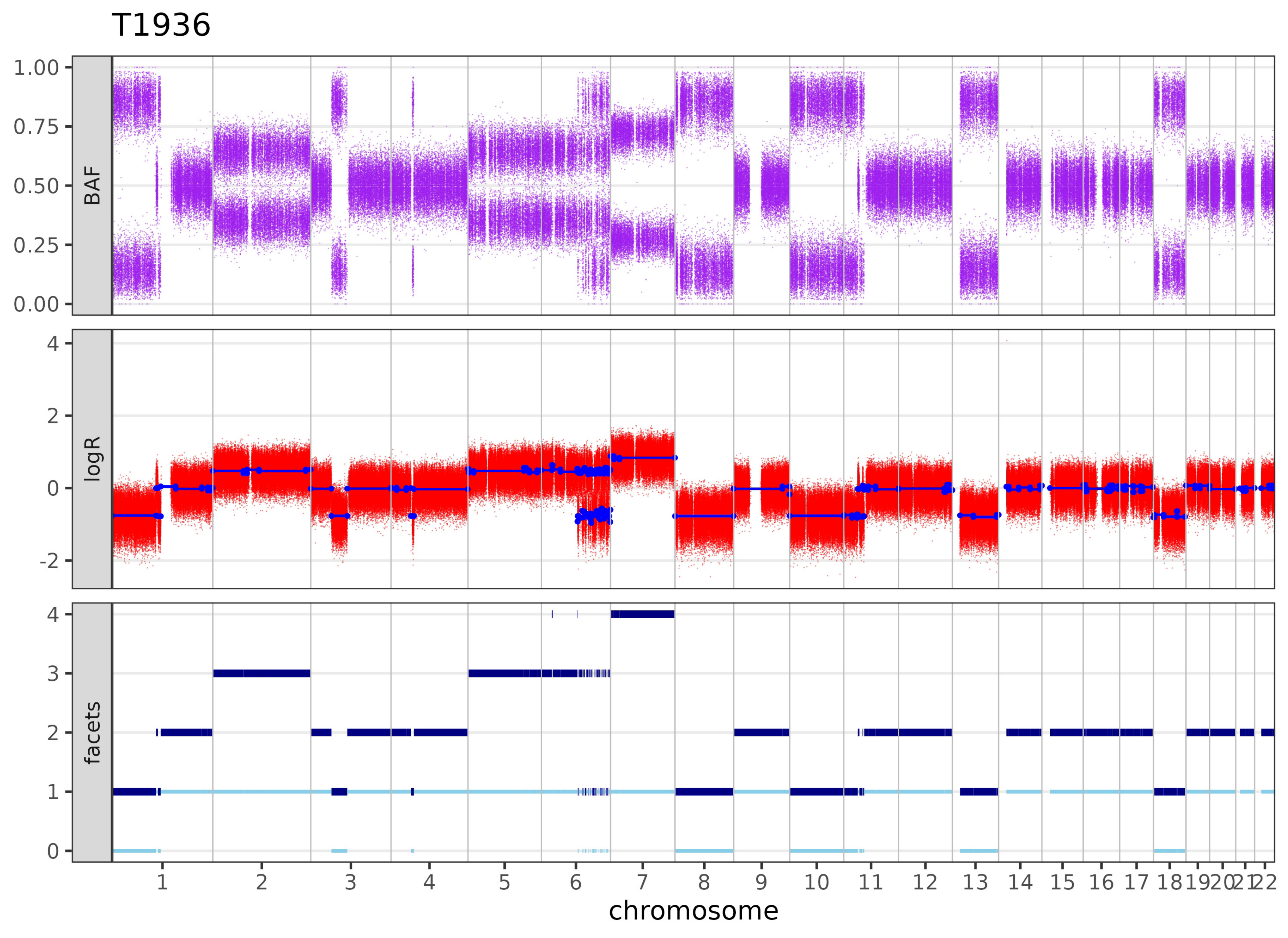

T1936
BAF
1.00
0.75
0.50
0.25
0.00
logR
4
2
0
-2
facets
4
3
2
1
0
1 2 3 4 5 6 7 8 9 10 11 12 13 14 15 16 17 18 19 20 21 22
chromosome

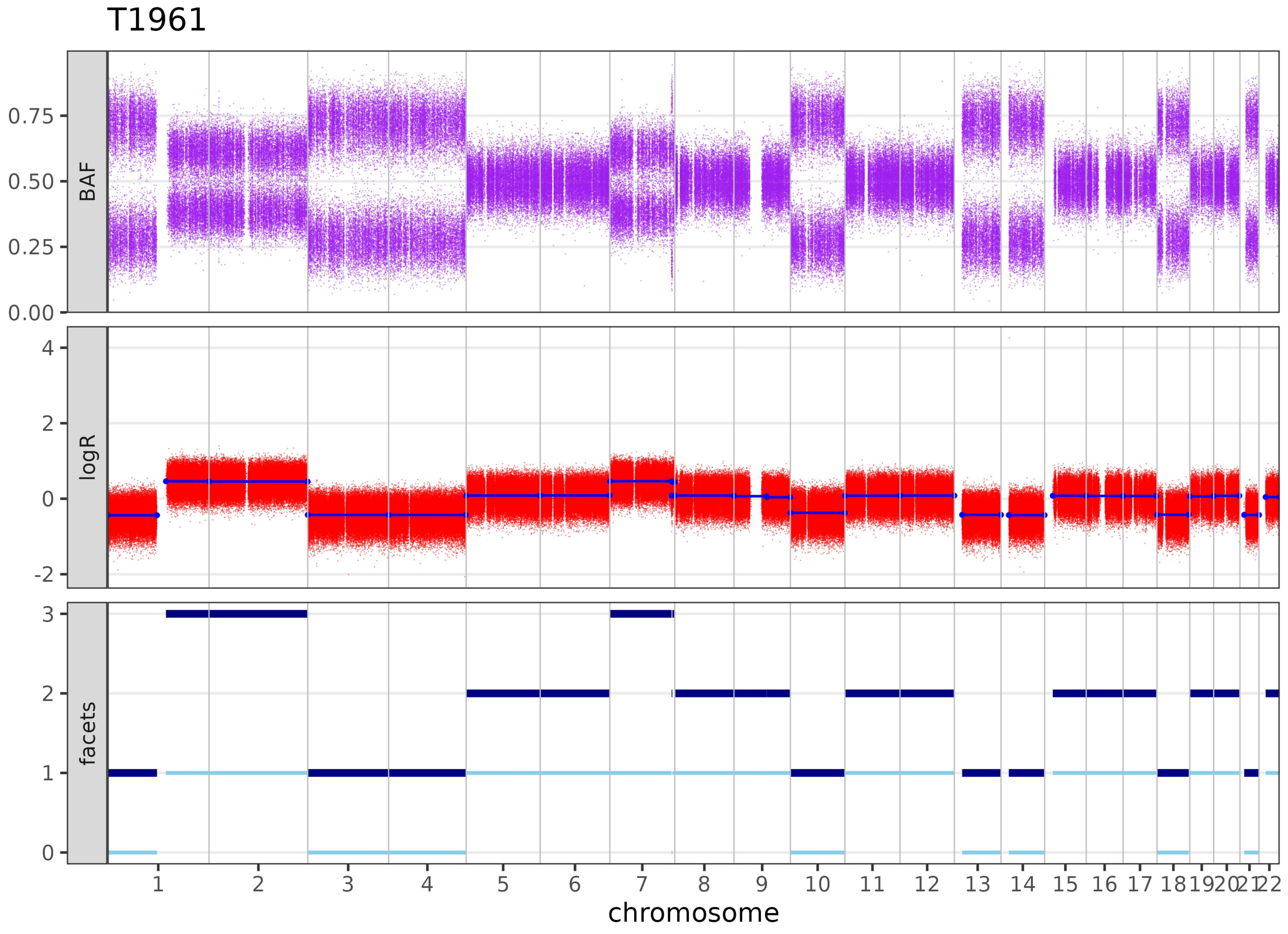

T1961
BAF
0.75
0.50
0.25
0.00
logR
4
2
0
-2
facets
3
2
1
chromosome

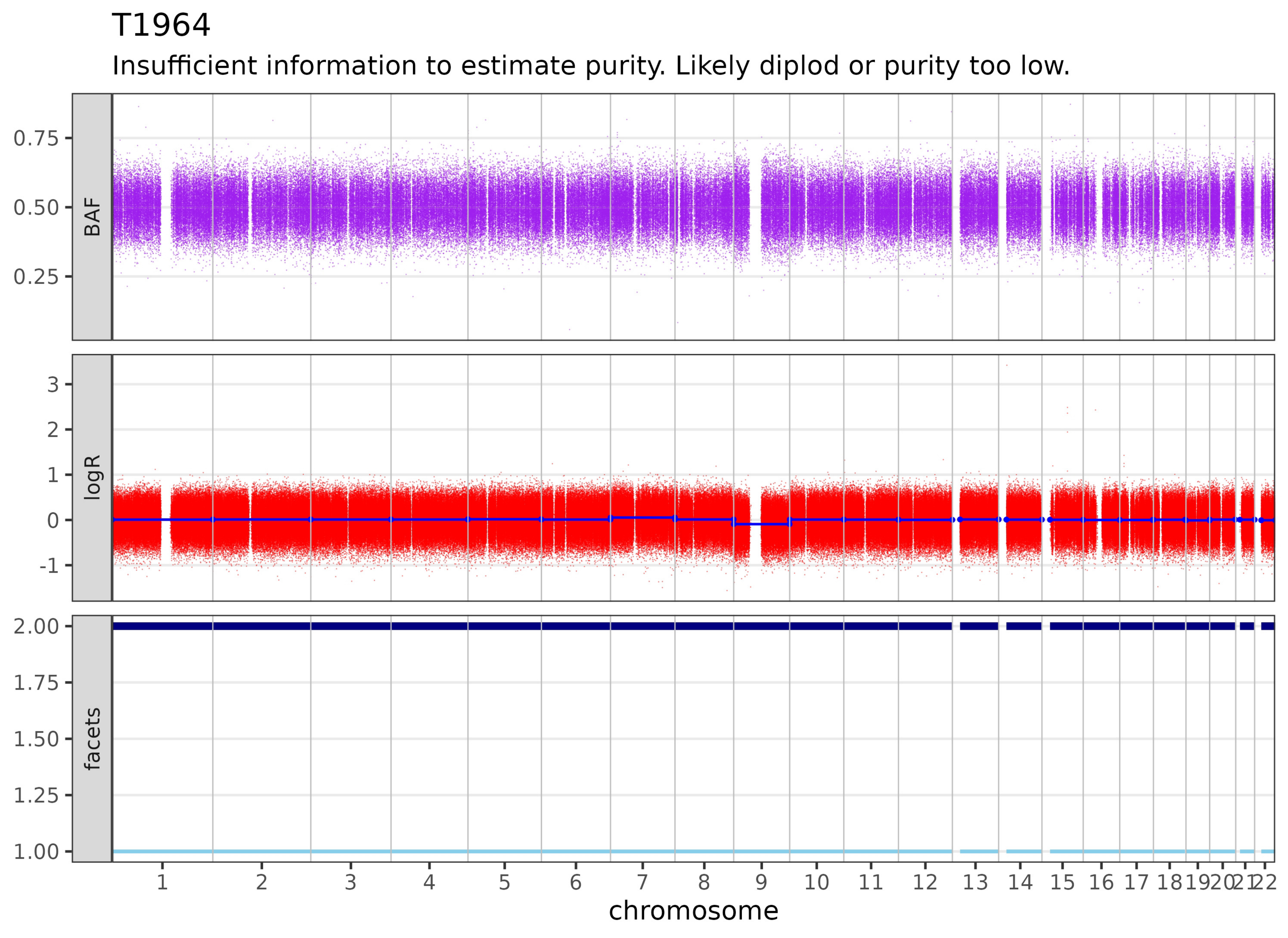

T1964
Insufficient information to estimate purity. Likely diplod or purity too low.
BAF
0.75
0.50
0.25
logR
3
2
1
0
-1
facets
2.00
1.75
1.50
1.25
1.00
chromosome

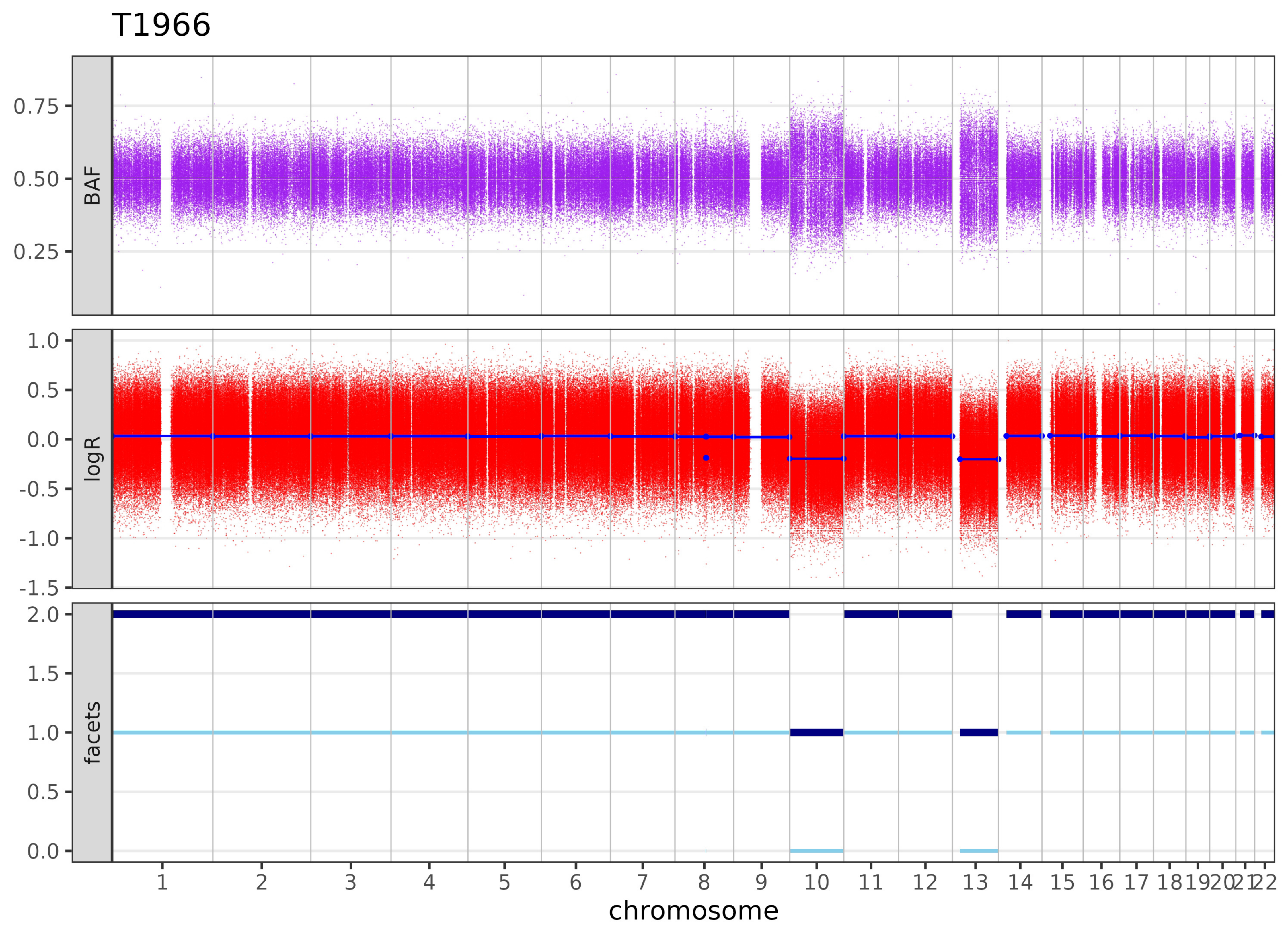
T1966
BAF
0.75
0.50
0.25
logR
1.0
0.5
0.0
-0.5
-1.0
-1.5
facets
2.0
1.5
1.0
0.5
0.0
1 2 3 4 5 6 7 8 9 10 11 12 13 14 15 16 17 18 19 20 21 22
chromosome

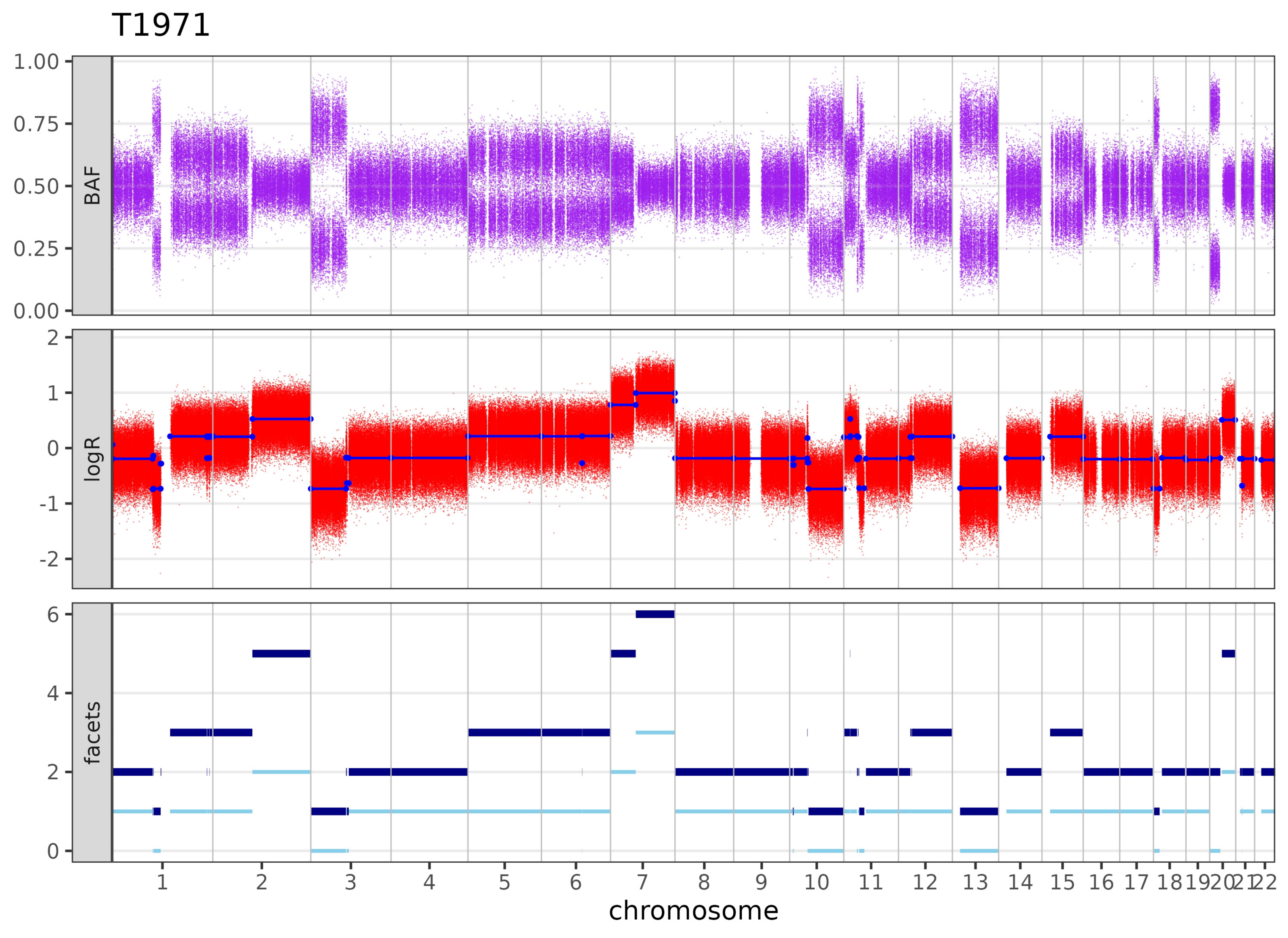

T1971
BAF
logR
facets
1.00
0.75
0.50
0.25
0.00
2
1
0
-1
-2
6
4
2
0
1 2 3 4 5 6 7 8 9 10 11 12 13 14 15 16 17 18 19 20 21 22
chromosome

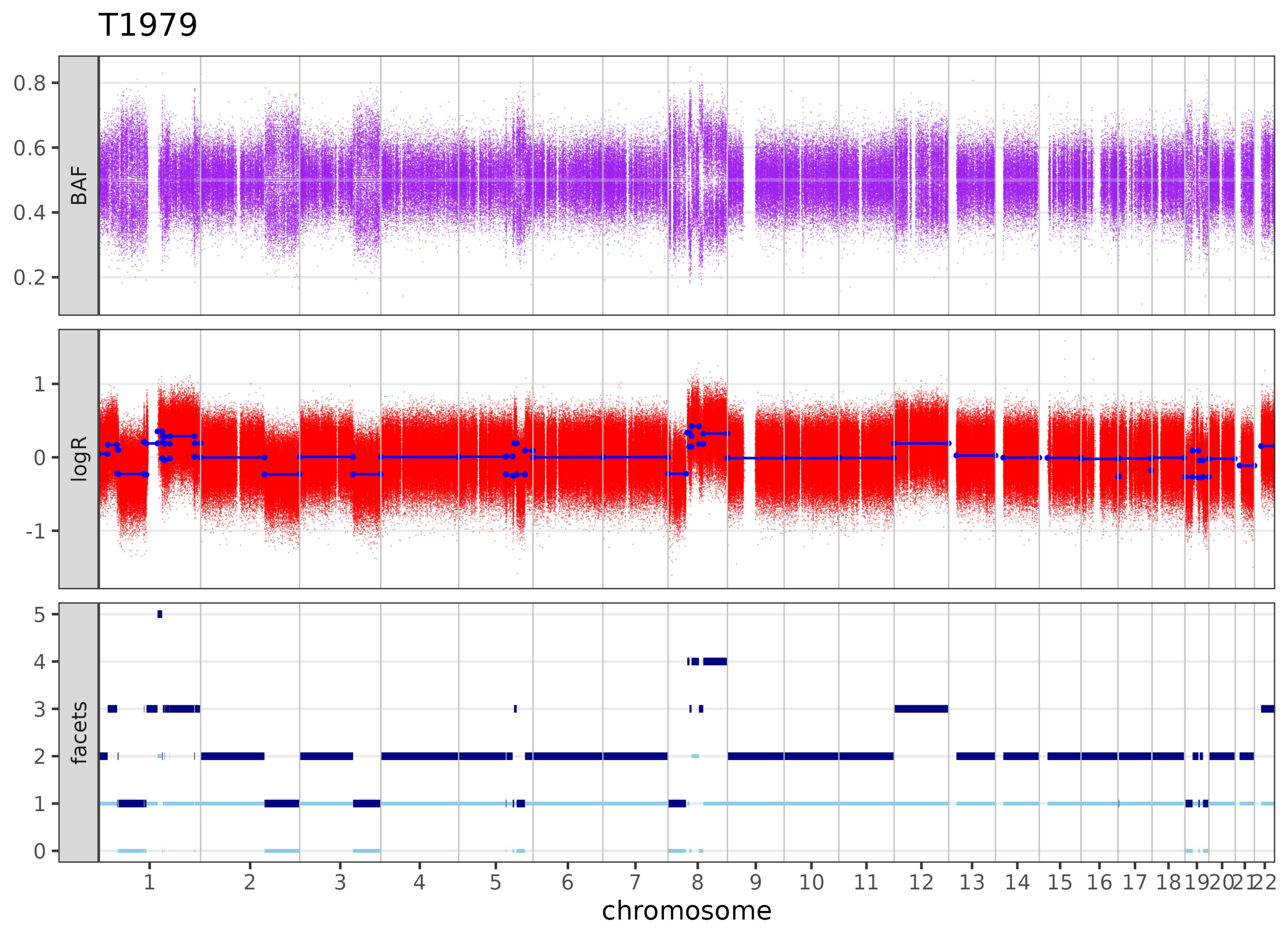
T1979
BAF
logR
facets
0.8
0.6
0.4
0.2
1
0
-1
5
4
3
2
1
0
1 2 3 4 5 6 7 8 9 10 11 12 13 14 15 16 17 18 19 20 21 22
chromosome

# T1992

Insufficient information to estimate purity. Likely diplod or purity too low.

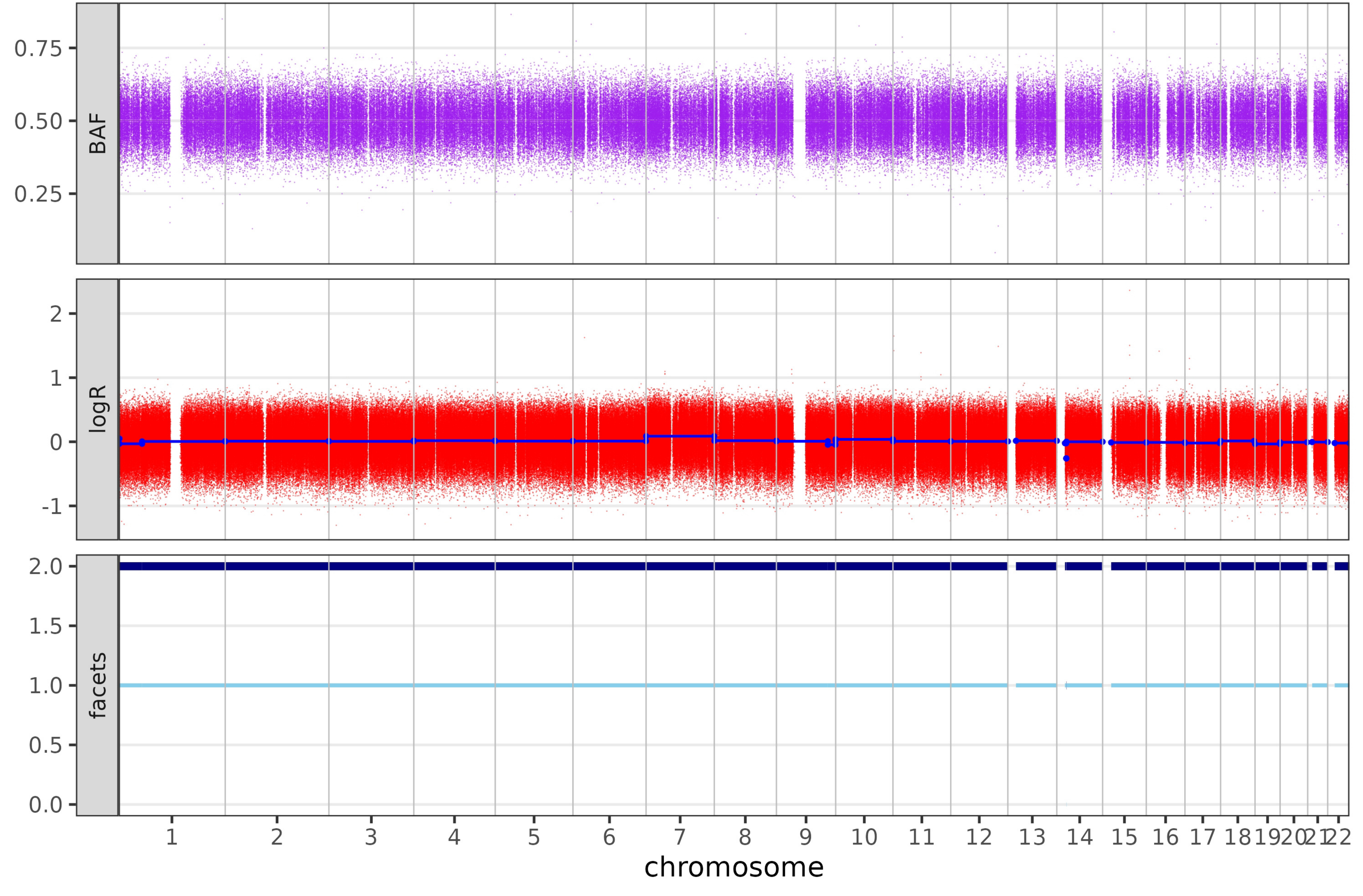

# T2011

Insufficient information to estimate purity. Likely diplod or purity too low.

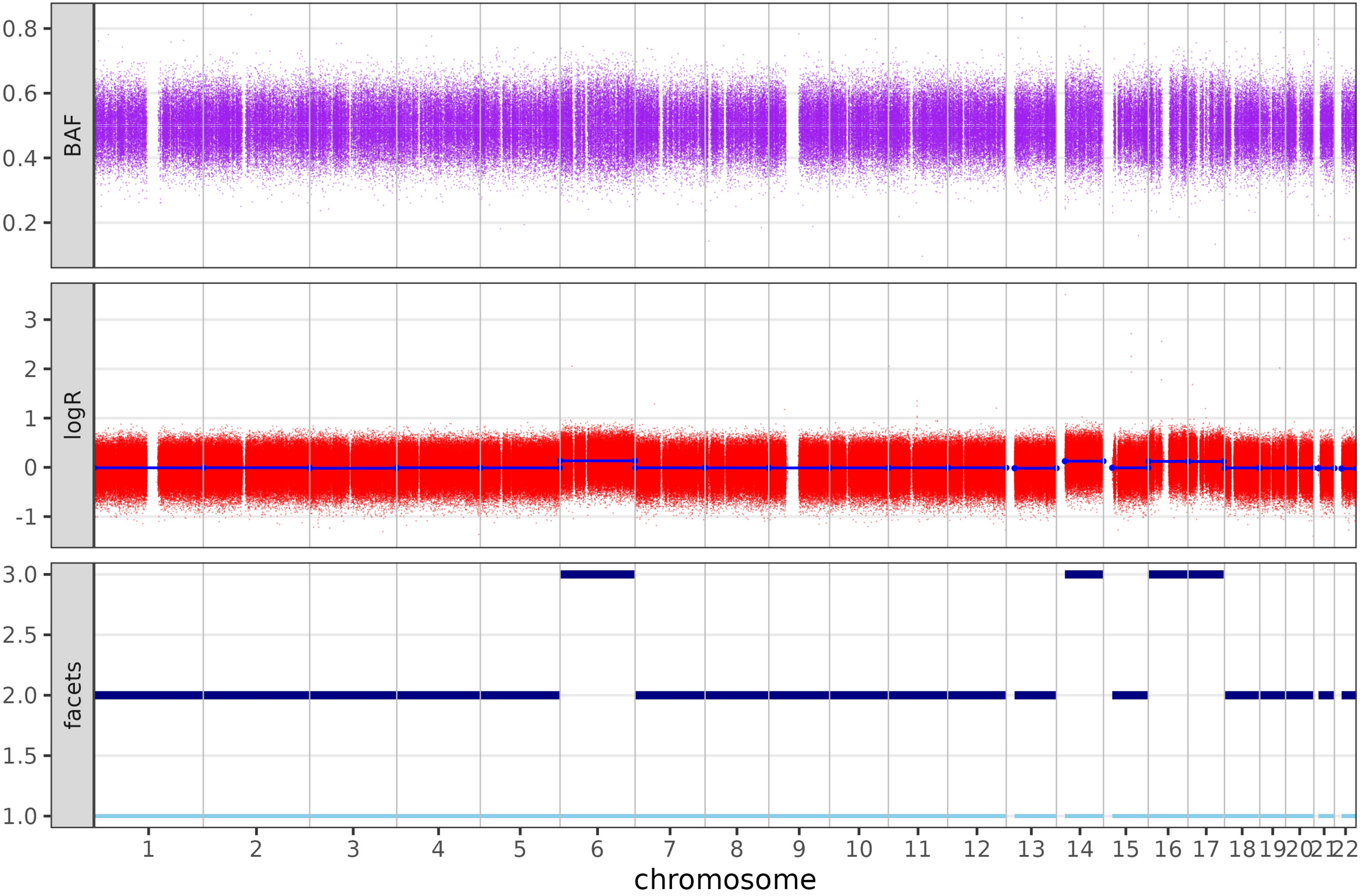

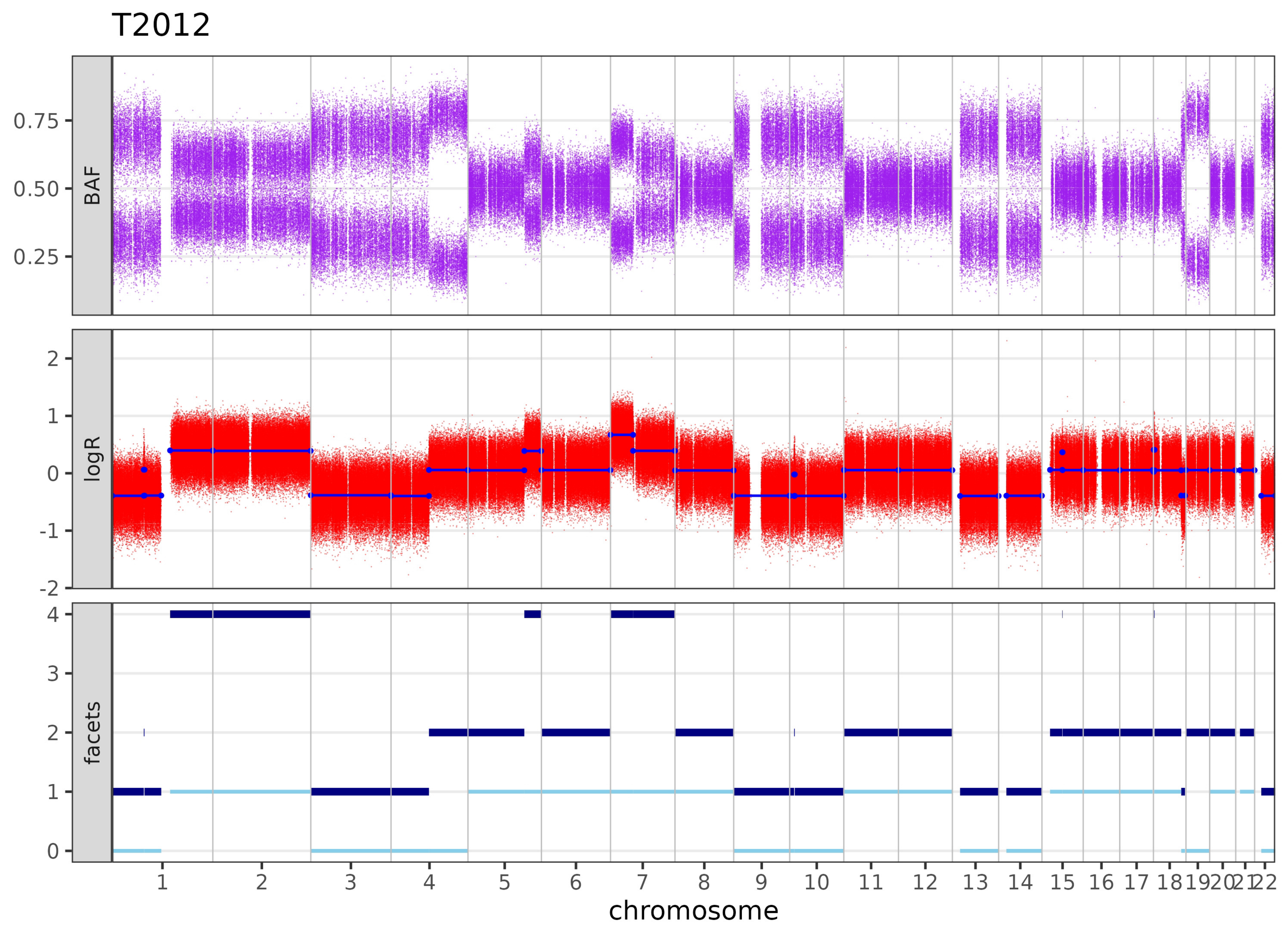

T2012
BAF
0.75
0.50
0.25
logR
2
1
0
-1
-2
facets
4
3
2
1
chromosome

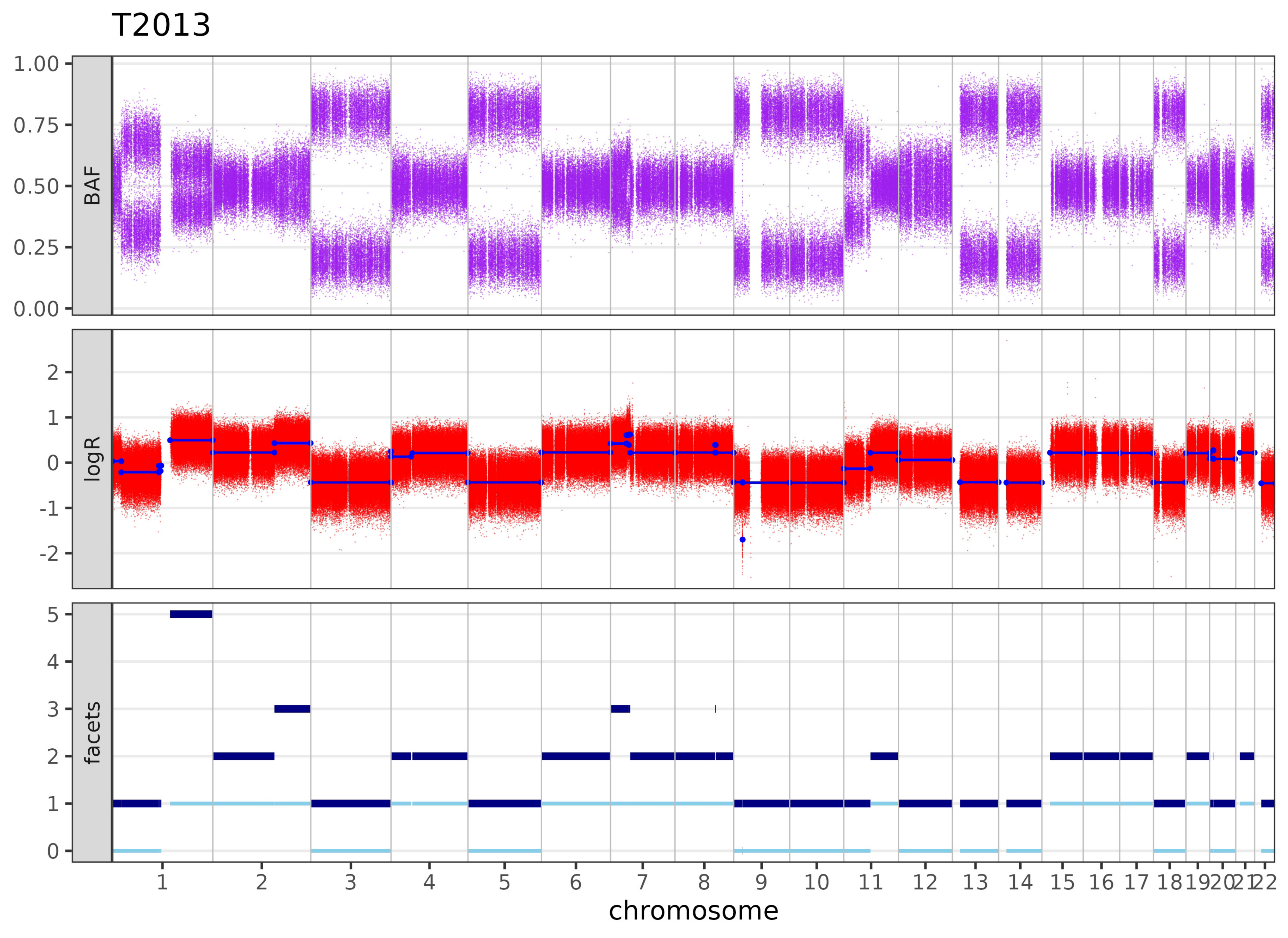
T2013
BAF
1.00
0.75
0.50
0.25
0.00
logR
2
1
0
-1
-2
facets
5
4
3
2
1
0
1 2 3 4 5 6 7 8 9 10 11 12 13 14 15 16 17 18 19 20 21 22
chromosome

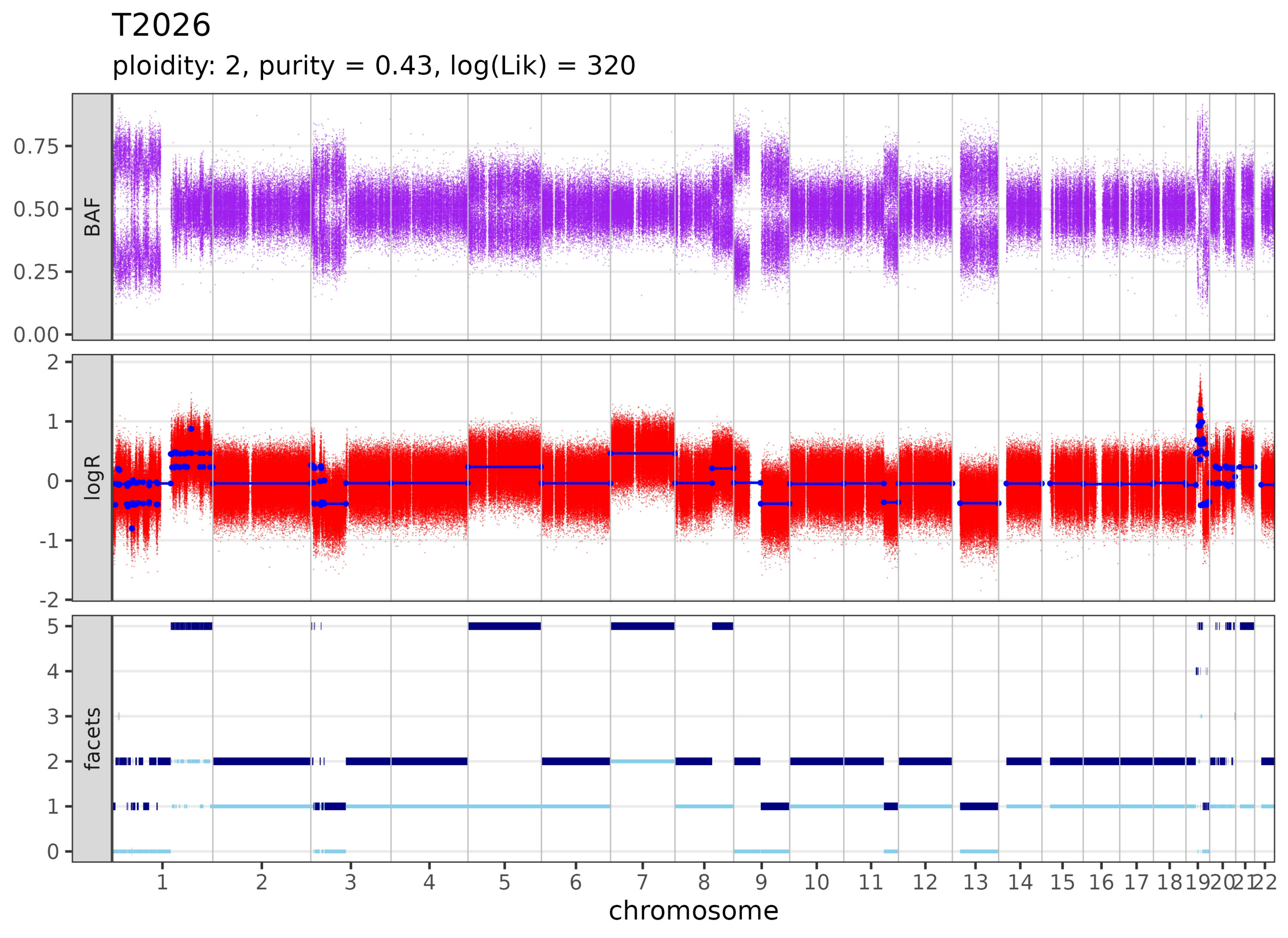

T2026
ploidity: 2, purity = 0.43, log(Lik) = 320
BAF
0.75
0.50
0.25
0.00
logR
2
1
0
-1
-2
facets
5
4
3
2
1
0
1 2 3 4 5 6 7 8 9 10 11 12 13 14 15 16 17 18 19 20 21 22
chromosome

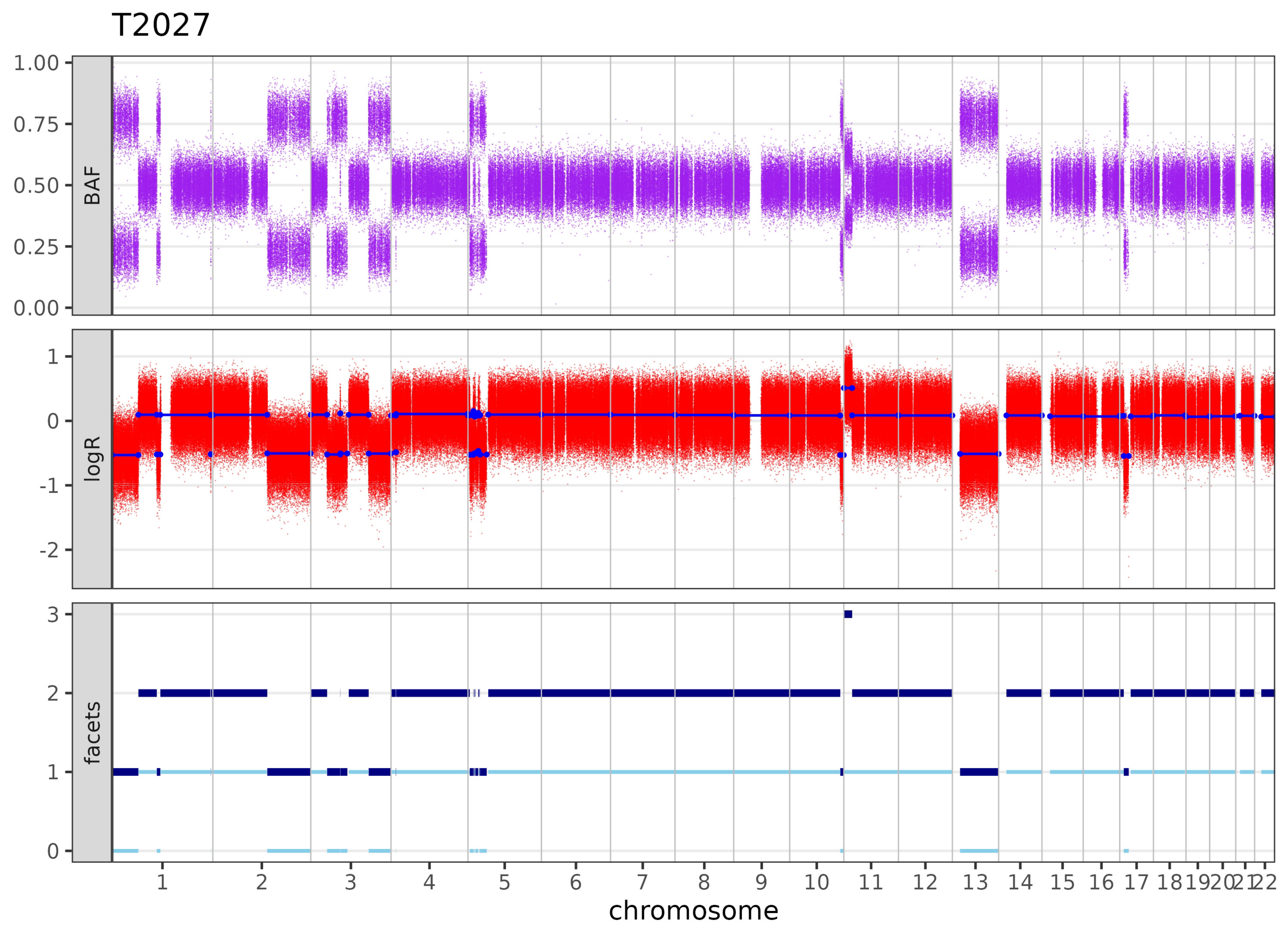

T2027
BAF
logR
facets
1.00
0.75
0.50
0.25
0.00
1
0
-1
-2
3
2
1
chromosome

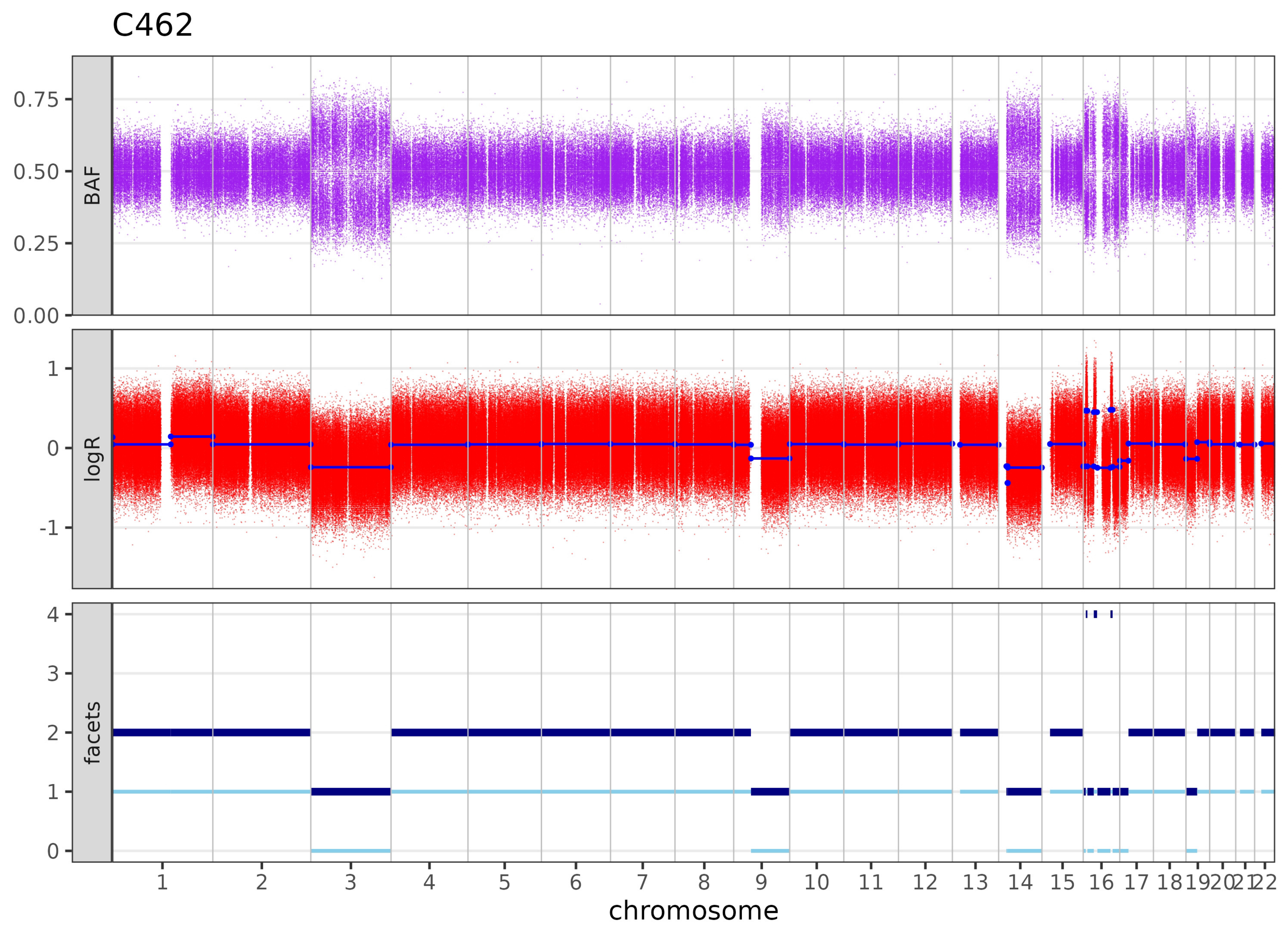
C462
BAF
0.75
0.50
0.25
0.00
logR
1
0
-1
facets
4
3
2
1
chromosome

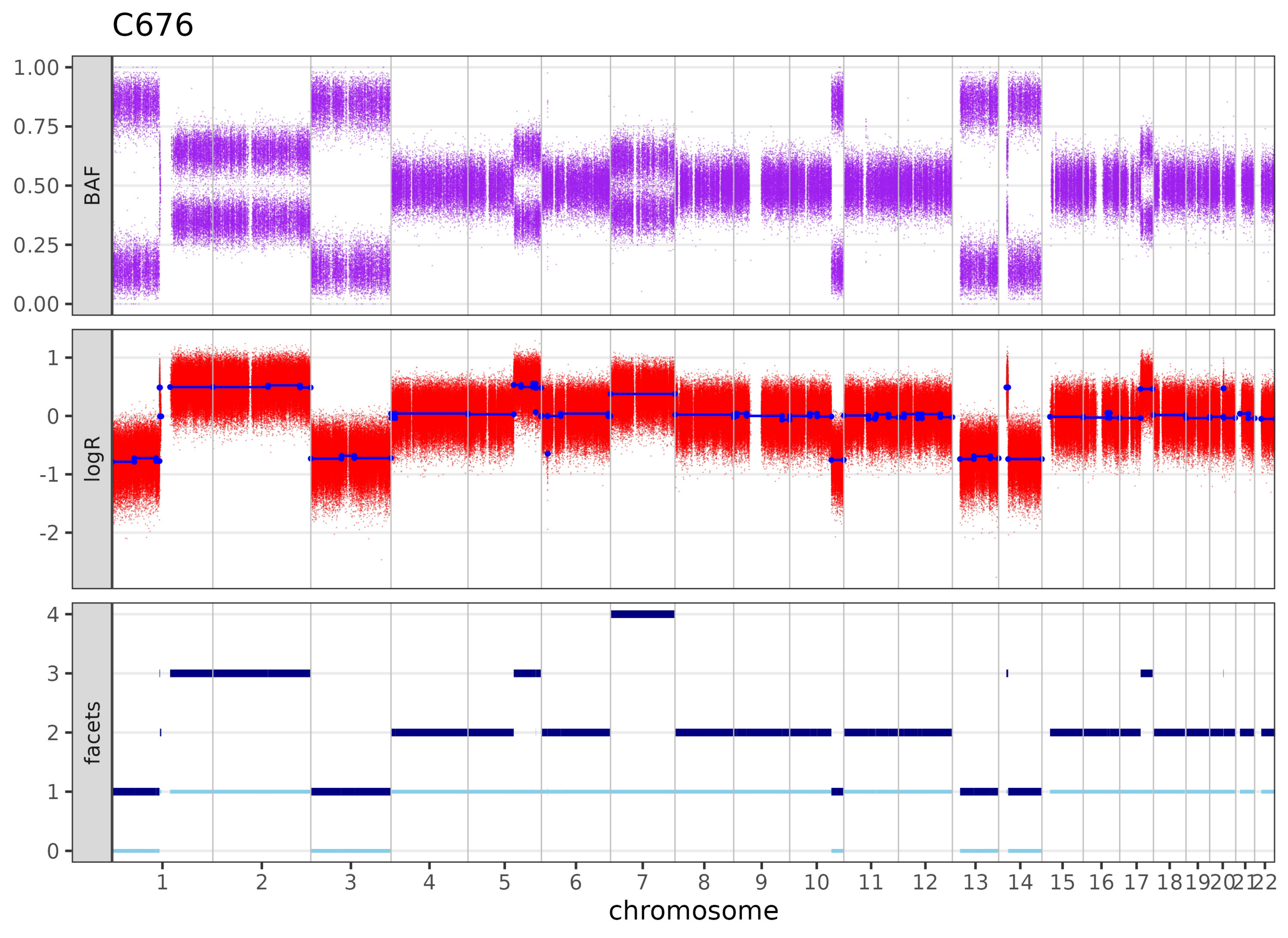

C676
BAF
1.00
0.75
0.50
0.25
0.00
logR
1
0
-1
-2
facets
4
3
2
1
0
1 2 3 4 5 6 7 8 9 10 11 12 13 14 15 16 17 18 19 20 21 22
chromosome

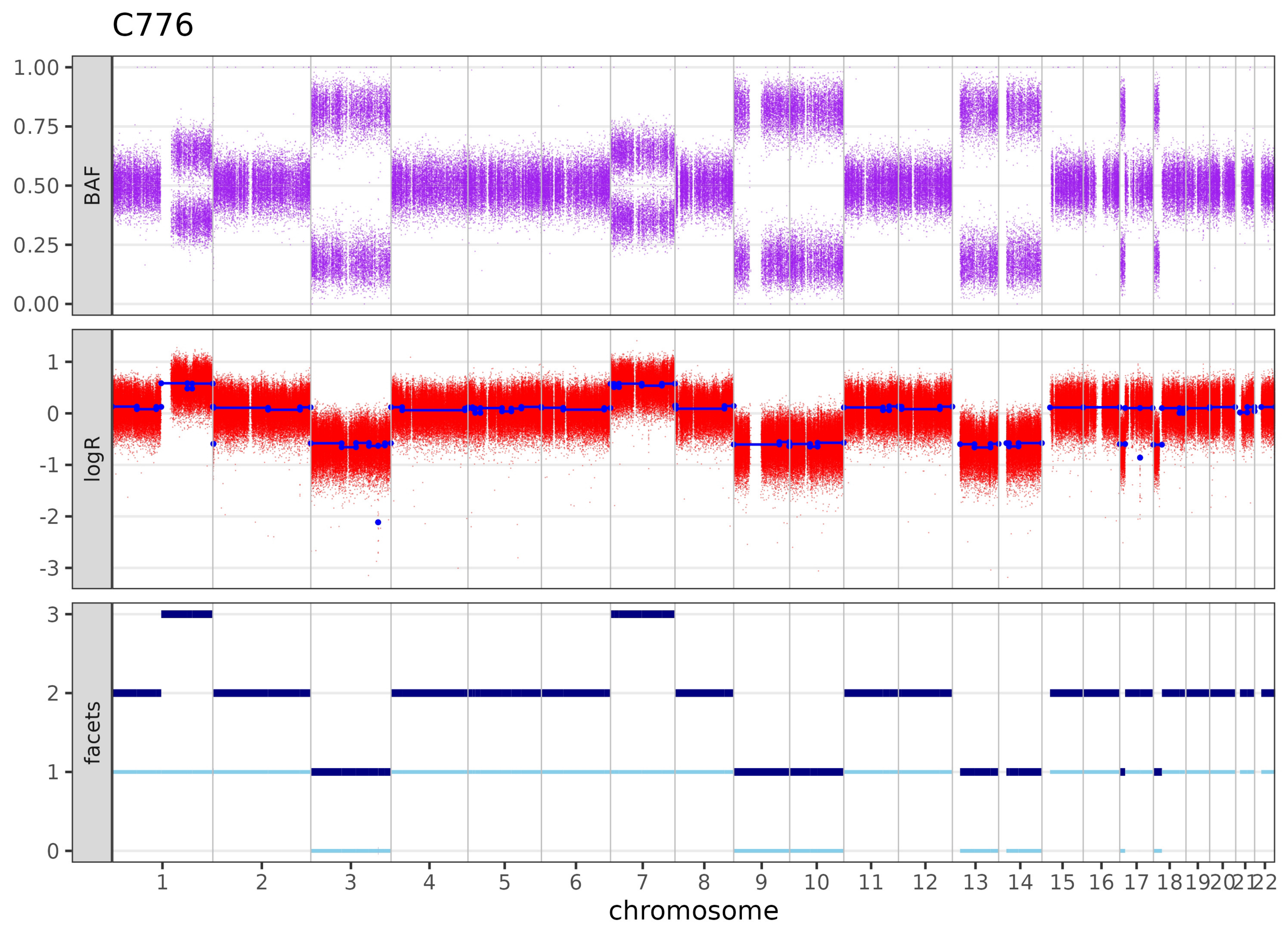
C776
BAF
logR
facets
chromosome
1.00
0.75
0.50
0.25
0.00
1
0
-1
-2
-3
3
2
1
